# Evaluation of Thiol-Mediated Uptake in VHL-Based PROTACs

**DOI:** 10.64898/2025.12.05.692502

**Authors:** Isabella Ferrara, Karl Gademann

## Abstract

Modification of biologically active molecules with 1,2-dithiolane derivatives constitutes a promising strategy to increase the cellular uptake of compounds by leveraging thiol-mediated uptake pathways. In this study, we evaluate the effect of introducing 1,2-dithiolane handles to von Hippel-Lindau (VHL)-based proteolysis targeting chimeras (PROTACs) on their cytotoxic properties. Starting from a previous molecular design, two sets of derivatives with 1,2-dithiolane handles and control compounds comprising the corresponding carbon-equivalents were synthesized and studied for their anticancer properties. Increased or equipotent cytotoxicity towards cancer cells (HeLa) was observed for several derivatives compared to the parent compound. Especially for lipoic acid derivatives, and the corresponding all-carbon derivatives, a significant increase in cytotoxicity compared to the unfunctionalized compound was observed. These results suggest effects other than or in addition to thiol-mediated uptake for acylated PROTAC derivatives.

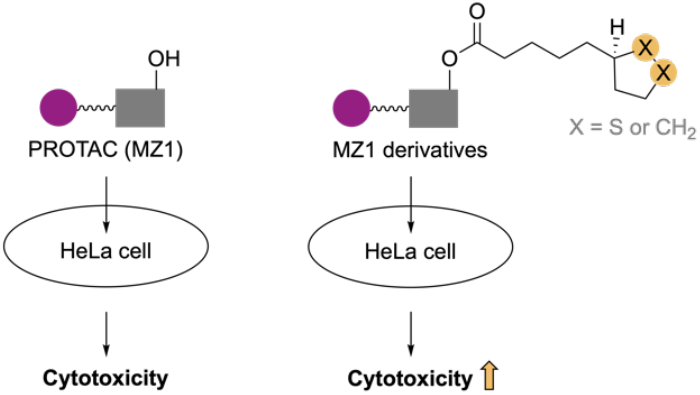

## Introduction

Thiol-mediated uptake (TMU) has emerged as a promising tool to increase cellular uptake into mammalian cells.^1–11^ In this context, modification of compounds with residues comprising cyclic disulfide moieties, such as 1,2-dithiolanes, was shown to be successful to increase uptake (Fig. 1A-B).^1–7,9,10,12^ The underlying principle of TMU leveraging 1,2-dithiolane handles was suggested to be the reactivity of the disulfide functionality with cellular exofacial thiols that promote covalent bond formation between the 1,2-dithiolane functionalized cargo molecule and the cell surface protein (Fig. 1A).^1–7,9,10^ With the nucleofuge remaining connected to the cargo in the case of 1,2-dithiolane moieties, covalent exchange-cascades are feasible.^1,4–6,9^ Following covalent bond formation, uptake of the cargo can be promoted for example by endocytosis-dependent mechanisms, or direct translocation among other mechanisms.^1,2,5,6,8–10^ However, the detailed mechanisms of TMU remain not yet fully understood and might likely vary depending on the system under investigation (*e.g*. substrate-dependent, cell-line-dependent, *etc*.).^1,2,5,8^

**Figure 1.**
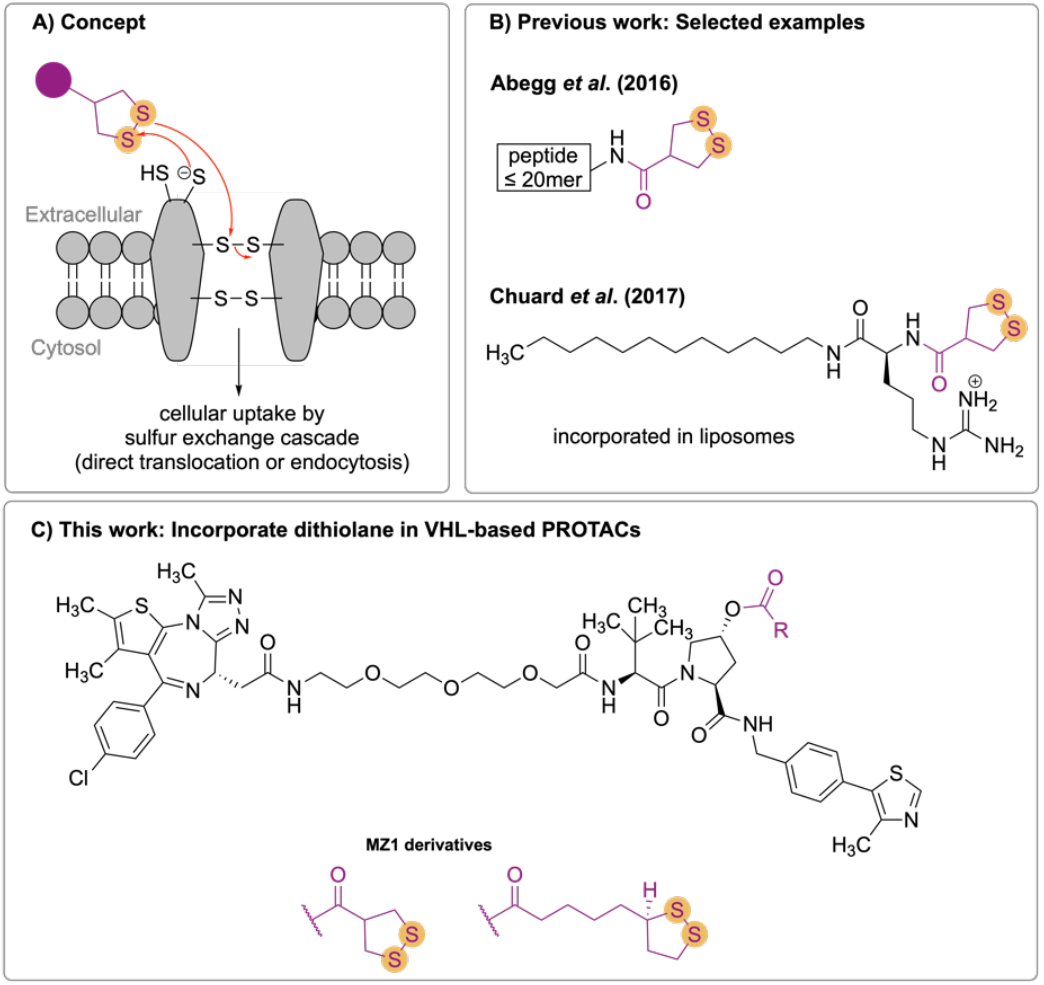
Thiol-mediated uptake leveraging 1,2-dithiolane handles: Concept (A),^1^ selected literature examples (B),^2,3^ and this work (C).

In this study, the applicability of TMU to the therapeutic field of protein-targeting chimeras (PROTACs) was evaluated. PROTACs are bifunctional entities that leverage the cellular ubiquitin-proteasome-system for degradation of the protein of interest (POI) by binding to the POI and an E3-ligase.^13–19^ By bringing the POI and E3-ligase in proximity, the POI is ubiquitylated and thereby tagged for degradation by the proteasome. PROTACs have emerged as a promising strategy to tackle malfunctioning or overexpressed proteins associated with causing diseases and various examples have reached clinical trials.^16–20^ One limitation of PROTACs is the often limited membrane permeability.^14,17,20–24^ Therefore, we envisaged that modification of PROTACs with 1,2-dithiolane handles could potentially allow to leverage TMU as a strategy to increase the cellular uptake of PROTACs into mammalian cells. We hypothesized that increased cellular uptake resulting in higher intracellular concentrations might result in improved anticancer potency. In support of the assumption that higher intracellular concentrations could increase potency, the Ciulli group recently reported an increased potency of the benchmark PROTAC MZ1 (**1**) when cells were pre-treated with an efflux inhibitor, and a significant shift of the potency curves of MZ1 in NanoBRET target engagement assays comparing permeabilized and live-cell modes.^25^

The literature-known von Hippel-Lindau (VHL)-based PROTACs MZ1 (**1**) and MZ3 (**2**)^26^ were selected as base designs for our study. The hydroxyproline moiety (Hyp-OH) within VHL-based PROTACs was reasoned as a suitable handle to introduce the 1,2-dithiolane moieties *via* an acyl linkage (Fig. 1C). However, the Hyp-OH moiety is required for binding of VHL-based PROTACs to VHL^15,17,20,26^ and thus, we anticipated that the active PROTAC after cellular uptake and hydrolysis of the ester moiety would be the unmodified parent compounds MZ1 and MZ3. In support of this hypothesis, the Hyp-OH functionality of VHL-based PROTACs has been successfully leveraged previously as a handle for modification *via* an acyl linkage to allow for targeted drug delivery for example in antibody-PROTAC conjugates^27,28^ and PROTAC-folate conjugates^29^.

We synthesized the asparagusic acid (AspA), and lipoic acid (LipA) derivatives of MZ1 and MZ3. In addition, the carbon-equivalents (All-C) of the AspA and LipA conjugates were prepared as control compounds that do not constitute substrates for thiol-mediated uptake. The cytotoxic properties of the 1,2-dithiolane derivatives, and the All-C analogs toward HeLa cells was tested by 3-(4,5-dimethylthiazol-2-yl)-2,5-diphenyltetrazolium bromide (MTT) assays.

## Results and Discussion

### Synthesis of the common intermediate (*S,R,S*)-AHPC.HCl salt

We started out with the preparation of the literature known compounds MZ1 (**1**) and MZ3 (**2**) *via* preparation of the common intermediate (*S,R,S*)-AHPC hydrochloride (**3**) following literature procedures from Steinebach *et al*.^30^ and Yan *et al*.^31^ with minor modifications as depicted in Fig. 2. Starting from Boc-protected 4-bromo-benzyl amine (**4**) and 4-methyl thiazole (**5**), a Pd-catalyzed coupling afforded compound **6** in 71% yield. Subsequent Boc-deprotection under acidic conditions and amide coupling of the resulting amine **7** with Boc-protected Hyp-OH yielded Boc-protected compound **8**. Acid-mediated Boc-deprotection and coupling of amine **9** with Boc-protected *tert*-leucine, followed by Boc deprotection of compound **10** provided the targeted VHL-binding ligand **3**.

**Figure 2.**
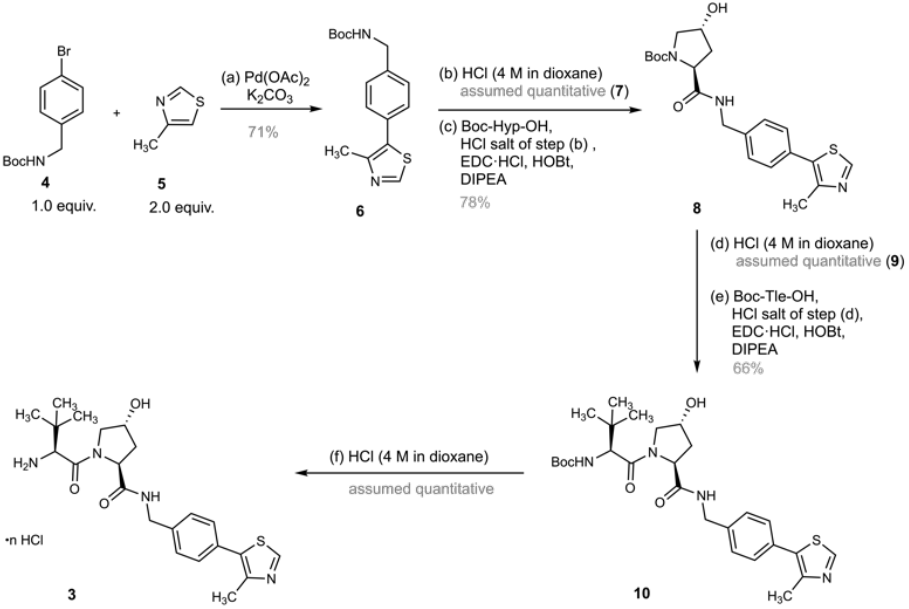
Preparation of the common intermediate (S,R,S)-AHPC.HCl salt (**3**): Reagents and conditions: (a) bromide **4** (1.0 equiv.), thiazole **5** (2.0 equiv.), Pd(OAc)_2_ (1 mol%), K_2_CO_3_ (2.0 equiv.), DMA (0.3 M), 130 °C, 5 h; (b) HCl (4 M in dioxane, 4.0 equiv.), CH_2_Cl_2_ (0.2 M), 25 °C, 24 h; (c) Boc-Hyp-OH (1.0 equiv.), HCl salt of step (b) (1.1 equiv.), EDC.HCl (1.3 equiv.), HOBt (1.3 equiv.), DIPEA (3.0 equiv.), CH_2_Cl_2_ (0.1 M), 0 to 25 °C, 18 h; (d) HCl (4 M in dioxane, 4.0 equiv.), CH_2_Cl_2_ (0.2 M), 25 °C, 5.5 h; (e) Boc-Tle-OH (1.0 equiv.), HCl salt of step (d) (1.1 equiv.), EDC.HCl (1.3 equiv.), HOBt (1.3 equiv.), DIPEA (3.0 equiv.), CH_2_Cl_2_ (0.1 M), 0 to 25 °C, 18 h; (f) HCl (4 M in dioxane, 20.0 equiv.), CH_2_Cl_2_ (0.2 M), 0 to 25 °C, 3.5–5 h.

### Synthesis of MZ1 and MZ1 derivatives

We continued with the 1-ethyl-3-(3-dimethylaminopropyl)carbodiimide (EDC)/1- hydroxybenzotriazole (HOBt) coupling procedure to couple the common intermediate **3** to the Boc protected linker affording compound **11** (Fig. 3). Subsequent Boc deprotection under acidic conditions, followed by coupling to the carboxylic acid derivative of JQ-1 afforded the compound MZ1 (**1**). The Hyp-OH moiety in MZ1 was esterified with the carboxylic acids of the targeted dithiolanes and the corresponding carbon-analogs adapting similar conditions as reported by Liu *et al*.^29^ for the acylation of the Hyp-OH during their preparation of PROTAC-Folate conjugates. Following the coupling reaction, the mixtures were filtered over DSC-18 SPE cartridges and purified by preparative reversed-phase HPLC to provide the dithiolane containing MZ1 conjugates MZ1-AspA **13** and MZ1-LipA **15**, and the corresponding All-C analogs MZ1-All-C-AspA **14** and MZ1-All-C-LipA **16**.

**Figure 3.**
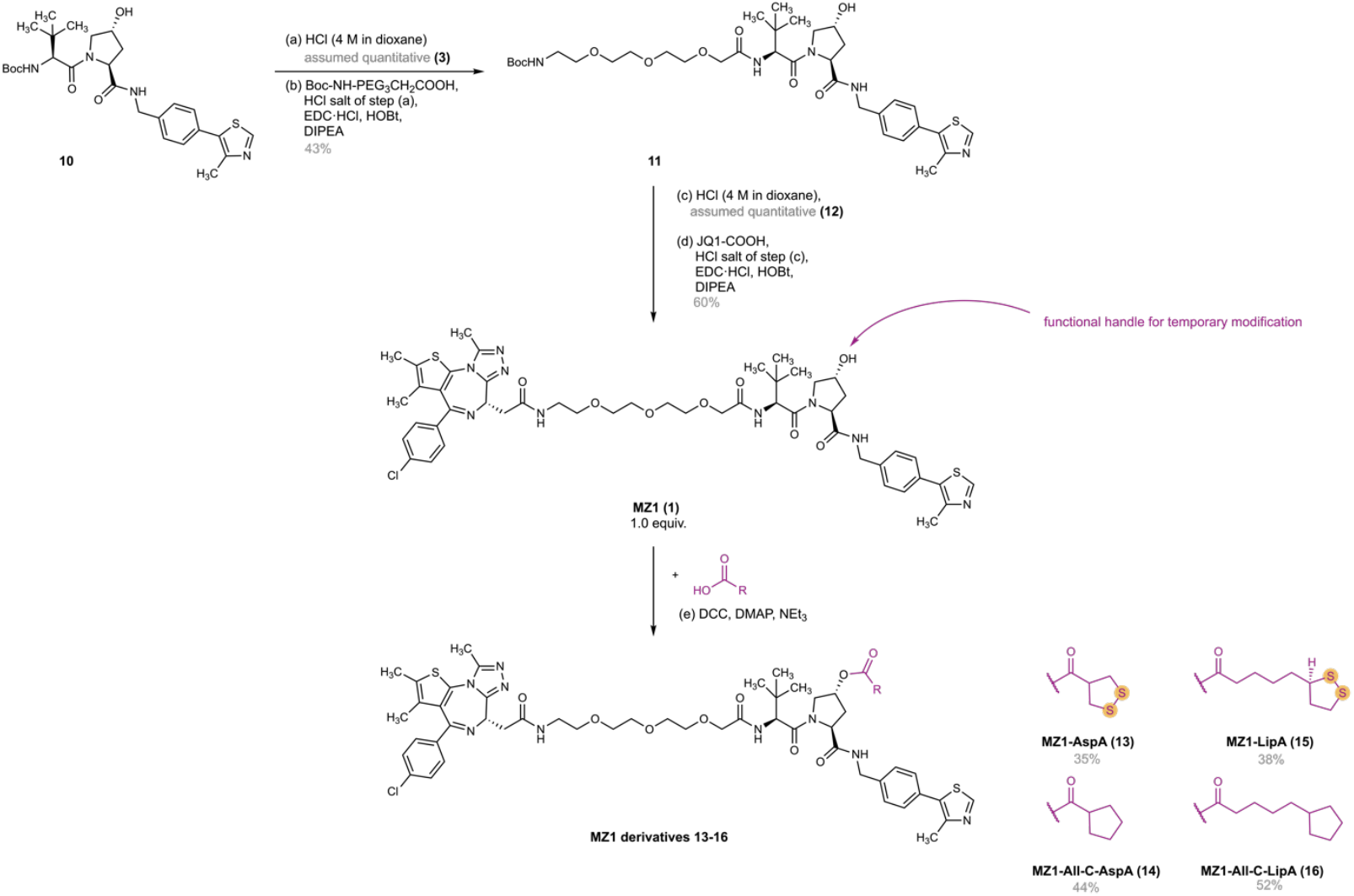
Synthesis of MZ1 and MZ1 derivatives **13-16.** Reagents and conditions: (a) HCl (4 M in dioxane, 20.0 equiv.), CH_2_Cl_2_ (0.2 M), 0 to 25 °C, 3.5 h; (b) Boc-NH-PEG_3_CH_2_COOH (1.0 equiv.), HCl salt of step (a) (1.1 equiv.), EDC.HCl (1.3 equiv.),HOBt (1.3 equiv.), DIPEA (3.0 equiv.), CH_2_Cl_2_ (0.1 M), 0 to 25 °C, 44 h; (c) HCl (4 M in dioxane, 4.0 equiv.), CH_2_Cl_2_ (0.2 M), 25 °C, 18 h; (d) JQ1-COOH (1.0 equiv.), HCl salt of step (c) (1.1 equiv.), EDC.HCl (2.6 equiv.), HOBt (2.6 equiv.), DIPEA (7.0 equiv.), CH_2_Cl_2_ (0.1 M), 0 to 25 °C, 27 h; (e) RCOOH (2.0–3.0 equiv.), DCC (1.5–3.2 equiv.), DMAP (0.1–0.2 equiv.), NEt_3_ (1.0-7.0 equiv.), CH_2_Cl_2_ (0.01 M), 25 °C, 41–73 h.

### Synthesis of MZ3 and MZ3 derivatives

The next target MZ3 (**2**) with an additional phenylalanine moiety was prepared *via* sequential Boc-deprotection, EDC/HOBt-mediated amide coupling reactions (see Fig. 4). The common VHL-binding ligand **3** was first coupled with Boc-protected phenylalanine to provide compound **17**, followed by Boc-deprotection and coupling to the Boc protected linker to access compound **19**. Subsequent acid-mediated Boc-deprotection and coupling to the JQ1-derived carboxylic acid then afforded MZ3 (**2**). Esterification of the Hyp-OH moiety in MZ3 was found to be sluggish with poor conversion of MZ3, and significant amounts of side-product formation, with the major side products putatively corresponding to the *N*-acyl-isourea side-products of dicyclohexylcarbodiimide (DCC) and the respective carboxylic acid. To account for the sluggish reactivity of MZ3, and the suggested consumption of reagents by side-product formation, a second batch of reagents was added in the reactions, and 4-dimethylaminopyridine (DMAP) loadings were increased. In the case of the MZ3-AspA conjugate, even direct exposure of MZ3 to freshly prepared AspA^32^ (4.5 equiv.; estimated purity less than 80%), DCC (3.4 equiv.), DMAP (2.0 equiv.), and NEt3 (10.2 equiv.) did not lead to full MZ3 conversion after 2 days. Therefore, a second batch of reagents (DCC, AspA, and NEt3) was added. For the synthesis of the All-C-AspA analog (**22**), a preliminary test reaction with DCC resulted in significant side-product formation likely corresponding to the *N*-acyl-isourea side-product of DCC and cyclopentanecarboxylic acid based on UHPLC-MS analysis and purification by preparative reversed-phase HPLC proved difficult. Therefore, the synthesis of the All-C-AspA analog (**22**) was carried out using diisopropylcarbodiimide (DIC) instead of DCC. No attempts were undertaken to optimize the reaction conditions for the esterification reactions of MZ3.

**Figure 4.**
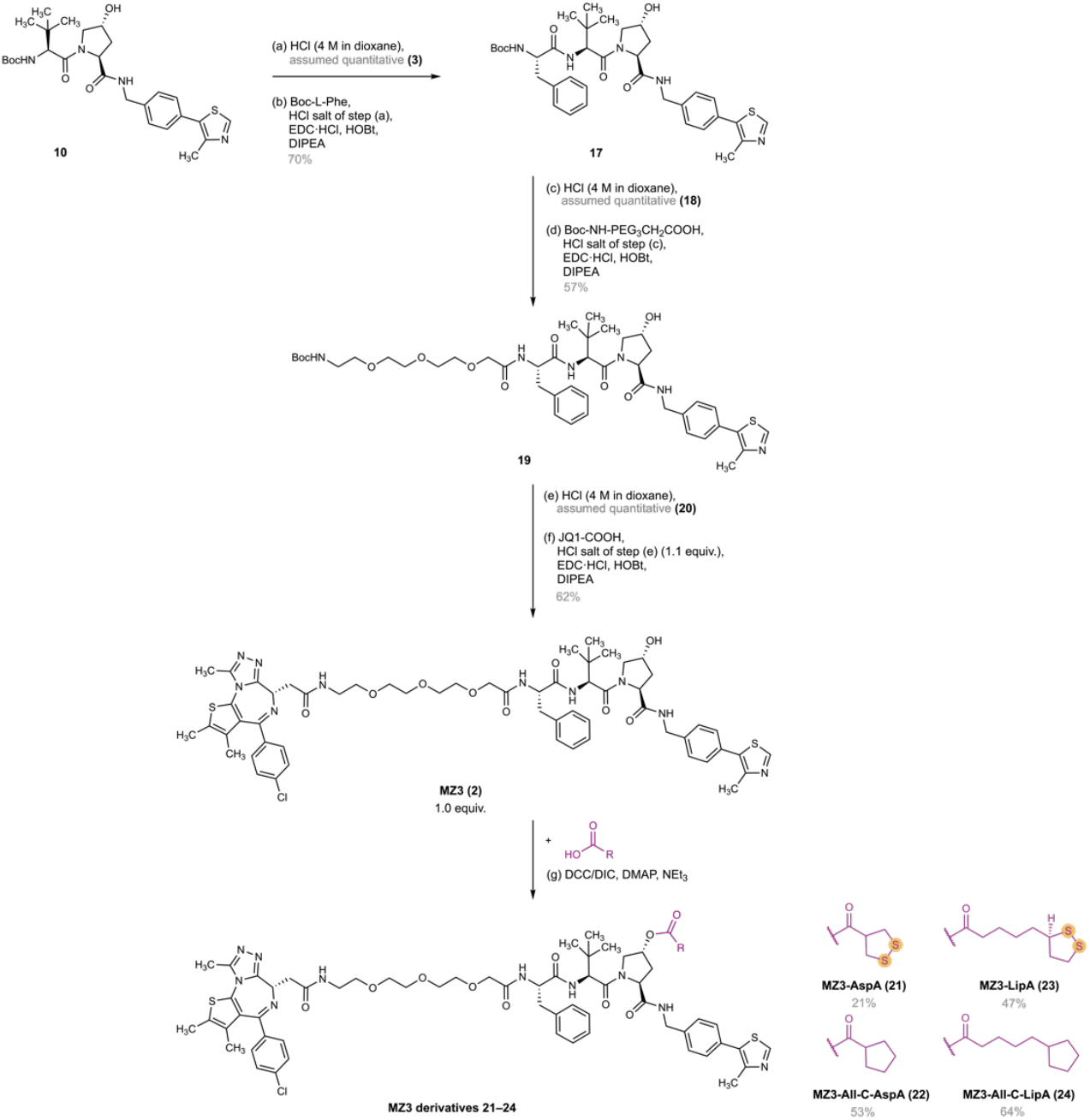
Synthesis of MZ3 and MZ3 derivatives **21-24.** (a) HCl (4 M in dioxane, 20.0 equiv.), CH_2_Cl_2_ (0.2 M), 0 to 25 °C, 5 h; (b) Boc-L-Phe (1.0 equiv.), HCl salt of step (a) (1.1 equiv.), EDC.HCl (1.3 equiv.), HOBt (1.3 equiv.), DIPEA (3.0 equiv.), CH_2_Cl_2_ (0.1 M), 0 to 25 °C, 21 h; (c) HCl (4 M in dioxane, 4.0 equiv.), CH_2_Cl_2_ (0.2 M), 25 °C, 16 h; (d) Boc-NH-PEG_3_CH_2_COOH (1.0 equiv.), HCl salt of step (c) (1.1 equiv.), EDC.HCl (2.6 equiv.), HOBt (2.6 equiv.), DIPEA (7.0 equiv.), CH_2_Cl_2_ (0.1 M), 0 to 25 °C, 25 h; (e) HCl (4 M in dioxane, 4.0 equiv.), CH_2_Cl_2_ (0.2 M), 0 to 25 °C, 70 h; (f) JQ1-COOH (1.0 equiv.), HCl salt of step (e) (1.1 equiv.), EDC.HCl (2.6 equiv.), HOBt (2.6 equiv.), DIPEA (7.0 equiv.), CH_2_Cl_2_ (0.1 M), 0 to 25 °C, 24 h; (g) for detailed procedures on the esterification, see Supporting information.

### Cell viability studies

To study the ability of the compounds to cause cytotoxicity in cancer cells, the effect of the compounds was tested in a cellular system using MTT assays. HeLa cells were treated with 2-fold serial dilutions of test compounds spanning a concentration range of 0.089–45.45 µM. After exposure for 72-73 h, the cell viability was analyzed by MTT assays. For the MZ1-derived series, similar or increased cytotoxicity was observed for the new derivatives compared to their respective parent compound (Fig. 5A-B). At the highest inhibitor concentration tested (45.45 µM), a cell viability of 15.7±5.4% (mean±SEM) was observed for MZ1. In comparison, higher cytotoxicity levels were reached for all MZ1-conjugates: MZ1-AspA (**13**): 2.1±0.2%; MZ1-All-C-AspA (**14**): 4.1±1.1%; MZ1-LipA (**15**): 5.4±0.6%; MZ1-All-C-LipA (**16**): 5.7±1.3% (mean±SEM). For the MZ1 derivatives **15** and **16**, an increase in potency was observed compared to the parent compound (Fig. 5B). Interestingly however, similar cell-viability curves were observed testing the MZ1-dithiolane conjugates and the corresponding control compounds, the All-C analogs that cannot utilize thiol-mediated uptake for cellular entry. These results suggest that uptake mechanisms other than or in addition to thiol-mediated uptake can play a role in this system.

**Figure 5.**
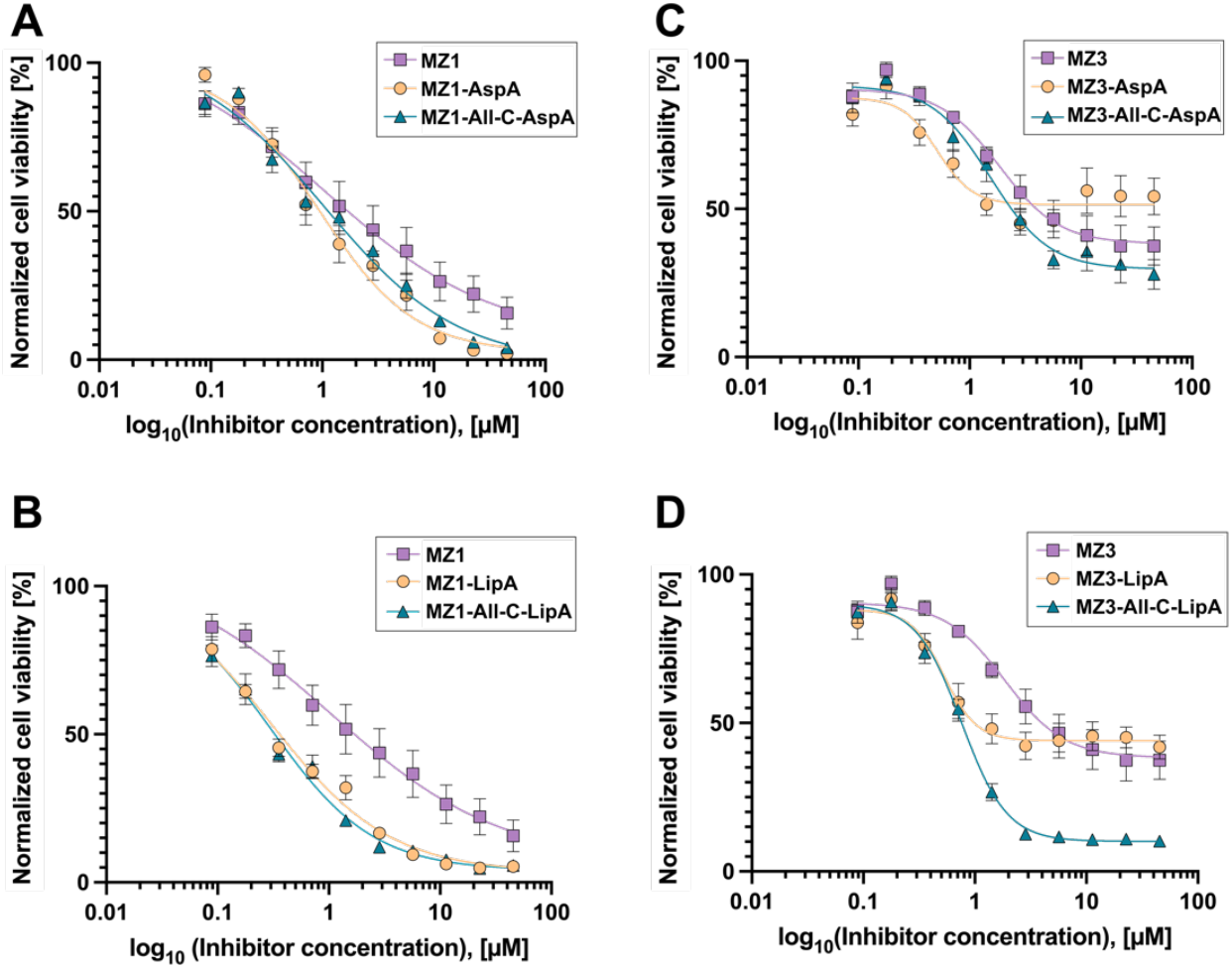
Cell viability curves of HeLa cells after exposure for 72-73 h to the indicated concentration of test compounds of the MZ1-derived series (A-B), and the MZ3-derived series (C-D) spanning a concentration range of 0.089–45.45 µM by means of MTT assay. Data refer to mean±SEM from experiments carried out in biological triplicate each in 2-3 technical replicates. Data analysis and visualization performed with GraphPad Prism^®^ version 10.3.1 (509)^33^ and version 10.4.1 (627)^34^. Data was normalized and fitted to a non-linear regression curve using the sigmoidal curve fit (sigmoidal, 4PL, X is concentration) without constraints. See Supporting Information for details.

For MZ3, lower levels of cytotoxicity were reached compared to MZ1 (Fig. 5A *vs*. C), which is in line with literature reports describing decreased cellular activity of MZ3 compared to MZ1.^21,26^ For the MZ3-derived series, interpretation of the results has to be considered with care as the high bottom plateaus reached for multiple compounds could indicate solubility limits reached in the concentration range tested (Fig. 5C-D) amongst other effects that might come into play at high concentrations of the compounds. Interestingly, for the MZ3-AllC-LipA derivative (**24**) a significantly higher cytotoxicity level was reached compared to MZ3 and the other MZ3 derivatives. Similar as in the MZ1-derived series, the apparent IC50 of the All-C-LipA derivative **24** was found to be significantly lower than for the parent compound MZ3. Surprisingly, higher apparent cytotoxicity was observed for the C-equivalent **24** compared to the dithiolane-conjugate MZ3-LipA (**23**) (Fig. 5D). However, it cannot be excluded that the MZ3-LipA derivative (**23**) reached a solubility limit before its full potential of cytotoxicity was observed. In this context, the similar initial trend of both curves suggests similar potencies for compound **23** and **24** but lower solubility limits for MZ3-LipA (**23**).

Unexpectedly, the cytotoxic properties of the C-analogs were similar or even exceeded the cytotoxicity observed for the corresponding dithiolane conjugates. This result suggests effects other than or in addition to thiol-mediated uptake to play a role for the uptake of the acylated MZ1/MZ3 derivatives. Especially the acylation with longer aliphatic chains such as LipA and the corresponding All-C equivalent was found to constitute a promising modification to increase the cytotoxicity of the parent compound. The reasons behind the increased cytotoxicity warrant further investigations. Supporting our observation, recent patent data suggest acyl modifications of the Hyp-OH moiety in MZ1 including relatively lipophilic moieties as lauric acid might increase cellular uptake by endocytosis-mediated mechanisms.^35^ Therefore, a potential explanation for the increased cytotoxicity of the LipA and All-C-LipA derivatives compared to the respective parent compounds could be an endocytosis-leveraging uptake mechanism of the acylated derivatives resulting in increased cellular uptake.

## Acknowledgements

The authors thank the Swiss National Science Foundation (212603) for financial support. The authors thankfully acknowledge Dr. Annabell Martin and Dr. Juan Tamez Fernandez of the group of Prof. Dr. Rivera-Fuentes (University of Zurich, Department of Chemistry), for their supervision and training of Isabella Ferrara to conduct the MTT assays. The authors further want to express their gratitude to Prof. Dr. Pablo Rivera-Fuentes for the possibility to conduct the MTT assays in his group. The authors thankfully acknowledge Annika Altorfer (University of Zurich, Department of Chemistry) for her work on the re-synthesis of molecules. 5-Cyclopentylpentanoic acid was previously synthesized by Dr. Inga S. Shchelik in the Gademann group at the University of Zurich.

## Author contributions

**Isabella Ferrara:** Conceptualization, methodology, investigation, formal analysis, writing– original draft, visualization, and project administration.

**Karl Gademann:** Conceptualization, funding acquisition, project administration, resources, supervision, validation, writing– review, and editing.

## Experimental procedures

Detailed information on the Experimental procedures and data on the characterization of compounds are provided in the Supporting Information.

## Data availability

Additional data in support of this manuscript has been made available by the authors via zenodo and can be accessed free of charge via DOI: 10.5281/zenodo.15275300.

## Notes

The authors declare no competing interests.

## Notes

### Competing Interest Statement

The authors have declared no competing interest.

