## Supplementary Information for "Evaluation of Thiol-Mediated Uptake in VHL-Based PROTACs"

### 1. Experimental procedures

#### 1.1. General

Unless indicated differently, all chemicals used were of reagent grade, purchased from commercial sources and used as received. Solvents used in the reactions were obtained from commercial sources and used as received or used from a mBraun MB-SPS solvent purification system equipped with a MB-KOL-A columns (2× for $CH_2Cl_2$). If not indicated differently, solvents used for setting up the reactions were anhydrous. Reactions were monitored *via* UHPLC-Ms and/or TLC (Silica gel 60 coated alumina plates, Merck® 60 F254, 20x20 cm, visualization by UV absorbance at 254 nm, or staining with aqueous $KMnO_4$-solution, and subsequent heating). Removal of solvents *in vacuo* was performed using a rotary evaporator at 40 °C water bath temperature unless noted otherwise. Purification by flash column chromatography was carried out on silica gel (0.040-0.063 mm, Merck).

**Nuclear magnetic resonance spectra (NMR):** $^1H$ NMR, $^{13}C$ NMR, and 2-D NMR spectra (COSY, TOCSY, HSQC, HMBC, NOESY) were recorded on the instruments Bruker Avance Avance NEO 500 instruments (500 MHz, 126 MHz, Probes: 5 mm BBI, 5 mm BBOF Cryoprobe, or 5 mm BBO), or Bruker Avance AVI-, AVII-, or Avance NEO 400 instruments (400 MHz, 101 MHz, Probes: 5 mm QNP, 5 mm BBO, 5 mm BBFO (iProbe), or 5 mm BBFO (SmartProbe)) in the indicated deuterated solvent at 25 °C. The spectra were analyzed using MestReNova *v14.1.2-25024*.[1] The chemical shift (δ) in ppm of the reference solvent peak was set to the value reported by Fulmer *et al.*[2]

**Ultra-high performance liquid chromatography mass spectrometry (UHPLC-MS)** was used to monitor reactions and analyze experiments and products. Samples were prepared in MeOH at a final concentration of ≤ 100 µg/mL, filtered (4 mm syringe filter, PTFE (hydrophilic), pore size: 0.22 µm, obtained from BGB Analytik AG). UHPLC-MS analysis was carried out using one of the following machines.
**UHPLC-MS A:** Measurements were run on an Ultimate 3000 LC system (Thermo Fisher Scientific) coupled to a triple quadrupole Quantum Ultra EMR MS (Thermo Fisher Scientific) on a reversed-phase column (Kinetex® EVO C18; 1.7 µm; 100 Å, 50 × 2.1 mm; Phenomenex). The equipment of the LC comprised an HPG-3400RS pump, a WPS-3000TRS autosampler, a TCC-3000RS column oven and a Vanquish DAD detector (all Thermo Fisher Scientific). The following solvent system was used: $H_2O$ + 0.1% HCOOH (A), MeCN + 0.1% HCOOH (B). Measurements were run with a flow rate of 0.4 mL/min. The equipment of the LC comprised an H-ESI II ion source (source temperature: 250 °C, capillary temperature: 270 °C, capillary voltage: 3.5 kV) and datasets were obtained at resolution 0.7 on Q3 in centroid mode.
**UHPLC-MS B**: Measurements were run on a Synapt G2 HR-ESI-QTOF-MS (Waters, Milford, USA) fitted with an electrospray ion source (ESI), coupled to an Aquity UPLC (Waters, Milford, USA). Samples were run on a reversed-phase column (Waters Acquity BEH C18; 1.7 µm; 50 × 2.1 mm). The equipment of the LC comprised a column manager, a sample manager, a binary solvent manager, and an eλ diode array detector (all Waters Aquity Ultra Performance LC). The following solvent system was used: $H_2O$ + 0.04% HCOOH + 0.02% TFA (A), MeCN + 0.04% HCOOH + 0.02% TFA (B). UV spectra were monitored over a range of 190 – 300 nm. Runs were performed using a linear gradient of 10-95% B over 3 min (then 2 min 95% B). The ESI was used in positive ionization mode (source temperature: 120 °C, capillary voltage: 3.0 kV, sampling cone 40 V, extraction cone 4 V, cone gas ($N_2$) 4 L/h, desolvation gas ($N_2$) 800 L/min, mass analyzer in resolution mode: mass range 100 – 2000 *m/z* with a scan rate of 1 Hz). Calibration of the mass was performed to an accuracy of under 2 ppm (50-2500 *m/z*) using a

5 mM aqueous solution of sodium formate and the lock masses of caffeine (*m/z* 195.0882, 0.7 ng/mL) and leucine-enkephalin (*m/z* 556.2771, 2 ng/mL).
**UHPLC-MS C:** Measurements were run on a Dionex Ultimate 3000 LC system (*Thermo Fisher Scientific*, Waltham, USA) connected to a QTOF *Compact* mass spectrometer (*Bruker Daltonics*, Bremen, Germany). The equipment of the LC comprised an HPG-3400RS pump, a WPS-3000 autosampler, a TCC-3100 column oven and a Vanquish DAD-3000RS detector (all Thermo Fisher Scientific). Separation was performed with an Kinetex® EVO C18 column (1.7 µm particle size; 100 Å, 50 × 2.1 mm; Phenomenex) kept at 40°C. The following solvent system was used: $H_2O$ + 0.1% HCOOH (A), MeCN + 0.1% HCOOH (B). Measurements were run with a flow rate of 0.4 mL/min. The mass spectrometer was operated in positive electrospray ionization mode (dry temperature: 200 °C, capillary voltage:4.5 kV, endplate offset: -0.5 kV).

**High resolution electrospray ionization mass spectrometry (HRMS): On-flow injection:** High resolution mass spectra were measured on a QExactive MS instrument with a heated ESI source (ThermoFisher Scientific, Bremen, Germany) following separation on the connected Dionex Ultimate 3000 UHPLC system (ThermoFischer Scientifics, Germering, Germany). Samples were prepared in the indicated solvent system (50 µg/mL) and injected with an *XRS* auto-sampler (*CTC*, Zwingen, Switzerland) (injection volume: 1 µL; flow rate: 120 µL/min). Ion source parameters: spray voltage: 3.0 kV, capillary temperature: 280 °C, sheath gas: 30 L/min, aux gas: 8 L/min, s-lens RF level: 55.0, aux gas temperature: 250 °C. Full scan MS was performed in alternating alternating (+)/(−)-ESI mode (mass ranges: 80–1200 *m/z*, 133–2000 *m/z*, or 200–3000 *m/z,* resolution: 70000 (full width half-maximum), automatic gain control target: $3\times10^6$, maximum allowed ion transfer time: 30 ms). Calibration of the mass was performed to an accuracy of less than 2 ppm using *Pierce*® ESI calibration solutions (*ThermoFisher Scientific*, Rockford, USA) and the lock masses of frequently encountered erucamide (*m/z* 338.34174, (+)-ESI) and palmitic acid (*m/*z 255.23295, (−)-ESI).

**Infrared spectra (IR)** were acquired on a *SpectrumTwo* FT-IR spectrometer (Perkin-Elmer) with a *Specac Golden Gate*™ attenuated total reflection (ATR) device.

**Preparative high performance liquid chromatography (Prep HPLC):** Purification *via* preparative HPLC was performed on a *Prominence* modular HPLC system (*Shimadzu*) coupled to an *SPD-20A* UV/Vis detector (*Shimadzu*). Purification conditions were first evaluated using the analytical instrument with a reversed-phase (RP) column (*Gemini NX* C18, 3 µm, 10 Å, 150 mm × 4.6 mm). Suitable conditions were then used on the preparative HPLC on a RP column (*Gemini NX* C18, 5 µm, 110 Å, 250 mm × 21.2 mm). The Equipment of the LC comprised a *CBM-20A* system controller, *LC-20A* solvent delivery unit, a *DGU-20A* degassing unit, and a *FRC-10A* fraction collector (all *Shimadzu*). Solvents and conditions are listed in the individual experiments. Samples were filtered over a Discovery® DSC-18 SPE cartridge prior to purification by RP prep-HPLC.

**Specific optical rotations** $[\alpha]_D^T$ were acquired on a *Jasco P-2000 Polarimeter* at the temperature indicated. Analyte solutions were measured in MeOH (c is given in g/100 mL). The dimension for $[\alpha]_D^T$ is ° $cm^3$ $dm^{-1}$ $g^{-1}$.

Isolated yields as indicated in the individual experiments refer to the calculated yields based on the obtained amounts of product in comparison to the maximum theoretical yields and are not corrected for the purity. Where indicated, the purity of the products was estimated based on $^1H$ NMR analysis. Where no purity is provided, no significant amount of impurity was observed based on $^1H$ NMR analysis.

### 1.2. Characterization and preparation of compounds

#### *1.2.1. Preparation and characterization of the Boc-protected common intermediate (S,R,S)-AHPC*

##### 1.2.1.1. Preparation of *tert*-butyl (4-(4-methylthiazol-5-yl)benzyl)carbamate (6)[3] (IF356)

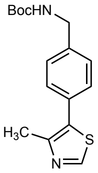

*tert*-Butyl (4-(4-methylthiazol-5-yl)benzyl)carbamate **6** was synthesized after Steinebach *et al.*[3] with minor modifications. To a suspension of *tert*-butyl (4-bromobenzyl)carbamate (**4**, 2.00 g, 6.99 mmol, 1.0 equiv.), $K_2CO_3$ (oven dried at 150 °C, 1.93 g, 13.98 mmol, 2.0 equiv.), and $Pd(OAc)_2$ (15.7 mg, 69.9 µmol, 1 mol%) in anhydrous DMA (20 mL, 0.3 M) was added 4-methylthiazole (**5**, 1.27 mL, 13.98 mmol, 2.0 equiv.). The reaction was heated at 130 °C for 5 h, before it was allowed to cool to 25 °C. The mixture was then diluted with water (80 mL), and the aqueous phase extracted with $CH_2Cl_2$ (3 × 80 mL). The combined organic phase was washed with brine (2 × 200 mL), dried over anhydrous $MgSO_4$, filtered, and concentrated *in vacuo* to provide the crude material. The crude material was further purified via silica gel column chromatography using an eluent of pentane/EtOAc = 7:3. The product was obtained as a colorless to pale yellow solid in a yield of 71% (1.50 g, 4.93 mmol).

##### 1.2.1.2. Characterization of *tert*-butyl (4-(4-methylthiazol-5-yl)benzyl)carbamate (6)[3] (IF356)

**$^{1}$H NMR** (500 MHz, DMSO-$d_6$) δ (ppm) 8.98 (s, 1H), 7.47 – 7.41 (m, 3H), 7.35 – 7.29 (m, 2H), 4.16 (d, $J$ = 6.2 Hz, 2H), 2.45 (s, 3H), 1.40 (s, 9H). **$^{13}$C NMR** (126 MHz, DMSO-$d_6$) δ (ppm) 155.8, 151.5, 147.8, 140.1, 131.1, 129.8, 128.9, 127.4, 77.9, 43.0, 28.3, 16.0. The NMR data is in agreement with the data reported in literature.[3] **HRMS ESI**(+) (MeOH) calculated for $C_{16}H_{21}O_2N_2S^+$ $[M+H]^+$: 305.13183, found: 305.13152. **R$_f$** (EtOAc/pentane = 3/7): 0.3.

#### 1.2.1.3. Preparation of *tert*-butyl (2*S*,4*R*)-4-hydroxy-2-((4-(4-methylthiazol-5-yl)benzyl)carbamoyl)pyrrolidine-1-carboxylate (8)[3–5] (IF357, IF358)

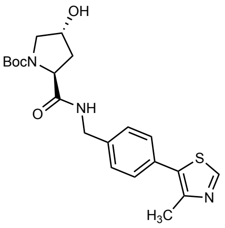

*tert*-Butyl (2*S*,4*R*)-4-hydroxy-2-((4-(4-methylthiazol-5-yl)benzyl) carbamoyl)pyrrolidine-1-carboxylate (**8**) was synthesized similar to Steinebach *et al.,* but with modifications as follows.[3] The Boc-deprotected HCl-salt of *tert*-Butyl (4-(4-methylthiazol-5-yl)benzyl)carbamate (**7**) was prepared by dissolving the Boc-protected amine **6** (1.48 g, 4.86 mmol, 1.0 equiv.) in anhydrous $CH_2Cl_2$ (30 mL, 0.2 M), followed by slow addition of HCl (4 M solution in 1,4-dioxane, 4.86 mL, 19.4 mmol, 4.0 equiv.) at 25 °C. The reaction was stirred at 25 °C for 24 h, before the solvent was removed *in vacuo* at 40 °C water bath temperature. The crude Boc-deprotected HCl salt **7** was obtained as a pale yellow-green solid that was used directly in the next step without purification. **$^1$H NMR** (500 MHz, DMSO-$d_6$) δ (ppm) 9.12 (s, 1H), 8.60 (s, 3H), 7.64 – 7.59 (m, 2H), 7.57 – 7.51 (m, 2H), 4.09 – 4.01 (m, 2H), 2.47 (s, 3H). **$^{13}$C NMR** (126 MHz, DMSO-$d_6$) δ (ppm)152.3, 147.7, 134.1, 131.3, 131.0, 129.6, 129.1, 41.7, 15.8. The $^1$H NMR spectroscopic data is in good agreement with literature reported data.[5] **HRMS ESI(+)** (MeOH) calculated for $C_{11}H_{13}N_2S^+$ $[M+H]^+$: 205.07940, found: 205.07957.

To a suspension of Boc-Hyp-OH (1.02 g, 4.42 mmol, 1.0 equiv.), EDC·HCl (1.10 g, 5.74 mmol, 1.3 equiv.), and HOBt (0.78 g, 5.74 mmol, 1.3 equiv.) in anhydrous $CH_2Cl_2$ (40 mL, 0.1 M) was added at 25 °C DIPEA (2.19 mL, 13.3 mmol, 3.0 equiv.) and the resulting mixture stirred for 5 min, before it was cooled to 0 °C. At 0 °C was added the crude Boc-deprotected HCl salt **7** as a solid (1.1 equiv.). The mixture was allowed to warm to 25 °C and stirred at 25 °C for 18 h. The mixture was then diluted with $CH_2Cl_2$ (100 mL), and the organic phase washed with brine (1 × 100 mL), and saturated aqueous $NaHCO_3$ solution (1 × 100 mL), dried over anhydrous $MgSO_4$, filtered, and concentrated in vacuo to provide the crude material as a colorless solid. Purification via NP column chromatography on silica gel using an eluent of $CH_2Cl_2$/MeOH = 95/5 afforded after concentration *in vacuo* the product **8** as a colorless solid in a yield of 78% (1.43 g, 3.42 mmol).

#### 1.2.1.4. Characterization of *tert*-butyl (2*S*,4*R*)-4-hydroxy-2-((4-(4-methylthiazol-5-yl)benzyl)carbamoyl) pyrrolidine-1-carboxylate (8)[3–5] (IF358)

**$^1$H NMR** (500 MHz, DMSO-$d_6$) δ (ppm) 8.98 (s, 1H), 8.53 – 8.42 (m, 1H), 7.46 – 7.33 (m, 4H), 5.03 – 4.96 (m, 1H), 4.40 – 4.32 (m, 1H), 4.29 – 4.13 (m, 3H), 3.49 – 3.35 (m, 1H), 3.31 – 3.25 (m, 1H), 2.44 (s, 3H), 2.13 – 2.00 (m, 1H), 1.90 – 1.79 (m, 1H), 1.41 (s, 3H), 1.25 (s, 6H). **$^{13}$C NMR** (126 MHz, DMSO-$d_6$) δ (ppm) (putatively mixture of rotamers) 172.5, 172.3, 154.0, 153.6, 151.5, 151.5, 147.8, 147.8, 139.6, 139.5, 131.1, 131.1, 130.0, 129.7, 128.8, 128.7, 128.1, 127.5, 78.7, 78.5, 68.5, 67.8, 58.9, 58.8, 55.1, 54.8, 41.8, 41.5, 39.6, 38.7, 28.1, 27.9, 15.9, 15.9. The NMR spectroscopic data is in agreement with literature reports.[3] *(Note: Steinebach et al. describe the existence of rotamers, however only report the $^{13}$C signals for the major rotamer, complicating direct comparison with our data.*[3] *The $^{13}$C NMR spectrum shown in literature however clearly displays two signal sets in agreement with our data.*[3]*)* **HRMS ESI(+)** (MeOH) calculated for $C_{21}H_{28}O_4N_3S^+$ $[M+H]^+$: 418.17950, found: 418.17962. **$R_f$** ($CH_2Cl_2$/MeOH = 95/5) = 0.2.

#### 1.2.1.5. Preparation of *tert*-butyl ((S)-1-((2*S*,4*R*)-4-hydroxy-2-((4-(4-methylthiazol-5-yl)benzyl) carbamoyl)pyrrolidin-1-yl)-3,3-dimethyl-1-oxobutan-2-yl)carbamate (10)[3–5] (IF359, IF360)

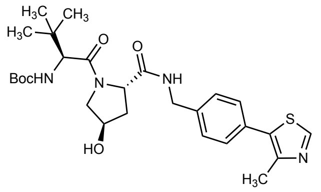

*tert*-Butyl ((S)-1-((2*S*,4*R*)-4-hydroxy-2-((4-(4-methylthiazol-5-yl)benzyl)carbamoyl)pyrrolidin-1-yl)-3,3-dimethyl-1-oxobutan-2-yl)carbamate (**10**) was synthesized similar to Steinebach *et al.,* but with modifications as follows.[3]

The Boc-deprotected HCl-salt of *tert*-Butyl (2*S*,4*R*)-4-hydroxy-2-((4-(4-methylthiazol-5-yl)benzyl) carbamoyl)pyrrolidine-1-carboxylate (**9**) was prepared by dissolving the Boc-protected amine **8** (1.43 g, 3.42 mmol, 1.0 equiv.) in anhydrous $CH_2Cl_2$ (21 mL, 0.2 M), followed by slow addition of HCl (4 M solution in 1,4-dioxane, 3.42 mL, 13.69 mmol, 4.0 equiv.) at 25 °C. The reaction was stirred at 25 °C for 5.5 h, before the solvent was removed *in vacuo* at 40 °C water bath temperature. The crude Boc-deprotected HCl salt **9** was obtained as a colorless solid that was used directly in the next step without purification.

**$^1$H NMR** (500 MHz, MeOD) δ (ppm) 9.79 (s, 1H), 9.00 (t, $J$ = 5.7 Hz, 1H), 7.60 – 7.46 (m, 4H), 4.63 – 4.57 (m, 1H), 4.56 – 4.47 (m, 3H), 3.47 – 3.39 (m, 1H), 3.34 – 3.32 (m, 1H), 2.59 (s, 3H), 2.54 – 2.45 (m, 1H), 2.13 – 2.04 (m, 1H). The $^1$H NMR spectroscopic data is in good agreement with literature reported data.[4,5] **HRMS ESI(+)** (MeOH) calculated for $C_{16}H_{20}O_2N_3S^+$ $[M+H]^+$: 318.12707, found: 318.12727.

To a suspension of Boc-Tle-OH (0.72 g, 3.11 mmol, 1.0 equiv.), EDC·HCl (0.78 g, 4.05 mmol, 1.3 equiv.), and HOBt (0.55 g, 4.05 mmol, 1.3 equiv.) in anhydrous $CH_2Cl_2$ (33 mL, 0.1 M) was added at 25 °C DIPEA (1.54 mL, 9.34 mmol, 3.0 equiv.) and the resulting mixture stirred for 5 min, before it was cooled to 0 °C. At 0 °C was added the crude Boc-deprotected HCl salt **9** as a solid (1.1 equiv.). The mixture was stirred at 0 °C for 15 min, before it was allowed to warm to 25 °C and stirred at 25 °C for 18 h. The mixture was then diluted with $CH_2Cl_2$ (50 mL), and the organic phase washed with brine (1 × 50 mL), and saturated aqueous $NaHCO_3$ solution (1 × 50 mL), dried over anhydrous $MgSO_4$, filtered, and concentrated in vacuo to provide the crude material as a colorless solid. Purification *via* NP column chromatography on silica gel using an eluent of $CH_2Cl_2$/MeOH = 95/5 afforded after concentration *in vacuo* the product **10** as a colorless solid in a yield of 66% (1.09 g, 2.05 mmol).

#### 1.2.1.6. Characterization of *tert*-butyl ((S)-1-((2*S*,4*R*)-4-hydroxy-2-((4-(4-methylthiazol-5-yl)benzyl) carbamoyl)pyrrolidin-1-yl)-3,3-dimethyl-1-oxobutan-2-yl)carbamate (10)[3–5] (IF360)

**$^1$H NMR** (400 MHz, DMSO-$d_6$) δ (ppm) 8.98 (s, 1H), 8.59 (t, $J$ = 6.0 Hz, 1H), 7.45 – 7.31 (m, 4H), 6.49 (d, $J$ = 9.3 Hz, 1H), 5.15 (d, $J$ = 3.5 Hz, 1H), 4.48 – 4.32 (m, 3H), 4.23 (dd, $J$ = 15.9, 5.5 Hz, 1H), 4.15 (d, $J$ = 9.3 Hz, 1H), 3.65 (dd, $J$ = 10.6, 4.0 Hz, 1H), 3.60 (d, $J$ = 10.8 Hz, 1H), 2.44 (s, 3H), 2.08 – 1.99 (m, 1H), 1.95 – 1.84 (m, 1H), 1.38 (s, 9H), 0.93 (s, 9H). **$^{13}$C NMR** (101 MHz, DMSO-$d_6$) δ (ppm) 172.0, 169.9, 155.4, 151.5, 147.8, 139.5, 131.2, 129.7, 128.7, 127.5, 78.1, 69.0, 58.8, 58.4, 56.4, 41.7, 37.9, 35.4, 28.2, 26.3, 16.0. The NMR spectroscopic data is in good agreement with literature reported data.[3] **HRMS ESI(+)** (MeOH) calculated for $C_{27}H_{39}O_5N_4S^+$ $[M+H]^+$: 531.26357, found: 531.26381. **$R_f$** ($CH_2Cl_2$/MeOH = 95/5) = 0.2.

#### 1.2.2. *Preparation and characterization of MZ1-derivatives*

##### 1.2.2.1. Preparation of *tert*-butyl ((S)-13-((2*S*,4*R*)-4-hydroxy-2-((4-(4-methylthiazol-5-yl)benzyl) carbamoyl)pyrrolidine-1-carbonyl)-14,14-dimethyl-11-oxo-3,6,9-trioxa-12-azapentadecyl)carbamate (11)[6–10] (IF363, IF364)

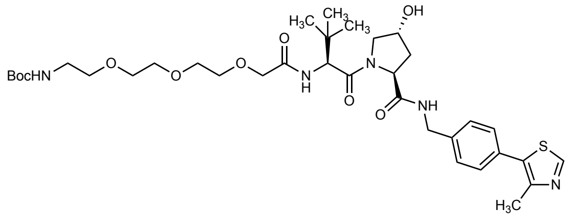

*tert*-Butyl ((*S*)-13-((2*S*,4*R*)-4-hydroxy-2-((4-(4-methylthiazol-5-yl)benzyl) carbamoyl) pyrrolidine-1-carbonyl)-14,14-dimethyl-11-oxo-3,6,9-trioxa-12-azapentadecyl)carbamate (**11**) was synthesized similar to Gui *et al.,*[6] and as described in the patent literature by Plewe *et al.*[8] but with modifications as follows.

The Boc-deprotected HCl-salt of *tert*-butyl ((*S*)-1-((2*S*,4*R*)-4-hydroxy-2-((4-(4-methylthiazol-5-yl)benzyl)carbamoyl)pyrrolidin-1-yl)-3,3-dimethyl-1-oxobutan-2-yl)carbamate (**3**, VH032 HCl salt, (*S,R,S*)-AHPC hydrochloride)[4–7,11,12] was prepared after Buckley *et al.*[11] and Crew *et al.*[12] with modifications as follows: The Boc-protected amine **10** (0.41 g, 0.77 mmol, 1.0 equiv.) was dissolved in anhydrous $CH_2Cl_2$ (3.4 mL, 0.2 M), and cooled to 0 °C. At 0 °C, to the solution was added HCl (4 M solution in 1,4-dioxane, 3.85 mL, 15.41 mmol, 20.0 equiv.), and the reaction allowed to warm to 25 °C and stirred at 25 °C for 3.5 h. The solvent was removed *in vacuo* at 40 °C water bath temperature. The crude Boc-deprotected HCl salt **3** was obtained as a colorless solid that was used directly in the next step without purification.
**$^1$H NMR** (500 MHz, MeOD) δ (ppm) 9.63 (s, 1H), 7.56 – 7.52 (m, 2H), 7.52 – 7.46 (m, 2H), 4.72 – 4.63 (m, 1H), 4.59 – 4.51 (m, 2H), 4.44 – 4.36 (m, 1H), 4.06 (s, 1H), 3.87 – 3.80 (m, 1H), 3.75 – 3.69 (m, 1H), 2.56 (s, 3H), 2.35 – 2.26 (m, 1H), 2.13 – 2.04 (m, 1H), 1.14 (s, 9H). The $^1$H NMR spectroscopic data is in good agreement with literature reported data.[5] **$^{13}$C NMR** (126 MHz, MeOD) δ (ppm) 174.1, 168.6, 155.4, 144.5, 141.8, 136.2, 130.5, 129.3, 129.1, 71.2, 61.0, 60.4, 58.1, 43.7, 39.1, 35.8, 26.7, 13.9. *Note: Upon comparing the $^{13}$C NMR data with the data reported by Yan et al.*[5] *a minor discrepancy was recognized as the authors report one less aromatic peak (136.2 was not listed), and one more aliphatic peak (49.0 absent in our data).*[5] *As the compound however should display 7 different aromatic $^{13}$C signals (2 with double intensity) our data is in line with the expected signal pattern.* **HRMS ESI(+)** (MeOH) calculated for $C_{22}H_{31}O_3N_4S^+$ $[M+H]^+$: 431.21114, found: 431.21034.

To a suspension of Boc-NH-PEG$_3$-CH$_2$COOH (0.22 g, 0.71 mmol, 1.0 equiv.), EDC·HCl (0.18 g, 0.91 mmol, 1.3 equiv.), and HOBt (0.12 g, 0.91 mmol, 1.3 equiv.) in anhydrous $CH_2Cl_2$ (7 mL, 0.1 M) was added at 25 °C DIPEA (0.35 mL, 2.10 mmol, 3.0 equiv.), and the resulting mixture cooled to 0 °C. At 0 °C was added the crude Boc-deprotected HCl salt **3** as a solid (1.1 equiv.). The mixture was stirred at 0 °C for 5 min, before it was allowed to warm to 25 °C and stirred at 25 °C for 44 h. The mixture was then diluted with $CH_2Cl_2$ (15 mL), and the organic phase washed with brine (1 × 15 mL), and saturated aqueous $NaHCO_3$ solution (1 × 15 mL), dried over anhydrous $MgSO_4$, filtered, and concentrated *in vacuo* to provide the crude material as an orange oil. Purification *via* NP column chromatography on silica gel using an eluent of $CH_2Cl_2$/MeOH = 95/5 afforded after concentration *in vacuo* the product **11** as a colorless solid in a yield of 43% (0.22 g, 0.30 mmol).

#### 1.2.2.2. Characterization of *tert*-butyl ((*S*)-13-((2*S*,4*R*)-4-hydroxy-2-((4-(4-methylthiazol-5-yl)benzyl) carbamoyl)pyrrolidine-1-carbonyl)-14,14-dimethyl-11-oxo-3,6,9-trioxa-12-azapentadecyl)carbamate (11)[6–10] (IF364)

**$^1$H NMR** (400 MHz, DMSO-$d_6$) δ (ppm) 8.98 (s, 1H), 8.59 (t, *J* = 6.0 Hz, 1H), 7.45 – 7.38 (m, 5H), 6.73 (t, *J* = 5.8 Hz, 1H), 5.15 (d, *J* = 3.5 Hz, 1H), 4.56 (d, *J* = 9.6 Hz, 1H), 4.48 – 4.32 (m, 3H), 4.25 (dd, *J* = 15.8, 5.6 Hz, 1H), 3.97 (s, 2H), 3.69 – 3.46 (m, 11H), 3.04 (q, *J* = 6.0 Hz, 2H), 2.44 (s, 3H), 2.06 (dd, *J* = 16.6, 4.7 Hz, 1H), 1.90 (ddd, *J* = 13.0, 8.8, 4.5 Hz, 1H), 1.36 (s, 9H), 0.94 (s, 9H). *(Note: multiplet reported at around 3.35 ppm[10] likely overlaps with water peak).* The $^1$H NMR data is in overall accordance with literature.[10] **$^1$H NMR** (400 MHz, $CDCl_3$) δ (ppm) 8.68 (s, 1H), 7.39 – 7.31 (m, 4H), 4.75 (t, *J* = 7.9 Hz, 1H), 4.62 – 4.43 (m, 3H), 4.39 – 4.28 (m, 1H), 4.17 – 3.92 (m, 3H), 3.73 – 3.47 (m, 12H), 3.34 – 3.22 (m, 2H), 2.52 (s, 3H), 2.19 – 2.06 (m, 1H), 1.43 (s, 9H), 0.96 (s, 9H). **$^1$H NMR** (500 MHz, MeOD) δ (ppm) 8.88 (s, 1H), 7.49 – 7.39 (m, 4H), 4.70 (s, 1H), 4.60 – 4.47 (m, 3H), 4.36 (d, *J* = 15.4 Hz, 1H), 4.11 – 4.01 (m, 2H), 3.89 – 3.85 (m, 1H), 3.81 (dd, *J* = 11.0, 3.8 Hz, 1H), 3.75 – 3.58 (m, 9H), 3.48 (t, *J* = 5.6 Hz, 2H), 3.19 (t, *J* = 5.7 Hz, 2H), 2.48 (s, 3H), 2.27 – 2.19 (m, 1H), 2.13 – 2.04 (m, 1H), 1.42 (s, 9H), 1.05 (s, 9H). **HRMS ESI(+)** (MeOH) calculated for $C_{35}H_{54}O_9N_5S^+$ $[M+H]^+$: 720.36368, found: 720.36346. **R$_f$** ($CH_2Cl_2$/MeOH = 95/5) = 0.1–0.2.

#### 1.2.2.3. Preparation of MZ1 (1)[7,9,11,13] (IF367, IF375)

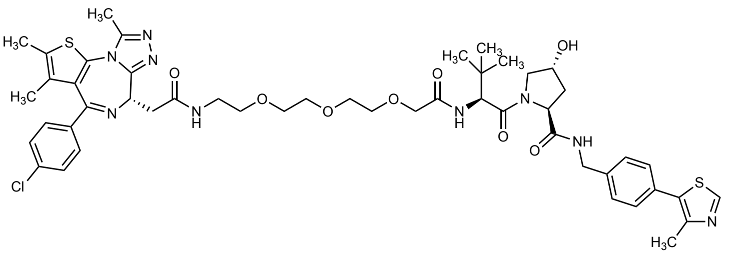

The Boc-deprotected HCl-salt of *tert*-butyl ((*S*)-13-((2*S*,4*R*)-4-hydroxy-2-((4-(4-methylthiazol-5-yl)benzyl)carbamoyl)pyrrolidine-1-carbonyl)-14,14-dimethyl-11-oxo-3,6,9-trioxa-12-azapentadecyl) carbamate (**12**)[7,9,11] was prepared after Buckley *et al.*[11] with modifications as follows: The Boc-protected amine **11** (0.289 g, 0.40 mmol, 1.0 equiv.) was dissolved in anhydrous $CH_2Cl_2$ (2.0 mL, 0.2 M). To the solution was added HCl (4 M solution in 1,4-dioxane, 0.40 mL, 1.61 mmol, 4.0 equiv.), and the reaction stirred at 25 °C for 18 h. The solvent was removed *in vacuo* at 40 °C water bath temperature, and the residue taken up in $CH_2Cl_2$ and concentrated again (2×). The crude Boc-deprotected HCl salt **12** was obtained as a colorless solid (sticky foam) that was used directly in the next step without purification.

**$^1$H NMR (500 MHz, MeOD)** δ (ppm) 9.82 (s, 1H), 7.58 – 7.48 (m, 4H), 4.70 (s, 1H), 4.61 – 4.37 (m, 4H), 4.12 – 4.05 (m, 2H), 3.90 (dt, *J* = 11.2, 1.7 Hz, 1H), 3.81 (dd, *J* = 11.0, 3.8 Hz, 1H), 3.77 – 3.66 (m, 11H), 3.13 (t, *J* = 5.0 Hz, 2H), 2.59 (s, 3H), 2.26 (ddt, *J* = 13.2, 7.5, 1.9 Hz, 1H), 2.08 (ddd, *J* = 13.5, 9.5, 4.4 Hz, 1H), 1.05 (s, 9H). The $^1$H NMR spectroscopic data is in good agreement with literature reported data.[9] **$^{13}$C NMR** (126 MHz, MeOD) δ (ppm) 174.4, 172.1, 171.9, 156.1, 143.4, 142.3, 136.9, 130.5, 129.4, 128.6, 71.9, 71.6, 71.4, 71.2, 71.1, 71.0, 67.9, 60.9, 58.4, 58.2, 43.6, 40.7, 39.1, 37.0, 27.0, 26.9, 13.5. **HRMS ESI(+)** (MeOH) calculated for $C_{30}H_{46}O_7N_5S^+$ $[M+H]^+$: 620.31125, found: 620.31198.

To a suspension of JQ1-COOH (0.15 g, 0.37 mmol, 1.0 equiv.), EDC·HCl (0.18 g, 0.95 mmol, 2.6 equiv.), and HOBt (0.12 g, 0.95 mmol, 2.6 equiv.) in anhydrous $CH_2Cl_2$ (4 mL, 0.1 M) was added at 25 °C DIPEA (0.42 mL, 2.55 mmol, 7.0 equiv.), and the resulting mixture stirred at

25°C for 15 min, before it was cooled to 0 °C. At 0 °C was added the crude Boc-deprotected HCl salt **12** as a solid (1.1 equiv.). The mixture was allowed to warm to 25 °C and stirred at 25 °C for 27 h. The mixture was then diluted with $CH_2Cl_2$ (15 mL), and the organic phase washed with brine (1 × 15 mL), and saturated aqueous $NaHCO_3$ solution (1 × 15 mL), dried over anhydrous $MgSO_4$, filtered, and concentrated in vacuo to provide the crude material as a colorless to pale yellow solid (foam). Purification of the crude material was performed by prep RP-HPLC using a gradient of 30 to 60% B (MeCN+0.1% FA) over 60 min (LC time program (time - %B): 0 min - 30%, 5 min - 30%, 65 min – 60 %, 95 min – 60%, 96 min – 100%, 106 min – 100%). Product containing fractions were combined and concentrated *in vacuo* at 40 °C water bath temperature and further dried *in vacuo* to provide the product MZ1 (**1**) ($R_t$: 37.0 min) as a colorless solid in a yield of 60% (221 mg, 0.22 mmol).

#### 1.2.2.4. Characterization of MZ1 (1)[7,9,11,13] (IF375)

**$^1$H NMR** (500 MHz, $CDCl_3$) δ (ppm) 8.67 (s, 1H), 7.93 (t, *J* = 5.6 Hz, 1H), 7.42 – 7.27 (m, 9H), 7.25 – 7.22 (m, 1H), 4.83 (t, *J* = 7.9 Hz, 1H), 4.75 – 4.63 (m, 3H), 4.56 – 4.46 (m, 2H), 4.34 – 4.24 (m, 2H), 4.15 – 4.08 (m, 2H), 3.74 – 3.47 (m, 14H), 3.37 – 3.29 (m, 2H), 2.61 (s, 3H), 2.50 (s, 3H), 2.49 – 2.42 (m, 1H), 2.39 (s, 3H), 2.18 – 2.10 (m, 1H), 1.66 (s, 3H), 0.98 (s, 9H). The $^1$H NMR spectroscopic data is in good agreement with literature.[13] **$^{13}$C NMR** (126 MHz, $CDCl_3$) δ (ppm) 171.4 (*likely two signals stacked*), 171.0, 171.0, 163.9, 156.0, 150.4, 149.9, 148.6, 138.4, 136.8, 136.7, 132.0, 131.8, 131.2, 131.0, 130.9, 130. 9, 130.1, 129.6, 128.9, 128.2, 71.8, 71.0, 70.6, 70.5, 70.5, 70.3, 70.2, 58.9, 57.2, 56.8, 54.3, 43.3, 39.9, 38.2, 36.4, 35.5, 26.6, 16.2, 14.6, 13.2, 11.9. **HRMS ESI(+)** (MeOH) calculated for $C_{49}H_{60}ClN_9O_8S_2Na^+$ $[M+Na]^+$: 1024.35870, found: 1024.35963.

#### 1.2.2.5. Esterification of MZ1

The esterification procedures were adapted from the procedure described by Liu *et al.*[14] with the specified modifications.

#### 1.2.2.6. Preparation of MZ1-AspA ester (13) (IF393)

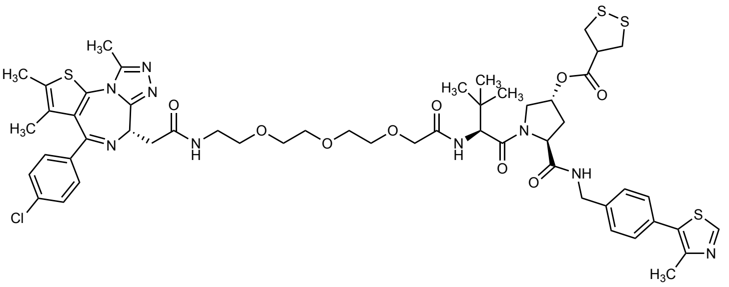

10xstock solution of DMAP: A stock solution of DMAP (3.7 mg) in anhydrous $CH_2Cl_2$ (1 mL) was prepared. MZ1 (**1**, 30.0 mg, 29.9 µmol, 1.0 equiv), and freshly prepared AspA (crude product, estimated purity: approx. 70%, 19.3 mg, 89.8 µmol, 3.0 equiv.), and DCC (18.5 mg, 89.8 µmol, 3.0 equiv.), were suspended in anhydrous $CH_2Cl_2$ (1.8 mL). To the suspension was added 200 µL of the 10xstock solution of DMAP in $CH_2Cl_2$ (5.98 µmol, 0.2 equiv.), followed by $NEt_3$ (25.2 µL, 180 µmol, 6.0 equiv.). The reaction mixture was sparged with $N_2$ for 2 min, and then stirred at 25 °C for 41 h. After 41 h, the solvent was removed *in vacuo* to provide the crude material. The crude material was resuspended in MeCN and filtered over a Discovery DSC-18 SPE cartridge (50 mg) and concentrated to provide the pre-purified material for further purification *via* RP prep-HPLC. Further purification was performed by prep RP-HPLC using a gradient of 50 to 80% B (MeCN+0.1% FA) over 60 min (LC time program (time - %B): 0 min - 50%, 5 min - 50%, 65 min – 80 %, 95 min – 80%, 96 min – 100%, 106 min – 100%). The product containing fractions were combined and concentrated *in vacuo* at 44 °C water bath temperature and further dried *in vacuo* to provide the product MZ1-AspA (**13**) ($R_t$: approx. 19.4 min) as a colorless solid in a yield of 35% (11.8 mg, 10.4 µmol). *Note: The purity of the crude AspA used is difficult to determine, as polymeric material does not, or only poorly dissolve in NMR solvent (turbid mixture). The estimated purity of 70% is thus, just a rough approximation. To account for the poor purity of the crude AspA mixture it was used in excess.*

#### 1.2.2.7. Characterization of MZ1-AspA ester (13) (IF393)

**$^1$H NMR** (500 MHz, $CDCl_3$) δ (ppm) 8.67 (s, 1H), 7.78 (t, $J$ = 6.0 Hz, 1H), 7.40 – 7.36 (m, 2H), 7.32 – 7.26 (m, 7H), 7.20 (t, $J$ = 5.8 Hz, 1H), 5.42 – 5.39 (m, 1H), 4.82 (t, $J$ = 7.8 Hz, 1H), 4.67 – 4.62 (m, 1H), 4.52 (d, $J$ = 9.1 Hz, 1H), 4.49 (dd, $J$ = 15.3, 6.6 Hz, 1H), 4.31 (dd, $J$ = 15.0, 5.6 Hz, 1H), 4.25 – 4.20 (m, 1H), 4.06 (s, 2H), 3.84 (dd, $J$ = 11.8, 4.3 Hz, 1H), 3.73 – 3.63 (m, 8H), 3.61 – 3.56 (m, 2H), 3.51 – 3.34 (m, 7H), 3.29 – 3.21 (m, 2H), 2.68 – 2.59 (m, 1H), 2.61 (s, 3H), 2.50 (s, 3H), 2.39 (s, 3H), 2.34 – 2.26 (m, 1H), 1.66 (s, 3H), 0.98 (s, 9H). **$^{13}$C NMR** (126 MHz, $CDCl_3$) δ (ppm) 171.60, 171.2, 170.8, 170.7, 170.2, 163.9, 155.9, 150.4, 150.0, 148.6, 138.3, 136.9, 136.8, 132.2, 131.8, 131.1, 131.0, 130.9, 130.8, 130.0, 129.5, 128.8, 128.2, 74.2, 71.1, 70.8, 70.6, 70.6, 70.3, 70.1, 58.7, 57.0, 54.4, 54.0, 50.8, 43.3, 41.6, 41.3, 39.8, 38.9, 35.3, 33.6, 26.6, 16.2, 14.6, 13.3, 11.9. **HRMS ESI(+)** (MeOH) calculated for $C_{53}H_{64}ClN_9O_9S_4Na^+$ $[M+Na]^+$: 1156.32906, found: 1156.32877. **Specific Rotation** $[\alpha]_D^{23°C}$ = +13.1 (c = 0.43, MeOH). **Specific Rotation** $[\alpha]_D^{23°C}$ = +8.9 (c = 0.37, MeOH). **FT-IR** ($CDCl_3$): $\nu$ ($cm^{-1}$) 3313w, 3070w, 2925m, 2870w, 1736m, 1656s, 1592m, 1531s, 1487m, 1432m, 1419m, 1369w, 1347w, 1268w, 1232w, 1175m, 1109m, 1091m, 1059w, 1036w, 1015w, 918w, 844w, 805w, 731m, 645w, 606w, 562w, 551w, 486w.

#### 1.2.2.8. Preparation of MZ1-All-C-AspA ester (14) (IF395)

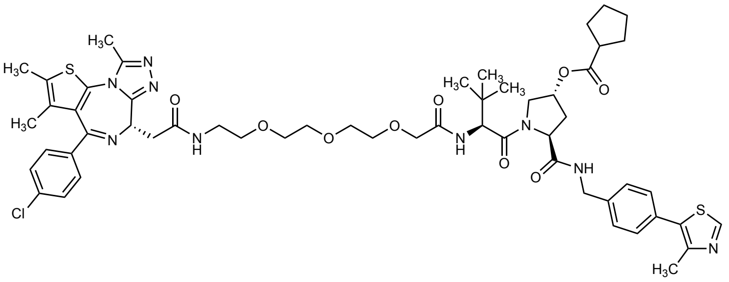

10xstock solution of DMAP: A stock solution of DMAP (3.7 mg) in anhydrous $CH_2Cl_2$ (1 mL) was prepared. MZ1 (**1**, 30.0 mg, 29.9 µmol, 1.0 equiv), and DCC (18.5 mg, 89.8 µmol, 3.0 equiv.), were suspended in anhydrous $CH_2Cl_2$ (1.8 mL). To the suspension was added cyclopentanecarboxylic acid (8.13 µL, 74.8 µmol, 2.5 equiv.), followed by addition of $NEt_3$ (29.4 µL, 209 µmol, 7.0 equiv.), and 200 µL of the 10xstock solution of DMAP in $CH_2Cl_2$ (5.98 µmol, 0.2 equiv.). The reaction mixture was stirred at 25 °C for 42 h. After 42 h, the solvent was removed *in vacuo* to provide the crude material. The crude material was resuspended in MeCN and filtered over a Discovery DSC-18 SPE cartridge (50 mg) and concentrated to provide the pre-purified material for further purification *via* RP prep-HPLC. Further purification was performed by prep RP-HPLC using a gradient of 50 to 95% B (MeCN+0.1% FA) over 60 min (LC time program (time - %B): 0 min - 50%, 5 min - 50%, 65 min – 95 %, 95 min – 95%, 96 min – 100%, 106 min – 100%). The product containing fractions were combined and concentrated *in vacuo* at 44 °C water bath temperature and further dried *in vacuo* to provide the product MZ1-All-C-AspA (**14**) ($R_t$: 19.4 min) as a colorless solid in a yield of 44% (14.6 mg, 13.3 µmol).

#### 1.2.2.9. Characterization of MZ1-All-C-AspA ester (14) (IF395)

**$^1$H NMR** (500 MHz, $CDCl_3$) δ (ppm) 8.67 (s, 1H), 7.68 (t, *J* = 6.0 Hz, 1H), 7.41 – 7.37 (m, 2H), 7.34 – 7.23 (m, 7H), 7.12 – 7.08 (m, 1H), 5.42 – 5.36 (m, 1H), 4.80 (dd, *J* = 8.2, 6.6 Hz, 1H), 4.66 – 4.61 (m, 1H), 4.60 (d, *J* = 9.3 Hz, 1H), 4.49 (dd, *J* = 15.0, 6.3 Hz, 1H), 4.35 (dd, *J* = 15.0, 5.7 Hz, 1H), 4.10 – 3.97 (m, 3H), 3.91 (dd, *J* = 11.4, 5.0 Hz, 1H), 3.73 – 3.34 (m, 14H (*Note: observed integral: ~16; should correspond to 14 Hs*)), 2.74 – 2.58 (m, 2H), 2.61 (s, 3H), 2.50 (s, 3H), 2.39 (d, *J* = 0.9 Hz, 3H), 2.27 – 2.19 (m, 1H), 1.89 – 1.79 (m, 2H), 1.79 – 1.69 (m, 2H), 1.68 – 1.65 (m, 3H), 1.67 – 1.62 (m, 2H), 1.57 – 1.50 (m, 2H), 0.97 (s, 9H). **$^{13}$C NMR** (126 MHz, $CDCl_3$) δ (ppm) 176.4, 171.1, 170.8, 170.8, 169.8, 163.8, 155.9, 150.4, 149.9, 148.6, 138.4, 136.9, 136.8, 132.2, 131.8, 131.1, 130.9, 130.7, 130.0, 129.6, 128.8, 128.2, 72.6, 71.1, 70.9, 70.6, 70.4, 70.1, 58.8, 56.7, 54.4, 53.9, 43.7, 43.3, 39.7, 39.0, 35.7, 33.7, 30.1, 30.0, 26.6, 25.9, 16.2, 14.6, 13.2, 11.9. *Note: Two C-signals not observed or potentially stack with other signals.* **HRMS ESI(+)** (MeOH) calculated for $C_{55}H_{69}ClN_9O_9S_2^+$ $[M+H]^+$: 1098.43427, found: 1098.43397. **Specific Rotation** $[\alpha]_D^{23°C}$ = +14.7 (c = 0.63, MeOH). **FT-IR** ($CDCl_3$): $\nu$ (cm$^{-1}$) 3306w, 3068w, 2952w, 2869w, 1730m, 1668s, 1644s, 1592m, 1528s, 1487m, 1432m, 1417s, 1367m, 1350m, 1309m, 1268m, 1231m, 1178m, 1145m, 1106s, 1089s, 1059m, 1036m, 1014m, 966w, 937w, 843m, 804w, 748w, 648w, 606w, 561w, 550w, 484w, 445w.

#### 1.2.2.10. Preparation of MZ1-(*R*)-LipA ester (15) (IF370)

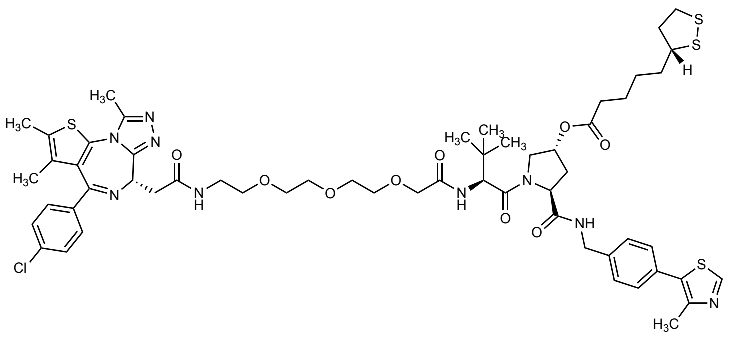

To a solution of MZ1 (**1**, 30.0 mg, 29.9 µmol, 1.0 equiv), *R*-lipoic acid (12.3 mg, 59.8 µmol, 2.0 equiv.), DCC (N,N'-Dicyclohexylcarbodiimide) (9.3 mg, 44.9 µmol, 1.5 equiv.), and DMAP (4-Dimethylaminopyridine) (0.4 mg, 3.0 µmol, 0.1 equiv.) in anhydrous $CH_2Cl_2$ (2.0 mL, 0.01 M), was added $NEt_3$ (4.2 µL, 29.9 µmol, 1.0 equiv.). The reaction mixture was stirred at 25 °C for 73 h. After 73 h, the solvent was removed in vacuo to provide the crude material. The crude material was resuspended in MeCN and filtered over a Discovery DSC-18 SPE cartridge (50 mg) and concentrated to provide the pre-purified material for further purification via RP prep-HPLC. Further purification was performed by prep RP-HPLC using a gradient of 50 to 95% B (MeCN+0.1% FA) over 60 min (LC time program (time - %B): 0 min - 50%, 5 min - 50%, 65 min – 95 %, 95 min – 60%, 96 min – 100%, 106 min – 100%). The product containing fraction was concentrated *in vacuo* at 40 °C water bath temperature and further dried *in vacuo* to provide the product MZ1-LipA (**15**) ($R_t$: 25.8 min) as a colorless solid in a yield of 38% (13.6 mg, 11.4 µmol).

#### 1.2.2.11. Characterization of MZ1-(*R*)-LipA ester (15) (IF370)

**$^1$H NMR** (500 MHz, $CDCl_3$) δ (ppm) 8.67 (s, 1H), 7.71 (t, *J* = 6.0 Hz, 1H), 7.42 – 7.36 (m, 2H), 7.33 – 7.24 (m, 7H), 7.18 (t, *J* = 5.6 Hz, 1H), 5.43 – 5.37 (m, 1H), 4.83 – 4.77 (m, 1H), 4.67 – 4.61 (m, 1H), 4.55 (d, *J* = 9.2 Hz, 1H), 4.48 (dd, *J* = 15.0, 6.2 Hz, 1H), 4.34 (dd, *J* = 15.1, 5.7 Hz, 1H), 4.11 – 3.97 (m, 3H), 3.89 (dd, *J* = 11.5, 4.8 Hz, 1H), 3.73 – 3.29 (m, 15H (*Note: observed integral:17H; should correspond to 15 Hs)*), 3.20 – 3.05 (m, 2H), 2.65 – 2.56 (m, 1H), 2.60 (s, 3H), 2.50 (s, 3H), 2.48 – 2.41 (m, 1H), 2.39 (s, 3H), 2.35 – 2.21 (m, 3H), 1.94 – 1.86 (m, 1H), 1.66 (s, 3H), 1.72 – 1.57 (m, 4H), 1.43 (dtd, *J* = 16.1, 9.0, 6.1 Hz, 2H), 0.98 (s, 9H). **$^{13}$C NMR** (126 MHz, $CDCl_3$) δ (ppm) 173.2, 171.0, 170.9, 170.8, 169.9, 163.8, 155.9, 150.4, 149.9, 148.6, 138.4, 136.9, 136.8, 132.2, 131.8, 131.1, 131.0, 130.9, 130.7, 130.0, 129.5, 128.8, 128.2, 72.9, 71.1, 70.8, 70.6 (supposably 2 signals stacked), 70.4, 70.1, 58.8, 56.8, 56.5, 54.4, 54.0, 43.3, 40.4, 39.7, 39.0, 38.6, 35.6, 34.7, 34.0, 33.7, 28.8, 26.6, 24.6, 16.2, 14.6, 13.3, 11.9. **HRMS ESI(+)** (MeOH) calculated for $C_{57}H_{73}ClN_9O_9S_4^+$ $[M+H]^+$: 1190.40971, found: 1190.41215. **Specific Rotation** $[\alpha]_D^{23°C}$ = +37.9 (c = 0.72, MeOH). **FT-IR** ($CDCl_3$): $\nu$ ($cm^{-1}$) 3312w, 3070w, 2925m, 2870w, 1734m, 1667s, 1646s, 1592m, 1531s, 1487m, 1433m, 1418m, 1378w, 1368w, 1350w, 1315w, 1303w, 1268w, 1231w, 1174m, 1111m, 1091m, 1060w, 1036w, 1014w, 966w, 918w, 845w, 805w, 731m, 645w, 606w, 574w, 551w, 513w, 486w, 446w.

#### 1.2.2.12. Preparation of MZ1-All-C-(*R*)-LipA ester (16) (IF378)

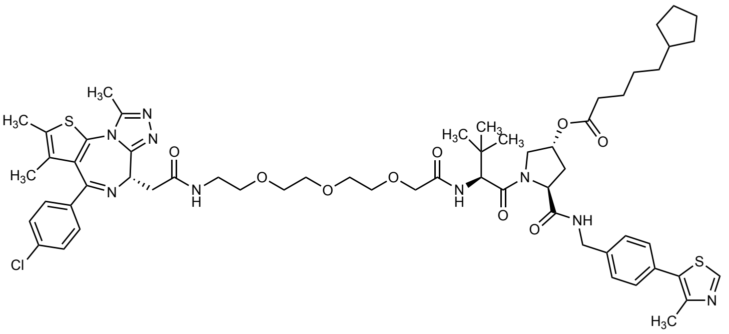

A stock solution of DMAP in $CH_2Cl_2$ was prepared by dissolving DMAP (3.7 mg) in anhydrous $CH_2Cl_2$ (1 mL). To a mixture of MZ1 (**1**, 29.1 mg, 29.0 µmol, 1.0 equiv.), 5-cyclopentylpentanoic acid (prepared by Dr. Inga S. Shchelik; 12.4 mg, 72.6 µmol, 2.5 equiv.), and DCC (19.3 mg, 93.7 µmol, 3.2 equiv.) in anhydrous $CH_2Cl_2$ (1.8 mL, 0.01 M) was added 200 µL of the stock solution of DMAP in $CH_2Cl_2$ (final amount: 6.1 µmol, 0.2 equiv.)., followed by $NEt_3$ (16.8 µL, 120 µmol, 4.1 equiv.). The reaction mixture was stirred at 25 °C for 64 h. After 64 h, the solvent was removed *in vacuo* to provide the crude material. The crude material was resuspended in MeCN and filtered over a Discovery DSC-18 SPE cartridge (50 mg) and concentrated to provide the pre-purified material for further purification *via* prep RP-HPLC. Further purification was performed by preparative RP-HPLC using a gradient of 50 to 95% B (MeCN+0.1% FA) over 60 min (LC time program (time - %B): 0 min - 50%, 5 min - 50%, 65 min – 95%, 95 min – 95%, 96 min – 100%, 106 min – 100%). The product containing fractions were combined and concentrated *in vacuo* at 40 °C water bath temperature and further dried *in vacuo* to provide the product MZ1-All-C-LipA (**16**) ($R_t$: 35.5±1.0 min) as a colorless solid in a yield of 52% (17.6 mg, 15.2 µmol).

#### 1.2.2.13. Characterization of MZ1-All-C-(*R*)-LipA ester (16) (IF378)

**$^1$H NMR** (500 MHz, $CDCl_3$) δ (ppm) 8.67 (s, 1H), 7.70 (t, *J* = 6.0 Hz, 1H), 7.41 – 7.37 (m, 2H), 7.33 – 7.22 (m, 7H), 7.18 – 7.14 (m, 1H), 5.43 – 5.36 (m, 1H), 4.84 – 4.78 (m, 1H), 4.64 (t, *J* = 7.0 Hz, 1H), 4.56 (d, *J* = 9.2 Hz, 1H), 4.48 (dd, *J* = 15.0, 6.3 Hz, 1H), 4.34 (dd, *J* = 15.0, 5.6 Hz, 1H), 4.10 – 3.99 (m, 3H), 3.91 (dd, *J* = 11.4, 5.0 Hz, 1H), 3.74 – 3.34 (m, 14H (*Note: observed integral: 17; should correspond to 14 Hs (see Note below)*)), 2.66 – 2.57 (m, 1H), 2.60 (s, 3H), 2.50 (s, 3H), 2.39 (s, 3H), 2.34 – 2.19 (m, 3H), 1.76 – 1.68 (m, 6H), 1.67 (s, 3H), 1.61 – 1.53 (m, 5H), 1.52 – 1.43 (m, 2H), 1.33 – 1.24 (m, 4H), 1.03 (tq, *J* = 5.7, 3.6, 2.7 Hz, 1H), 0.97 (s, 9H). *Note: An overall sum of H signals of approx. 82 was observed; This should correspond to 76 signals.* **$^{13}$C NMR** (126 MHz, $CDCl_3$) δ (ppm) 173.6, 171.1, 170.9, 170.8, 169.9, 163.8, 155.9, 150.4, 149.9, 148.6, 138.4, 136.9, 136.8, 132.2, 131.8, 131.1, 131.0, 130.9, 130.7, 130.0, 129.5, 128.8, 128.2, 72.7, 71.1, 70.9, 70.6 (likely 2 signals stacked), 70.4, 70.1, 58.8, 56.8, 54.4, 53.9, 43.3, 40.1, 39.7, 39.0, 35.9, 35.6, 34.3, 33.6, 32.8, 28.4, 26.6, 25.3, 25.1, 16.2, 14.6, 13.2, 11.9. **HRMS ESI(+)** (MeOH) calculated for $C_{59}H_{76}ClN_9O_9S_2Na^+$ $[M+Na]^+$: 1176.47882, found: 1176.47852. **Specific Rotation** $[\alpha]_D^{23°C}$ = +17.4 (c = 0.64, MeOH). **FT-IR** ($CDCl_3$): $\nu$ ($cm^{-1}$) 3306w, 3071w, 2939m, 2867w, 1735m, 1647s, 1592m, 1530s, 1487m, 1433m, 1418m, 1378m, 1368m, 1349m, 1315w, 1303w, 1270m, 1230m, 1172m, 1106m, 1091m, 1061w, 1035w, 1015w, 966w, 915m, 843w, 805w, 729s, 645w, 606w, 562w, 550w, 524w, 485w, 446w, 424w.

#### *1.2.3. Preparation and characterization of MZ3-derivatives*

##### 1.2.3.1. Preparation of *tert*-butyl ((*S*)-1-(((*S*)-1-((2*S*,4*R*)-4-hydroxy-2-((4-(4-methylthiazol-5-yl)benzyl) carbamoyl)pyrrolidin-1-yl)-3,3-dimethyl-1-oxobutan-2-yl)amino)-1-oxo-3-phenylpropan-2-yl)carbamate (17)[13,15] (IF361, IF362)

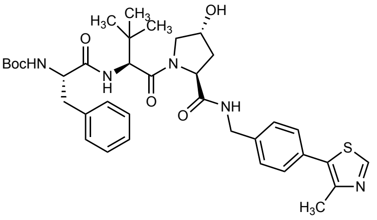

*tert*-Butyl ((*S*)-1-(((*S*)-1-((2*S*,4*R*)-4-hydroxy-2-((4-(4-methylthiazol-5-yl)benzyl)carbamoyl)pyrrolidin-1-yl)-3,3-dimethyl-1-oxobutan-2-yl)amino)-1-oxo-3-phenylpropan-2-yl)carbamate (**17**) was synthesized similar to Galdeano *et al.*[15] but with modifications as follows.

The Boc-deprotected HCl-salt of *tert*-butyl ((*S*)-1-((2*S*,4*R*)-4-hydroxy-2-((4-(4-methylthiazol-5-yl)benzyl)carbamoyl)pyrrolidin-1-yl)-3,3-dimethyl-1-oxobutan-2-yl)carbamate (**3**, VH032 HCl salt, (*S,R,S*)-AHPC hydrochloride)[4–7,11,12] was prepared after Buckley *et al.*[11] and Crew *et al.*[12] with modifications as follows: The Boc-protected amine **10** (0.409 g, 0.771 mmol, 1.0 equiv.) was dissolved in anhydrous $CH_2Cl_2$ (3.4 mL, 0.2 M), and cooled to 0 °C. At 0 °C, to the solution was added HCl (4 M solution in 1,4-dioxane, 3.85 mL, 15.4 mmol, 20.0 equiv.), and the reaction allowed to warm to 25 °C and stirred at 25 °C for 5 h. The solvent was removed *in vacuo* at 40 °C water bath temperature. The crude Boc-deprotected HCl salt **3** was obtained as a colorless solid that was used directly in the next step without purification.

**$^{1}$H NMR** (500 MHz, MeOD) δ (ppm) 9.52 (s, 1H), 7.54 – 7.51 (m, 2H), 7.50 – 7.46 (m, 2H), 4.70 – 4.65 (m, 1H), 4.58 – 4.52 (m, 2H), 4.40 (d, *J* = 15.7 Hz, 1H), 4.06 (s, 1H), 3.85 – 3.80 (m, 1H), 3.75 – 3.70 (m, 1H), 2.55 (s, 3H), 2.34 – 2.26 (m, 1H), 2.13 – 2.06 (m, 1H), 1.14 (s, 9H). The $^{1}$H NMR spectroscopic data is in reasonable agreement with literature reported data.[5]
**$^{13}$C NMR** (126 MHz, MeOD) δ (ppm) 174.1, 168.6, 155.1, 145.2, 141.6, 135.8, 130.4, 129.5, 129.2, 71.2, 61.0, 60.4, 58.1, 43.7, 39.2, 35.8, 26.6, 14.2. *Note: Upon comparing the $^{13}$C NMR data with the data reported by Yan et al.*[5] *a minor discrepancy was recognized as the authors report one less aromatic peak (135.8 was not listed), and one more aliphatic peak (49.0 absent in our data).*[5] *As the compound however should display 7 different aromatic $^{13}$C signals (2 with double intensity) our data is in line with the expected signal pattern.* **HRMS ESI(+)** (MeOH) calculated for $C_{22}H_{31}O_3N_4S^+$ $[M+H]^+$: 431.21114, found: 431.21133.

To a suspension of Boc-L-Phe (0.186 g, 0.701 mmol, 1.0 equiv.), EDC·HCl (0.175 g, 0.911 mmol, 1.3 equiv.), and HOBt (0.124 g, 0.918 mmol, 1.3 equiv.) in anhydrous $CH_2Cl_2$ (7 mL, 0.1 M) was added dropwise at 25 °C DIPEA (348 µL, 2.10 mmol, 3.0 equiv.), and the resulting mixture cooled to 0 °C. At 0 °C was added the crude Boc-deprotected HCl salt **3** as a solid (0.771 mmol, 1.1 equiv.). The mixture was stirred at 0 °C for 5 min, before it was allowed to warm to 25 °C and stirred at 25 °C for 21 h. The mixture was then diluted with $CH_2Cl_2$ (15 mL), and the organic phase washed with brine (1 × 15 mL), and saturated aqueous $NaHCO_3$ solution (1 × 15 mL), dried over anhydrous $MgSO_4$, filtered, and concentrated *in vacuo* to provide the crude material as a pale-yellow oil. Purification *via* NP column chromatography on silica gel using an eluent of $CH_2Cl_2$/MeOH = 95/5 afforded after concentration *in vacuo* the product **17** as a colorless solid in a yield of 70% (0.334 g, 0.493 mmol).

##### 1.2.3.2. Characterization of *tert*-butyl ((*S*)-1-(((*S*)-1-((2*S*,4*R*)-4-hydroxy-2-((4-(4-methylthiazol-5-yl) benzyl)carbamoyl)pyrrolidin-1-yl)-3,3-dimethyl-1-oxobutan-2-yl)amino)-1-oxo-3-phenylpropan-2-yl)carbamate (17)[13,15] (IF362)

**$^1$H NMR** (500 MHz, DMSO-*d$_6$*) δ (ppm) 8.98 (s, 1H), 8.58 (t, *J* = 6.1 Hz, 1H), 7.63 (d, *J* = 9.4 Hz, 1H), 7.43 – 7.37 (m, 4H), 7.28 – 7.22 (m, 4H), 7.20 – 7.14 (m, 1H), 7.11 (d, *J* = 8.7 Hz, 1H), 5.18 – 5.13 (m, 1H), 4.57 – 4.53 (m, 1H), 4.47 – 4.39 (m, 2H), 4.37 (s, 1H), 4.27 – 4.17 (m, 2H), 3.71 – 3.66 (m, 1H), 3.59 (d, *J* = 10.7 Hz, 1H), 3.00 – 2.94 (m, 1H), 2.78 – 2.71 (m, 1H), 2.44 (s, 3H), 2.08 – 2.01 (m, 1H), 1.95 – 1.87 (m, 1H), 1.29 (s, 9H), 0.95 (s, 9H).
**$^{13}$C NMR** (126 MHz, DMSO-*d$_6$*) δ (ppm) 171.9, 171.1, 169.2, 155.3, 151.5, 147.7, 139.5, 138.2, 131.2, 129.7, 129.2, 128.7, 128.0, 127.5, 126.1, 78.2, 68.9, 58.7, 56.5, 56.3, 55.7, 41.7, 37.9, 36.9, 35.9, 28.1, 26.2, 16.0. (*Note: One additional C signal at 26.3 ppm was observed that could not be assigned and could likely come from an impurity*). **HRMS ESI(+)** (MeOH) calculated for $C_{36}H_{48}O_6N_5S^+$ $[M+H]^+$: 678.33198, found: 678.33183. **R$_f$** ($CH_2Cl_2$/MeOH = 95/5): 0.2.

##### 1.2.3.3. Preparation of *tert*-butyl ((13*S*,16*S*)-13-benzyl-16-((2*S*,4*R*)-4-hydroxy-2-((4-(4-methylthiazol-5-yl) benzyl)carbamoyl)pyrrolidine-1-carbonyl)-17,17-dimethyl-11,14-dioxo-3,6,9-trioxa-12,15-diazaoctadecyl)carbamate (19) (IF389, IF390)

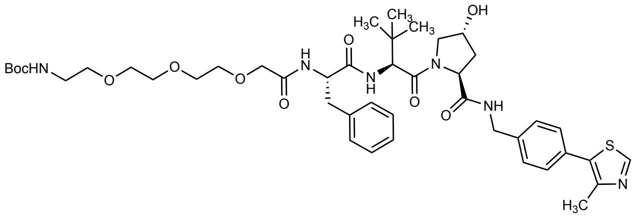

*tert*-Butyl ((13*S*,16*S*)-13-benzyl-16-((2*S*,4*R*)-4-hydroxy-2-((4-(4-methylthiazol-5-yl)benzyl)carbamoyl) pyrrolidine-1-carbonyl)-17,17-dimethyl-11,14-dioxo-3,6,9-trioxa-12,15-diazaoctadecyl) carbamate (**19**) was synthesized analogous to *tert*-butyl ((*S*)-13-((2*S*,4*R*)-4-hydroxy-2-((4-(4-methylthiazol-5-yl)benzyl)carbamoyl)pyrrolidine-1-carbonyl)-14,14-dimethyl-11-oxo-3,6,9-trioxa-12-azapentadecyl)carbamate (**11**) with modifications as follows.

The Boc-deprotected HCl-salt of *tert*-butyl ((*S*)-1-(((*S*)-1-((2*S*,4*R*)-4-hydroxy-2-((4-(4-methylthiazol-5-yl)benzyl)carbamoyl)pyrrolidin-1-yl)-3,3-dimethyl-1-oxobutan-2-yl)amino)-1-oxo-3-phenylpropan-2-yl)carbamate (**18**) was prepared similar to the corresponding TFA salt prepared by Zengerle *et al.*[13] but with modifications as follows: Boc-protected amine **17** (0.349 g, 0.515 mmol, 1.0 equiv.) was dissolved in anhydrous $CH_2Cl_2$ (2.5 mL, 0.2 M). To the solution was added HCl (4 M solution in 1,4-dioxane, 0.515 mL, 2.06 mmol, 4.0 equiv.), and the reaction stirred at 25 °C for 16 h. After 16 h, the reaction was diluted with $CH_2Cl_2$, and the solvent was removed *in vacuo* at 40 °C water bath temperature. The residue was taken up in $CH_2Cl_2$ and the solvent removed again *in vacuo* (2×). The crude Boc-deprotected HCl salt **18** was obtained as a colorless solid that was used directly in the next step without purification.
**$^1$H NMR** (500 MHz, MeOD) δ (ppm) 9.67 (s, 1H), 7.57 – 7.52 (m, 2H), 7.51 – 7.47 (m, 2H), 7.40 – 7.35 (m, 2H), 7.32 – 7.28 (m, 1H), 7.26 – 7.22 (m, 2H), 4.67 (s, 1H), 4.63 – 4.57 (m, 1H), 4.57 – 4.51 (m, 2H), 4.42 (d, *J* = 15.7 Hz, 1H), 4.24 (dd, *J* = 7.7, 6.1 Hz, 1H), 3.90 – 3.85 (m, 1H), 3.82 (dd, *J* = 10.8, 3.9 Hz, 1H), 3.20 (dd, *J* = 14.1, 6.1 Hz, 1H), 3.03 (dd, *J* = 14.2, 7.7 Hz, 1H), 2.57 (s, 3H), 2.28 (ddt, *J* = 13.2, 7.6, 1.8 Hz, 1H), 2.11 (ddd, *J* = 13.3, 9.0, 4.5 Hz,

1H), 1.06 (s, 9H). **$^{13}$C NMR** (126 MHz, MeOD) δ (ppm) 174.5, 171.3, 169.4, 155.6, 144.3, 142.0, 136.4, 135.3, 130.5, 130.4, 130.3, 129.3, 129.0, 128.9, 71.0, 60.8, 59.3, 58.1, 55.3, 43.7, 39.1, 38.6, 36.7, 27.0, 13.8. **HRMS ESI(+)** (MeOH) calculated for $C_{31}H_{40}O_4N_5S^+$ [M+H]$^+$: 578.27955, found: 578.27986.

To a suspension of Boc-NH-PEG$_3$-CH$_2$COOH (0.146 g, 0.475 mmol, 1.0 equiv.), EDC·HCl (0.233 g, 1.217 mmol, 2.6 equiv.), and HOBt (0.164 g, 1.217 mmol, 2.6 equiv.) in anhydrous $CH_2Cl_2$ (4.6 mL, 0.1 M) was added at 25 °C DIPEA (542 µL, 3.28 mmol, 7.0 equiv.), and the resulting mixture stirred at 25 °C for 15 min, before it was cooled to 0 °C. At 0 °C was added the crude Boc-deprotected HCl salt **18** as a solid (0.515 mmol, 1.1 equiv.). The mixture was stirred at 0 °C for 5 min, before it was allowed to warm to 25 °C and stirred at 25 °C for 25 h. The mixture was then diluted with $CH_2Cl_2$ (15 mL), and the organic phase washed with brine (2 × 15 mL), dried over anhydrous $MgSO_4$, filtered, and concentrated *in vacuo* to provide the crude material as a pale brown sticky solid. Purification *via* NP column chromatography on silica gel using an eluent of $CH_2Cl_2$/MeOH = 95/5 afforded after concentration *in vacuo* the product **19** as a colorless solid in a yield of 57% (0.232 g, 0.268 mmol).

##### 1.2.3.4. Characterization of *tert*-butyl ((13*S*,16*S*)-13-benzyl-16-((2*S*,4*R*)-4-hydroxy-2-((4-(4-methylthiazol-5-yl)benzyl)carbamoyl)pyrrolidine-1-carbonyl)-17,17-dimethyl-11,14-dioxo-3,6,9-trioxa-12,15-diazaoctadecyl)carbamate (19) (IF390)

**$^1$H NMR** (500 MHz, $CDCl_3$) δ (ppm) 8.67 (s, 1H), 7.48 – 7.38 (m, 2H), 7.38 – 7.30 (m, 4H), 7.29 – 7.22 (m, 2H (*overlaps with residual solvent signal of $CDCl_3$*)), 7.20 – 7.13 (m, 3H), 7.09 – 7.01 (m, 1H), 5.19 – 5.10 (m, 1H), 4.79 – 4.71 (m, 1H), 4.71 – 4.61 (m, 1H), 4.58 – 4.44 (m, 3H), 4.35 (dd, *J* = 14.9, 5.4 Hz, 1H), 4.03 – 3.96 (m, 1H), 3.95 – 3.85 (m, 2H), 3.64 – 3.54 (m, 7H), 3.53 – 3.47 (m, 4H), 3.30 – 3.18 (m, 2H), 3.13 – 3.07 (m, 1H), 3.05 – 2.97 (m, 1H), 2.58 – 2.52 (m, 1H), 2.51 (s, 3H), 2.21 – 2.11 (m, 1H), 1.43 (s, 9H), 0.90 (s, 9H). **$^{13}$C NMR** (126 MHz, $CDCl_3$) δ (ppm) 171.4, 171.2, 170.9, 170.7, 156.3, 150.4, 148.6, 138.2, 136.6, 131.7, 131.1, 129.7, 129.5, 128.7, 128.3, 127.0, 79.5, 71.1, 70.7, 70.4, 70.3, 70.2, 58.6, 58.0, 57.0, 54.4, 43.4, 40.5, 37.4, 36.0, 35.4, 28.6, 26.6, 16.2. *Note: Two C-signals were not observed or are potentially stacking with other C-signals in the area of the PEG-linker signals.* **FT-IR** ($CDCl_3$): $\nu$ (cm$^{-1}$) 3310m, 3066w, 3029w, 2957m, 2927m, 2873m, 1654s, 1621s, 1527s, 1482m, 1443m, 1418m, 1392m, 1366m, 1337m, 1277m, 1247m, 1170m, 1150m, 1107m, 1031w, 1017w, 967w, 937w, 921w, 854w, 733m, 701m, 645w, 584w, 569w, 551w, 495w. **HRMS ESI(+)** (MeOH) calculated for $C_{44}H_{62}O_{10}N_6SNa^+$ [M+Na]$^+$: 889.41403, found: 889.41346. **R$_f$** ($CH_2Cl_2$/MeOH = 95/5): 0.3. **Specific Rotation** $[\alpha]_D^{23°C}$ = –21.0 (c = 1.00, MeOH).

#### 1.2.3.5. Preparation of MZ3 (2)[13] (IF394, IF397)

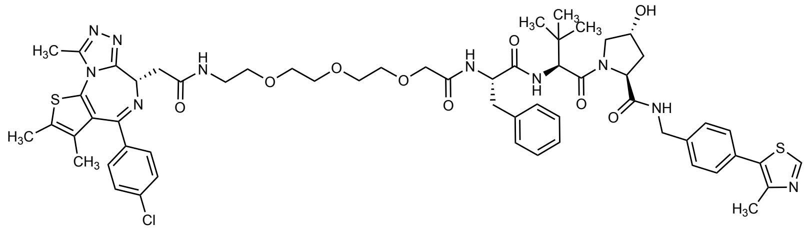

The Boc-deprotected HCl-salt of *tert*-butyl ((13*S*,16*S*)-13-benzyl-16-((2*S*,4*R*)-4-hydroxy-2-((4-(4-methylthiazol-5-yl)benzyl) carbamoyl)pyrrolidine-1-carbonyl)-17,17-dimethyl-11,14-dioxo-3,6,9-trioxa-12,15-diazaoctadecyl)carbamate (**20**) was prepared analogous to the sequence for the preparation of MZ1 after Buckley *et al.*[11] (see above) with modifications as follows: The Boc-protected amine **19** (0.214 g, 0.247 mmol, 1.0 equiv.) was dissolved in anhydrous $CH_2Cl_2$ (1.4 mL, 0.2 M) and the mixture cooled to 0 °C. At 0 °C, to the solution was added HCl (4 M solution in 1,4-dioxane, 247 µL, 0.986 mmol, 4.0 equiv.), and the reaction allowed to warm to 25 °C. The reaction was stirred at 25 °C for 70 h. After 70 h, the reaction was diluted with $CH_2Cl_2$ (2 mL), and the solvent was removed *in vacuo* at 40 °C water bath temperature. The residue was taken up in $CH_2Cl_2$ and concentrated again (2×). The crude Boc-deprotected HCl salt **20** was obtained as a colorless solid that was used directly in the next step without purification.

**$^1$H NMR** (500 MHz, MeOD) δ (ppm) 9.61 (s, 1H), 7.56 – 7.52 (m, 2H), 7.51 – 7.47 (m, 2H), 7.31 – 7.26 (m, 2H), 7.24 – 7.17 (m, 3H), 4.80 (dd, *J* = 8.7, 5.6 Hz, 1H), 4.63 (s, 1H), 4.62 – 4.51 (m, 3H), 4.41 (d, *J* = 15.6 Hz, 1H), 4.02 – 3.91 (m, 2H), 3.88 – 3.84 (m, 1H), 3.81 (dd, *J* = 10.9, 3.9 Hz, 1H), 3.73 – 3.67 (m, 4H), 3.67 – 3.61 (m, 4H), 3.60 – 3.55 (m, 2H), 3.18 – 3.09 (m, 3H), 2.95 (dd, *J* = 13.9, 8.7 Hz, 1H), 2.56 (s, 3H), 2.31 – 2.23 (m, 1H), 2.10 (ddd, *J* = 13.3, 9.0, 4.5 Hz, 1H), 1.04 (s, 9H). **$^{13}$C NMR** (126 MHz, MeOD) δ (ppm) 174.5, 173.0, 172.4, 171.7, 155.4, 144.6, 141.9, 138.0, 136.2, 130.4, 130.4, 129.6, 129.3, 129.1, 127.9, 71.7, 71.5, 71.3, 71.2, 71.1, 71.0, 67.9, 60.8, 59.1, 58.0, 55.2, 43.7, 40.7, 39.1, 38.7, 36.8, 27.0, 13.9. **HRMS ESI(+)** (MeOH) calculated for $C_{39}H_{55}O_8N_6S^+$ $[M+H]^+$: 767.37966, found: 767.38035.

To a suspension of JQ1-COOH (94.6 mg, 0.224 mmol, 1.0 equiv.), EDC·HCl (112 mg, 0.583 mmol, 2.6 equiv.), and HOBt (78.8 mg, 0.583 mmol, 2.6 equiv.) in anhydrous $CH_2Cl_2$ (2.5 mL, 0.1 M) was added dropwise at 25 °C DIPEA (259 µL, 1.57 mmol, 7.0 equiv.), and the resulting mixture stirred at 25°C for 15 min, before it was cooled to 0 °C. At 0 °C was added the crude Boc-deprotected HCl salt **20** as a solid (0.247 mmol, 1.1 equiv.). The mixture was allowed to warm to 25 °C and stirred at 25 °C for 24 h. The mixture was then diluted with $CH_2Cl_2$ (10 mL), and the organic phase washed with brine (1 × 10 mL), and saturated aqueous $NaHCO_3$ solution (1 × 10 mL), dried over anhydrous $MgSO_4$, filtered, and concentrated *in vacuo* to provide the crude material as a colorless solid (foam). For purification the crude was split on several runs: Purification of the crude material was performed by prep RP-HPLC using a gradient of 30 to 60% B (MeCN+0.1% FA) over 60 min (LC time program (time - %B): 0 min - 30%, 5 min - 30%, 65 min – 60 %, 95 min – 60%, 96 min – 100%, 106 min – 100%). Product containing fractions were combined and concentrated *in vacuo* at 44 °C water bath temperature, and further dried *in vacuo* to provide the product MZ3 (**2**) ($R_t$: 45.0 min) as a colorless solid in a yield of 62% (159.2 mg, 0.139 mmol).

#### 1.2.3.6. Characterization of MZ3 (2)[13] (IF397)

**$^1$H NMR (500 MHz, $CDCl_3$)** δ (ppm) 8.67 (s, 1H), 7.86 – 7.80 (m, 1H), 7.67 (d, *J* = 8.0 Hz, 1H), 7.40 – 7.29 (m, 7H), 7.25 – 7.12 (m, 7H), 7.03 (d, *J* = 8.7 Hz, 1H), 4.75 (t, *J* = 7.7 Hz, 1H), 4.70 (t, *J* = 7.0 Hz, 1H), 4.70 – 4.62 (m, 1H), 4.60 (d, *J* = 8.8 Hz, 1H), 4.51 (dd, *J* = 15.1,

6.6 Hz, 1H), 4.48 – 4.45 (m, 1H), 4.30 (dd, $J$ = 15.0, 5.3 Hz, 1H), 4.03 (d, $J$ = 11.0 Hz, 1H), 3.99 – 3.87 (m, 2H), 3.74 – 3.39 (m, 16H), 3.19 – 3.10 (m, 2H), 2.63 (s, 3H), 2.51 (s, 3H), 2.50 – 2.45 (m, 1H), 2.39 (s, 3H), 2.15 – 2.08 (m, 1H), 1.64 (s, 3H), 0.90 (s, 9H). The $^{1}$H NMR spectroscopic data is in overall agreement with literature.[13] *Note: 1xH less in the area 3.19–3.10, and 1xH more in the area of 3.74–3.39 was observed as compared to literature.*[13] **$^{13}$C NMR (126 MHz, $CDCl_3$)** δ (ppm) 171.3, 171.2, 171.2, 170.9, 170.9, 164.1, 156.0, 150.4, 150.0, 148.6, 138.4, 137.1, 136.9, 136.6, 132.1, 131.8, 131.2, 131.1, 131.0, 130.8, 130.0, 129.6, 129.4, 128.8, 128.6, 128.3, 126.6, 70.9, 70.9, 70.4, 70.3, 70.2, 70.1, 70.0, 58.9, 57.7, 57.2, 54.7, 54.3, 43.3, 39.7, 38.4, 36.6, 36.1, 35.8, 26.5, 16.2, 14.5, 13.2, 11.9. **HRMS ESI(+)** (MeOH) calculated for $C_{58}H_{69}ClN_{10}O_9S_2Na^+$ $[M+Na]^+$: 1171.42711, found: 1171.42711.

#### 1.2.3.7. Esterification of MZ3

The esterification procedures were adapted from the procedure described by Liu *et al.*[14] with the specified modifications.

#### 1.2.3.8. Preparation of MZ3-AspA ester (21) (IF403)

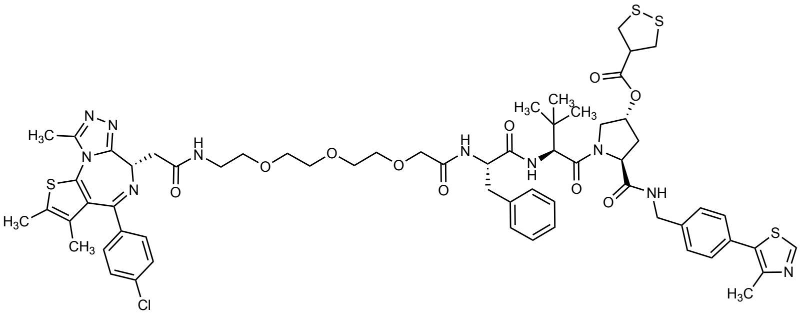

MZ3 (**2**, 19.6 mg, 17.0 µmol, 1.0 equiv), freshly prepared asparagusic acid (crude product, estimated purity: approx. 80%, 14.3 mg, 76.2 µmol, 4.5 equiv.), and DCC (11.8 mg, 57.1 µmol, 3.4 equiv.), were suspended in anhydrous $CH_2Cl_2$ (0.7 mL). To the suspension was added a solution of DMAP (4.25 mg, 34.8 µmol, 2.0 equiv.) in anhydrous $CH_2Cl_2$ (0.5 mL), followed by addition of $NEt_3$ (24.4 µL, 174 µmol, 10.2 equiv.). The reaction mixture was stirred at 25 °C. After 29 h, anhydrous $CH_2Cl_2$ (1.3 mL) was added to adjust for evaporation of the solvent. After 48 h, a second batch of reagents was added as follows: DCC (10.6 mg, 51.1 µmol, 3.0 equiv.) was added, followed by addition of a suspension of asparagusic acid (crude product, estimated purity: approx. 80%, 25.6 mg, 136 µmol, 8.0 equiv.) in anhydrous $CH_2Cl_2$ (0.5 mL), and $NEt_3$ (24.4 µL, 174 µmol, 10.2 equiv.). After 4 days, the solvent was removed *in vacuo* to provide the crude material. The crude material was resuspended in MeCN, filtered over a Discovery DSC-18 SPE cartridge (50 mg) and concentrated to provide the pre-purified material for further purification. Further purification was performed by prep RP-HPLC using a gradient of 50 to 70% B (MeCN+0.1% FA) over 60 min (LC time program (time - %B): 0 min - 50%, 5 min - 50%, 65 min – 70%, 95 min – 70%, 96 min – 100%, 106 min – 100%). The product containing fractions were combined and concentrated *in vacuo* at 44 °C water bath temperature, and further dried *in vacuo* to provide the product MZ3-AspA (**21**) ($R_t$: approx. 27.7 min) as a colorless solid in a yield of 21% (4.7 mg, 3.6 µmol). *Note: The purity of the crude AspA used is difficult to determine, as polymeric material does not, or only poorly dissolve in NMR solvent (turbid mixture). The estimated purity of 80% is thus, just a rough approximation. To account for the poor purity of the crude AspA mixture it was used in excess.*

#### 1.2.3.9. Characterization of MZ3-AspA ester (21) (IF403)

**$^1$H NMR** (500 MHz, $CDCl_3$) δ (ppm) 8.67 (s, 1H), 7.61 (t, *J* = 5.9 Hz, 1H), 7.41 – 7.36 (m, 3H), 7.35 – 7.28 (m, 6H), 7.25 – 7.18 (m, 5H), 7.13 (t, *J* = 5.6 Hz, 1H), 7.01 (d, *J* = 8.2 Hz, 1H), 5.42 – 5.37 (m, 1H), 4.81 (t, *J* = 7.7 Hz, 1H), 4.73 – 4.67 (m, 1H), 4.65 (t, *J* = 6.9 Hz, 1H), 4.52 (dd, *J* = 14.9, 6.4 Hz, 1H), 4.40 (d, *J* = 8.3 Hz, 1H), 4.35 (dd, *J* = 14.9, 5.4 Hz, 1H), 4.18 (d, *J* = 11.9 Hz, 1H), 4.01 (s, 2H), 3.76 (dd, *J* = 11.8, 4.2 Hz, 1H), 3.65 – 3.31 (m, 17H (*Note: observed integral: 19; should correspond to 17 Hs*)), 3.28 – 3.17 (m, 3H), 3.10 (dd, *J* = 14.3, 8.6 Hz, 1H), 2.75 (ddd, *J* = 13.3, 7.4, 5.5 Hz, 1H), 2.62 (s, 3H), 2.50 (s, 3H), 2.39 (s, 3H), 2.31 – 2.23 (m, 1H), 1.67 (s, 3H), 0.88 (s, 9H). **$^{13}$C NMR** (126 MHz, $CDCl_3$) δ (ppm) 171.7, 171.5, 171.1, 171.0, 170.9, 170.3, 163.8, 156. 0, 150.4, 149.9, 148.7, 138.2, 137.1, 136.9, 136.9, 132.3, 131.8, 131.2, 131.1, 130.9, 130.8, 130.0, 129.7, 129.5, 128.8, 128.7, 128.4, 127.0, 74.2, 70.9, 70.7, 70.7, 70.5, 70.5, 70.1, 58.5, 58.1, 54.6, 54.5, 54.0, 50.8, 43.5, 41.5, 41.4, 39.7, 39.1, 36.8, 35.2, 33.0, 26.5, 16.2, 14.5, 13.2, 11.9. **HRMS ESI(+)** (MeOH) calculated for $C_{62}H_{73}ClN_{10}O_{10}S_4Na^+$ $[M+Na]^+$: 1303.39747, found: 1303.39677. **FT-IR** ($CDCl_3$): $\nu$ ($cm^{-1}$) 3321m, 3064w, 2925w, 2870w, 1736m, 1653s, 1592m, 1529s, 1487m, 1443m, 1419m, 1370w, 1347w, 1270w, 1236w, 1176m, 1112m, 1091m, 1061w, 1032w, 1015w, 920w, 844w, 805w, 732m, 703w, 645w, 580w, 550w, 487w.

#### 1.2.3.10. Preparation of MZ3-All-C-AspA ester (22) (IF404)

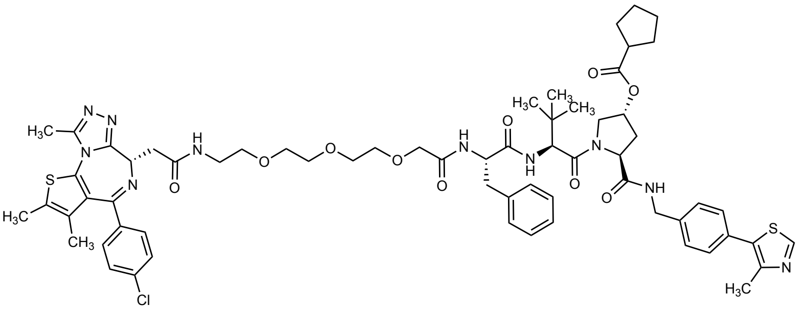

MZ3 (**2**, 21.5 mg, 18.7 µmol, 1.0 equiv) was suspended in anhydrous $CH_2Cl_2$ (0.8 mL). To the suspension was added $NEt_3$ (26.3 µL,187 µmol, 10.0 equiv.), DIC (8.7 µL, 56.1 µmol, 3.0 equiv.), and a solution of DMAP (4.57 mg, 37.4 µmol, 2.0 equiv.) in anhydrous $CH_2Cl_2$ (0.5 mL). To the mixture was then added cyclopentanecarboxylic acid (6.1 µL, 56.1 µmol, 3.0 equiv.) The reaction mixture was stirred at 25 °C. After 28 h, anhydrous $CH_2Cl_2$ (1.3 mL) was added to adjust for evaporation of the solvent, and then, a second batch of cyclopentanecarboxylic acid (6.1 µL, 56.1 µmol, 3.0 equiv.), and DIC (8.7 µL, 56.1 µmol, 3.0 equiv.) was added. After 64 h, the solvent was removed *in vacuo* to provide the crude material. The crude material was resuspended in MeCN, filtered over a Discovery DSC-18 SPE cartridge (50 mg) and concentrated to provide the pre-purified material for further purification. Further purification was performed by prep RP-HPLC using a gradient of 50 to 75% B (MeCN+0.1% FA) over 60 min (LC time program (time - %B): 0 min - 50%, 5 min - 50%, 65 min – 75%, 95 min – 75%, 96 min – 100%, 106 min – 100%). The product containing fractions were combined and concentrated *in vacuo* at 44 °C water bath temperature and further dried *in vacuo* to provide the product MZ3-All-C-AspA (**22**) ($R_t$: approx. 27.9 min) as a colorless solid in a yield of 53% (12.3 mg, 9.9 µmol).

#### 1.2.3.11. Characterization of MZ3-All-C-AspA ester (22) (IF404)

**$^1$H NMR** (500 MHz, $CDCl_3$) δ (ppm) 8.67 (s, 1H), 7.51 (t, *J* = 5.9 Hz, 1H), 7.42 – 7.36 (m, 3H), 7.35 – 7.27 (m, 6H), 7.25 – 7.16 (m, 5H), 7.10 (d, *J* = 6.0 Hz, 1H), 6.91 (d, *J* = 8.7 Hz, 1H), 5.42 – 5.35 (m, 1H), 4.77 (dd, *J* = 8.2, 6.3 Hz, 1H), 4.71 – 4.62 (m, 2H), 4.54 – 4.45 (m, 2H), 4.35 (dd, *J* = 14.9, 5.5 Hz, 1H), 4.01 – 3.97 (m, 2H), 3.98 – 3.89 (m, 1H), 3.85 (dd, *J* = 11.4, 5.0 Hz, 1H), 3.67 – 3.48 (m, 12H (*Note: observed integral: 13; should correspond to 12 Hs*)), 3.47 – 3.34 (m, 2H), 3.17 (dd, *J* = 14.1, 6.1 Hz, 1H), 3.10 (dd, *J* = 14.1, 8.6 Hz, 1H), 2.73 – 2.64 (m, 2H), 2.62 (s, 3H), 2.51 (s, 3H), 2.39 (s, 3H), 2.25 – 2.16 (m, 1H), 1.89 – 1.68 (m, 6H), 1.66 (s, 3H), 1.62 – 1.50 (m, 2H), 0.87 (s, 9H). **$^{13}$C NMR** (126 MHz, $CDCl_3$) δ (ppm) 176.5, 171.2, 170.9, 170.8, 170.6, 170.5, 163.8, 155.9, 150.4, 149.9, 148.7, 138.2, 137.1, 136.9, 136.8, 132.3, 131.8, 131.1, 131.1, 130.9, 130.7, 130.0, 129.6, 129.4, 128.8, 128.7, 128.4, 127.0, 72.6, 70.9, 70.7, 70.6, 70.5, 70.4, 70.1, 58.7, 57.6, 54.5, 54.5, 53.8, 43.7, 43.4, 39.6, 39.1, 36.9, 35.8, 33.1, 30.2, 30.0, 26.5, 26.0, 26.0, 16.2, 14.6, 13.2, 11.9. **HRMS ESI(+)** (MeOH) calculated for $C_{64}H_{77}ClN_{10}O_{10}S_2Na^+$ $[M+Na]^+$: 1267.48463, found:1267.48612. **Specific Rotation** $[\alpha]_D^{23°C}$ = +4.97 (c = 0.85, MeOH). **Specific Rotation** $[\alpha]_D^{23°C}$ = +12.5 (c = 0.55, MeOH). **FT-IR** ($CDCl_3$): $\nu$ ($cm^{-1}$) 3315w, 3064w, 2955m, 2926m, 2871w, 1732m, 1658s, 1588w, 1531s, 1488m, 1437m, 1419m, 1376w, 1351w, 1311w, 1267w, 1231w, 1184m, 1147m, 1110m, 1091m, 1034w, 1015w, 845w, 805w, 732w, 701w, 645w, 606w, 567w.

#### 1.2.3.12. Preparation of MZ3-LipA ester (23) (IF398)

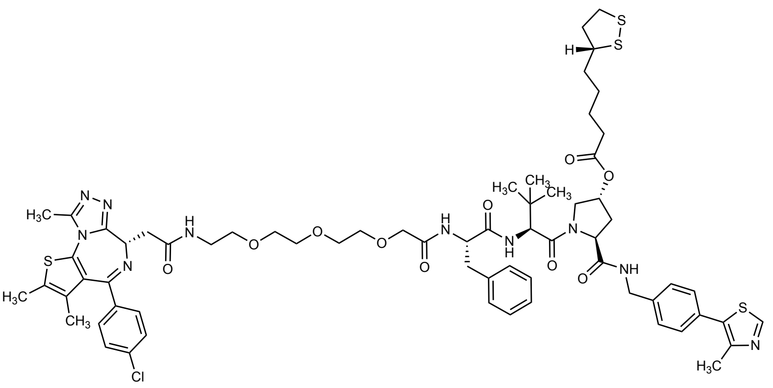

A stock solution of DMAP was prepared by dissolving 3.19 mg DMAP in anhydrous $CH_2Cl_2$ (1 mL). MZ3 (**2**, 30.0 mg, 26.1 µmol, 1.0 equiv.), *R*-lipoic acid (13.5 mg, 65.2 µmol, 2.5 equiv.), and DCC (16.1 mg, 78.3 µmol, 3.0 equiv.), were suspended in anhydrous $CH_2Cl_2$ (1.6 mL). To the mixture was added $NEt_3$ (25.7 µL, 183 µmol, 7.0 equiv.), followed by addition of 200 µL of the stock-solution of DMAP (effective amount added: 0.638 mg, 5.22 µmol, 0.2 equiv.). The reaction mixture was stirred at 25 °C. After 50 h, a second batch of reagents was added as follows: To the mixture was added $NEt_3$ (25.7 µL, 183 µmol, 7.0 equiv.), followed by addition of a solution of DMAP (3.19 mg, 26.1 µmol, 1.0 equiv.) in 300 µL anhydrous $CH_2Cl_2$. Then was added DCC (16.1 mg, 78.3 µmol, 3.0 equiv.), and *R*-lipoic acid (13.5 mg, 65.2 µmol, 2.5 equiv.). After 6 days, the solvent was removed *in vacuo* to provide the crude material. The crude material was resuspended in MeCN, filtered over a Discovery DSC-18 SPE cartridge (50 mg) and concentrated to provide the pre-purified material for further purification. Further purification was performed by prep RP-HPLC using a gradient of 50 to 75% B (MeCN+0.1% FA) over 60 min (LC time program (time - %B): 0 min - 50%, 5 min - 50%, 65 min – 75%, 95 min – 75%, 96 min – 100%, 106 min – 100%). The product containing fractions were combined and concentrated *in vacuo* at 44 °C water bath temperature and further dried *in vacuo* to provide the product MZ3-LipA (**23**) ($R_t$: approx. 36.4 min) as a colorless solid in a yield of 47% (16.4 mg, 12.2 µmol).

#### 1.2.3.13. Characterization of MZ3-LipA ester (23) (IF398)

**$^1$H NMR** (500 MHz, $CDCl_3$) δ (ppm) 8.67 (s, 1H), 7.55 (t, *J* = 5.9 Hz, 1H), 7.42 (d, *J* = 7.9 Hz, 1H), 7.44 – 7.15 (m, 14H (*partially overlaps with residual solvent peak*)), 6.95 (d, *J* = 8.6 Hz, 1H), 5.42 – 5.37 (m, 1H), 4.79 (dd, *J* = 8.2, 6.5 Hz, 1H), 4.70 – 4.62 (m, 2H), 4.50 (dd, *J* = 14.9, 6.4 Hz, 1H), 4.45 (d, *J* = 8.6 Hz, 1H), 4.35 (dd, *J* = 14.9, 5.5 Hz, 1H), 4.04 – 3.95 (m, 3H), 3.84 (dd, *J* = 11.5, 4.9 Hz, 1H), 3.66 – 3.49 (m, 13H (*Note: observed integral: 14; should correspond to 13 Hs*)), 3.44 – 3.35 (m, 2H), 3.23 – 3.04 (m, 4H), 2.74 – 2.65 (m, 1H), 2.61 (s, 3H), 2.50 (s, 3H), 2.48 – 2.41 (m, 1H), 2.39 (s, 3H), 2.37 – 2.17 (m, 3H), 1.93 – 1.83 (m, 1H), 1.66 (s, 3H), 1.72 – 1.58 (m, 4H (*Note: integral observed: 9; should correspond to 4 Hs*)), 1.52 – 1.37 (m, 2H), 0.87 (s, 9H). **$^{13}$C NMR** (126 MHz, $CDCl_3$) δ (ppm) 173.2, 171.1, 171.0, 170.8, 170.8, 170.5, 163.9, 155.9, 150.4, 149.9, 148.6, 138.2, 137.1, 136.8, 136.8, 132.3, 131.8, 131.1, 131.1, 130.9, 130.7, 130.0, 129.6, 129.4, 128.8, 128.7, 128.4, 127.0, 72.8, 70.9, 70.6, 70.6, 70.5, 70.4, 70.1, 58.6, 57.7, 56.5, 54.5, 54.5, 53.8, 43.4, 40.4, 39.6, 39.1, 38.6, 36.8, 35.6, 34.8, 34.0, 33.2, 28.9, 26.5, 24.6, 16.2, 14.6, 13.2, 11.9. **HRMS ESI(+)** (MeOH) calculated for $C_{66}H_{81}ClN_{10}O_{10}S_4Na^+$ $[M+Na]^+$: 1359.46007, found: 1359.45858. **Specific Rotation** $[\alpha]_D^{24°C}$ = +30.96 (c = 0.53, MeOH). **FT-IR** ($CDCl_3$): $\nu$ (cm$^{-1}$) 3312m, 3065w, 2926m, 2869w, 1735m, 1655s, 1592m, 1531s, 1488m, 1435m, 1419m, 1375w, 1351w, 1317w, 1268w, 1232m, 1174m, 1110m, 1091m, 1033w, 1014w, 916w, 844w, 805w, 731m, 702w, 645w, 605w, 571w, 550w.

#### 1.2.3.14. Preparation of MZ3-All-C-LipA ester (24) (IF399)

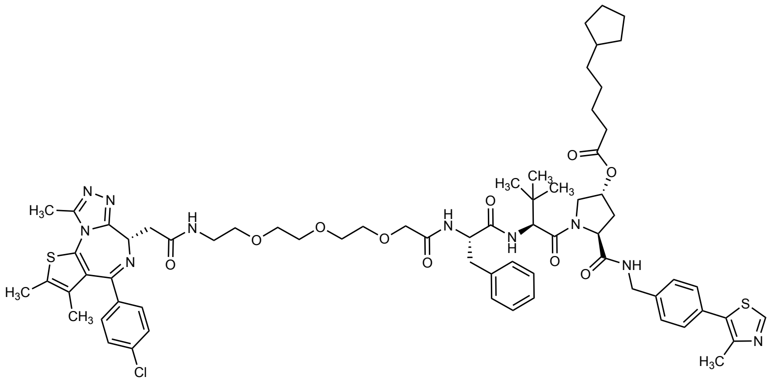

A stock solution of DMAP was prepared by dissolving 3.19 mg DMAP in anhydrous $CH_2Cl_2$ (1 mL). MZ3 (**2**, 30.0 mg, 26.1 µmol, 1.0 equiv.), 5-cyclopentylpentanoic acid (prepared by Dr. Inga S. Shchelik; 11.1 mg, 65.2 µmol, 2.5 equiv.), and DCC (16.6 mg, 80.6 µmol, 3.1 equiv.) were suspended in anhydrous $CH_2Cl_2$ (1.6 mL). To the mixture was added $NEt_3$ (25.7 µL, 183 µmol, 7.0 equiv.), followed by addition of 200 µL of the stock-solution of DMAP (effective amount added: 0.638 mg, 5.22 µmol, 0.2 equiv.). The reaction mixture was stirred at 25 °C. After 48 h, a second batch of reagents was added as follows: To the mixture was added $NEt_3$ (25.7 µL, 183 µmol, 7.0 equiv.), followed by addition of a solution of DMAP (3.19 mg, 26.1 µmol, 1.0 equiv.) in 300 µL anhydrous $CH_2Cl_2$. Then was added DCC (16.1 mg, 78.3 µmol, 3.0 equiv.), and a solution of 5-cyclopentylpentanoic acid (prepared by Dr. Inga S. Shchelik; 13.5 mg, 79.6 µmol, 3.1 equiv.) in 200 µL anhydrous $CH_2Cl_2$. After 68 h, the solvent was removed *in vacuo* to provide the crude material. The crude material was resuspended in MeCN, filtered over a Discovery DSC-18 SPE cartridge (50 mg) and concentrated to provide the pre-purified material for further purification *via* prep RP-HPLC. The crude was split on two runs for the purification by prep RP-HPLC using a gradient of 50 to 95% B (MeCN+0.1% FA) over 60 min (LC time program (time - %B): 0 min - 50%, 5 min - 50%, 65 min – 95%, 95 min – 95%, 96 min – 100%, 106 min – 100%). The product containing fractions were combined and concentrated *in vacuo* at 44 °C water bath temperature and further dried *in vacuo* to provide the product MZ3-All-C-LipA (**24**) ($R_t$: 39.3 min) as a colorless solid in a yield of 64% (21.7 mg, 16.7 µmol).

#### 1.2.3.15. Characterization of MZ3-All-C-LipA ester (24) (IF399)

**$^1$H NMR** (500 MHz, $CDCl_3$) δ (ppm) 8.67 (s, 1H), 7.54 (t, *J* = 6.0 Hz, 1H), 7.41 (d, *J* = 8.0 Hz, 1H), 7.39 – 7.22 (m, 10H), 7.21 – 7.16 (m, 3H), 7.15 – 7.11 (m, 1H), 6.93 (d, *J* = 8.6 Hz, 1H), 5.41 – 5.36 (m, 1H), 4.78 (dd, *J* = 8.2, 6.3 Hz, 1H), 4.70 – 4.62 (m, 2H), 4.50 (dd, *J* = 14.9, 6.4 Hz, 1H), 4.45 (d, *J* = 8.6 Hz, 1H), 4.35 (dd, *J* = 14.9, 5.5 Hz, 1H), 4.01 – 3.98 (m, 2H), 3.98 – 3.93 (m, 1H), 3.85 (dd, *J* = 11.4, 5.1 Hz, 1H), 3.66 – 3.49 (m, 12H), 3.44 – 3.34 (m, 2H), 3.18 (dd, *J* = 14.1, 6.1 Hz, 1H), 3.10 (dd, *J* = 14.2, 8.7 Hz, 1H), 2.71 (dt, *J* = 13.5, 6.0 Hz, 1H), 2.62 (s, 3H), 2.50 (s, 3H), 2.39 (s, 3H), 2.34 – 2.24 (m, 2H), 2.24 – 2.16 (m, 1H), 1.75 – 1.67 (m, 3H (*Note: observed integral: 4; supposably correspond to 3 Hs)),* 1.66 (s, 3H), 1.61 – 1.53 (m, 4H), 1.52 – 1.43 (m, 2H), 1.34 – 1.24 (m, 4H (*Note: observed integral: 5; supposably correspond to 4 Hs*)), 1.06 – 0.99 (m, 2H), 0.86 (s, 9H). **$^{13}$C NMR** (126 MHz, $CDCl_3$) δ (ppm) 173.6, 171.2, 170.9, 170.8, 170.7, 170.5, 163.8, 155.9, 150.4, 149.9, 148.6, 138.2, 137.1, 136.8, 136.8, 132.3, 131.8, 131.1, 131.1, 130.9, 130.7, 130.0, 129.6, 129.4, 128.8, 128.7, 128.4, 127.0, 72.7, 70.9, 70.7, 70.6, 70.5, 70.4, 70.1, 58.6, 57.6, 54.5, 54.5, 53.7, 43.4, 40.1, 39.6, 39.1, 36.8, 35.9, 35.6, 34.3, 33.1, 32.8, 28.5, 26.5, 25.3, 25.1, 16.2, 14.6, 13.2, 11.9. **HRMS ESI(+)** (MeOH) calculated for $C_{68}H_{85}ClN_{10}O_{10}S_2Na^+$ $[M+Na]^+$: 1323.54723, found: 1323.54717. **Specific Rotation** $[\alpha]_D^{23°C}$ = +9.24 (c = 0.70, MeOH). **FT-IR** ($CDCl_3$): $\nu$ ($cm^{-1}$) 3314m, 3066w, 2936m, 2866m, 1736m, 1656s, 1592m, 1532s, 1488m, 1436m, 1418m, 1370w, 1351w, 1271w, 1231m, 1097m, 1033w, 1015w, 845w, 805w, 732w, 701w, 606w.

#### 1.2.4. *Preparation of AspA*

The crude AspA was prepared in 3 steps from 3-bromo-2-(bromomethyl)propanoic acid as previously reported by Tirla *et al.*[16]

##### 1.2.4.1. Preparation of 3-(acetylthio)-2-((acetylthio)methyl)propanoic acid[16] (IF379)

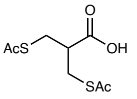

3-(Acetylthio)-2-((acetylthio)methyl)propanoic acid was prepared after Tirla *et al.*[16] with minor modifications. 3-Bromo-2-(bromomethyl)propanoic acid (1.50 g, 6.10 mmol, 1.0 equiv.) was dissolved in 1 M aqueous NaOH (6.1 mL), followed by addition of potassium thioacetate (1.79 g, 15.4 mmol, 2.5 equiv.), and milliQ water (12 mL). The reaction was stirred at 25 °C. After 24 h, the reaction mixture was acidified to a pH of approx. 2-3 by slow addition of 1 M aqueous HCl (approx. 8 mL) under vigorous stirring. The reaction mixture is then extracted with EtOAc (3 × 25 mL), and the combined organic phase washed with acidified brine (pH 1, 2 × 75 mL), dried over anhydrous $MgSO_4$, filtered and concentrated *in vacuo* to provide the crude material as an orange oil in a crude yield of approx. ≤ 91% (1.31 g) in a purity of approx. ≤ 94% (based on $^1H$ NMR analysis by comparing the integrals of the $^1H$ signal of the product and the impurity peaks in the vinylic region). The crude material was used in the next step without further purification.

##### 1.2.4.2. Characterization of 3-(acetylthio)-2-((acetylthio)methyl)propanoic acid[16] (IF379)

**$^1$H NMR** (400 MHz, $CDCl_3$) δ (ppm) 3.26 – 3.12 (m, 4H), 2.97 – 2.86 (m, 1H), 2.35 (s, 6H). The $^1H$ NMR data is in good agreement with literature.[16] **$^{13}$C NMR** (101 MHz, $CDCl_3$) δ (ppm) 195.1, 177.9, 45.4, 30.7, 29.4. **HRMS ESI(–)** (MeOH) calculated for $C_8H_{11}O_4S_2^-$ [M-H$^+$]$^-$: 235.01042, found: 235.01030.

##### 1.2.4.3. Preparation of 3-mercapto-2-(mercaptomethyl) propanoic acid[16] (IF385)

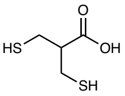

3-Mercapto-2-(mercaptomethyl)propanoic acid was prepared after Tirla *et al.*[16] with minor modifications. The crude 3-(acetylthio)-2-((acetylthio)methyl)propanoic acid (≤94% purity, 200 mg, 0.80 mmol, 1.0 equiv.) was dissolved in 1 M aqueous NaOH (1.5 mL). The solution was cooled to 0 °C, before addition of NaOH pellets (freshly pestled) (102 mg, 2.55 mmol, 3.2 equiv.). After the pellets have dissolved, the solution was allowed to warm to 25 °C and stirred overnight. After 24 h, the reaction mixture was cooled to 0 °C and ice was added to the reaction mixture, before it was acidified with 16% aqueous HCl to a pH of 0-1 (approx. 0.7 mL used). A white emulsion formed that was extracted with EtOAc (3 × 10 mL). The combined organic layer was washed with acidified brine (pH ~ 1, 2 × 30 mL), dried over anhydrous $MgSO_4$, filtered, and concentrated *in vacuo* to provide the crude product as a yellow oil (118 mg). The crude material was used directly in the next step without further purification. *Note: Significant amounts of other species, and residual EtOAc are present in the $^1H$ NMR spectrum.*

#### 1.2.4.4. Preparation of AspA[16] (IF392)

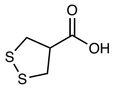

AspA was prepared after Tirla *et al.*[16] with minor modifications. The crude 3-mercapto-2-(mercaptomethyl)propanoic acid (118 mg) was dissolved in anhydrous DMSO (6.5 mL) and stirred at 75 °C overnight in an open flask. After 15 h the reaction mixture was allowed to cool to 25 °C, and then acidified with 1 M aqueous HCl until pH 2 (~1.3 mL used). The mixture was then extracted with EtOAc (4 × 5 mL), and the combined organic layer washed with acidified brine (pH 1, 2 × 20 mL), dried over anhydrous $MgSO_4$, filtered and concentrated *in vacuo* at 40°C water bath. The crude product was obtained as a yellow oil in a crude yield of approx. < 70% (81.5 mg) over 2 steps in an estimated purity of approx. < 70%. *Note: The purity of the product is difficult to determine based on the formation of polymerized material that is poorly soluble in the NMR solvent. For reactions using AspA, the purity was estimated with about ≤ 70% as indicated in the individual experiments. To avoid substantial polymerization and for operational simplicity, the crude AspA was not concentrated to dryness, but used in the next step (acylation reactions of MZ1, or MZ3) without any further purification and with residual amounts of solvents (DMSO, EtOAc) present. Therefore, the actual yield is lower.*

#### 1.2.4.5. Characterization of AspA[16] (IF392)

**$^1$H NMR** (400 MHz, $CDCl_3$) δ (ppm) 3.56 – 3.42 (m, 3H), 3.37 – 3.24 (m, 2H). The $^1$H NMR data is in overall agreement with literature.[16] **$^{13}$C NMR** (101 MHz, $CDCl_3$) δ (ppm) 176.7, 50.5, 41.5.

#### 1.3. Preliminary cell viability studies *via* MTT assays

Cell viability was studied using a 3-(4,5-dimethylthiazol-2-yl)-2,5-diphenyltetrazolium bromide (MTT) cytotoxicity assay in HeLa cells.

10 mM DMSO stock solutions of inhibitors were prepared freshly the day of the assay or prepared, stored at -20°C, and used within 2 weeks. All assay materials, including cells, reactants, reagents, and consumables were provided by Prof. Dr. Rivera-Fuentes (Department of Chemistry, University of Zurich). Assays were conducted following the procedures established in the group of Prof. Dr. Rivera-Fuentes and under the supervision of Dr. Annabell Martin (Department of Chemistry, University of Zurich), and Dr. Juan Tamez Fernandez (Department of Chemistry, University of Zurich).

**Cell culture:**
HeLa cells were grown analogous to a previously published protocol by Martin and Rivera-Fuentes.[17] HeLa cells (CLS, 300194) were grown in T75 flasks for cell culture (TC flask T75, Stand., Vent. Cap; Sarstedt AG&Co. KG, Cat No. 83.3911.002) in Dulbecco's Modified Eagle Medium (DMEM) supplemented with foetal bovine serum (FBS) (10%) and penicillin-streptomycin (1%) (Pen Strep, gibco™, Cat. No.: 15070-063) at 37 °C in an atmosphere of 95% humidity and 5% $CO_2$. Cells were grown to a confluency of approx. 90%, before medium was removed, and the cells washed with phosphate buffered saline (PBS, pH 7.4, gibco, Cat. No.: 10010-015) (2 × 15 mL), detached with trypsin (1.5 mL) (0.25 % Trypsin-EDTA (1x), gibco™, Cat. No.: 25200-056) for 3 min at 37 °C (atmosphere: 95% humidity, 5% $CO_2$), and taken up in DMEM again (9 mL, 10% FBS, 1% penicillin-streptomycin). Cells were then counted using an EVE™ automatic cell counter (NanoEnTek) and seeded in a 96 well plate at a density of $4 \times 10^3$ cells per well in a volume of 100 µL per well (see MTT assay).

**MTT assay:**
$4 \times 10^3$ HeLa cells in 100 µL DMEM medium (10% FBS, 1% penicillin-streptomycin) were dispensed in a 96-well flat bottom cell-culture plate (Greiner Bio-One, Cellstar®, Cat No. 655180; 96 well cell culture plate, sterile, F-bottom, with lid) from B2 to G12. The cells were incubated at 37°C (atmosphere: 95% humidity, 5% $CO_2$) for 24 h.

10 Sequential 2-fold serial dilutions of inhibitor spanning a range of 10 mM to 0.0195 mM were prepared in DMSO (5 µL of each concentration), and diluted with 95 µL DMEM (10% FBS, 1% penicillin-streptomycin). 10 µL of each of the inhibitor serial dilutions was added in triplicate to the cells grown for 24 h (Inhibitor concentration range upon incubation: 45.45 µM to 0.0888 µM; total volume: 110 µL; DMSO concentration: 0.45% (v/v)). As a growth control cells in the column B2 to G2 were treated with DMSO only.

After 72-73 h, the cell viability was assessed using an MTT assay. 10 µL of MTT solution (0.5% (w/v) in FluoroBrite™ DMEM, gibco™, Cat No. A18967-01) was added to each well and the cells incubated for 3 h at 37°C (atmosphere: 95% humidity, 5% $CO_2$) before the medium was removed carefully. The precipitate was dissolved in isopropanol (100 µL per well) and the plates shaken for 30 min to ensure homogeneity. Absorbance was then measured at 550 nm using a UV/VIS microplate reader (Multiskan Sky Microplate Spectrophotometer, Thermoscientific). Values obtained from wells with isopropanol only (wells A2-G2) were subtracted as background, and the absorbance of growth control wells containing no inhibitor was set to 100% cell viability.

Results were analyzed and visualized in GraphPad Prism® (Prism 10 for Windows 64-bit) version 10.3.1 (509)[18] and version 10.4.1 (627)[19].The data was normalized in Prism (each subcolumn separately) setting 0% as 0 and 100% as the largest value in each subcolumn. Normalized data was then fitted to a nonlinear regression curve using the sigmoidal curve fit (sigmoidal, 4PL, X is concentration) without any constraints in Prism 10[18,19]. Experiments were carried out in biological triplicate on different days with each biological replicate being performed in 3 technical replicates. Outliers due to edge effects in the 96 well plate were excluded after visual inspection resulting in 2–3 technical replicates per biological replicate that were used for data processing (Raw data used for data analysis is provided in the tables below (see section 1.3.1)). $IC_{50}$ values were not determined as in many cases no top- and/or bottom plateau was reached within the concentration range tested. Further for the MZ3-series effects of limited solubility at the higher concentration ranges tested cannot be excluded and thus, also here no IC50 values are provided herein.

#### *1.3.1. Raw data MTT assays*

*Table 1: Raw data of the absorbance at 550 nm after blank (isopropanol) subtraction of the biological and technical replicates of the MTT assay with MZ1 used for further data analysis.*

| MZ1 (IF375) | | | | | | | | | |
|---|---|---|---|---|---|---|---|---|---|
| Inhibitor concentration [µM] | Absorbance 550 nm Biological replicate 1 | | | Absorbance 550 nm Biological replicate 2 | | | Absorbance 550 nm Biological replicate 3 | | |
| 0 | 0.6639 | 0.7474 | 0.7314 | 0.8612 | 0.8339 | 0.9408 | –[a] | 0.5448 | 0.6841 |
| 45.45 | 0.1761 | 0.3157 | 0.272 | 0.05060 | 0.05100 | 0.06220 | –[a] | 0.02460 | 0.01730 |
| 22.73 | 0.2528 | 0.3913 | 0.3173 | 0.1058 | 0.1066 | 0.1102 | –[a] | 0.03510 | 0.05150 |
| 11.36 | 0.4313 | 0.3371 | 0.2779 | 0.1304 | 0.1584 | 0.1416 | –[a] | 0.06740 | 0.06660 |
| 5.68 | 0.4186 | 0.5625 | 0.444 | 0.1863 | 0.2482 | 0.2193 | –[a] | 0.09000 | 0.09770 |
| 2.84 | 0.5508 | 0.5255 | 0.5266 | 0.2377 | 0.2982 | 0.2820 | –[a] | 0.1298 | 0.1372 |
| 1.42 | 0.6146 | 0.5805 | 0.5906 | 0.2849 | 0.3811 | 0.3463 | –[a] | 0.2030 | 0.1678 |
| 0.71 | 0.593 | 0.5959 | 0.6194 | 0.4592 | 0.5163 | 0.4955 | –[a] | 0.2302 | 0.2012 |
| 0.36 | 0.6805 | 0.6961 | 0.6402 | 0.5703 | 0.6580 | 0.5989 | –[a] | 0.2939 | 0.3041 |
| 0.18 | 0.6925 | 0.7097 | 0.7209 | 0.6350 | 0.7262 | 0.7490 | –[a] | 0.4525 | 0.4285 |
| 0.09 | 0.7018 | 0.7742 | 0.8003 | 0.6714 | 0.6804 | 0.7385 | –[a] | 0.4522 | 0.4693 |

[a]*Values were excluded for further data analysis as there were edge effects observed in the 96 well plate by visual inspection.*

*Table 2: Raw data of the absorbance at 550 nm after blank (isopropanol) subtraction of the biological and technical replicates of the MTT assay with MZ1-AspA used for further data analysis.*

| MZ1-AspA (IF393) | | | | | | | | | |
|---|---|---|---|---|---|---|---|---|---|
| Inhibitor concentration [µM] | Absorbance 550 nm Biological replicate 1 | | | Absorbance 550 nm Biological replicate 2 | | | Absorbance 550 nm Biological replicate 3 | | |
| 0 | –[a] | 0.5909 | 0.3959 | 0.6541 | 0.8001 | 0.8964 | –[a] | 0.4258 | 0.5181 |
| 45.45 | –[a] | 0.0128 | 0.0171 | 0.01200 | 0.01630 | 0.01970 | –[a] | 0.01410 | 0.009400 |
| 22.73 | –[a] | 0.0241 | 0.0262 | 0.02930 | 0.02180 | 0.01660 | –[a] | 0.01110 | 0.02130 |
| 11.36 | –[a] | 0.0582 | 0.0666 | 0.06010 | 0.04800 | 0.06230 | –[a] | 0.01480 | 0.03480 |
| 5.68 | –[a] | 0.2332 | 0.2371 | 0.1374 | 0.1663 | 0.1803 | –[a] | 0.03970 | 0.04070 |
| 2.84 | –[a] | 0.276 | 0.2997 | 0.2521 | 0.2182 | 0.2691 | –[a] | 0.1013 | 0.1023 |
| 1.42 | –[a] | 0.3219 | 0.3696 | 0.3435 | 0.3026 | 0.3172 | –[a] | 0.1190 | 0.1048 |
| 0.71 | –[a] | 0.4028 | 0.4095 | 0.4793 | 0.4524 | 0.4771 | –[a] | 0.1725 | 0.1554 |
| 0.36 | –[a] | 0.4803 | 0.5271 | 0.5876 | 0.6073 | 0.5285 | –[a] | 0.3888 | 0.3634 |
| 0.18 | –[a] | 0.5709 | 0.5956 | 0.7079 | 0.6695 | 0.6834 | –[a] | 0.4881 | 0.5031 |
| 0.09 | –[a] | 0.5807 | 0.6054 | 0.7228 | 0.7213 | 0.7475 | –[a] | 0.6069 | 0.6126 |

[a]*Values were excluded for further data analysis as there were edge effects observed in the 96 well plate by visual inspection.*

*Table 3: Raw data of the absorbance at 550 nm after blank (isopropanol) subtraction of the biological and technical replicates of the MTT assay with MZ1-All-C-AspA used for further data analysis.*

| MZ1-All-C-AspA (IF395) | | | | | | | | | |
|---|---|---|---|---|---|---|---|---|---|
| Inhibitor concentration [µM] | Absorbance 550 nm Biological replicate 1 | | | Absorbance 550 nm Biological replicate 2 | | | Absorbance 550 nm Biological replicate 3 | | |
| 0 | 0.577 | 0.5713 | –[a] | 0.8637 | 0.7882 | 0.7302 | 0.5468 | 0.4793 | –[a] |
| 45.45 | 0.0207 | 0.0481 | –[a] | 0.01740 | 0.005200 | 0.02270 | 0.01370 | 0.03910 | –[a] |
| 22.73 | 0.0367 | 0.0548 | –[a] | 0.02780 | 0.03500 | 0.03760 | 0.02540 | 0.03700 | –[a] |
| 11.36 | 0.1075 | 0.0979 | –[a] | 0.1117 | 0.07620 | 0.08800 | 0.05390 | 0.05280 | –[a] |
| 5.68 | 0.2379 | 0.2296 | –[a] | 0.1889 | 0.1730 | 0.1590 | 0.07170 | 0.07030 | –[a] |
| 2.84 | 0.3312 | 0.3329 | –[a] | 0.2708 | 0.2680 | 0.2667 | 0.1014 | 0.1013 | –[a] |
| 1.42 | 0.3947 | 0.3861 | –[a] | 0.3456 | 0.3654 | 0.3652 | 0.1624 | 0.1619 | –[a] |
| 0.71 | 0.4172 | 0.3885 | –[a] | 0.3924 | 0.3543 | 0.3600 | 0.2400 | 0.2323 | –[a] |
| 0.36 | 0.4588 | 0.4828 | –[a] | 0.5461 | 0.5229 | 0.5183 | 0.2810 | 0.2709 | –[a] |
| 0.18 | 0.5209 | 0.5255 | –[a] | 0.7557 | 0.6596 | 0.6634 | 0.4903 | 0.4619 | –[a] |
| 0.09 | 0.5725 | 0.5526 | –[a] | 0.6553 | 0.5717 | 0.5951 | 0.4762 | 0.4445 | –[a] |

[a]*Values were excluded for further data analysis as there were edge effects observed in the 96 well plate by visual inspection.*

*Table 4: Raw data of the absorbance at 550 nm after blank (isopropanol) subtraction of the biological and technical replicates of the MTT assay with MZ1-LipA used for further data analysis.*

| MZ1-LipA (IF370) | | | | | | | | | |
|---|---|---|---|---|---|---|---|---|---|
| Inhibitor concentration [µM] | Absorbance 550 nm Biological replicate 1 | | | Absorbance 550 nm Biological replicate 2 | | | Absorbance 550 nm Biological replicate 3 | | |
| 0 | –[a] | 0.6632 | 0.6188 | 0.7747 | 0.7615 | 0.7407 | –[a] | 0.5130 | 0.6415 |
| 45.45 | –[a] | 0.0237 | 0.0289 | 0.04220 | 0.03710 | 0.03630 | –[a] | 0.04390 | 0.03590 |
| 22.73 | –[a] | 0.0195 | 0.0164 | 0.03730 | 0.05680 | 0.04960 | –[a] | 0.03060 | 0.02390 |
| 11.36 | –[a] | 0.0215 | 0.0197 | 0.05260 | 0.05520 | 0.03290 | –[a] | 0.03970 | 0.06790 |
| 5.68 | –[a] | 0.0546 | 0.0517 | 0.05610 | 0.06520 | 0.06790 | –[a] | 0.07770 | 0.05570 |
| 2.84 | –[a] | 0.1328 | 0.1476 | 0.09340 | 0.09740 | 0.09200 | –[a] | 0.1116 | 0.08720 |
| 1.42 | –[a] | 0.297 | 0.3094 | 0.2097 | 0.2044 | 0.2123 | –[a] | 0.1340 | 0.1309 |
| 0.71 | –[a] | 0.2953 | 0.3005 | 0.2583 | 0.2620 | 0.2586 | –[a] | 0.1796 | 0.2022 |
| 0.36 | –[a] | 0.3029 | 0.3783 | 0.3496 | 0.3339 | 0.3145 | –[a] | 0.2283 | 0.2263 |
| 0.18 | –[a] | 0.3854 | 0.3721 | 0.5654 | 0.5281 | 0.5139 | –[a] | 0.3309 | 0.3648 |
| 0.09 | –[a] | 0.4863 | 0.4833 | 0.5981 | 0.7185 | 0.6784 | –[a] | 0.3878 | 0.3886 |

[a]*Values were excluded for further data analysis as there were edge effects observed in the 96 well plate by visual inspection.*

*Table 5: Raw data of the absorbance at 550 nm after blank (isopropanol) subtraction of the biological and technical replicates of the MTT assay with MZ1-All-C-LipA used for further data analysis.*

| MZ1-All-C-LipA (IF378) | | | | | | | | | |
|---|---|---|---|---|---|---|---|---|---|
| Inhibitor concentration [µM] | Absorbance 550 nm Biological replicate 1 | | | Absorbance 550 nm Biological replicate 2 | | | Absorbance 550 nm Biological replicate 3 | | |
| 0 | 0.574 | 0.5676 | –[a] | 0.7262 | 0.7778 | 0.7007 | 0.6664 | 0.6509 | –[a] |
| 45.45 | 0.0149 | 0.0315 | –[a] | 0.02130 | 0.08830 | 0.06880 | 0.01990 | 0.03050 | –[a] |
| 22.73 | 0.0268 | 0.0207 | –[a] | 0.02420 | 0.04380 | 0.05390 | 0.02120 | 0.02950 | –[a] |
| 11.36 | 0.0268 | 0.0214 | –[a] | 0.03470 | 0.04510 | 0.05370 | 0.1087 | 0.06660 | –[a] |
| 5.68 | 0.0329 | 0.0409 | –[a] | 0.05330 | 0.06130 | 0.1054 | 0.07450 | 0.1294 | –[a] |
| 2.84 | 0.0442 | 0.0456 | –[a] | 0.08890 | 0.07360 | 0.06910 | 0.1027 | 0.1326 | –[a] |
| 1.42 | 0.1301 | 0.1066 | –[a] | 0.1530 | 0.1403 | 0.1414 | 0.1304 | 0.1666 | –[a] |
| 0.71 | 0.3207 | 0.2779 | –[a] | 0.2791 | 0.2344 | 0.2569 | 0.1812 | 0.2379 | –[a] |
| 0.36 | 0.295 | 0.2926 | –[a] | 0.3032 | 0.2637 | 0.3154 | 0.2620 | 0.2567 | –[a] |
| 0.18 | 0.3694 | 0.3346 | –[a] | 0.3886 | 0.4027 | 0.4234 | 0.5936 | 0.5117 | –[a] |
| 0.09 | 0.5312 | 0.4522 | –[a] | 0.5718 | 0.6092 | 0.4322 | 0.4888 | 0.4621 | –[a] |

[a]*Values were excluded for further data analysis as there were edge effects observed in the 96 well plate by visual inspection.*

*Table 6: Raw data of the absorbance at 550 nm after blank (isopropanol) subtraction of the biological and technical replicates of the MTT assay with MZ3 used for further data analysis.*

| MZ3 (IF397) | | | | | | | | | |
|---|---|---|---|---|---|---|---|---|---|
| Inhibitor concentration [µM] | Absorbance 550 nm Biological replicate 1 | | | Absorbance 550 nm Biological replicate 2 | | | Absorbance 550 nm Biological replicate 3 | | |
| 0 | 0.7089 | 0.6885 | 0.6289 | 0.5990 | 0.7146 | 0.9064 | 0.6361 | 0.6536 | –[a] |
| 45.45 | 0.5778 | 0.5766 | 0.5695 | 0.1907 | 0.2422 | 0.2465 | 0.1434 | 0.1359 | –[a] |
| 22.73 | 0.5964 | 0.6143 | 0.565 | 0.2223 | 0.2120 | 0.2378 | 0.1182 | 0.1281 | –[a] |
| 11.36 | 0.6852 | 0.5713 | 0.6107 | 0.2325 | 0.2471 | 0.2557 | 0.1786 | 0.1513 | –[a] |
| 5.68 | 0.6393 | 0.6705 | 0.6277 | 0.3377 | 0.2965 | 0.3707 | 0.1940 | 0.1596 | –[a] |
| 2.84 | 0.7046 | 0.7285 | 0.6907 | 0.4257 | 0.3963 | 0.4917 | 0.2358 | 0.2304 | –[a] |
| 1.42 | 0.7358 | 0.7417 | 0.7078 | 0.5163 | 0.5330 | 0.5995 | 0.4132 | 0.4004 | –[a] |
| 0.71 | 0.8041 | 0.767 | 0.7405 | 0.6364 | 0.7576 | 0.7826 | 0.5147 | 0.4977 | –[a] |
| 0.36 | 0.8072 | 0.7827 | 0.8076 | 0.7137 | 0.8532 | 0.7970 | 0.6288 | 0.5888 | –[a] |
| 0.18 | 1.054 | 0.9337 | 0.9381 | 0.7552 | 0.6756 | 0.8685 | 0.6746 | 0.6748 | –[a] |
| 0.09 | 0.8626 | 0.8363 | 0.8569 | 0.6207 | 0.8518 | 0.7478 | 0.6296 | 0.5566 | –[a] |

[a]*Values were excluded for further data analysis as there were edge effects observed in the 96 well plate by visual inspection.*

*Table 7: Raw data of the absorbance at 550 nm after blank (isopropanol) subtraction of the biological and technical replicates of the MTT assay with MZ3-AspA used for further data analysis.*

| MZ3-AspA (IF403) | | | | | | | | | |
|---|---|---|---|---|---|---|---|---|---|
| Inhibitor concentration [µM] | Absorbance 550 nm Biological replicate 1 | | | Absorbance 550 nm Biological replicate 2 | | | Absorbance 550 nm Biological replicate 3 | | |
| 0 | 0.7438 | 0.8379 | 0.8681 | 0.7490 | 0.8511 | 0.8528 | –[a] | 0.4149 | 0.5184 |
| 45.45 | 0.5913 | 0.6388 | 0.6804 | 0.2633 | 0.3372 | 0.3666 | –[a] | 0.2306 | 0.2294 |
| 22.73 | 0.6428 | 0.6817 | 0.6619 | 0.3411 | 0.3530 | 0.3368 | –[a] | 0.1988 | 0.1785 |
| 11.36 | 0.7199 | 0.707 | 0.6804 | 0.3117 | 0.3353 | 0.3367 | –[a] | 0.2009 | 0.2079 |
| 5.68 | 0.3599 | 0.5281 | 0.5334 | 0.3319 | 0.3044 | 0.3194 | –[a] | 0.2051 | 0.2184 |
| 2.84 | 0.4583 | 0.5029 | 0.5118 | 0.2890 | 0.3083 | 0.3598 | –[a] | 0.1856 | 0.1695 |
| 1.42 | 0.5427 | 0.5382 | 0.5462 | 0.3729 | 0.3527 | 0.3606 | –[a] | 0.2272 | 0.2134 |
| 0.71 | 0.6911 | 0.6655 | 0.7001 | 0.4594 | 0.4168 | 0.4461 | –[a] | 0.3080 | 0.2888 |
| 0.36 | 0.7682 | 0.7638 | 0.7504 | 0.5875 | 0.6003 | 0.5346 | –[a] | 0.3263 | 0.3131 |
| 0.18 | 0.8536 | 0.829 | 0.8618 | 0.7609 | 0.6585 | 0.6216 | –[a] | 0.4797 | 0.4245 |
| 0.09 | 0.4946 | 0.7991 | 0.774 | 0.6549 | 0.7420 | 0.6788 | –[a] | 0.3940 | 0.4021 |

[a]*Values were excluded for further data analysis as there were edge effects observed in the 96 well plate by visual inspection.*

*Table 8: Raw data of the absorbance at 550 nm after blank (isopropanol) subtraction of the biological and technical replicates of the MTT assay with MZ3-All-C-AspA used for further data analysis.*

| MZ3-All-C-AspA (IF404) | | | | | | | | | |
|---|---|---|---|---|---|---|---|---|---|
| Inhibitor concentration [µM] | Absorbance 550 nm Biological replicate 1 | | | Absorbance 550 nm Biological replicate 2 | | | Absorbance 550 nm Biological replicate 3 | | |
| 0 | 0.9098 | 0.9074 | 0.8056 | 0.8207 | 0.8267 | 0.7969 | 0.5519 | 0.4124 | –[a] |
| 45.45 | 0.4374 | 0.4054 | 0.3482 | 0.1656 | 0.1653 | 0.1310 | 0.08950 | 0.1514 | –[a] |
| 22.73 | 0.4896 | 0.4897 | 0.422 | 0.1900 | 0.1716 | 0.1541 | 0.1095 | 0.1138 | –[a] |
| 11.36 | 0.5442 | 0.5088 | 0.4816 | 0.1752 | 0.2046 | 0.1456 | 0.1718 | 0.1765 | –[a] |
| 5.68 | 0.3909 | 0.4209 | 0.2994 | 0.2550 | 0.2011 | 0.1780 | 0.2118 | 0.2227 | –[a] |
| 2.84 | 0.51 | 0.5586 | 0.4765 | 0.4027 | 0.3756 | 0.3429 | 0.2471 | 0.2070 | –[a] |
| 1.42 | 0.678 | 0.7248 | 0.7345 | 0.5437 | 0.5345 | 0.5978 | 0.2996 | 0.2720 | –[a] |
| 0.71 | 0.8021 | 0.7982 | 0.7245 | 0.6869 | 0.6600 | 0.5921 | 0.3724 | 0.3619 | –[a] |
| 0.36 | 0.8789 | 0.851 | 0.8484 | 0.6614 | 0.7611 | 0.7059 | 0.5965 | 0.5299 | –[a] |
| 0.18 | 0.815 | 0.8273 | 0.811 | 0.9065 | 0.6973 | 0.8234 | 0.6410 | 0.6779 | –[a] |
| 0.09 | 0.8049 | 0.78 | 0.4665 | 0.8078 | 0.8574 | 0.7178 | 0.6763 | 0.6383 | –[a] |

[a]*Values were excluded for further data analysis as there were edge effects observed in the 96 well plate by visual inspection.*

*Table 9: Raw data of the absorbance at 550 nm after blank (isopropanol) subtraction of the biological and technical replicates of the MTT assay with MZ3-LipA used for further data analysis.*

| MZ3-LipA (IF398) | | | | | | | | | |
|---|---|---|---|---|---|---|---|---|---|
| Inhibitor concentration [µM] | Absorbance 550 nm Biological replicate 1 | | | Absorbance 550 nm Biological replicate 2 | | | Absorbance 550 nm Biological replicate 3 | | |
| 0 | –[a] | 0.5595 | 0.601 | 0.8547 | 0.8404 | 0.5600 | –[a] | 0.8323 | 0.8380 |
| 45.45 | –[a] | 0.3387 | 0.3534 | 0.4030 | 0.4090 | 0.3703 | –[a] | 0.2203 | 0.3014 |
| 22.73 | –[a] | 0.3517 | 0.3546 | 0.4612 | 0.4977 | 0.3576 | –[a] | 0.2820 | 0.3031 |
| 11.36 | –[a] | 0.3577 | 0.3996 | 0.4765 | 0.4683 | 0.3984 | –[a] | 0.2297 | 0.2839 |
| 5.68 | –[a] | 0.3619 | 0.4213 | 0.4685 | 0.4109 | 0.3926 | –[a] | 0.2044 | 0.2429 |
| 2.84 | –[a] | 0.3315 | 0.3664 | 0.4437 | 0.4479 | 0.4028 | –[a] | 0.2324 | 0.2128 |
| 1.42 | –[a] | 0.4025 | 0.3928 | 0.4788 | 0.4599 | 0.4901 | –[a] | 0.2659 | 0.2644 |
| 0.71 | –[a] | 0.4998 | 0.4845 | 0.5681 | 0.5523 | 0.5107 | –[a] | 0.3195 | 0.3150 |
| 0.36 | –[a] | 0.5 | 0.5111 | 0.8631 | 0.8517 | 0.7439 | –[a] | 0.4910 | 0.5262 |
| 0.18 | –[a] | 0.5805 | 0.5767 | 0.9981 | 1.037 | 0.9266 | –[a] | 0.6299 | 0.6847 |
| 0.09 | –[a] | 0.6407 | 0.6091 | 0.8170 | 0.9365 | 0.7699 | –[a] | 0.4925 | 0.6036 |

[a]*Values were excluded for further data analysis as there were edge effects observed in the 96 well plate by visual inspection.*

*Table 10: Raw data of the absorbance at 550 nm after blank (isopropanol) subtraction of the biological and technical replicates of the MTT assay with MZ3-All-C-LipA used for further data analysis.*

| MZ3-All-C-LipA (IF399) | | | | | | | | | |
|---|---|---|---|---|---|---|---|---|---|
| Inhibitor concentration [µM] | Absorbance 550 nm Biological replicate 1 | | | Absorbance 550 nm Biological replicate 2 | | | Absorbance 550 nm Biological replicate 3 | | |
| 0 | 0.6295 | 0.7138 | –[a] | 0.7084 | 0.8618 | 0.6673 | 0.8337 | 0.8051 | –[a] |
| 45.45 | 0.0564 | 0.07 | –[a] | 0.07760 | 0.07800 | 0.1024 | 0.09120 | 0.09950 | –[a] |
| 22.73 | 0.0843 | 0.0823 | –[a] | 0.07730 | 0.1109 | 0.07930 | 0.08290 | 0.08430 | –[a] |
| 11.36 | 0.0986 | 0.0916 | –[a] | 0.08200 | 0.07520 | 0.07870 | 0.09000 | 0.06840 | –[a] |
| 5.68 | 0.0863 | 0.0828 | –[a] | 0.07500 | 0.08570 | 0.09030 | 0.1081 | 0.1139 | –[a] |
| 2.84 | 0.1112 | 0.1097 | –[a] | 0.08450 | 0.08020 | 0.07830 | 0.1012 | 0.1168 | –[a] |
| 1.42 | 0.2571 | 0.2248 | –[a] | 0.2019 | 0.2153 | 0.1607 | 0.2094 | 0.1828 | –[a] |
| 0.71 | 0.3133 | 0.2965 | –[a] | 0.5331 | 0.5337 | 0.4631 | 0.4816 | 0.4546 | –[a] |
| 0.36 | 0.4879 | 0.4635 | –[a] | 0.7251 | 0.5823 | 0.7088 | 0.5977 | 0.5476 | –[a] |
| 0.18 | 0.5518 | 0.5548 | –[a] | 0.6711 | 0.8832 | 0.9523 | 0.7789 | 0.7271 | –[a] |
| 0.09 | 0.5964 | 0.5994 | –[a] | 0.7821 | 0.8603 | 0.7332 | 0.6717 | 0.6240 | –[a] |

[a]*Values were excluded for further data analysis as there were edge effects observed in the 96 well plate by visual inspection.*

### 1.4. UHPLC-MS/UV spectra

The UHPLC-MS/UV spectra for the final MZ1-, and MZ3-derivatives were acquired on UHPLC-MS B using a linear gradient of 10-95% B over 3 min (then 2 min 95% B). The samples were prepared using MeOH as the solvent. Data visualization and processing was performed using the MestReNova *v14.1.2-25024*[1] Mass plugin. The total absorbance chromatograms are shown in the appendix.

### 3. Appendix

#### 3.1. NMR spectra

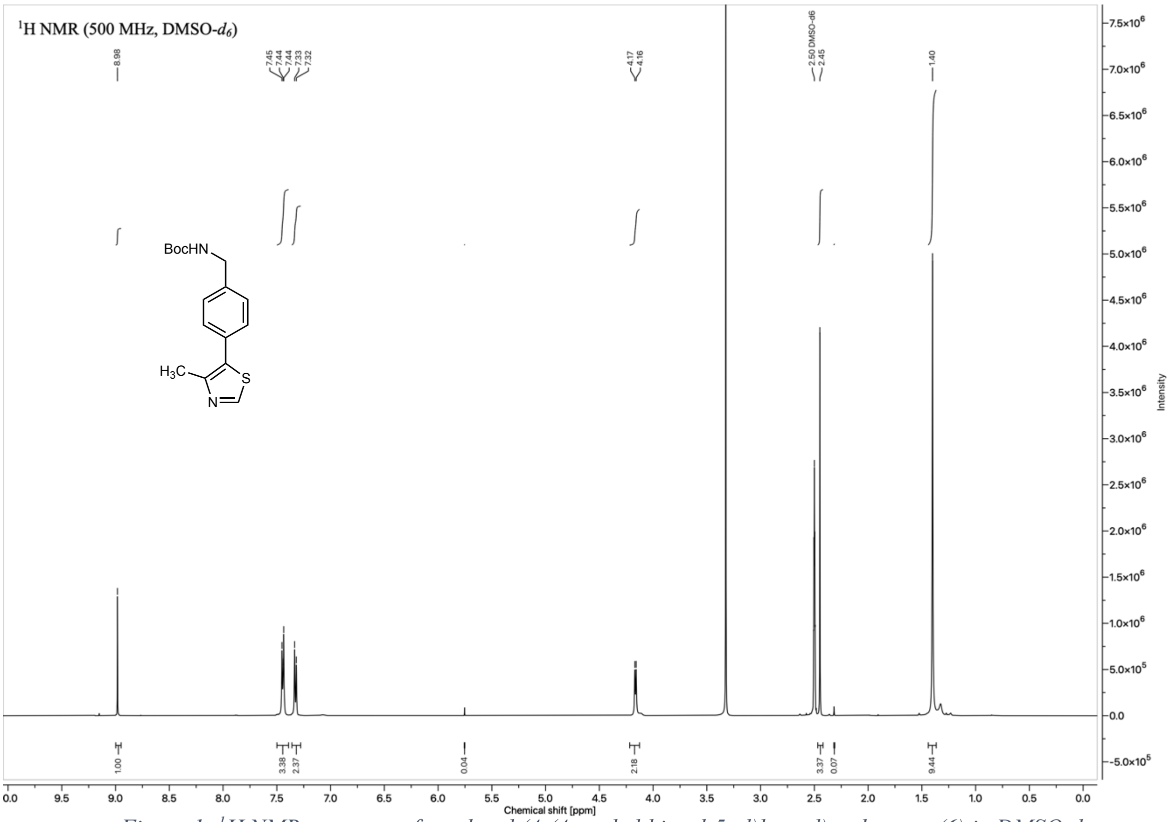

*Figure 1: ¹H NMR spectrum of tert-butyl (4-(4-methylthiazol-5-yl)benzyl)carbamate (**6**) in DMSO-$d_6$*

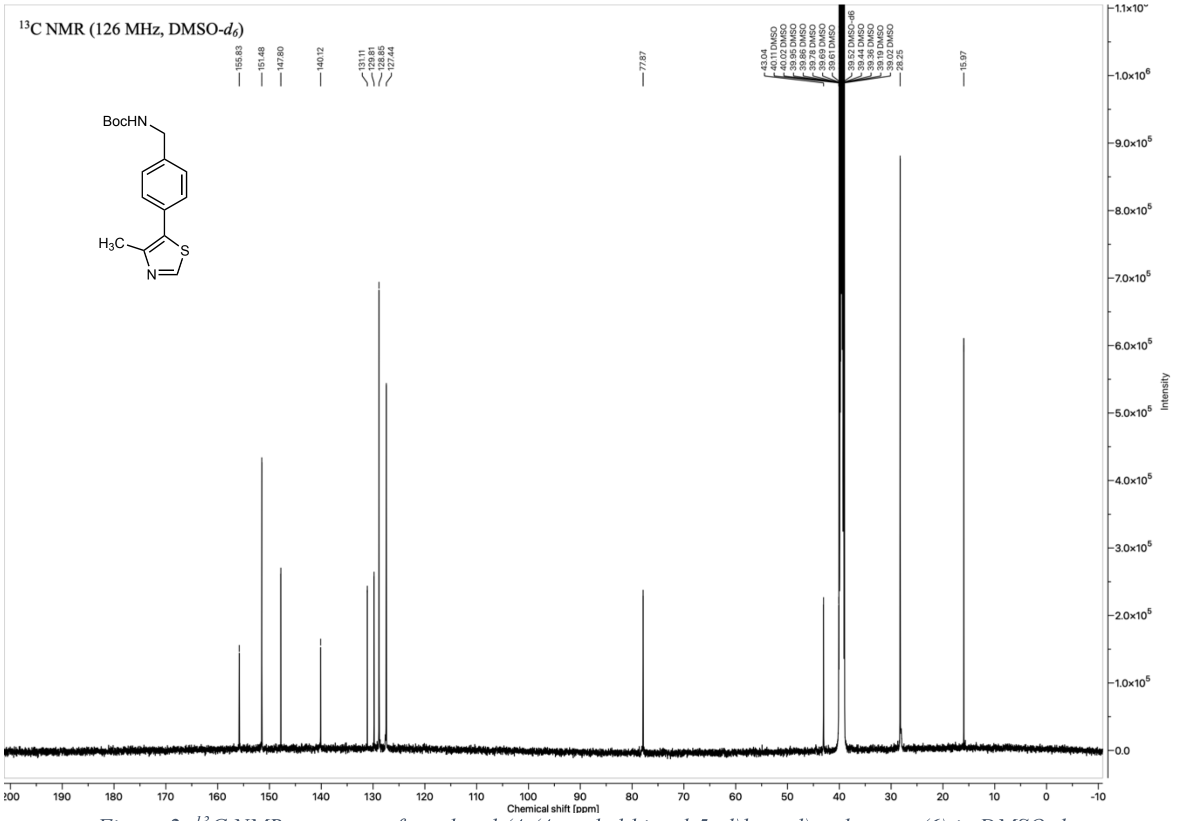

*Figure 2: ¹³C NMR spectrum of tert-butyl (4-(4-methylthiazol-5-yl)benzyl)carbamate (**6**) in DMSO-$d_6$*

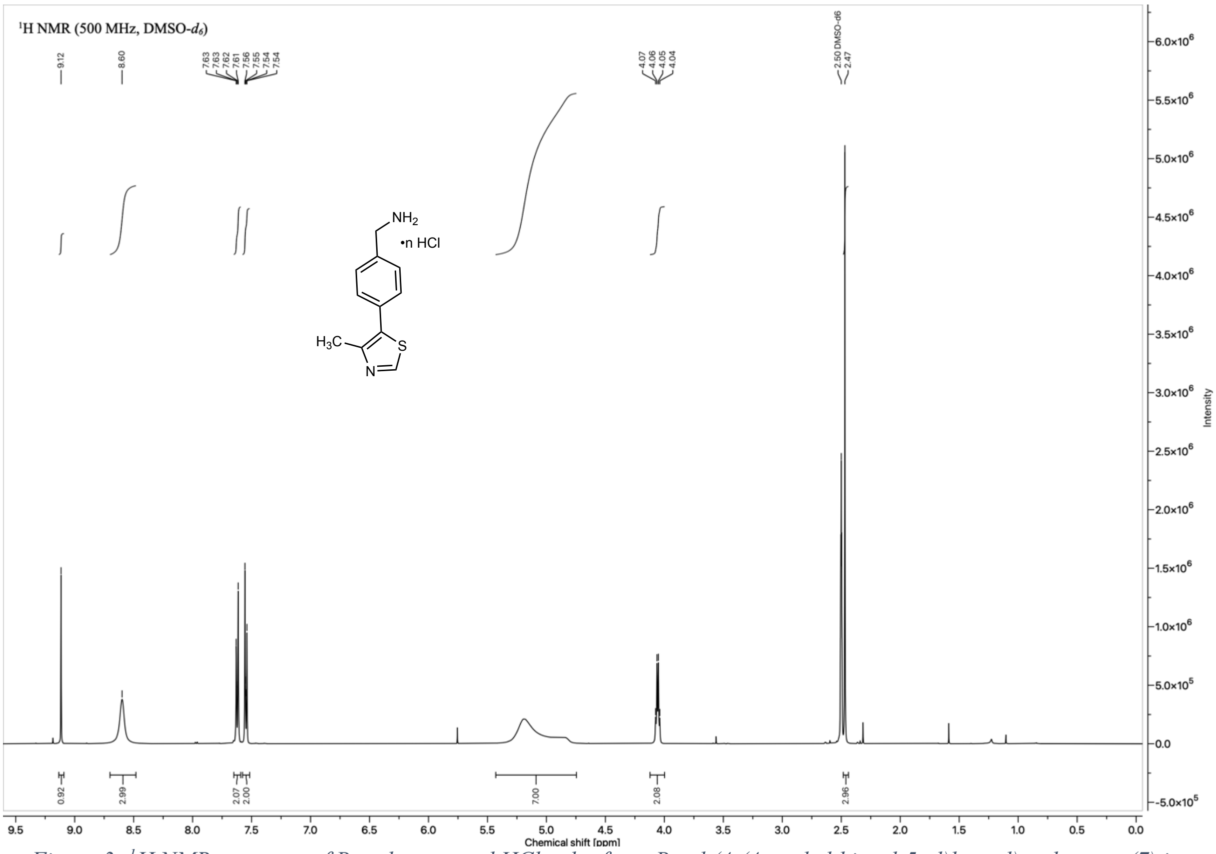

*Figure 3: $^{1}H$ NMR spectrum of Boc-deprotected HCl-salt of tert-Butyl (4-(4-methylthiazol-5-yl)benzyl)carbamate (7) in DMSO-$d_6$*

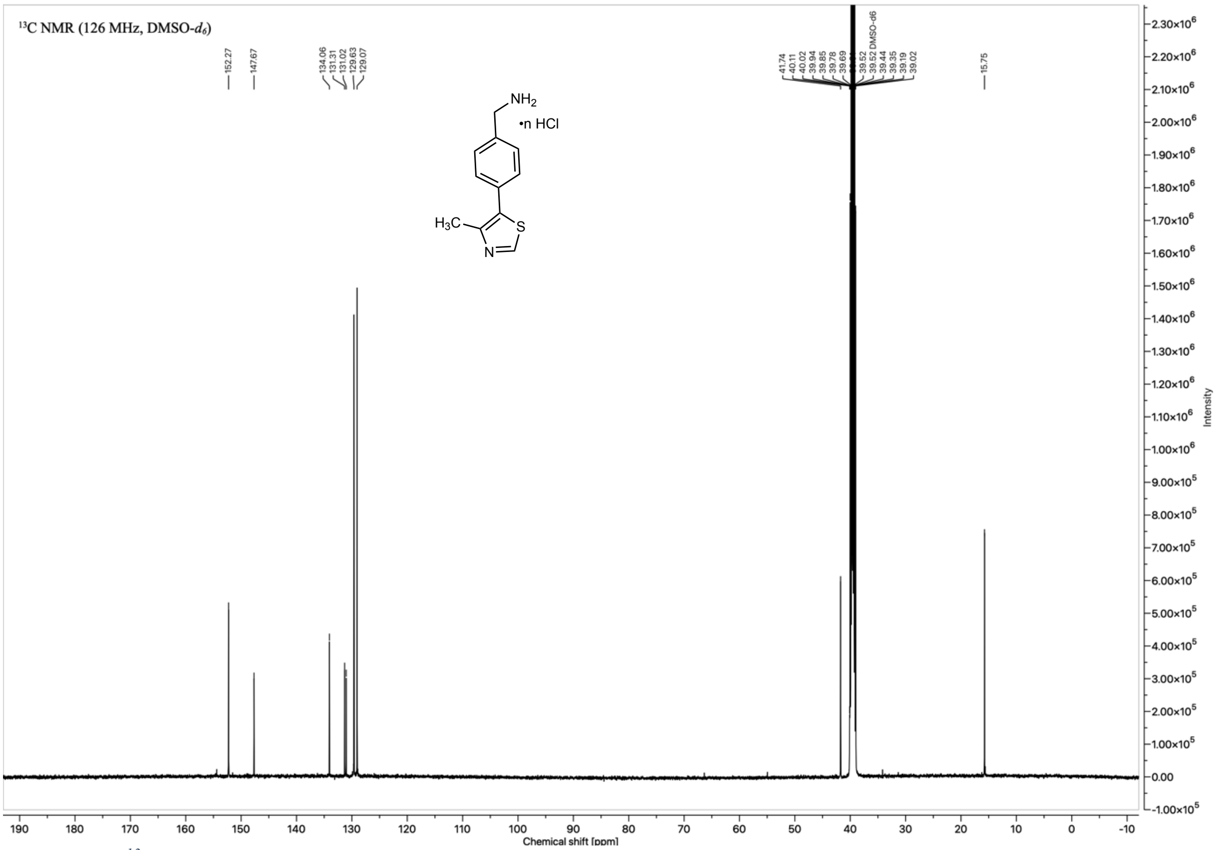

*Figure 4: $^{13}C$ NMR spectrum of Boc-deprotected HCl-salt of tert-Butyl (4-(4-methylthiazol-5-yl)benzyl)carbamate (7) in DMSO-$d_6$*

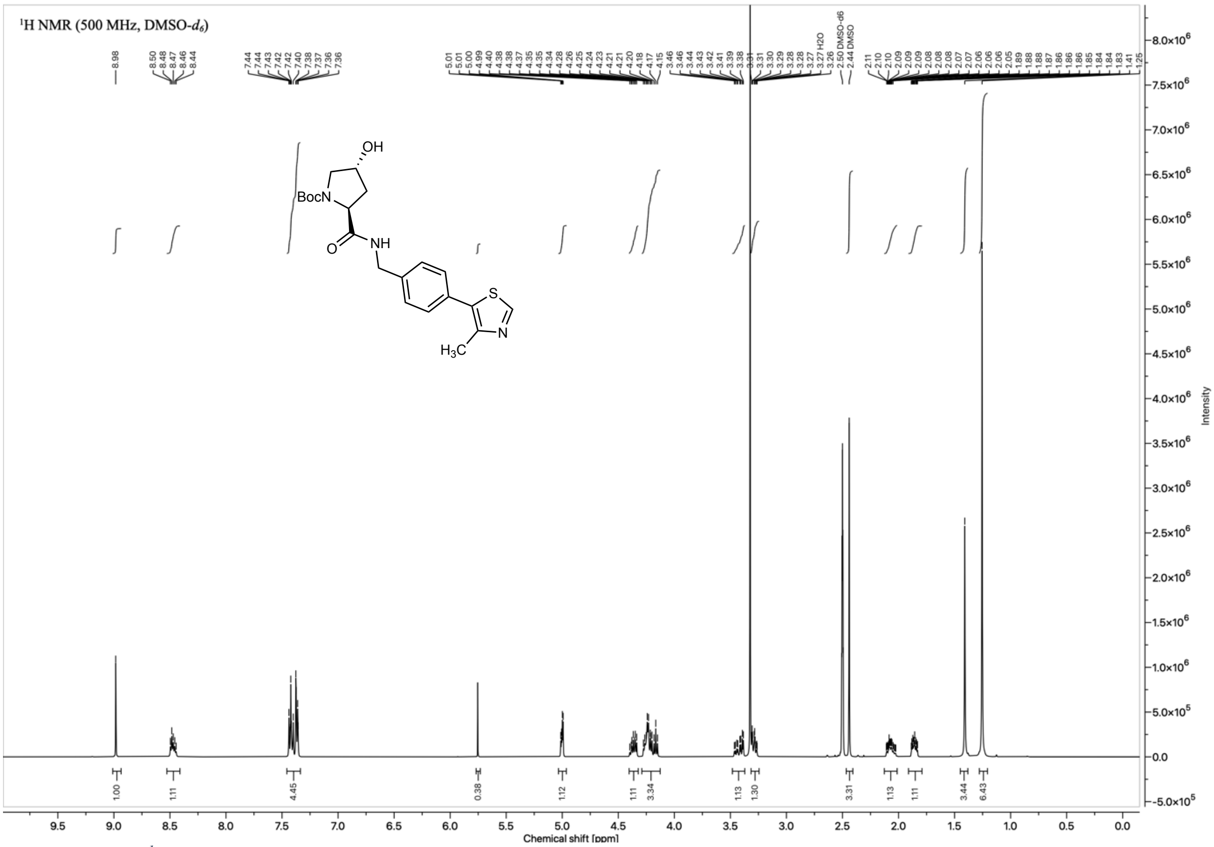

*Figure 5: $^{1}H$ NMR spectrum of tert-Butyl (2S,4R)-4-hydroxy-2-[[4-(4-methylthiazol-5-yl)phenyl] methylcarbamoyl] pyrrolidine-1-carboxylate (**8**) in DMSO-d$_6$.*

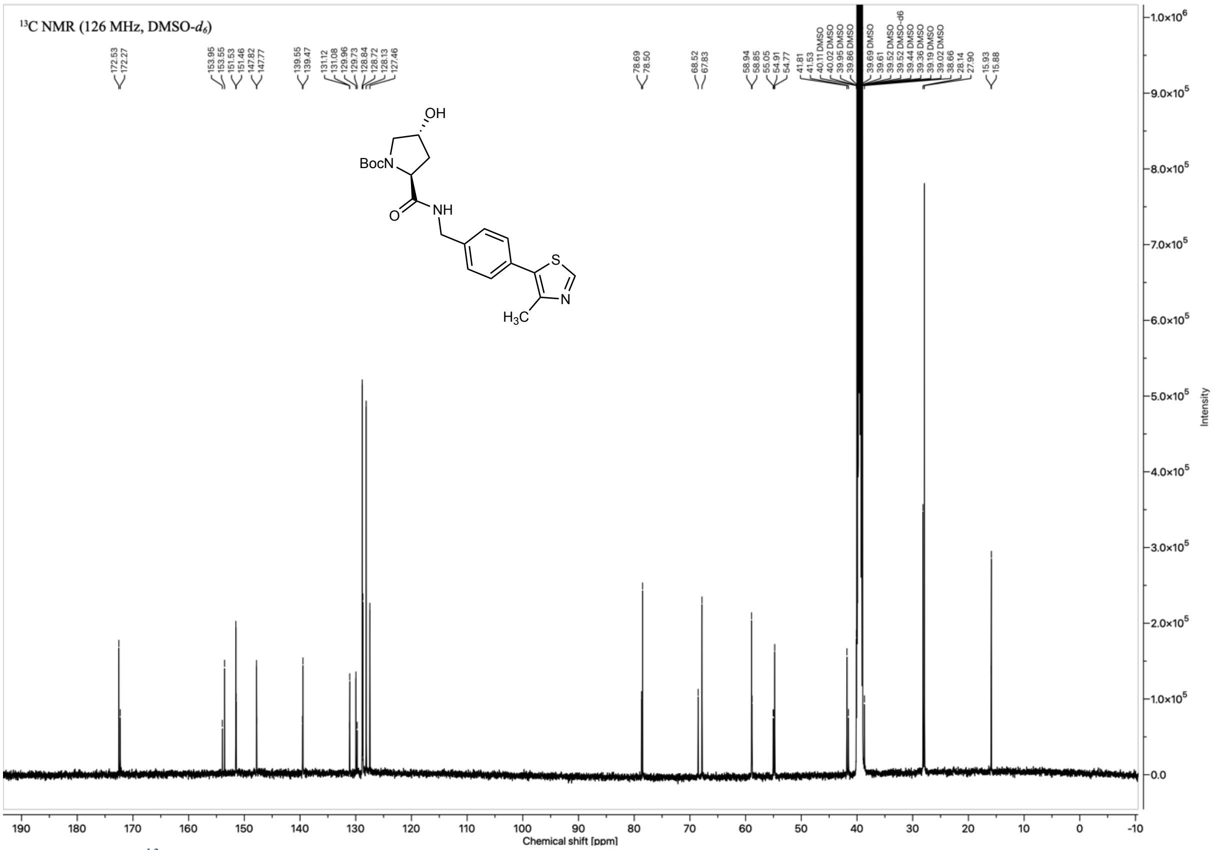

*Figure 6: $^{13}C$ NMR spectrum of tert-Butyl (2S,4R)-4-hydroxy-2-[[4-(4-methylthiazol-5-yl)phenyl] methylcarbamoyl] pyrrolidine-1-carboxylate (**8**) in DMSO-d$_6$.*

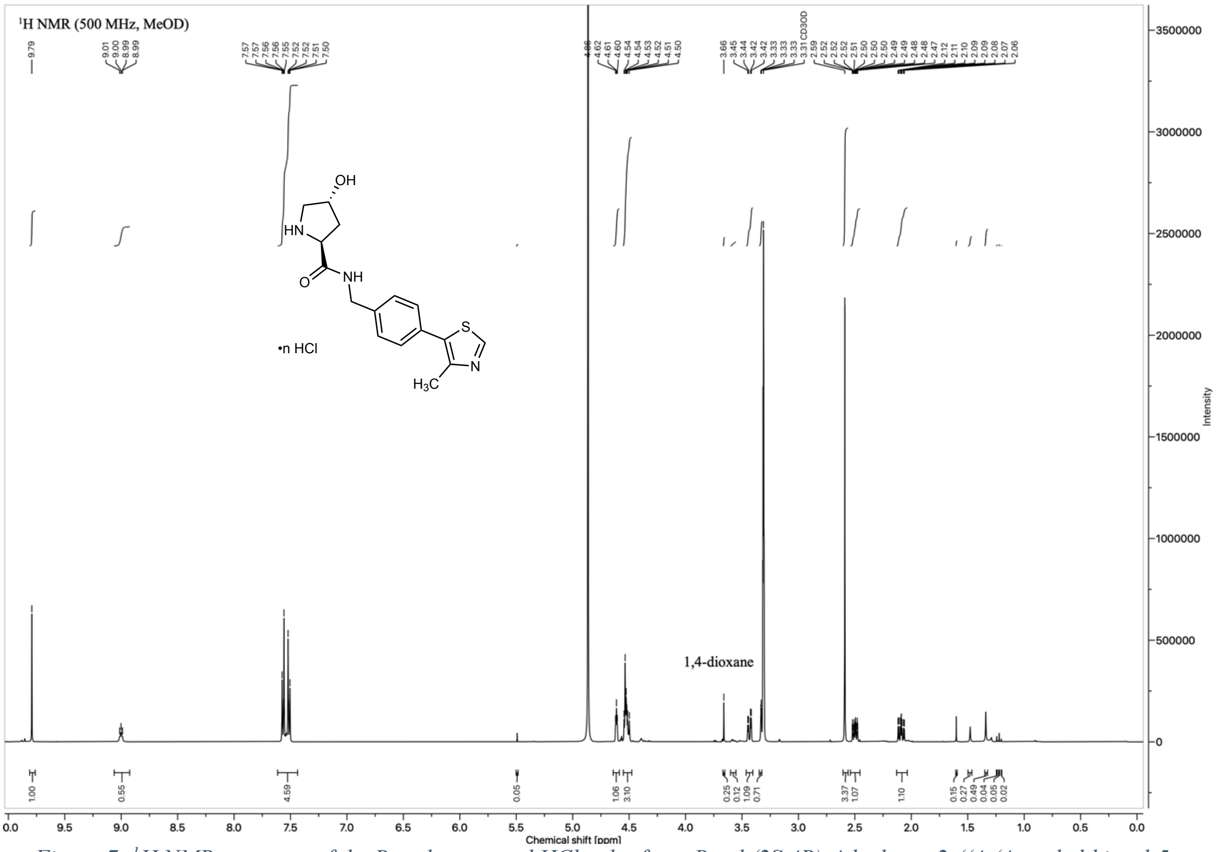

*Figure 7: $^{1}$H NMR spectrum of the Boc-deprotected HCl-salt of tert-Butyl (2S,4R)-4-hydroxy-2-((4-(4-methylthiazol-5-yl)benzyl) carbamoyl)pyrrolidine-1-carboxylate (**9**) in MeOD*

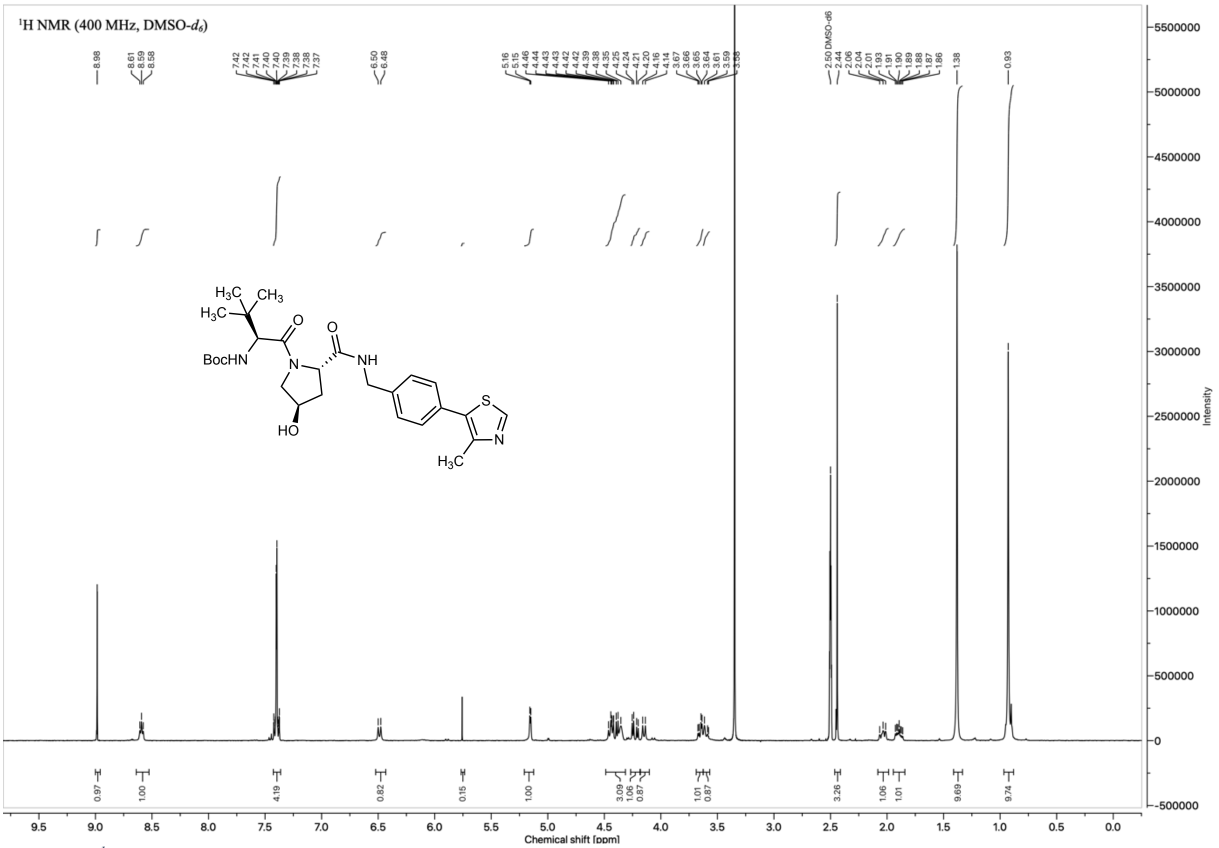

*Figure 8: $^{1}$H NMR spectrum of tert-butyl ((S)-1-((2S,4R)-4-hydroxy-2-((4-(4-methylthiazol-5-yl)benzyl)carbamoyl)pyrrolidin-1-yl)-3,3-dimethyl-1-oxobutan-2-yl)carbamate (**10**) in DMSO-$d_6$.*

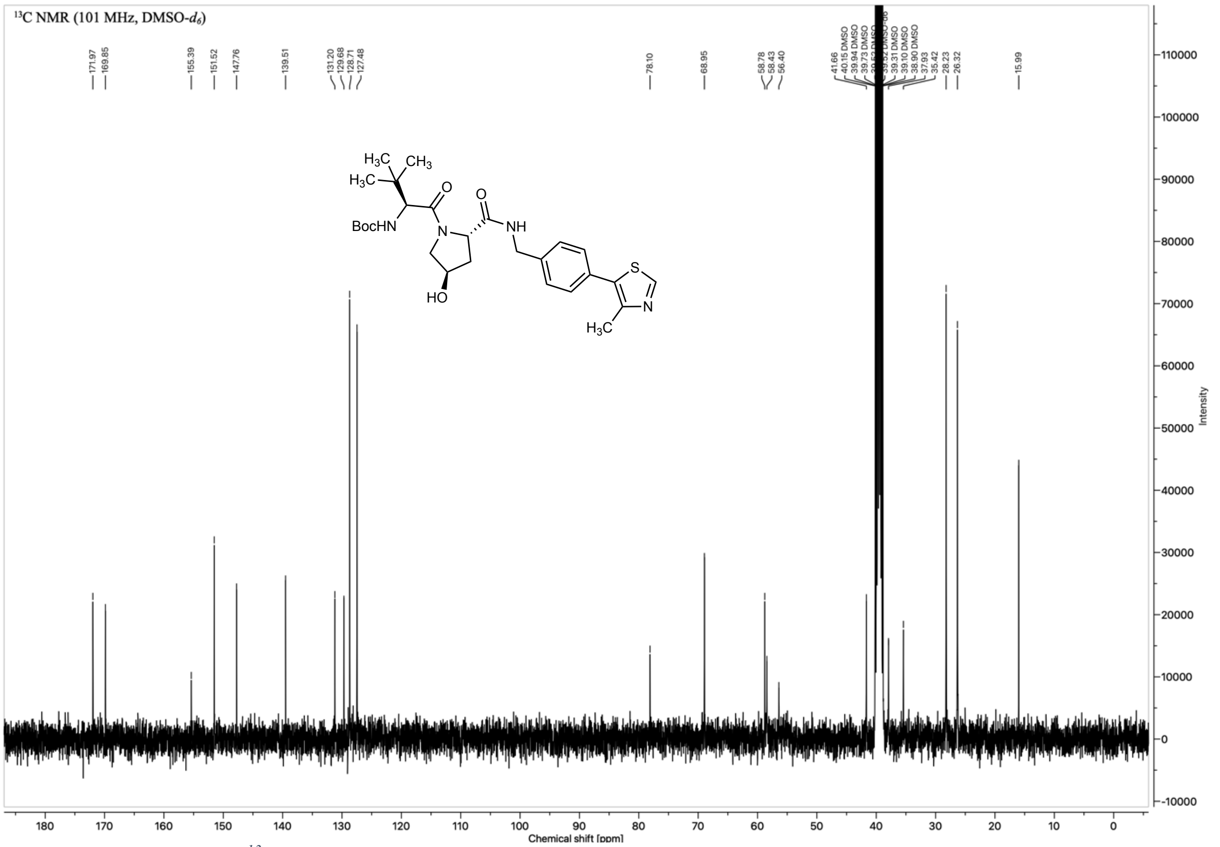

*Figure 9: $^{13}C$ NMR spectrum of tert-butyl ((S)-1-((2S,4R)-4-hydroxy-2-((4-(4-methylthiazol-5-yl)benzyl)carbamoyl)pyrrolidin-1-yl)-3,3-dimethyl-1-oxobutan-2-yl)carbamate (**10**) in DMSO-d$_6$.*

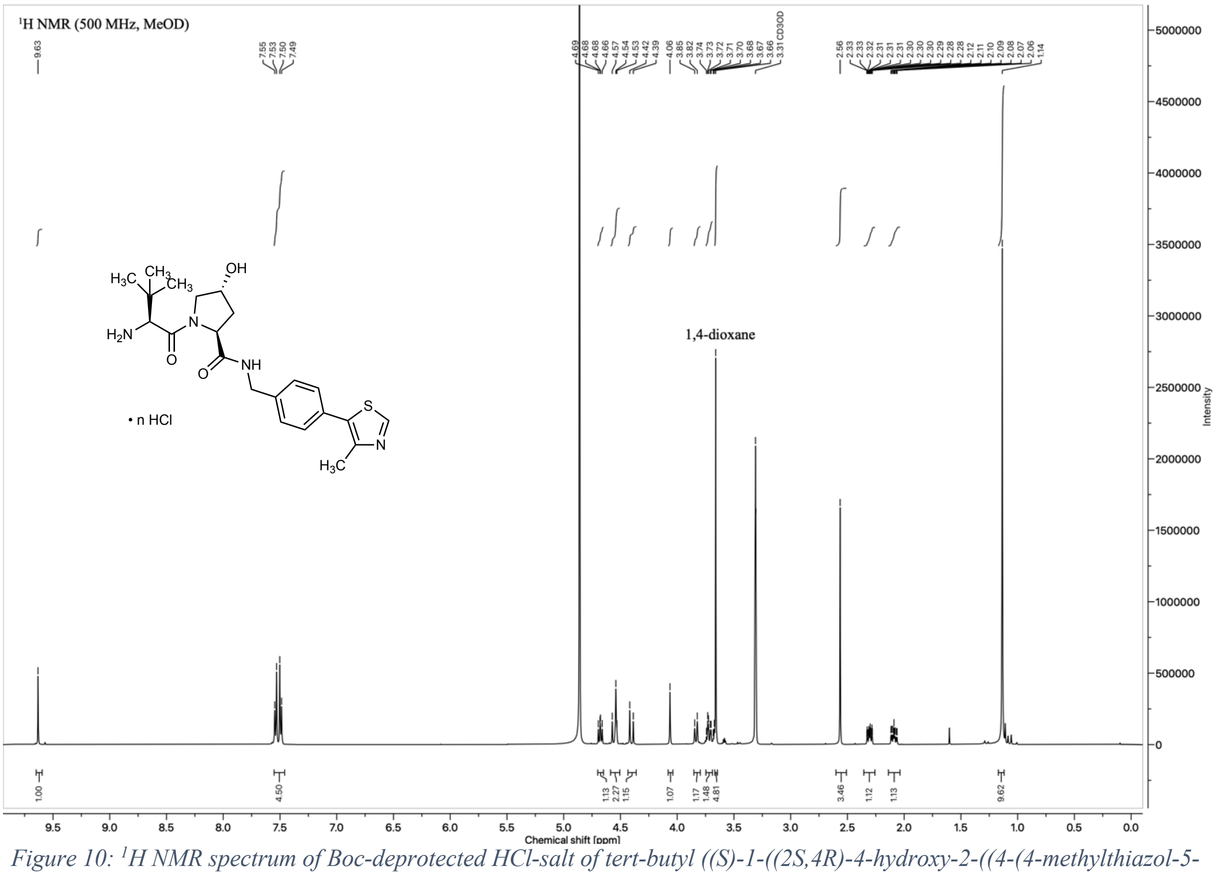

*Figure 10: $^{1}H$ NMR spectrum of Boc-deprotected HCl-salt of tert-butyl ((S)-1-((2S,4R)-4-hydroxy-2-((4-(4-methylthiazol-5-yl)benzyl)carbamoyl)pyrrolidin-1-yl)-3,3-dimethyl-1-oxobutan-2-yl)carbamate (**3**, (S,R,S)-AHPC•HCl salt) in $CD_3OD$*

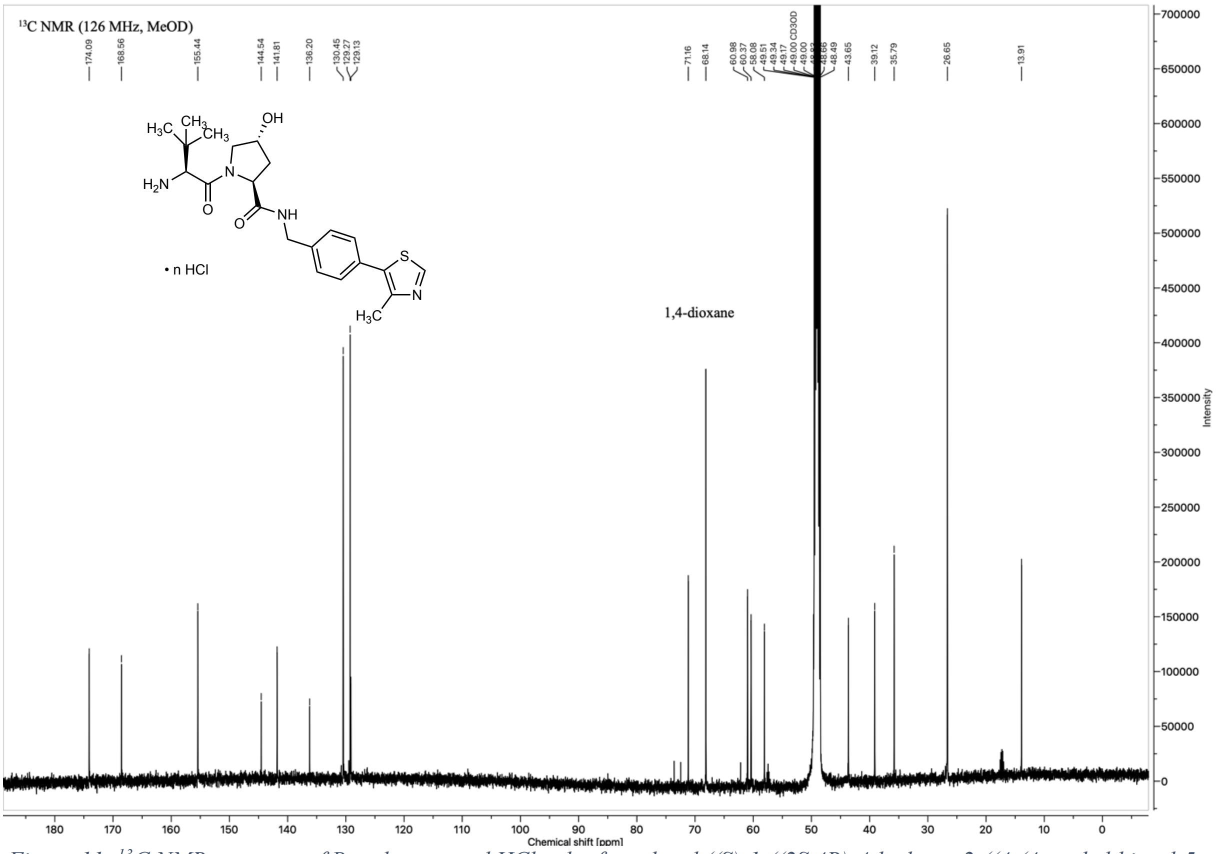

*Figure 11: $^{13}$C NMR spectrum of Boc-deprotected HCl-salt of tert-butyl ((S)-1-((2S,4R)-4-hydroxy-2-((4-(4-methylthiazol-5-yl)benzyl)carbamoyl)pyrrolidin-1-yl)-3,3-dimethyl-1-oxobutan-2-yl)carbamate (**3**, (S,R,S)-AHPC•HCl salt) in $CD_3OD$*

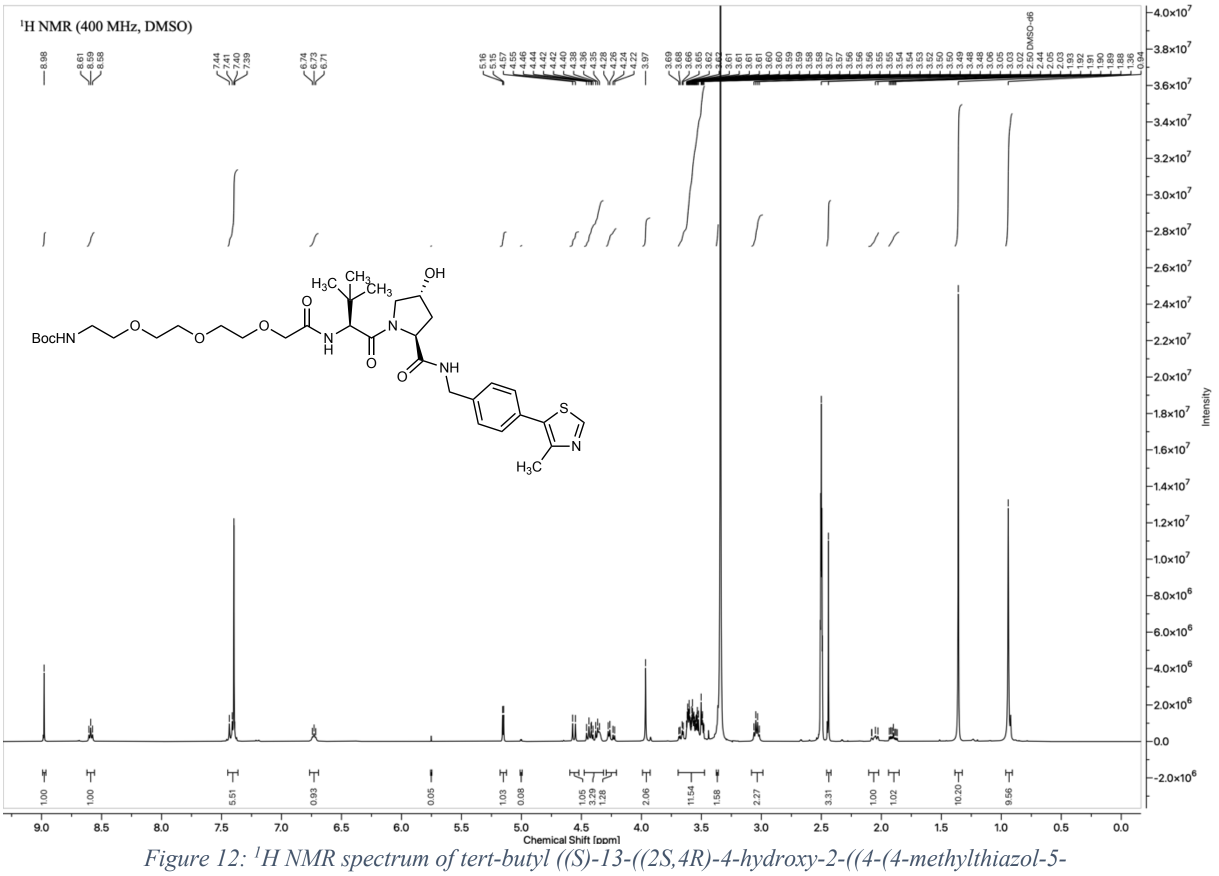

*Figure 12: $^{1}$H NMR spectrum of tert-butyl ((S)-13-((2S,4R)-4-hydroxy-2-((4-(4-methylthiazol-5-yl)benzyl)carbamoyl)pyrrolidine-1-carbonyl)-14,14-dimethyl-11-oxo-3,6,9-trioxa-12-azapentadecyl)carbamate (**11**) in DMSO-$d_6$*

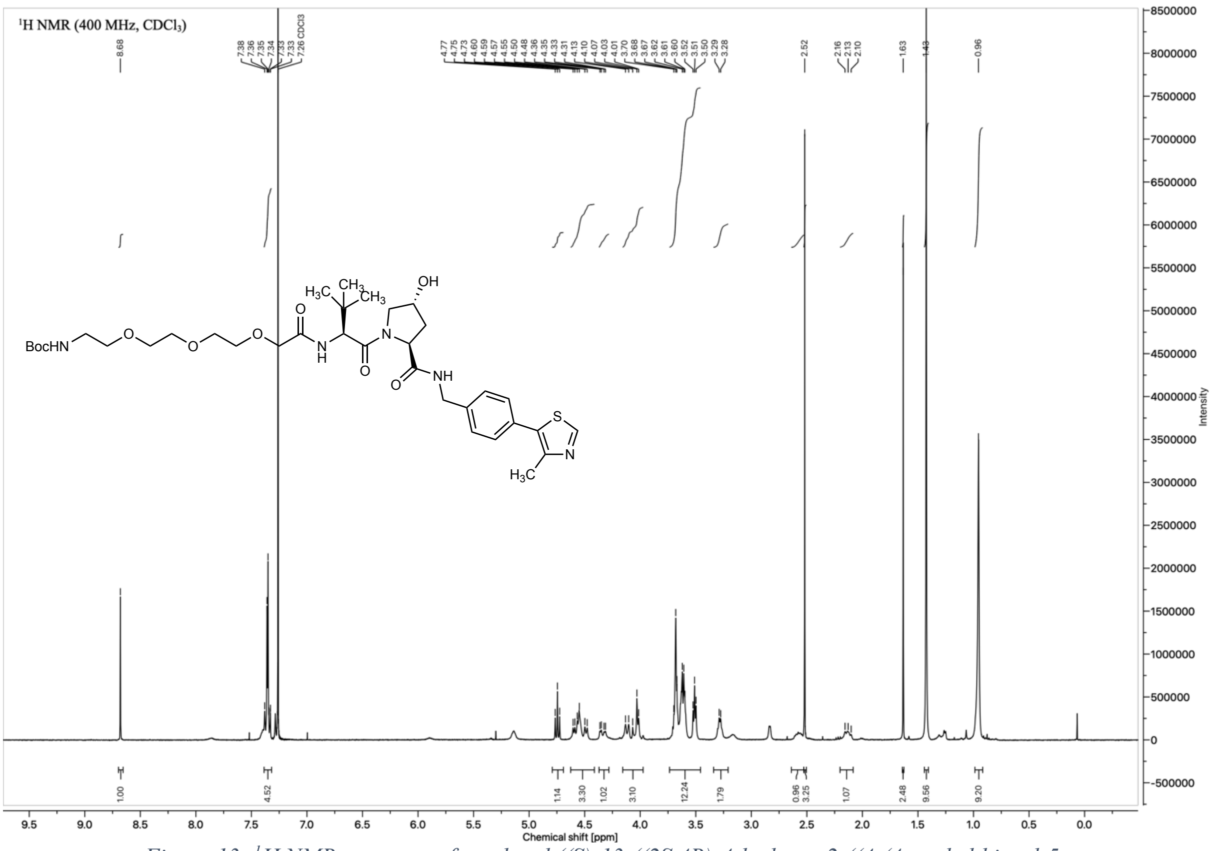

*Figure 13: [1]H NMR spectrum of tert-butyl ((S)-13-((2S,4R)-4-hydroxy-2-((4-(4-methylthiazol-5-yl)benzyl)carbamoyl)pyrrolidine-1-carbonyl)-14,14-dimethyl-11-oxo-3,6,9-trioxa-12-azapentadecyl)carbamate (**11**) in $CDCl_3$*

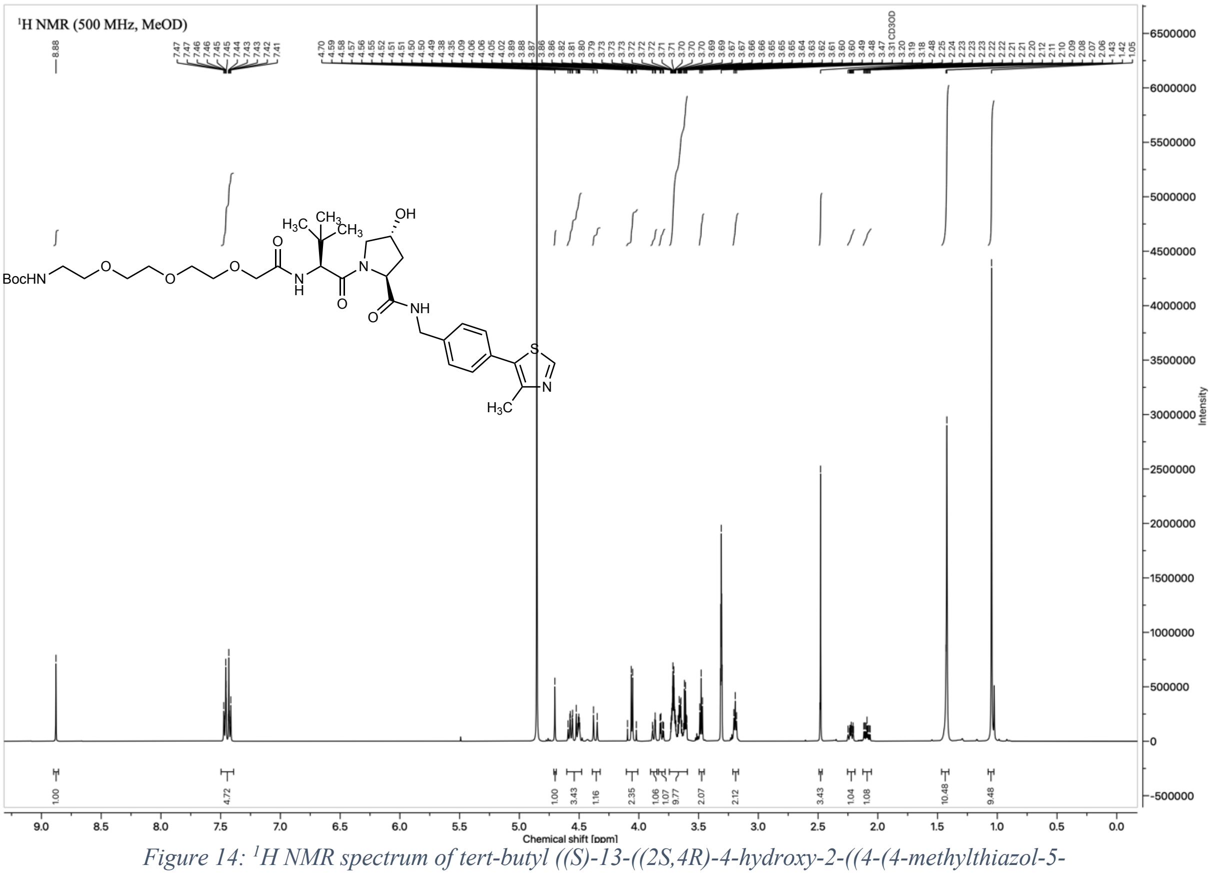

*Figure 14: [1]H NMR spectrum of tert-butyl ((S)-13-((2S,4R)-4-hydroxy-2-((4-(4-methylthiazol-5-yl)benzyl)carbamoyl)pyrrolidine-1-carbonyl)-14,14-dimethyl-11-oxo-3,6,9-trioxa-12-azapentadecyl)carbamate (**11**) in $CD_3OD$*

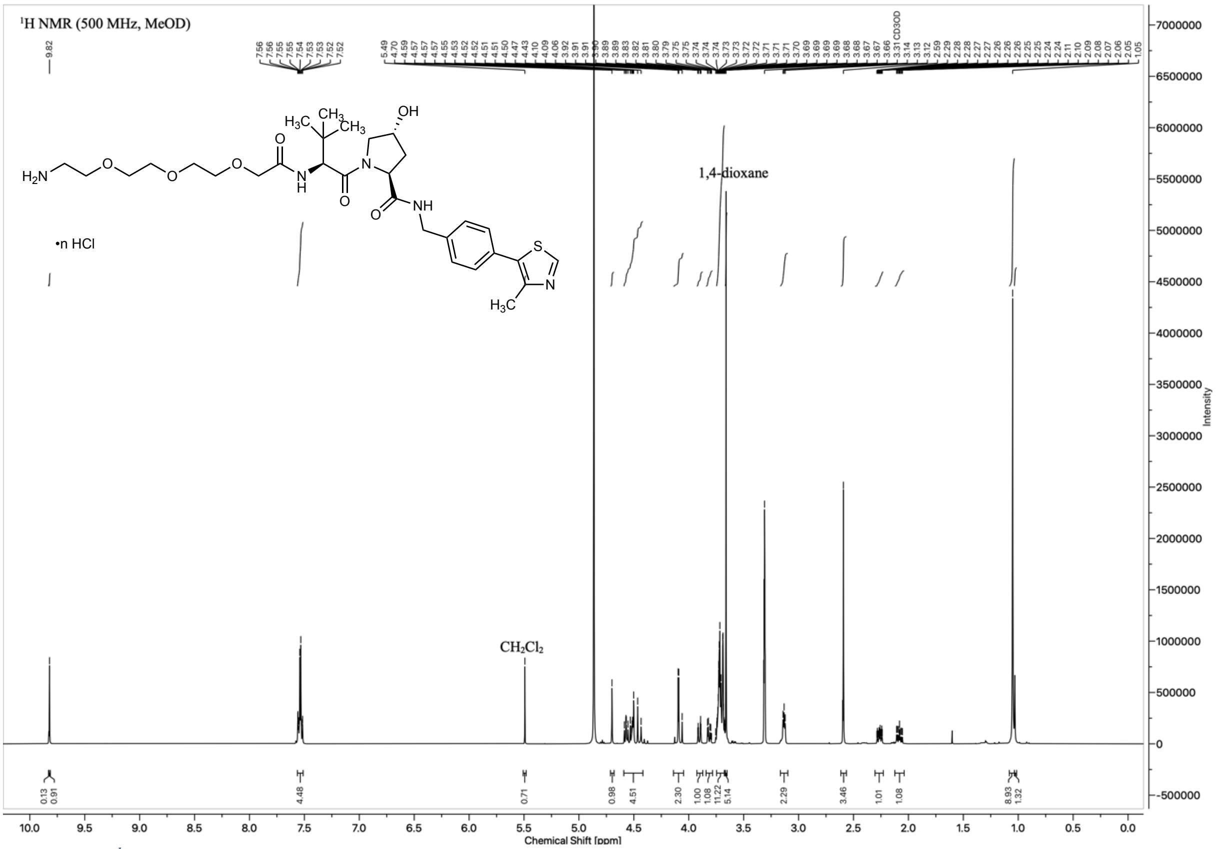

*Figure 15: [1]H NMR spectrum of Boc-deprotected HCl-salt of tert-butyl ((S)-13-((2S,4R)-4-hydroxy-2-((4-(4-methylthiazol-5-yl)benzyl)carbamoyl)pyrrolidine-1-carbonyl)-14,14-dimethyl-11-oxo-3,6,9-trioxa-12-azapentadecyl)carbamate (**12**) in $CD_3OD$*

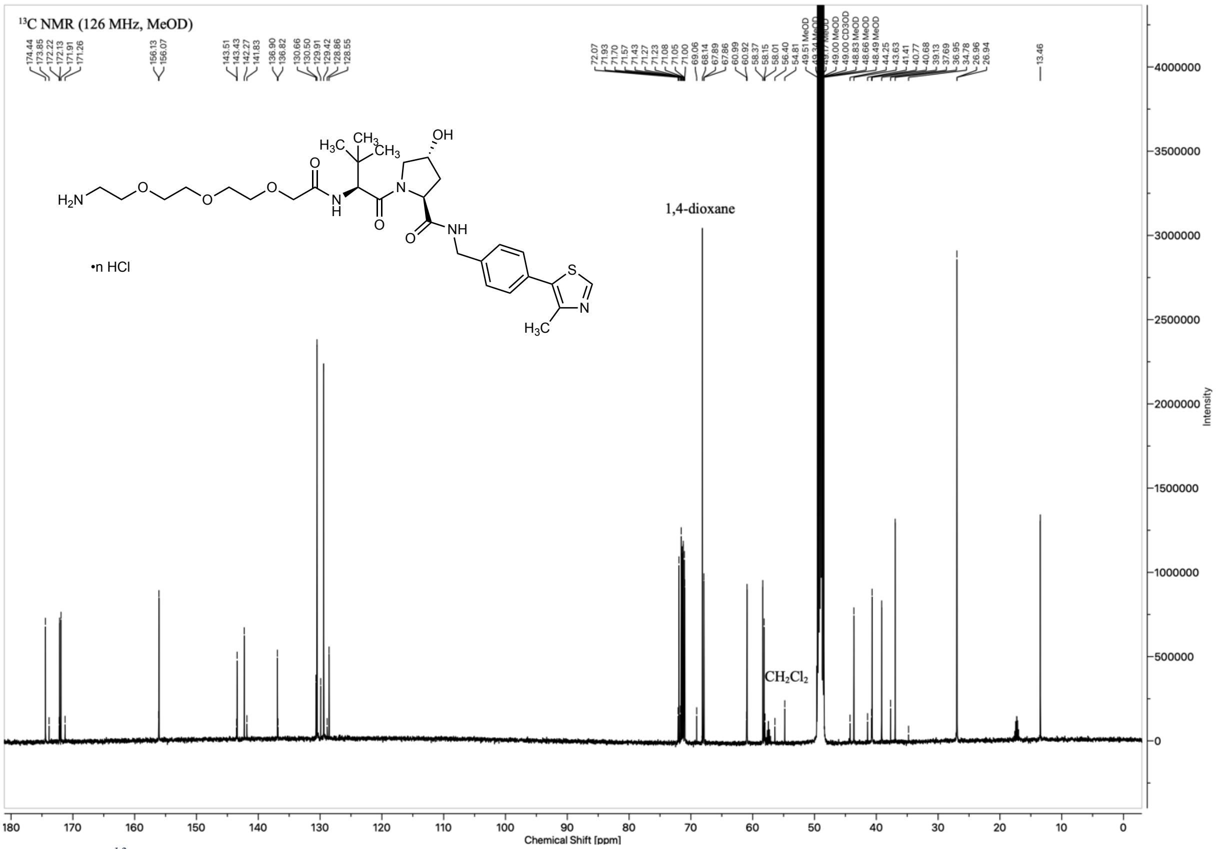

*Figure 16: [13]C NMR spectrum of Boc-deprotected HCl-salt of tert-butyl ((S)-13-((2S,4R)-4-hydroxy-2-((4-(4-methylthiazol-5-yl)benzyl)carbamoyl)pyrrolidine-1-carbonyl)-14,14-dimethyl-11-oxo-3,6,9-trioxa-12-azapentadecyl)carbamate (**12**) in $CD_3OD$*

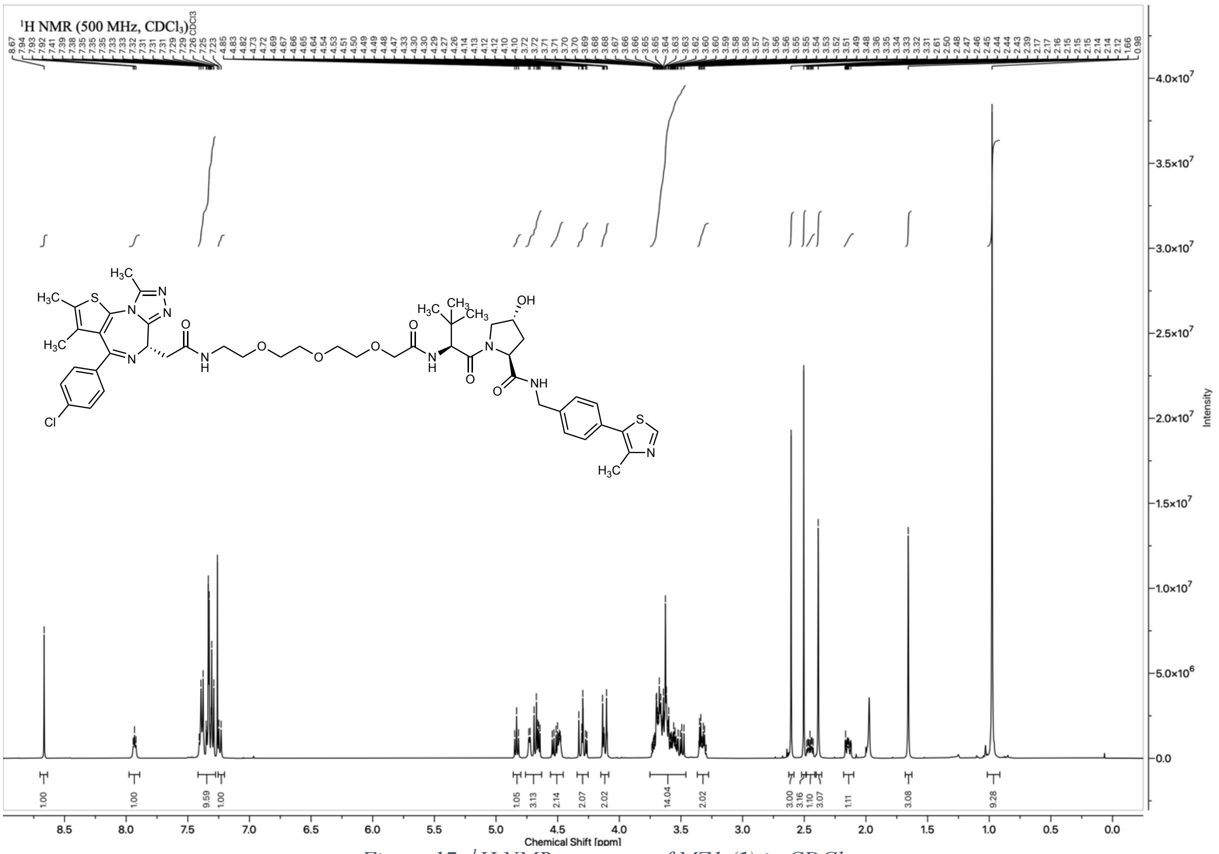

*Figure 17: $^{1}H$ NMR spectrum of MZ1 (**1**) in $CDCl_3$*

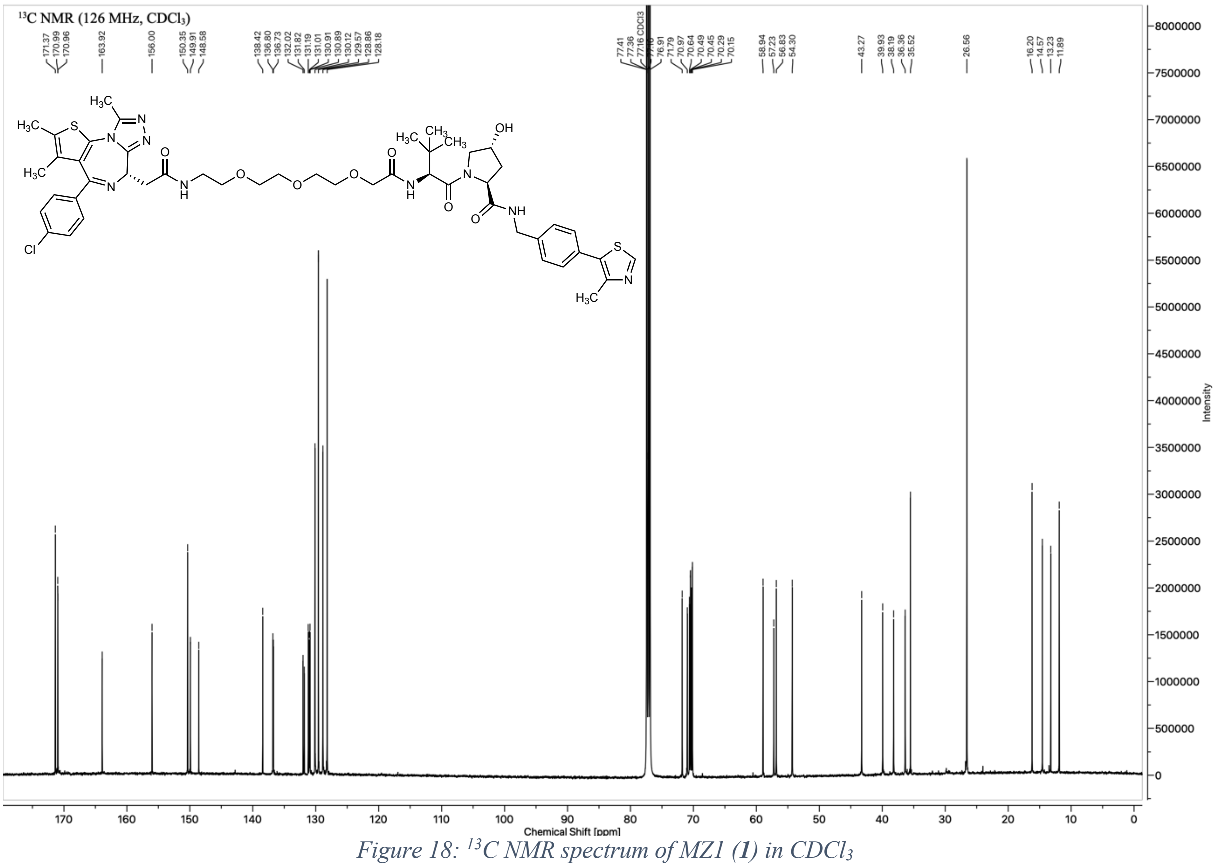

*Figure 18: $^{13}C$ NMR spectrum of MZ1 (**1**) in $CDCl_3$*

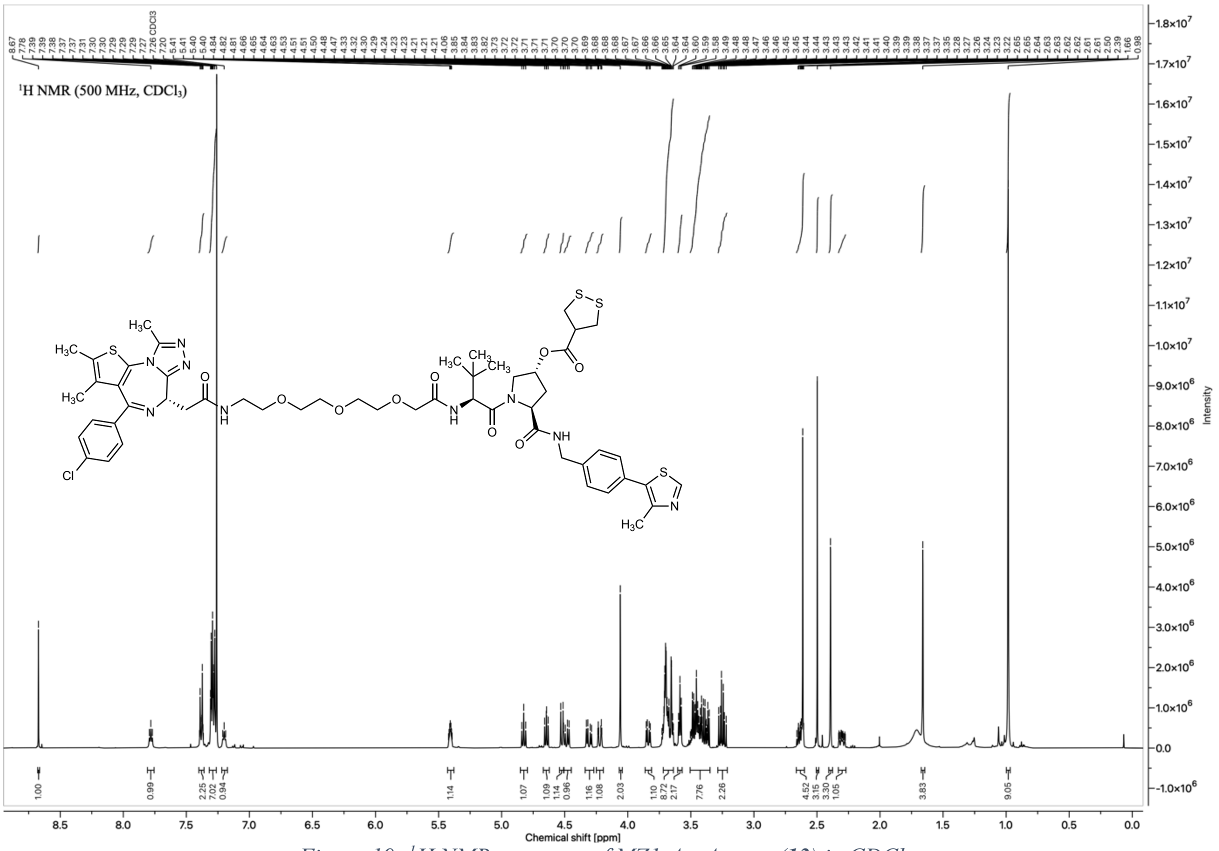

*Figure 19: $^{1}H$ NMR spectrum of MZ1-AspA ester (**13**) in $CDCl_3$*

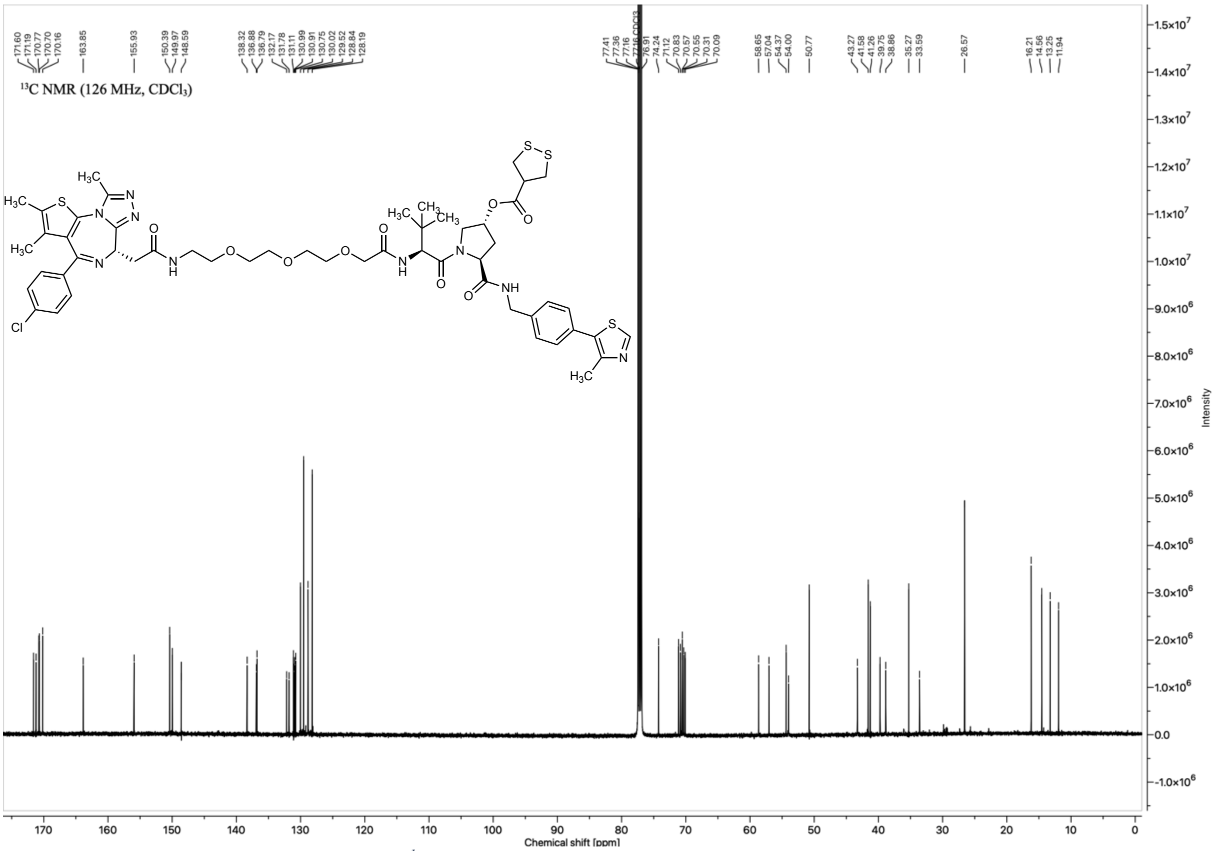

*Figure 20: $^{1}H$ NMR spectrum of MZ1-AspA ester (**13**) in $CDCl_3$*

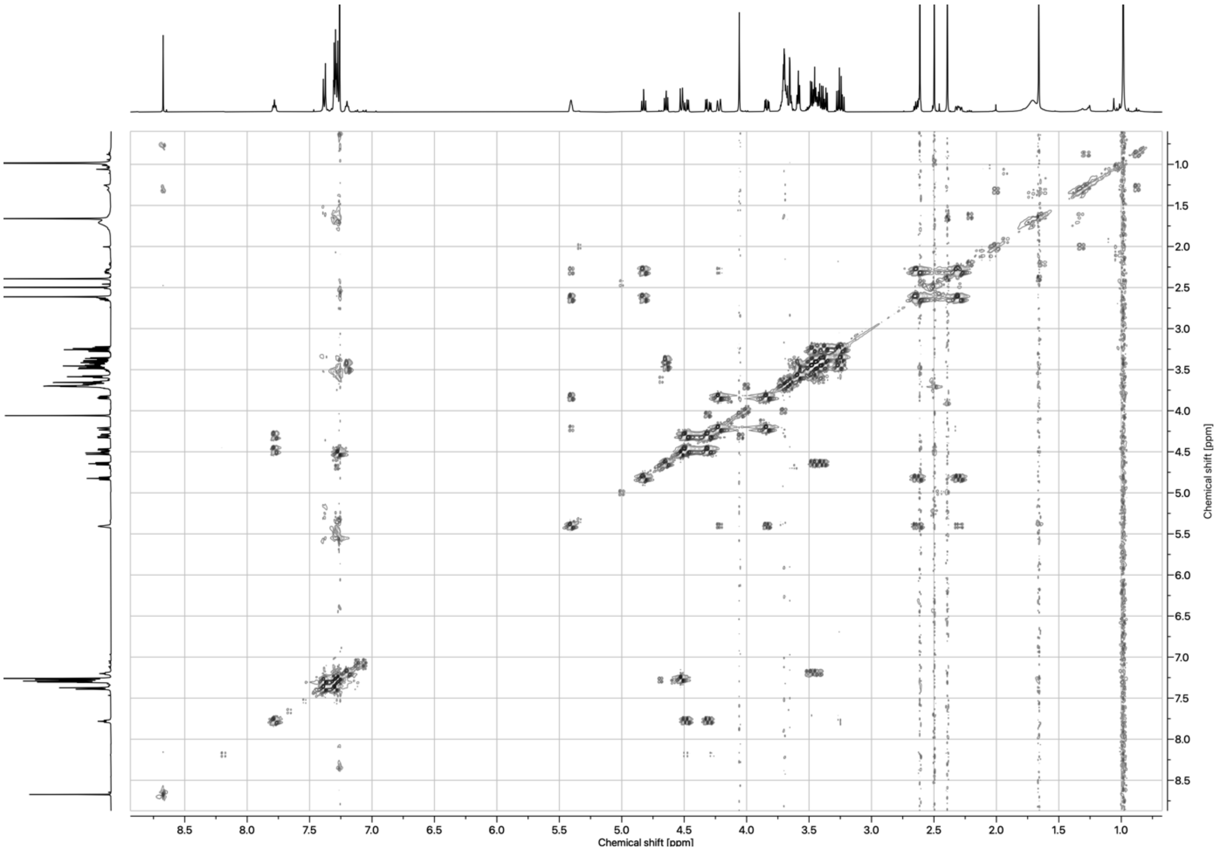

*Figure 21: COSY spectrum of MZ1-AspA ester (**13**) in $CDCl_3$*

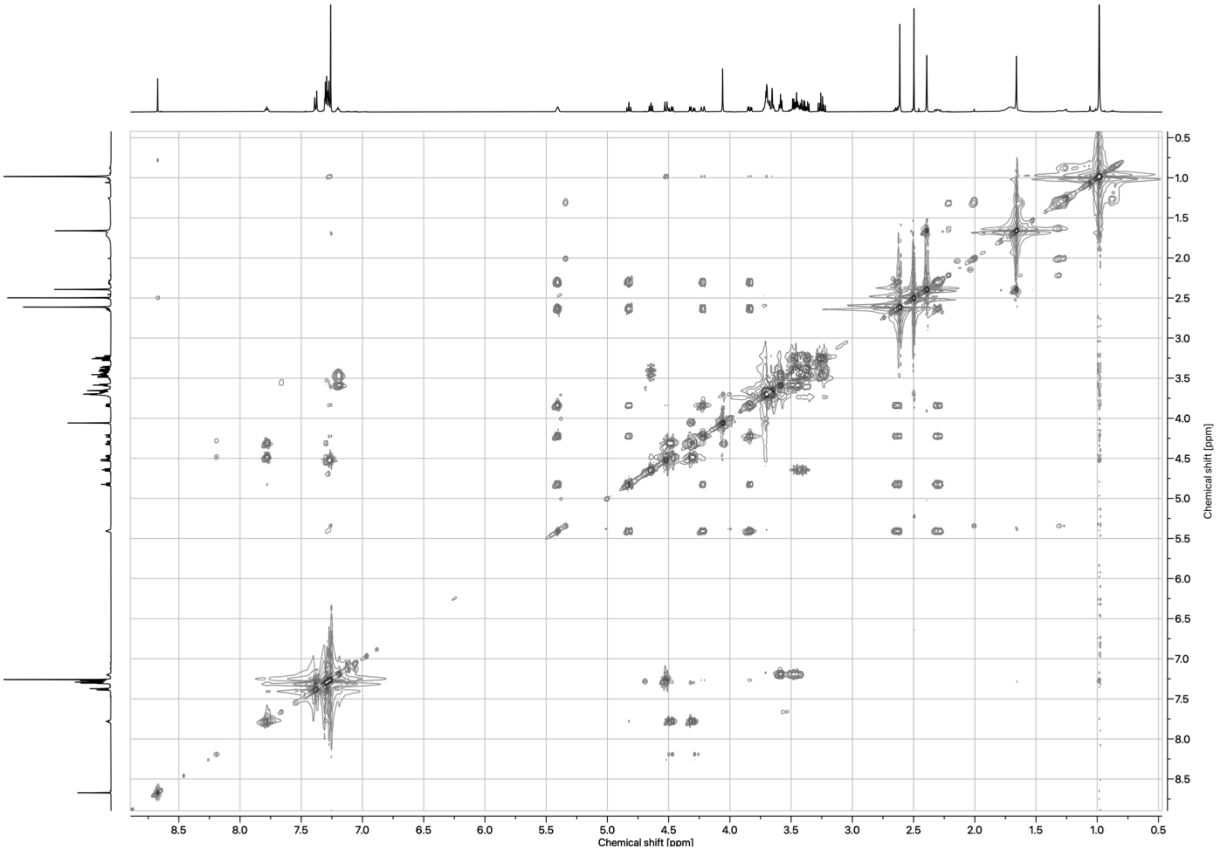

*Figure 22: TOCSY spectrum of MZ1-AspA ester (**13**) in $CDCl_3$*

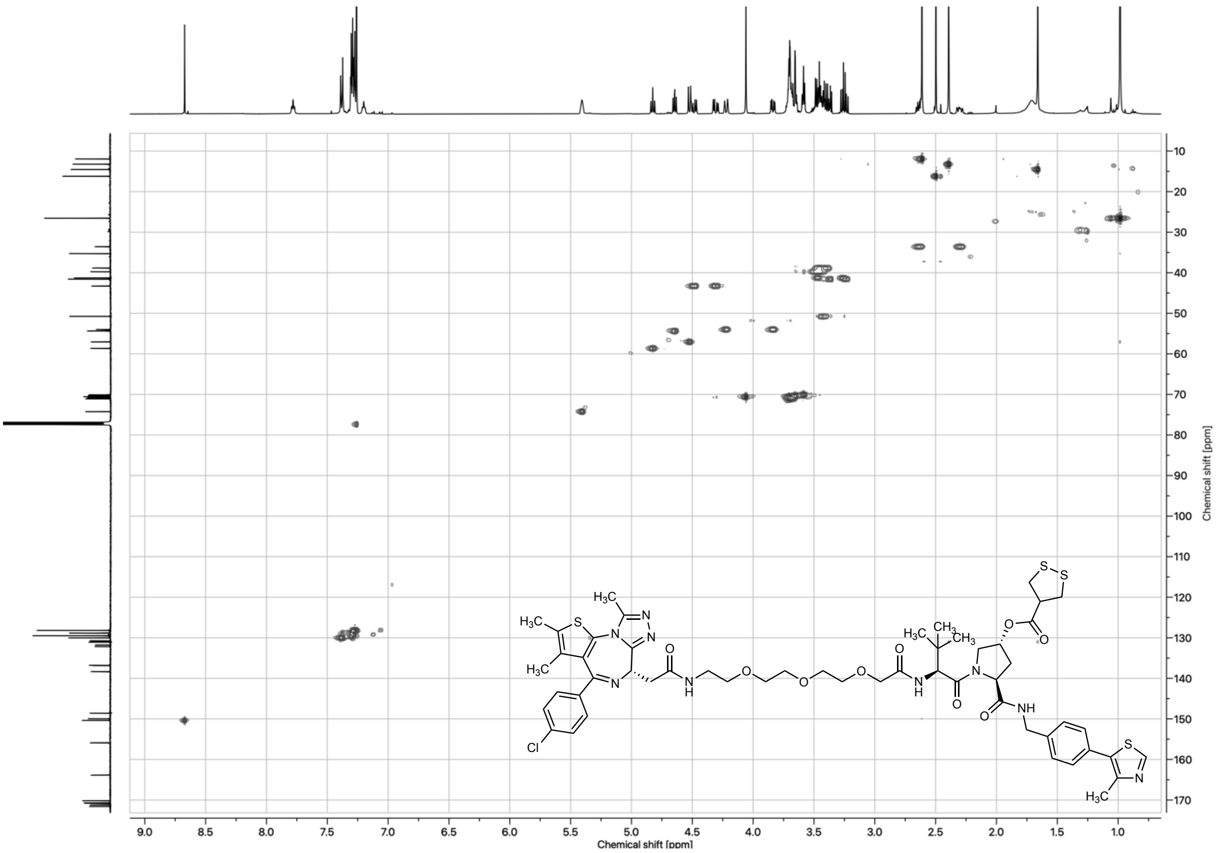

*Figure 23: HSQC spectrum of MZ1-AspA ester (**13**) in $CDCl_3$*

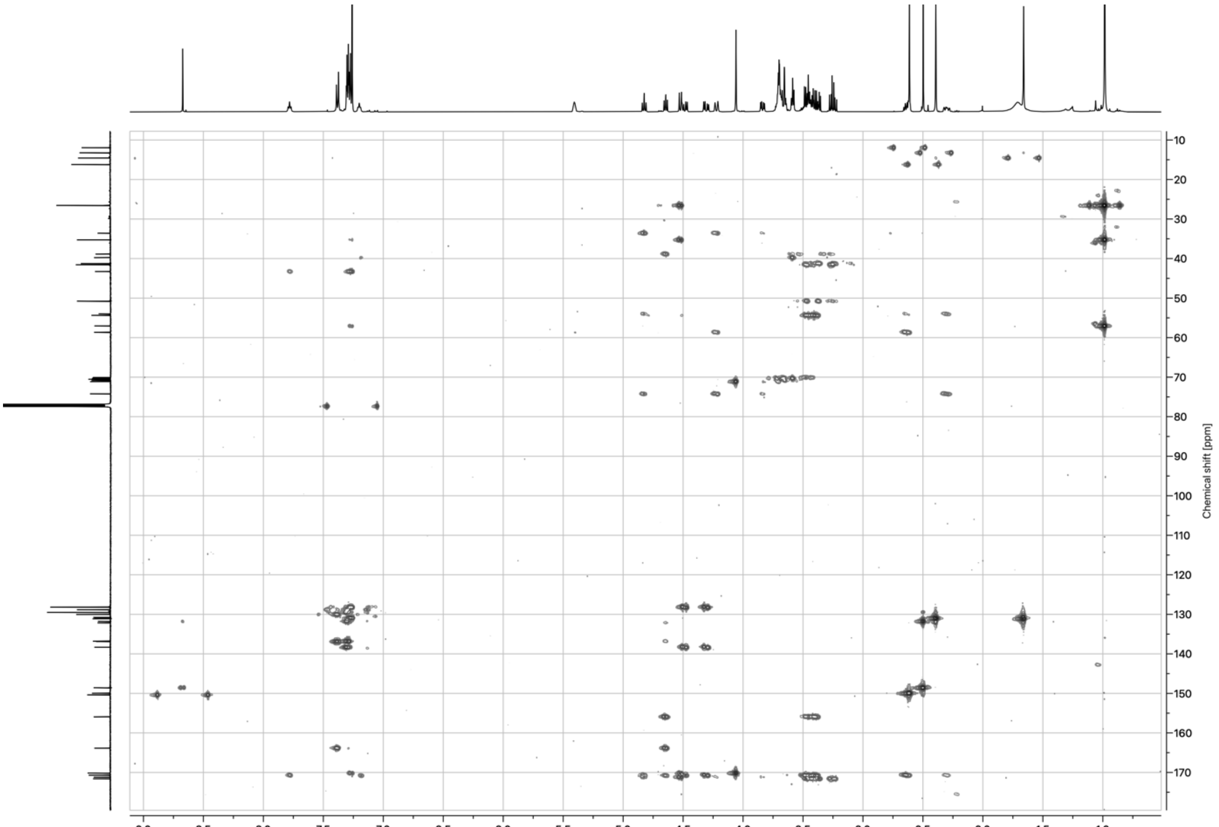

*Figure 24: HMBC spectrum of MZ1-AspA ester (**13**) in $CDCl_3$*

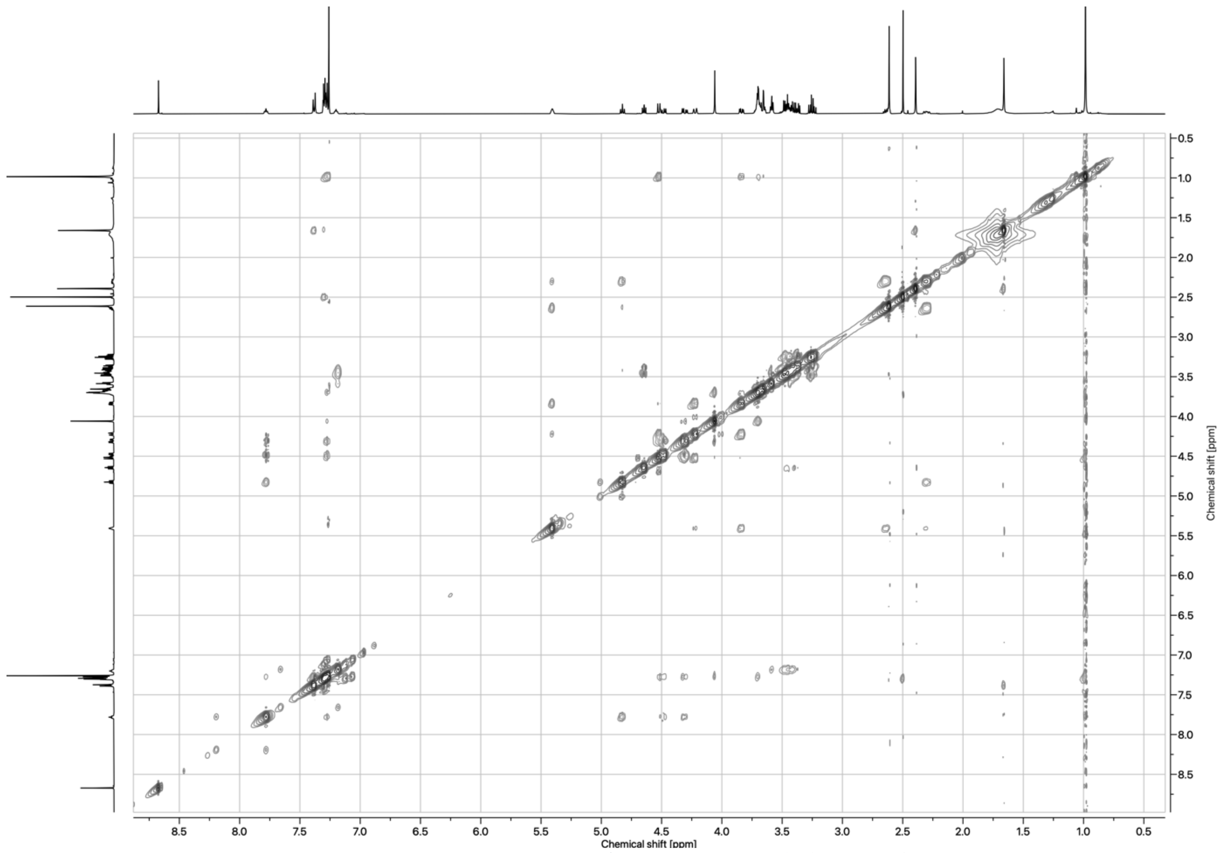

*Figure 25: NOESY spectrum of MZ1-AspA ester (**13**) in $CDCl_3$*

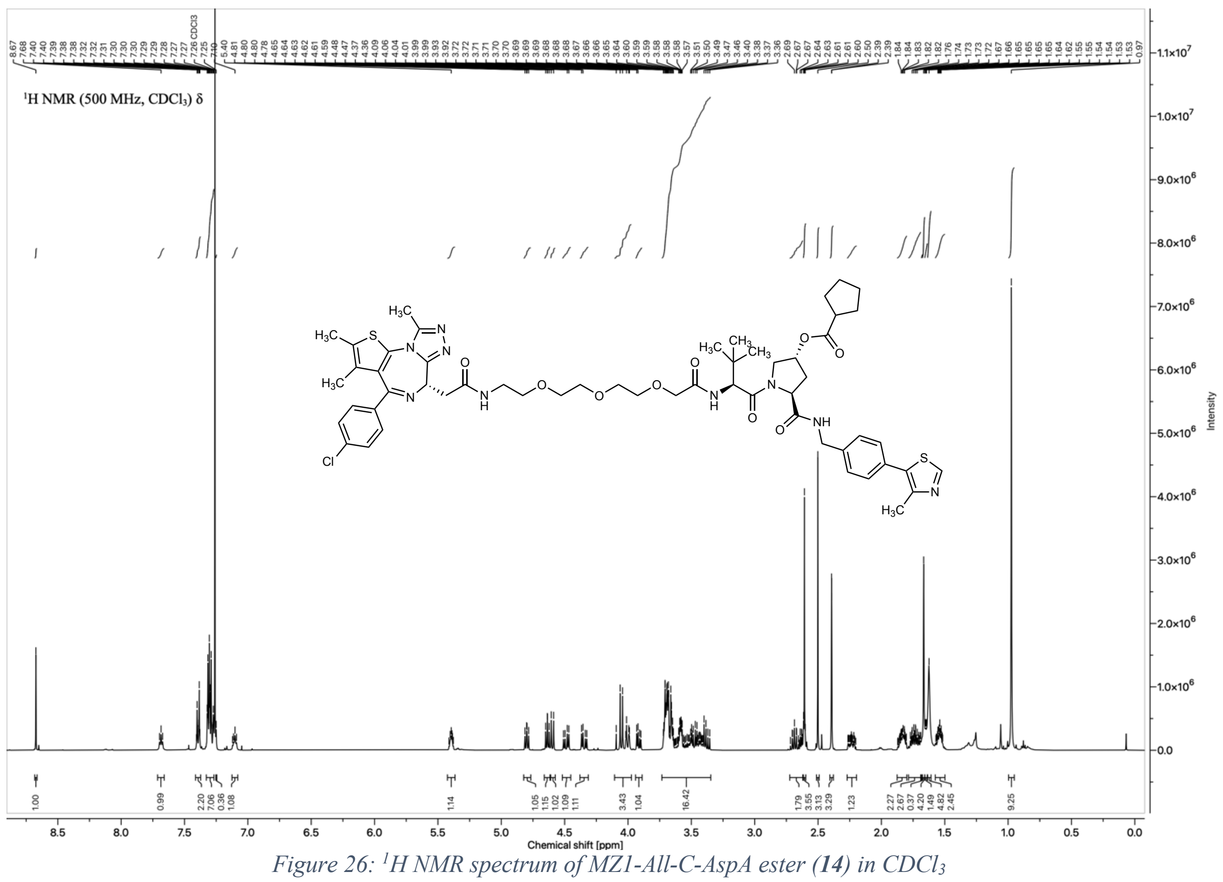

*Figure 26: $^1H$ NMR spectrum of MZ1-All-C-AspA ester (**14**) in $CDCl_3$*

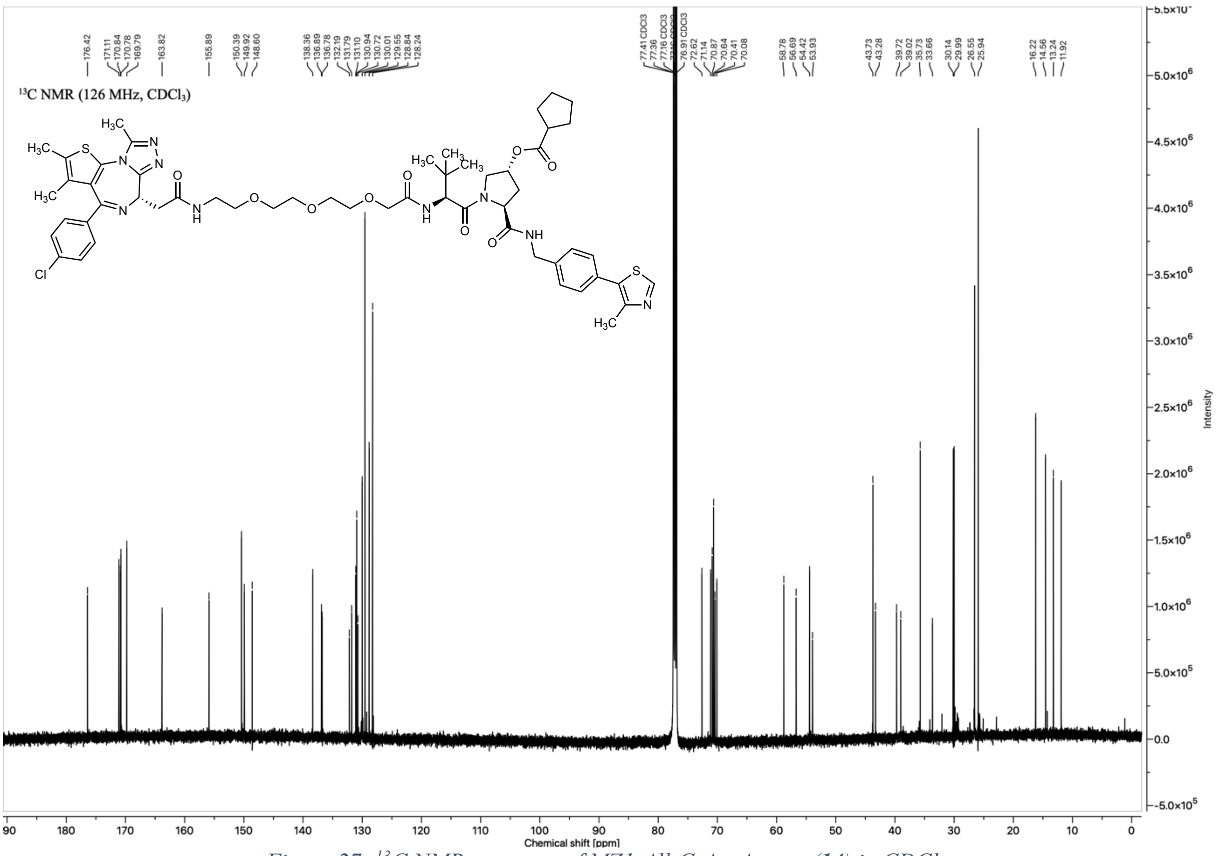

*Figure 27: $^{13}C$ NMR spectrum of MZ1-All-C-AspA ester (**14**) in $CDCl_3$*

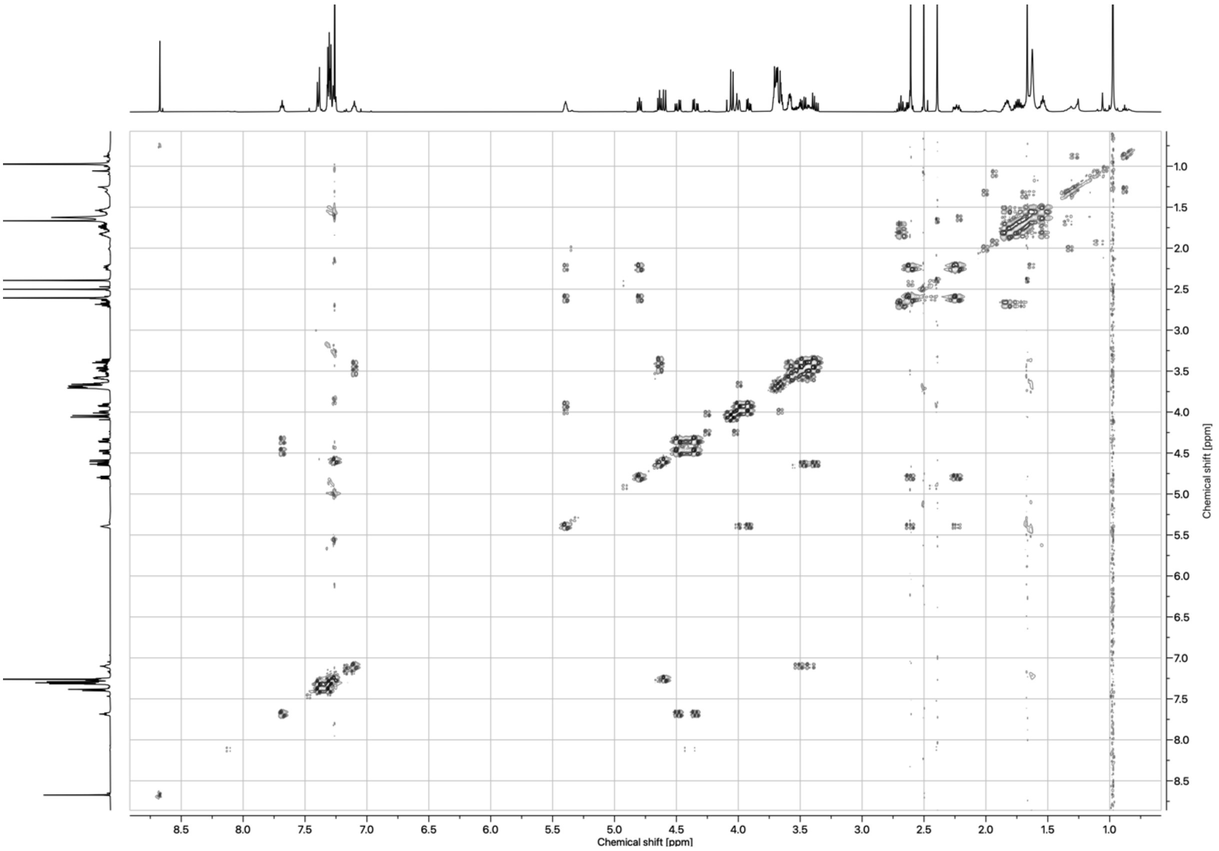

*Figure 28: COSY spectrum of MZ1-All-C-AspA ester (**14**) in $CDCl_3$*

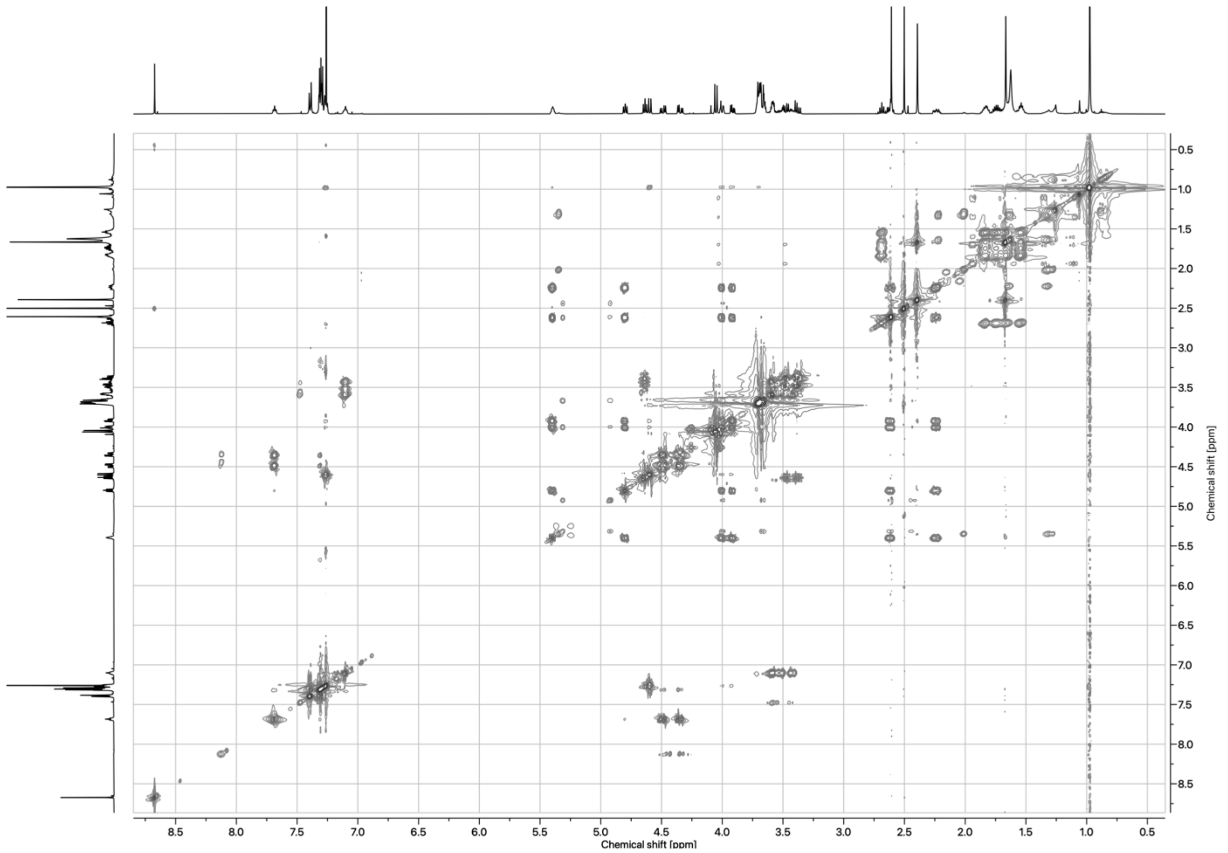

*Figure 29: TOCSY spectrum of MZ1-All-C-AspA ester (**14**) in $CDCl_3$*

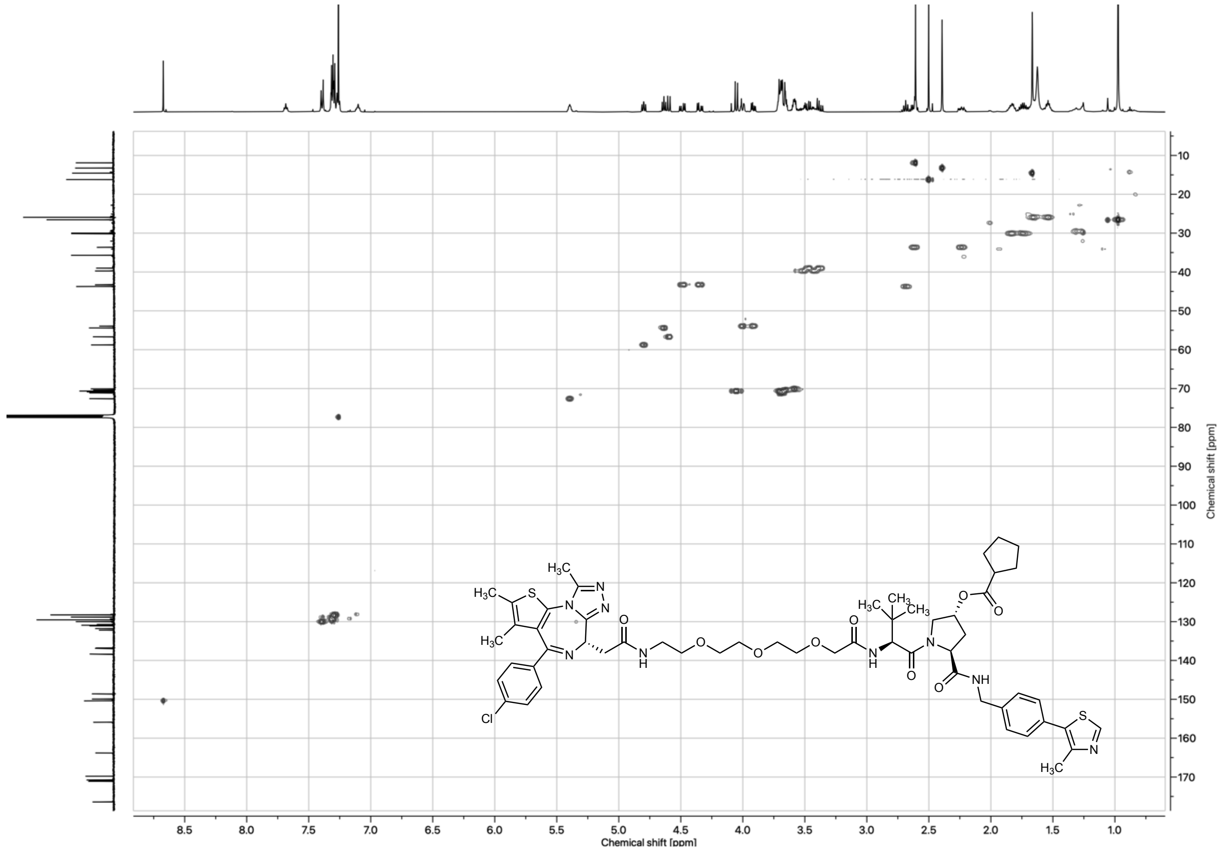

*Figure 30: HSQC spectrum of MZ1-All-C-AspA ester (**14**) in $CDCl_3$*

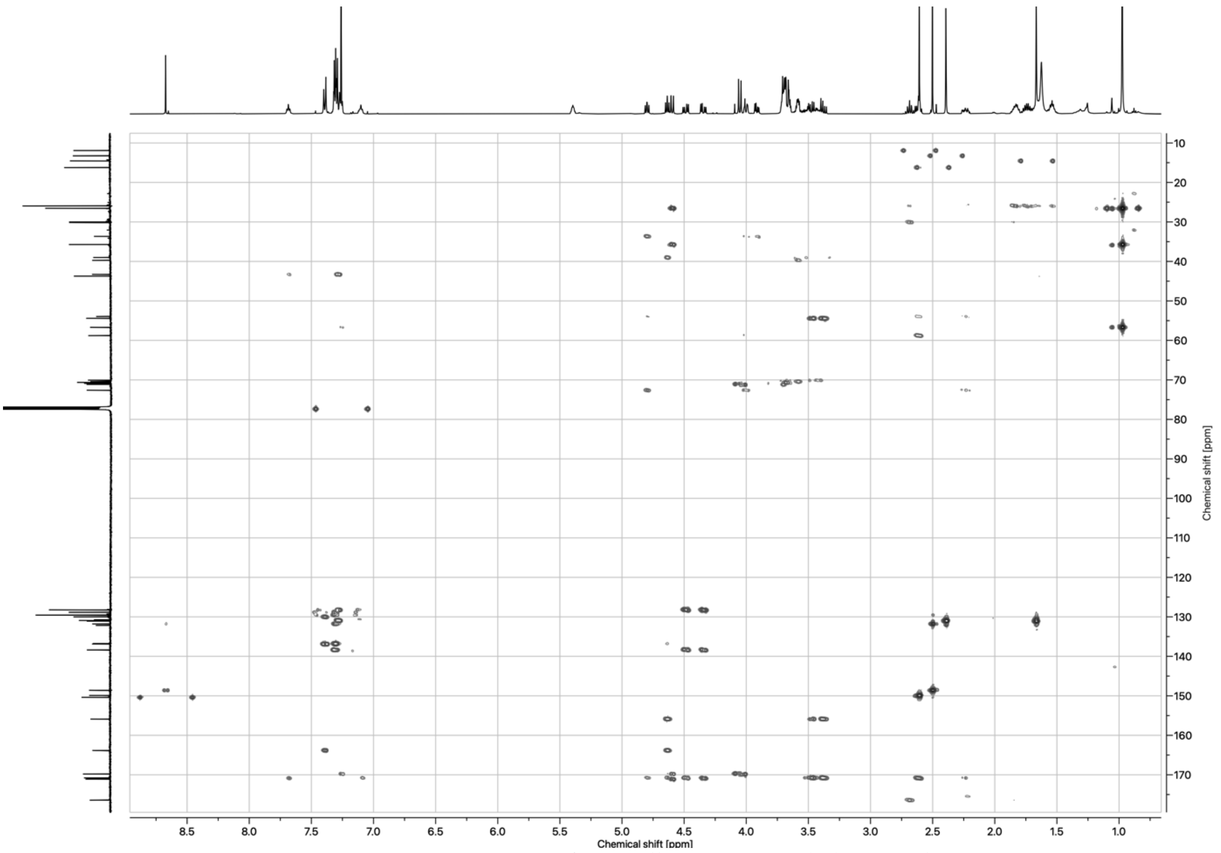

*Figure 31: HMBC spectrum of MZ1-All-C-AspA ester (**14**) in $CDCl_3$*

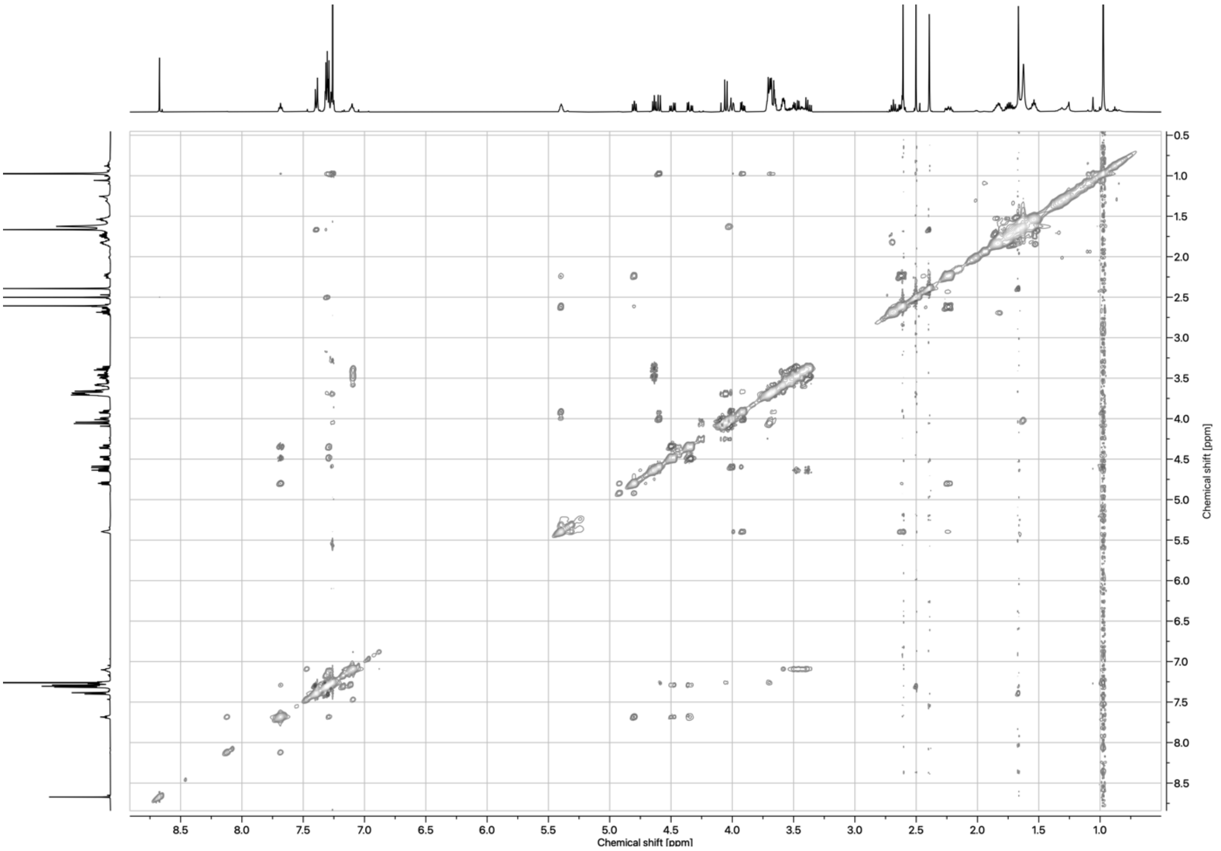

*Figure 32: NOESY spectrum of MZ1-All-C-AspA ester (**14**) in $CDCl_3$*

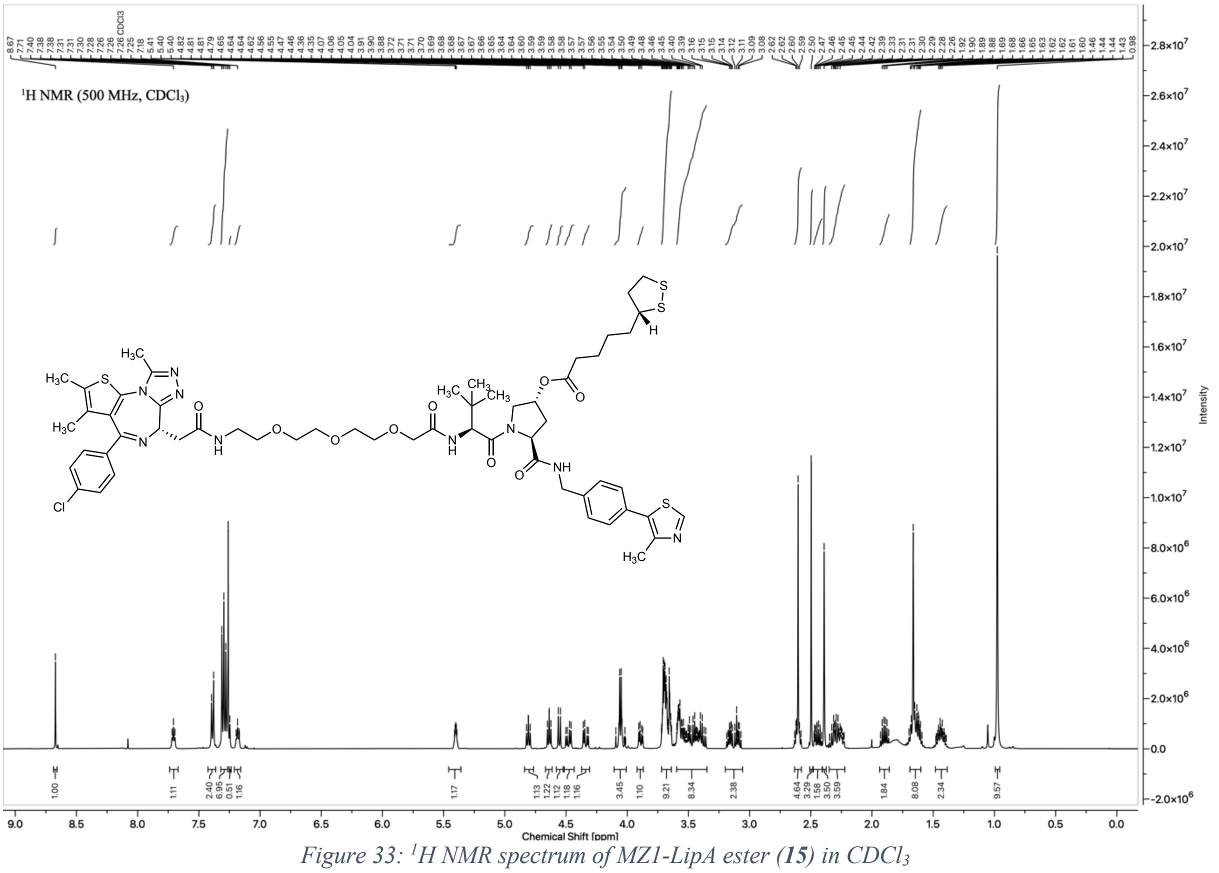

*Figure 33: $^{1}$H NMR spectrum of MZ1-LipA ester (**15**) in $CDCl_3$*

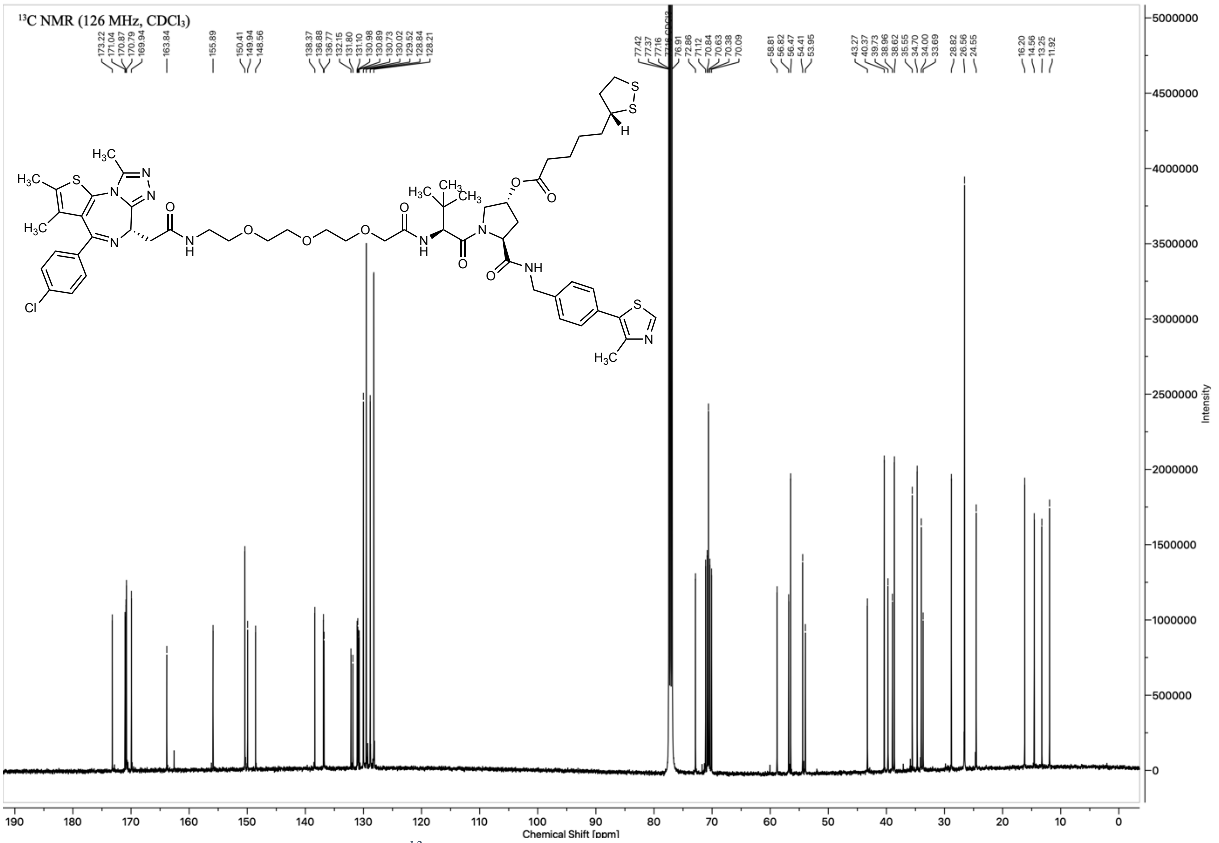

*Figure 34: $^{13}$C NMR spectrum of MZ1-LipA ester (**15**) in $CDCl_3$*

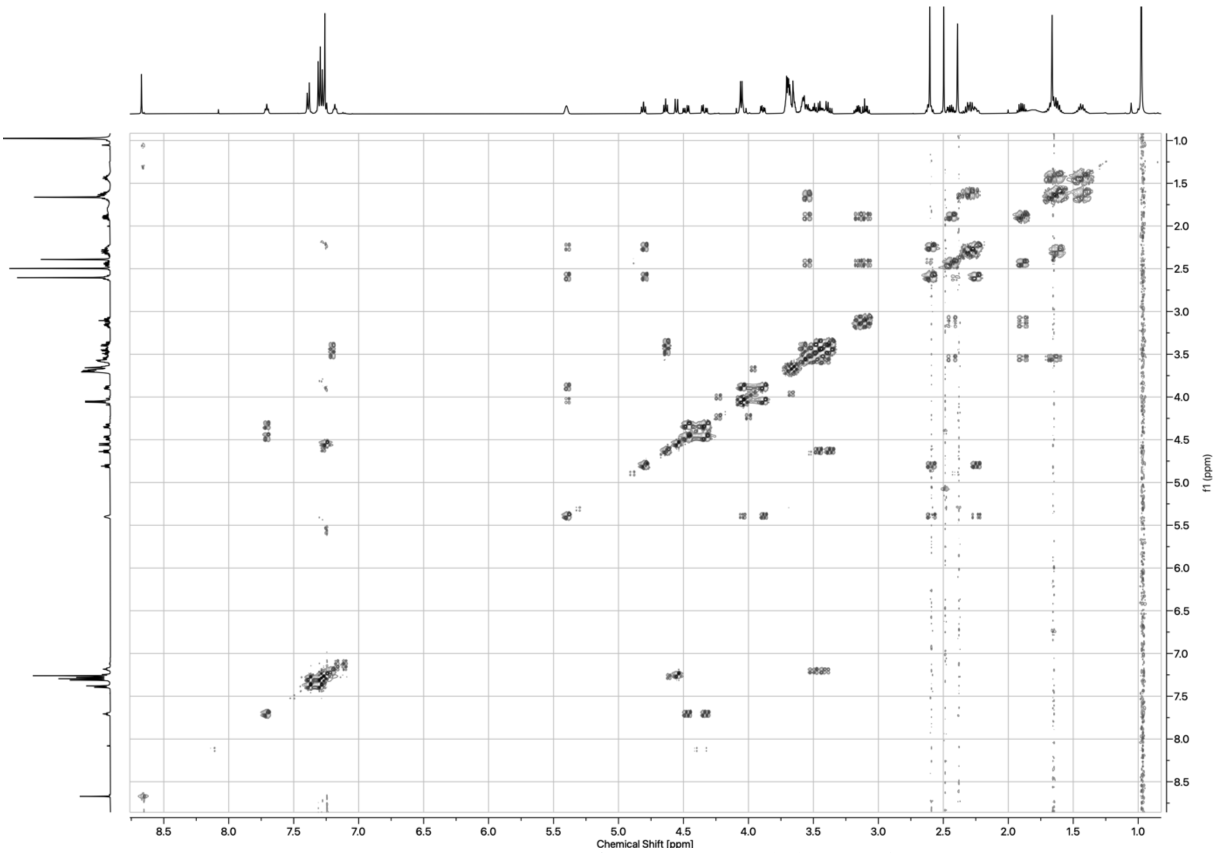

*Figure 35: COSY spectrum of MZ1-LipA ester (**15**) in $CDCl_3$*

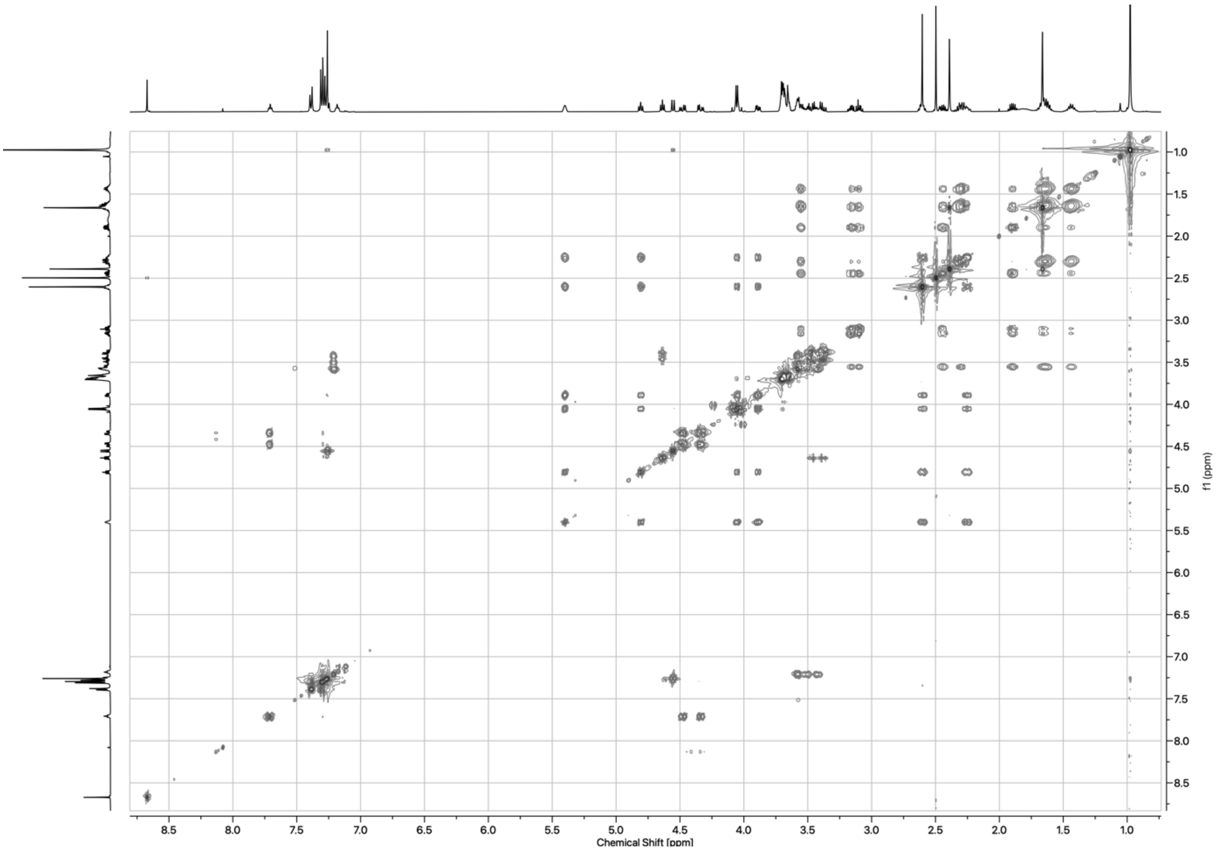

*Figure 36: TOCSY spectrum of MZ1-LipA ester (**15**) in $CDCl_3$*

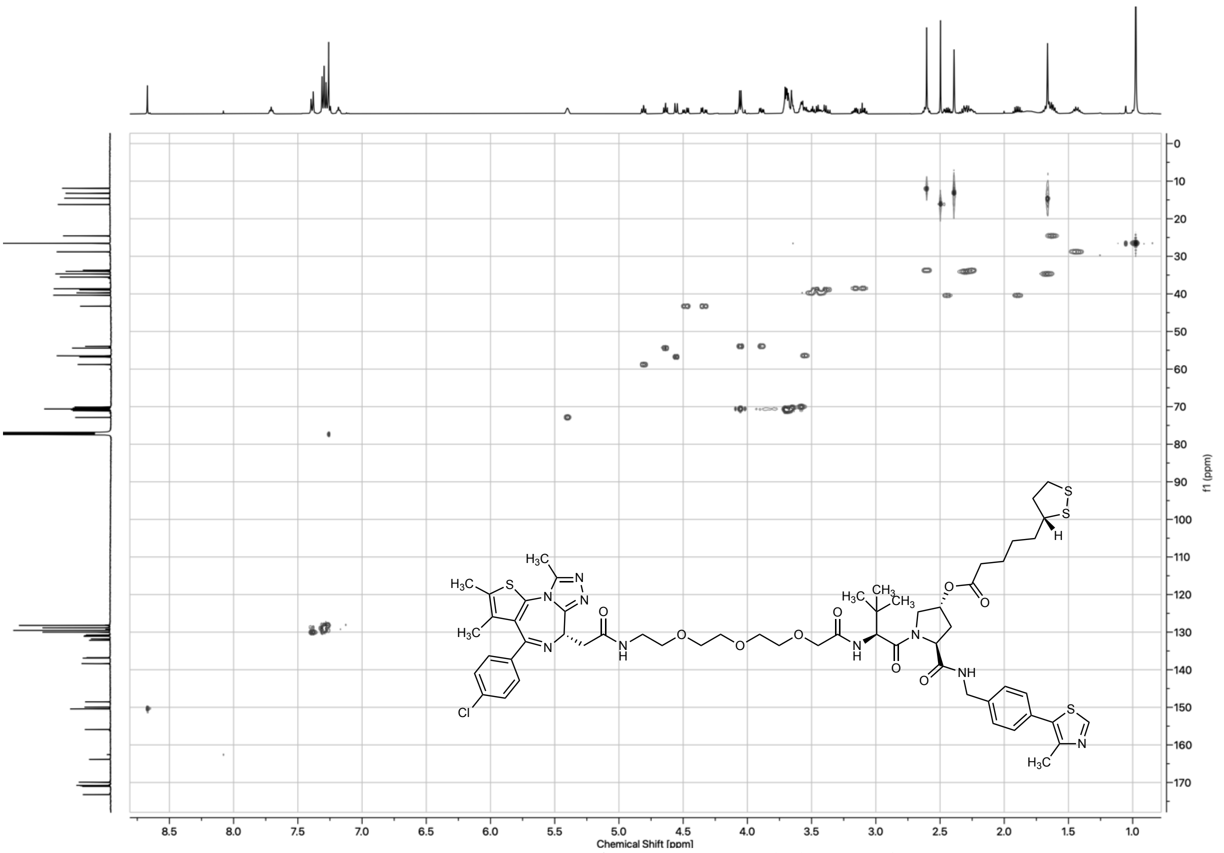

*Figure 37: HSQC spectrum of MZ1-LipA ester (**15**) in $CDCl_3$*

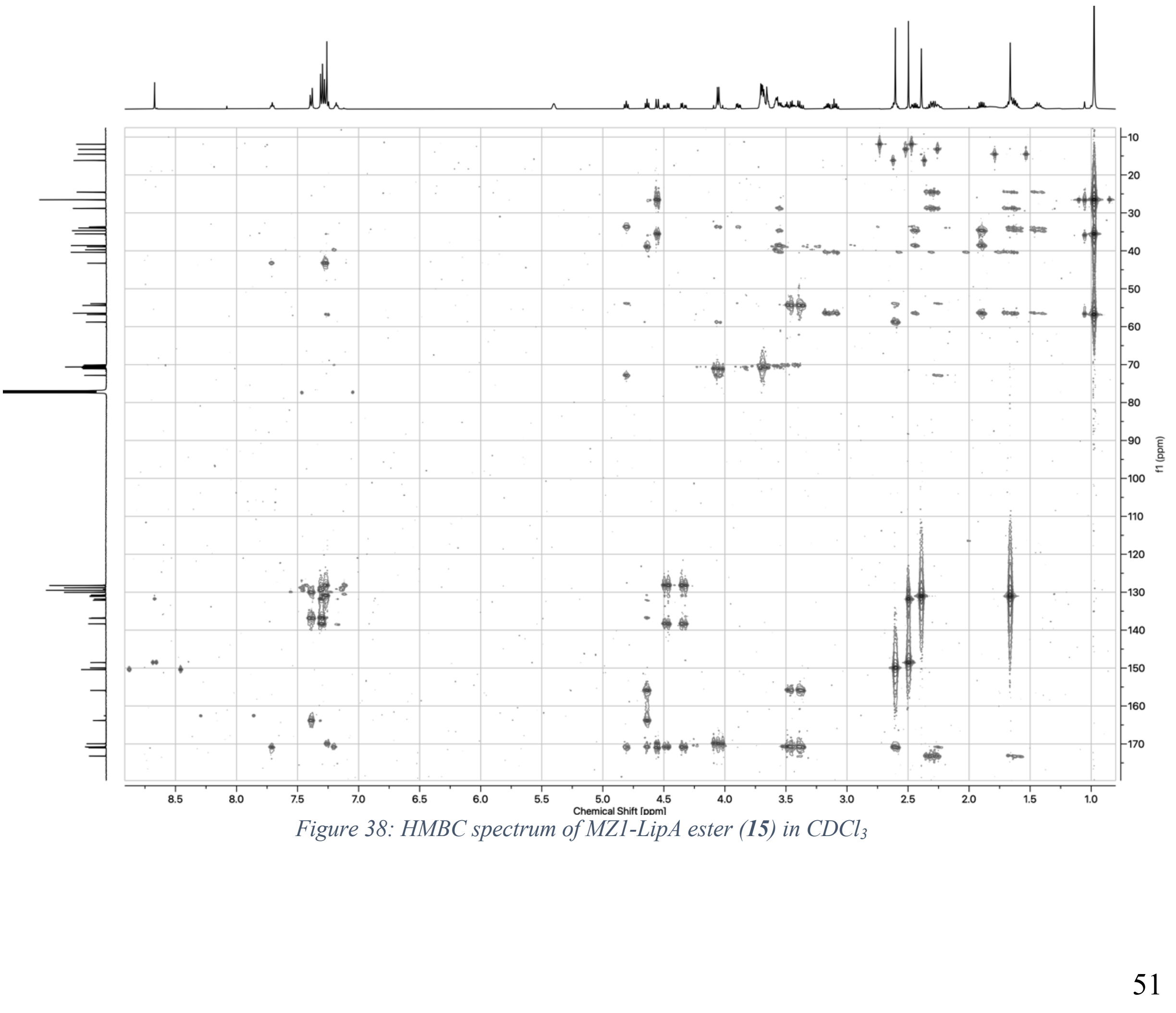

*Figure 38: HMBC spectrum of MZ1-LipA ester (**15**) in $CDCl_3$*

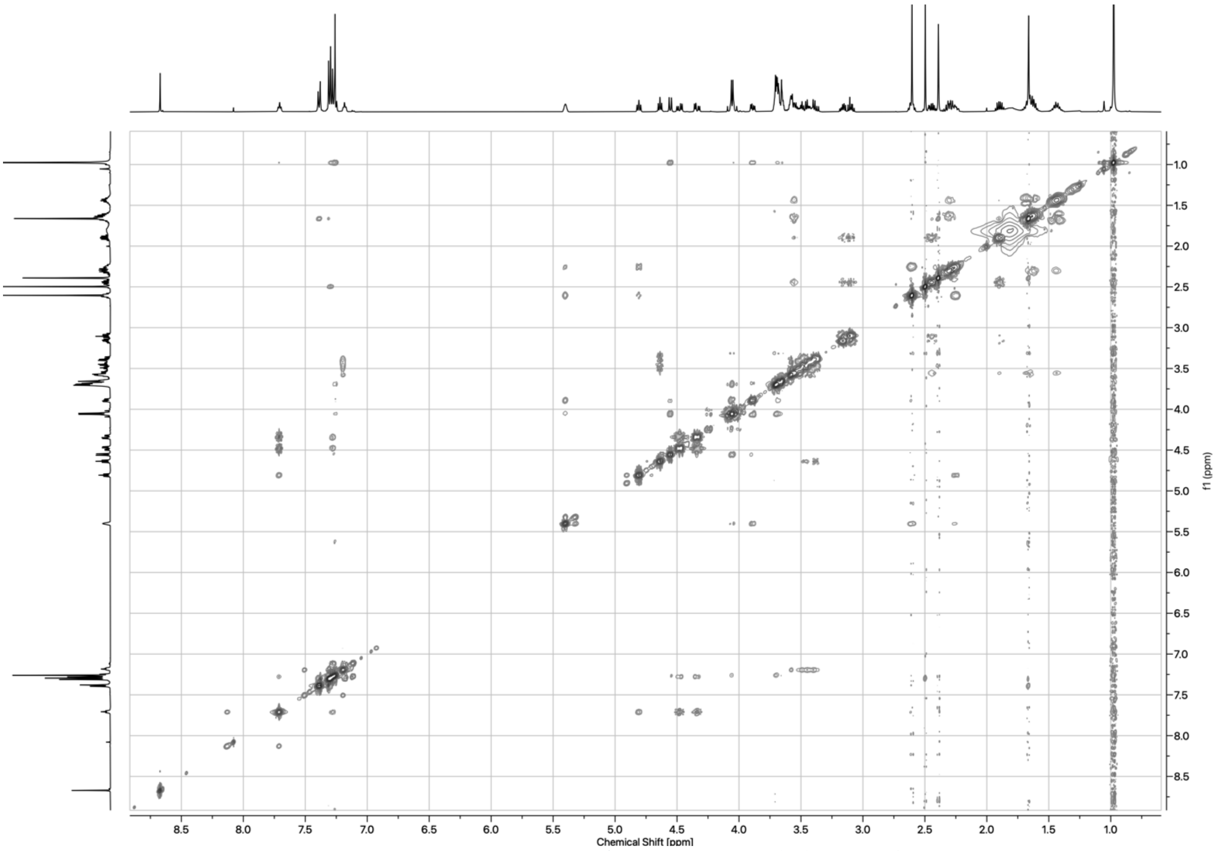

*Figure 39: NOESY spectrum of MZ1-LipA ester (**15**) in $CDCl_3$*

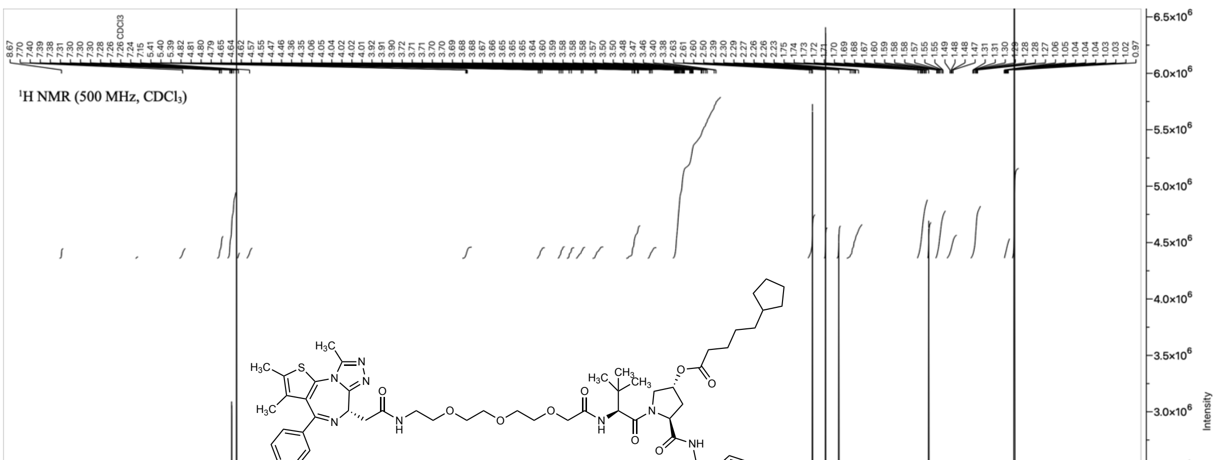

*Figure 40: $^{1}H$ NMR spectrum of MZ1-All-C-LipA ester (**16**) in $CDCl_3$*

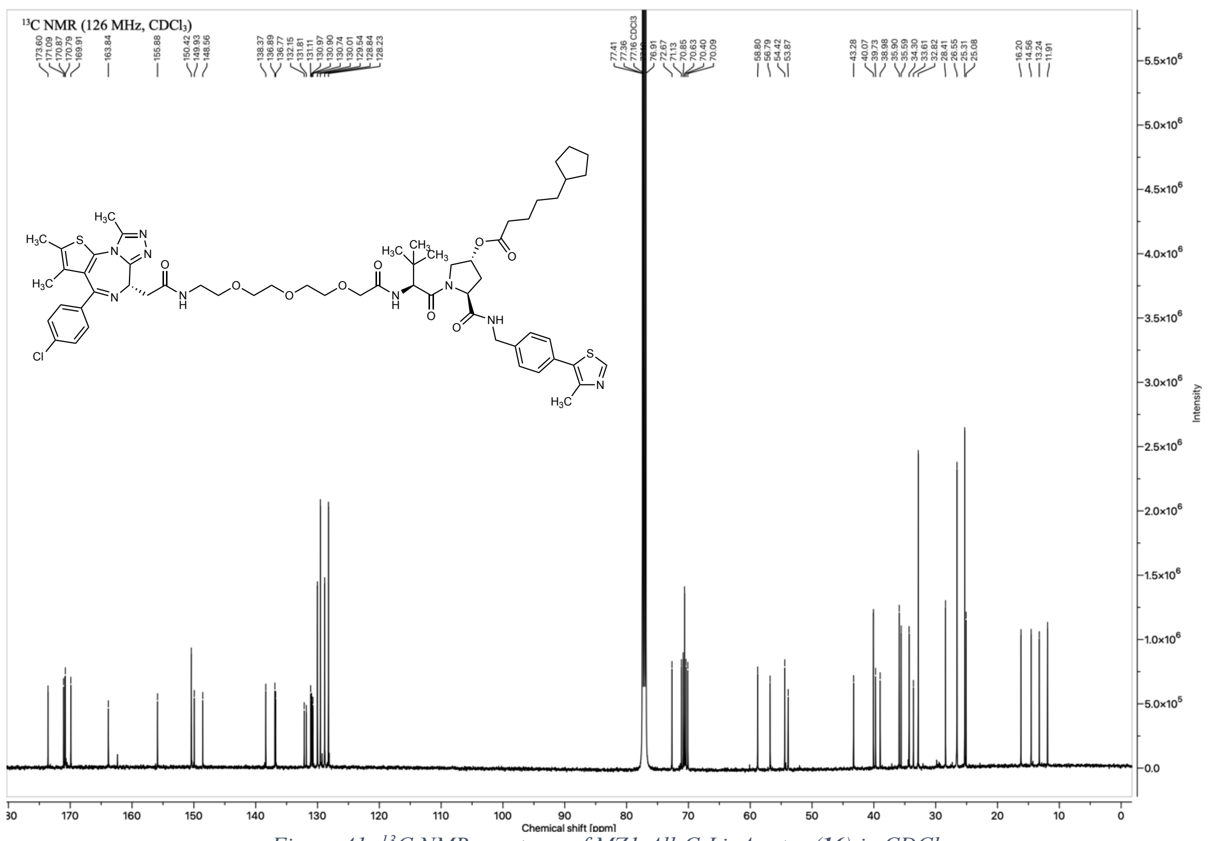

*Figure 41: $^{13}$C NMR spectrum of MZ1-All-C-LipA ester (**16**) in $CDCl_3$*

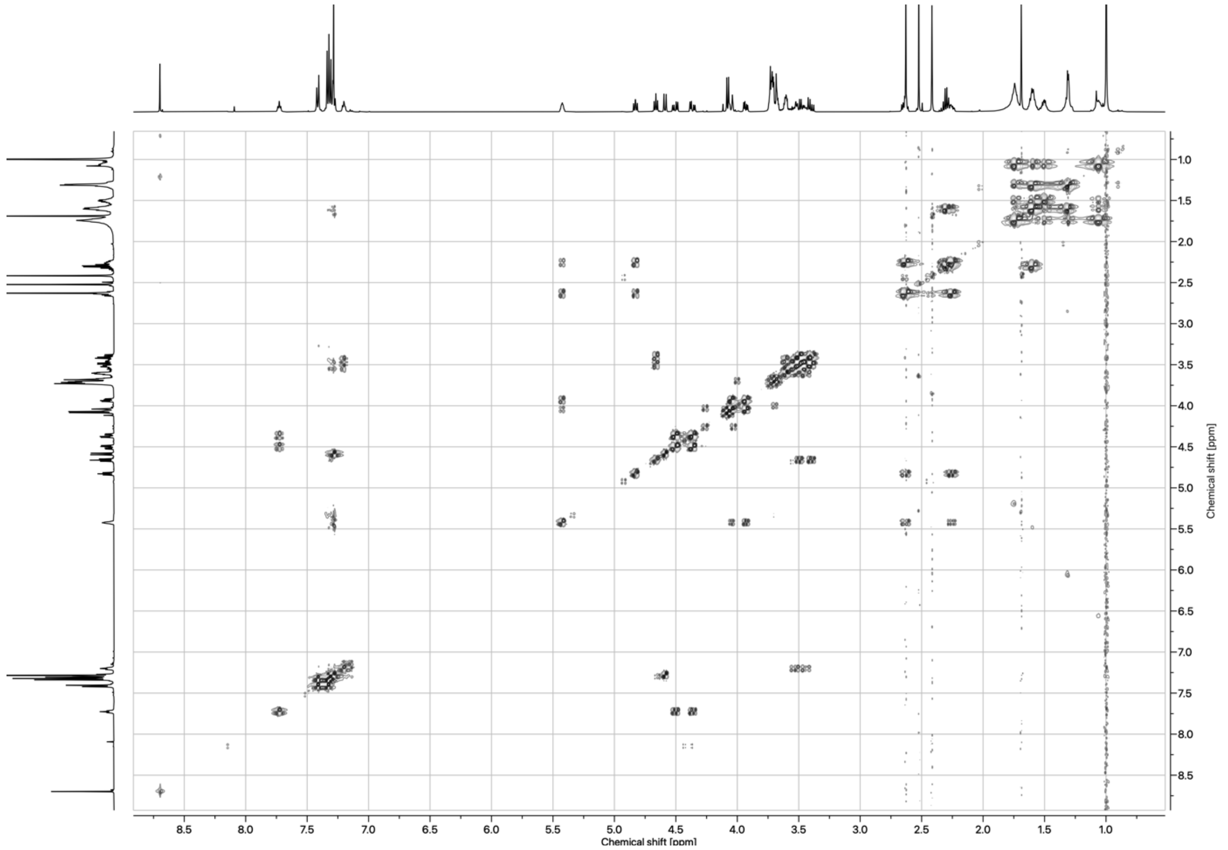

*Figure 42: COSY spectrum of MZ1-All-C-LipA ester (**16**) in $CDCl_3$*

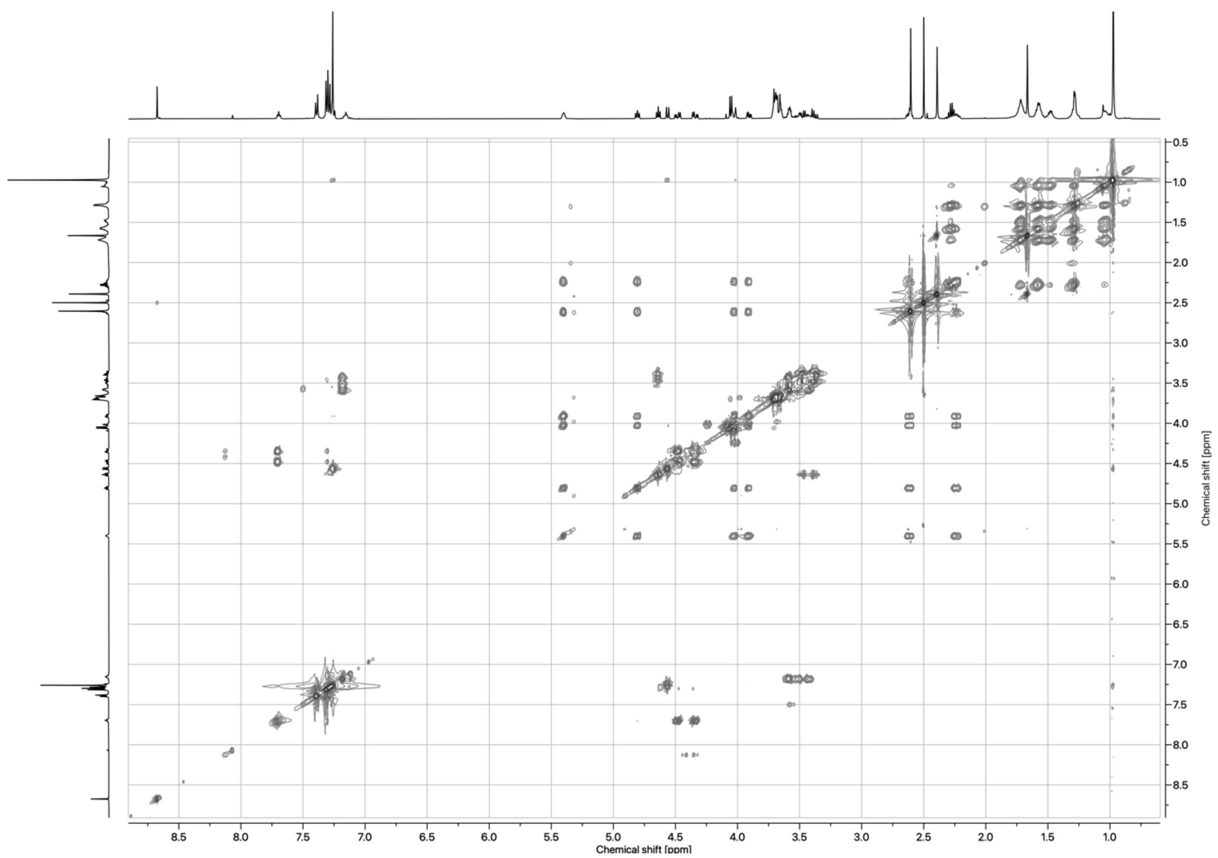

*Figure 43: TOCSY spectrum of MZ1-All-C-LipA ester (**16**) in $CDCl_3$*

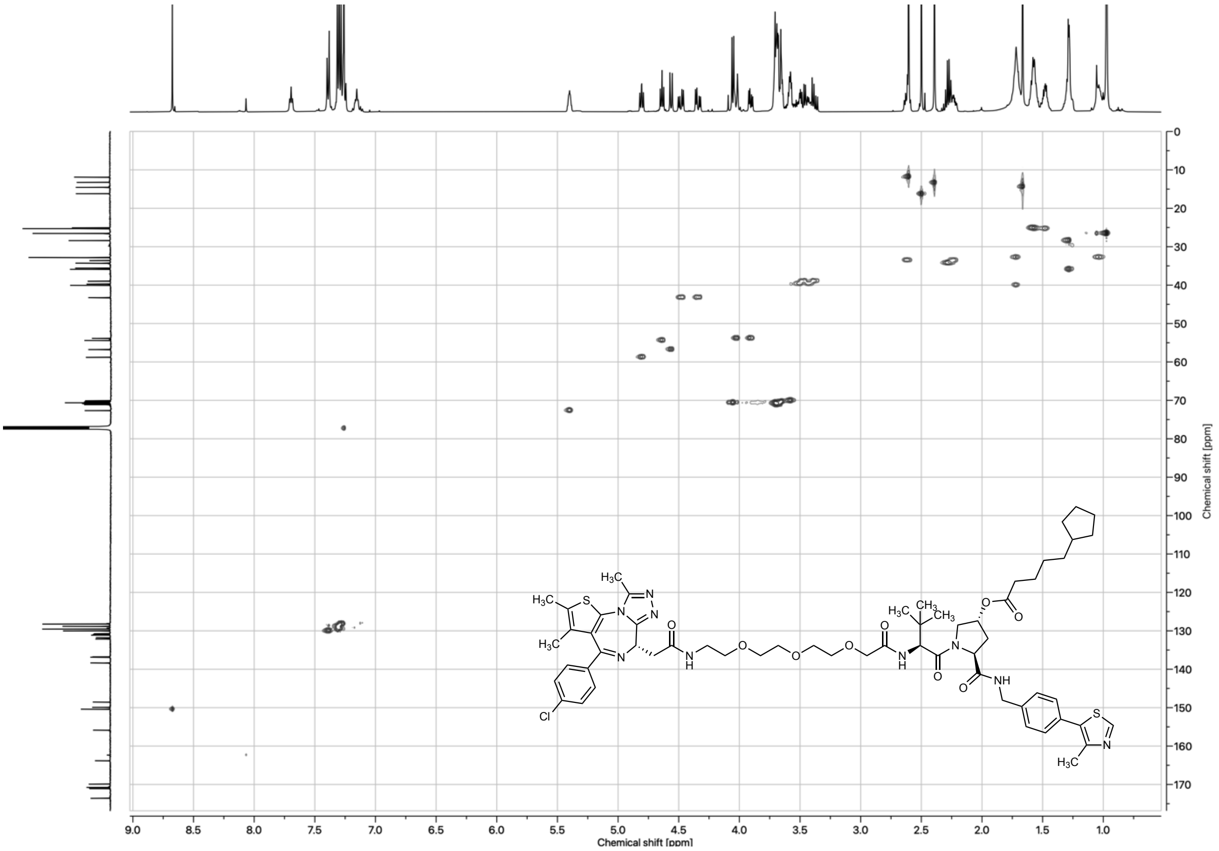

*Figure 44: HSQC spectrum of MZ1-All-C-LipA ester (**16**) in $CDCl_3$*

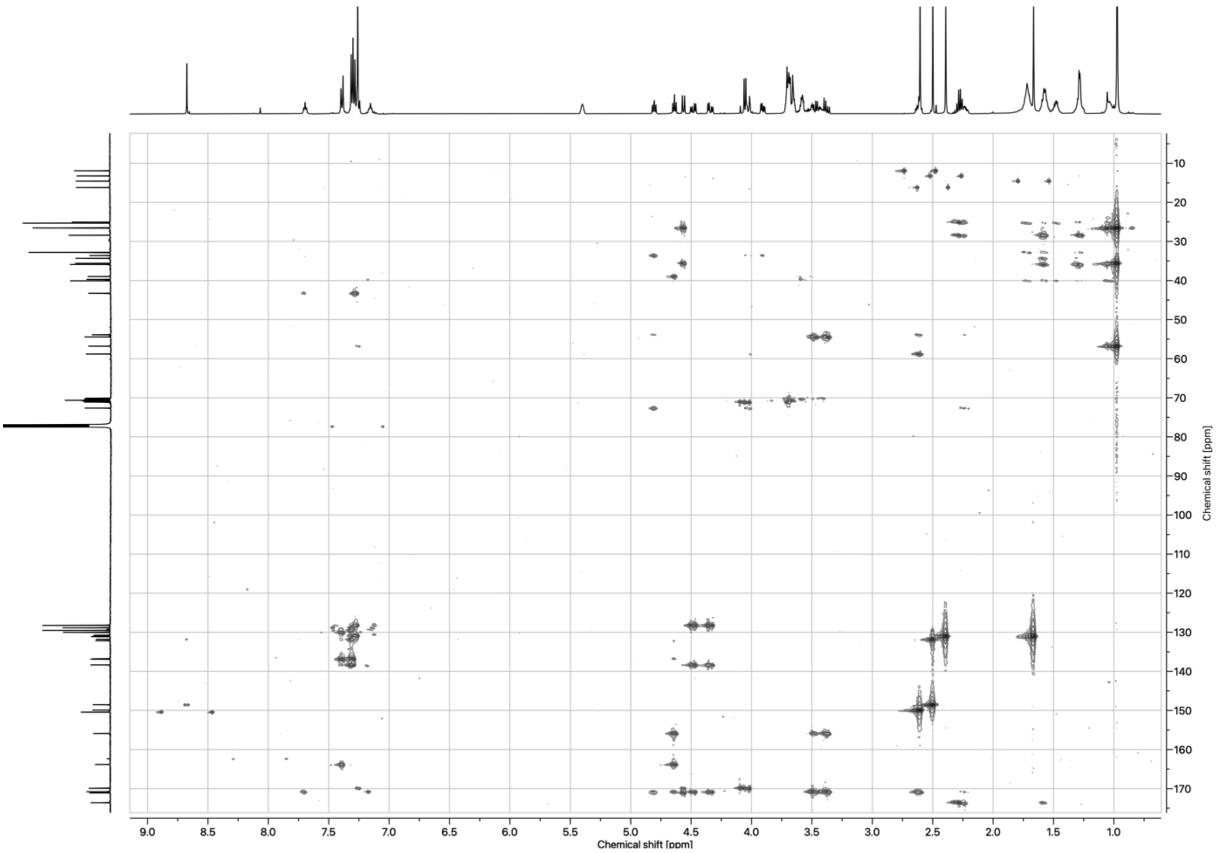

*Figure 45: HMBC spectrum of MZ1-All-C-LipA ester (**16**) in $CDCl_3$*

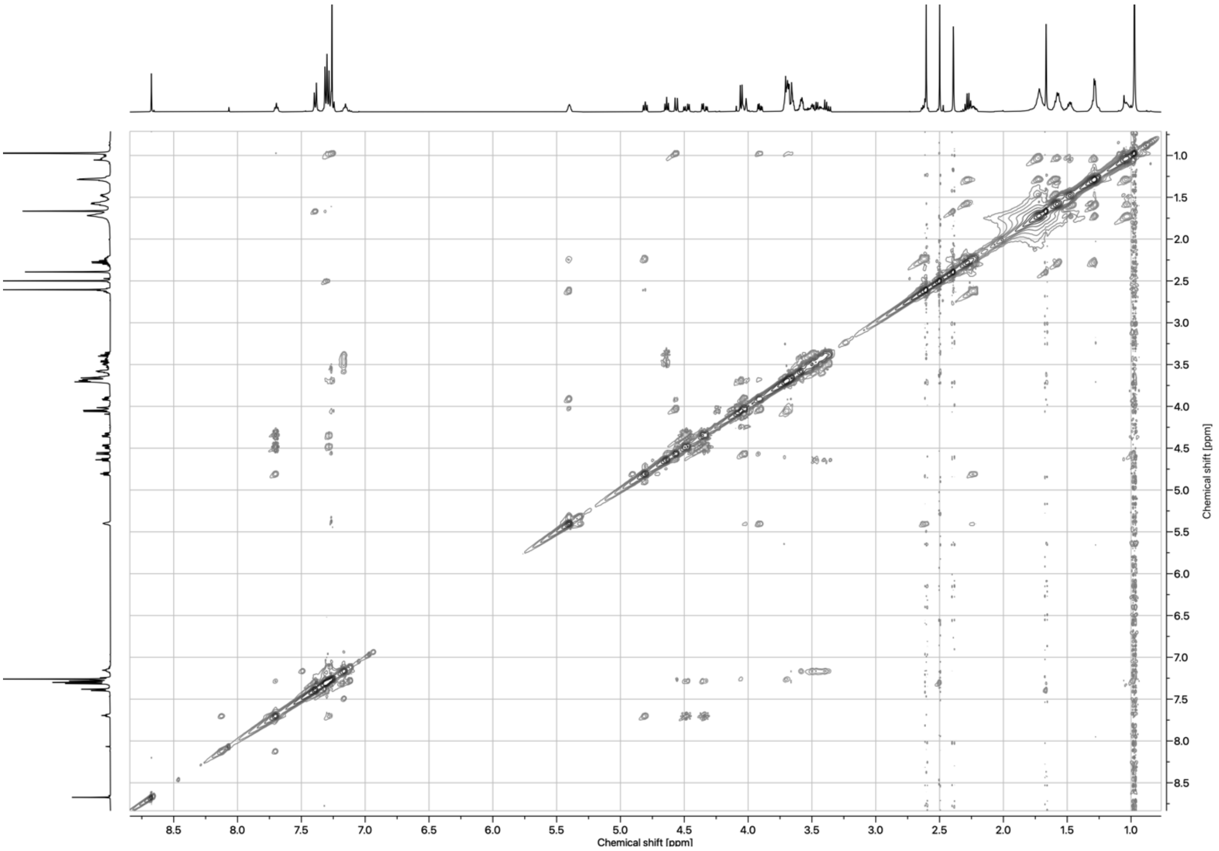

*Figure 46: NOESY spectrum of MZ1-All-C-LipA ester (**16**) in $CDCl_3$*

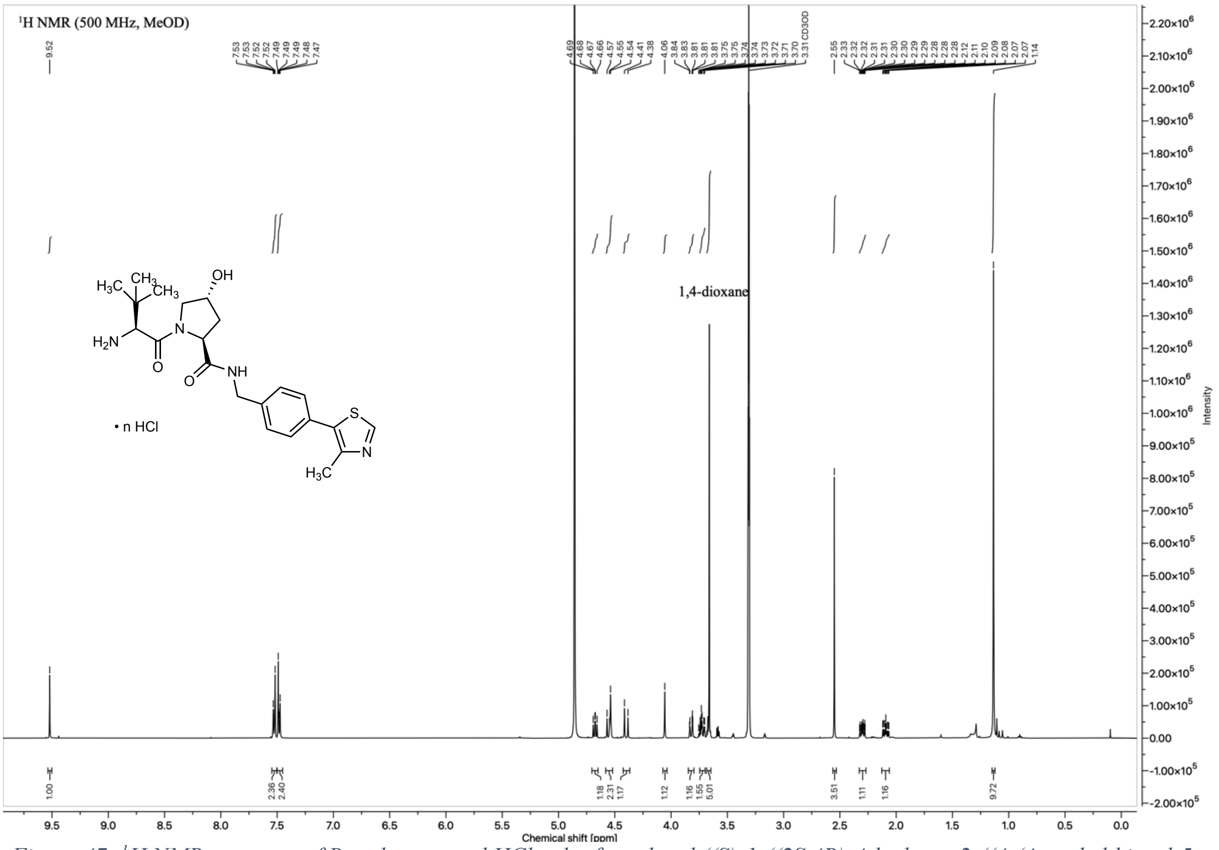

*Figure 47: $^{1}H$ NMR spectrum of Boc-deprotected HCl-salt of tert-butyl ((S)-1-((2S,4R)-4-hydroxy-2-((4-(4-methylthiazol-5-yl)benzyl)carbamoyl)pyrrolidin-1-yl)-3,3-dimethyl-1-oxobutan-2-yl)carbamate (**3**, (S,R,S)-AHPC•HCl salt) in $CD_3OD$*

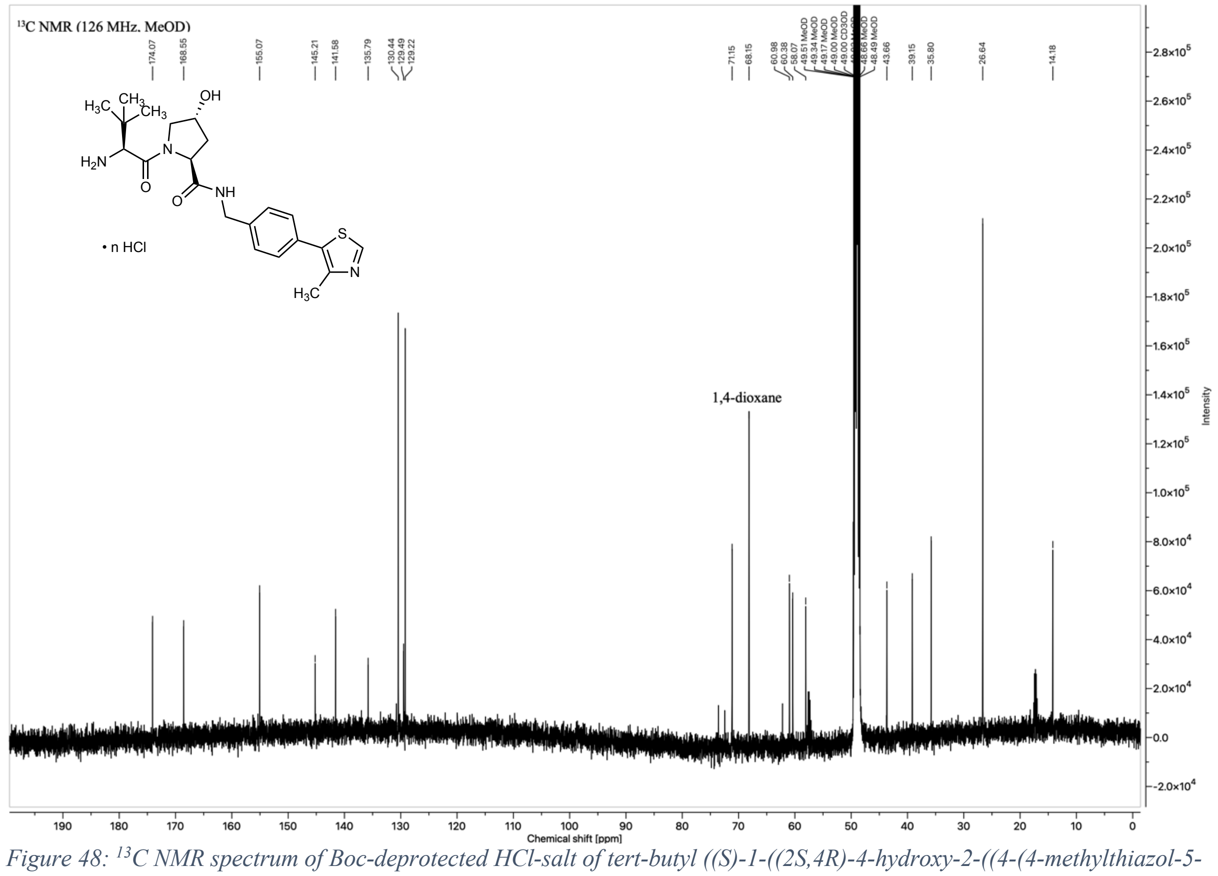

*Figure 48: $^{13}C$ NMR spectrum of Boc-deprotected HCl-salt of tert-butyl ((S)-1-((2S,4R)-4-hydroxy-2-((4-(4-methylthiazol-5-yl)benzyl)carbamoyl)pyrrolidin-1-yl)-3,3-dimethyl-1-oxobutan-2-yl)carbamate (**3**, (S,R,S)-AHPC•HCl salt) in $CD_3OD$*

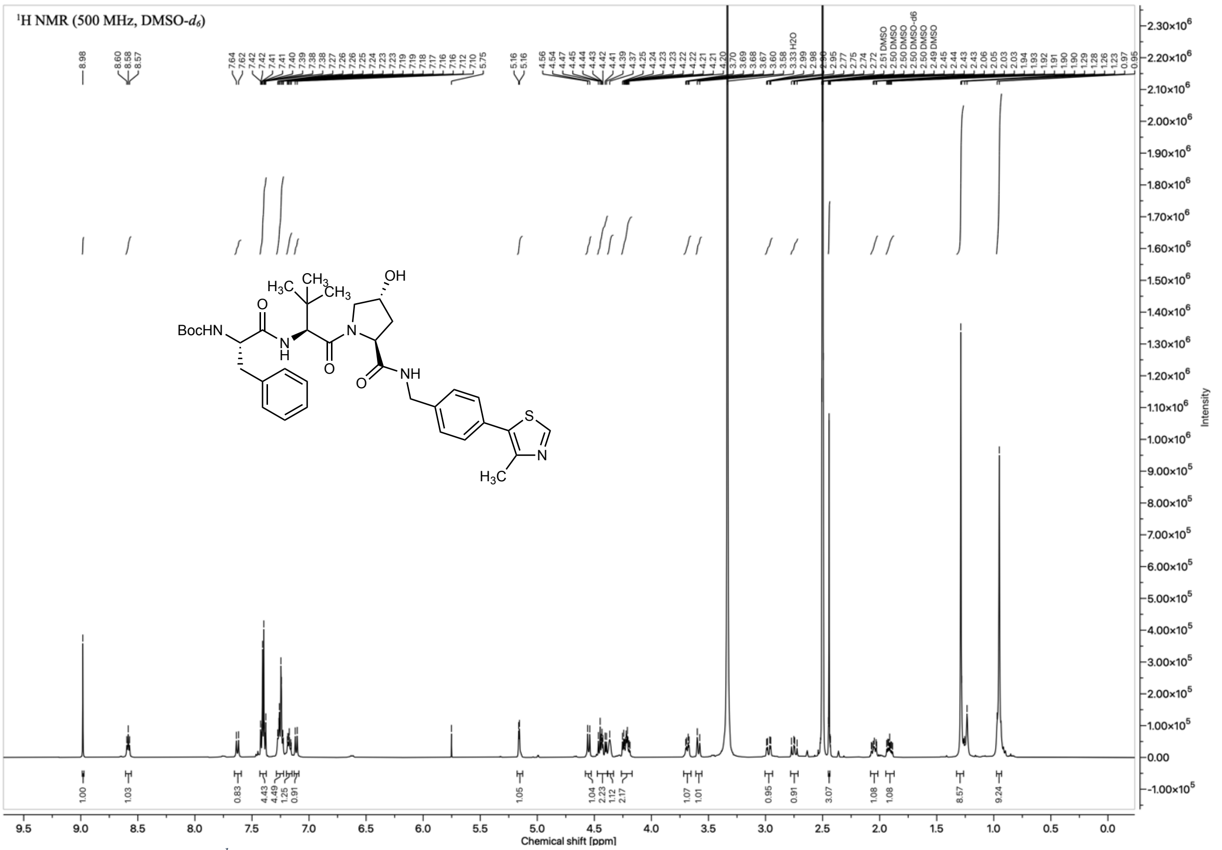

*Figure 49: $^{1}$H NMR spectrum of tert-butyl ((S)-1-(((S)-1-((2S,4R)-4-hydroxy-2-((4-(4-methylthiazol-5-yl)benzyl)carbamoyl)pyrrolidin-1-yl)-3,3-dimethyl-1-oxobutan-2-yl)amino)-1-oxo-3-phenylpropan-2-yl)carbamate (**17**) in DMSO-$d_6$*

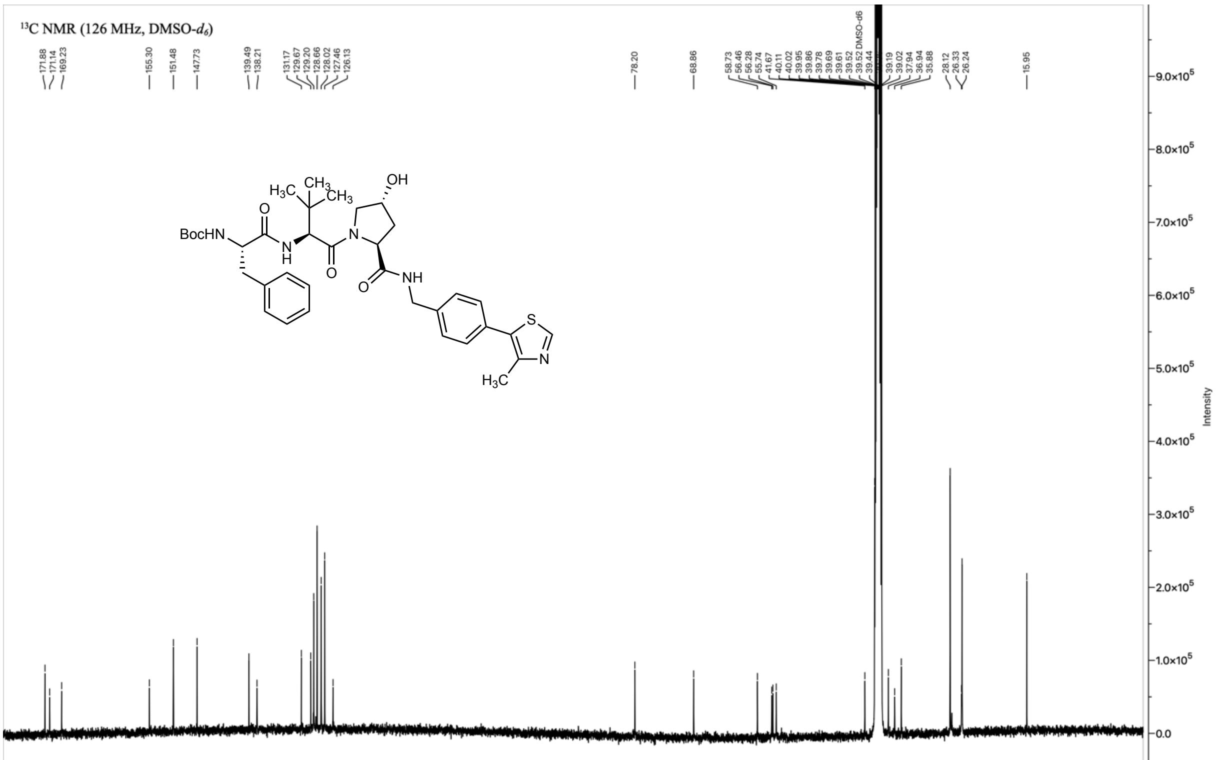

*Figure 50: $^{13}$C NMR spectrum of tert-butyl ((S)-1-(((S)-1-((2S,4R)-4-hydroxy-2-((4-(4-methylthiazol-5-yl)benzyl)carbamoyl)pyrrolidin-1-yl)-3,3-dimethyl-1-oxobutan-2-yl)amino)-1-oxo-3-phenylpropan-2-yl)carbamate (**17**) in DMSO-$d_6$*

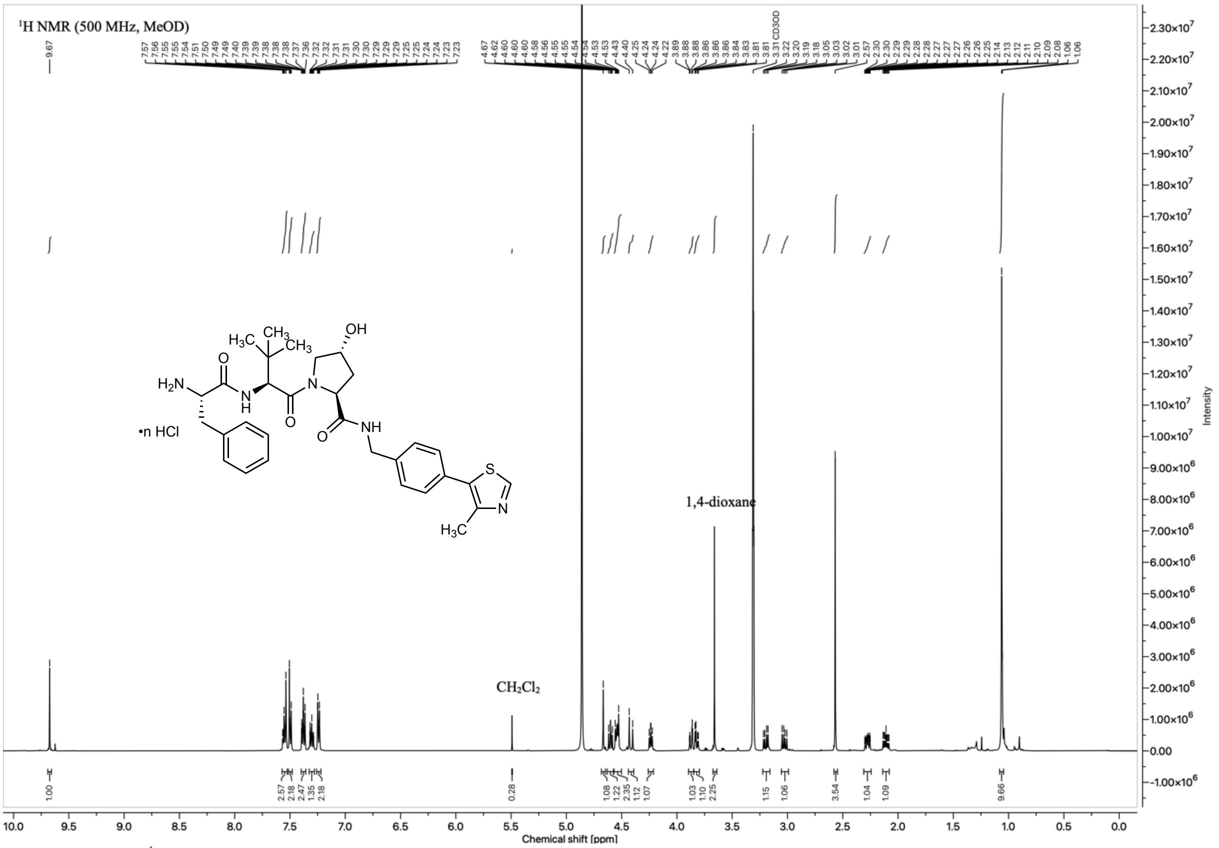

*Figure 51: $^1$H NMR spectrum of the Boc-deprotected HCl-salt of tert-butyl ((S)-1-(((S)-1-((2S,4R)-4-hydroxy-2-((4-(4-methylthiazol-5-yl)benzyl)carbamoyl)pyrrolidin-1-yl)-3,3-dimethyl-1-oxobutan-2-yl)amino)-1-oxo-3-phenylpropan-2-yl)carbamate (**18**) in $CD_3OD$.*

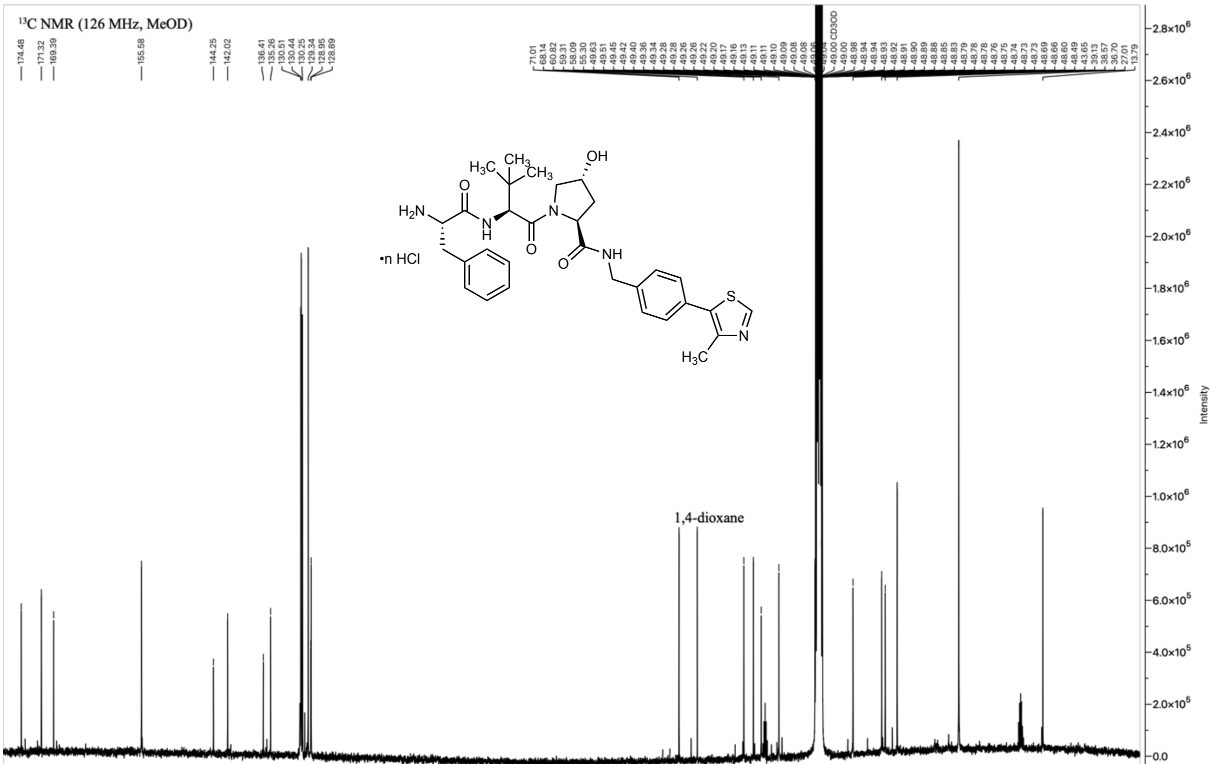

*Figure 52: $^{13}$C NMR spectrum of the Boc-deprotected HCl-salt of tert-butyl ((S)-1-(((S)-1-((2S,4R)-4-hydroxy-2-((4-(4-methylthiazol-5-yl)benzyl)carbamoyl)pyrrolidin-1-yl)-3,3-dimethyl-1-oxobutan-2-yl)amino)-1-oxo-3-phenylpropan-2-yl)carbamate (**18**) in $CD_3OD$.*

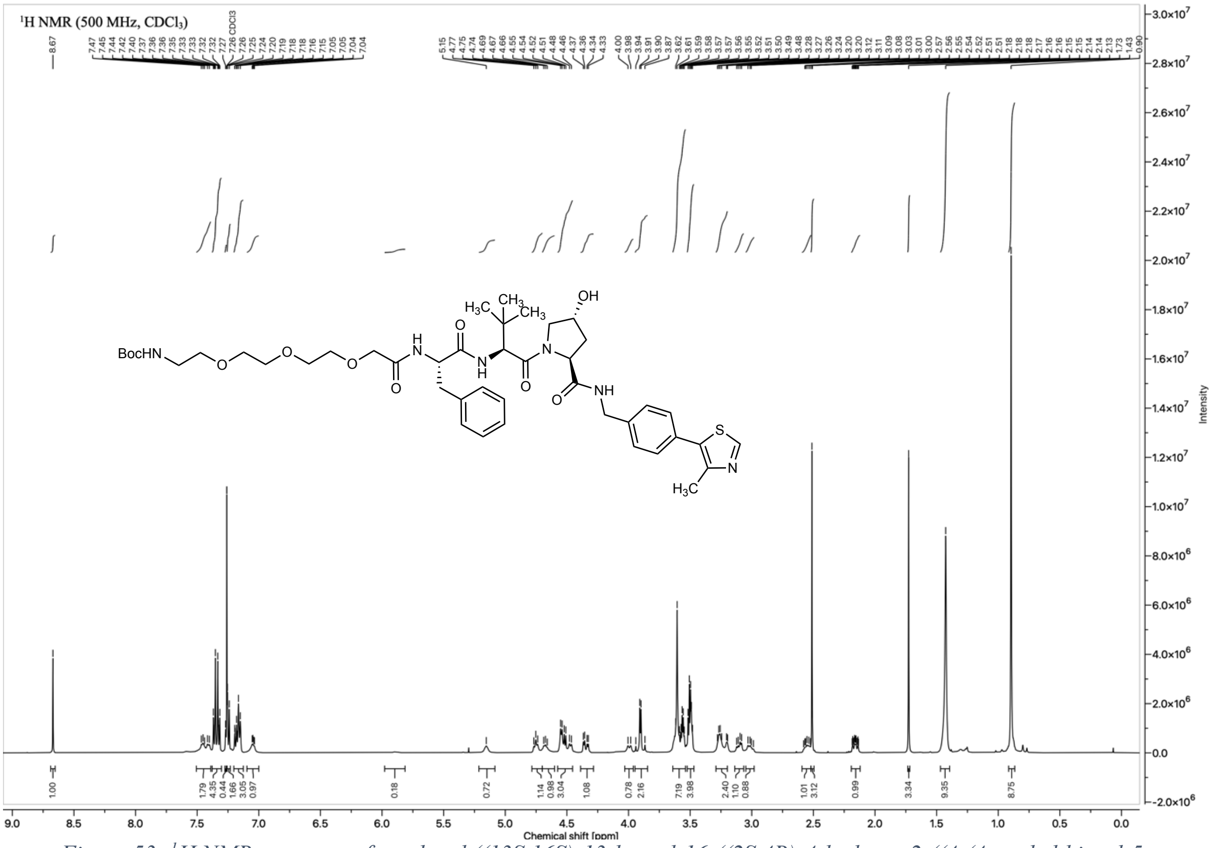

*Figure 53: [1]H NMR spectrum of tert-butyl ((13S,16S)-13-benzyl-16-((2S,4R)-4-hydroxy-2-((4-(4-methylthiazol-5-yl)benzyl)carbamoyl)pyrrolidine-1-carbonyl)-17,17-dimethyl-11,14-dioxo-3,6,9-trioxa-12,15-diazaoctadecyl)carbamate (**19**) in $CDCl_3$*

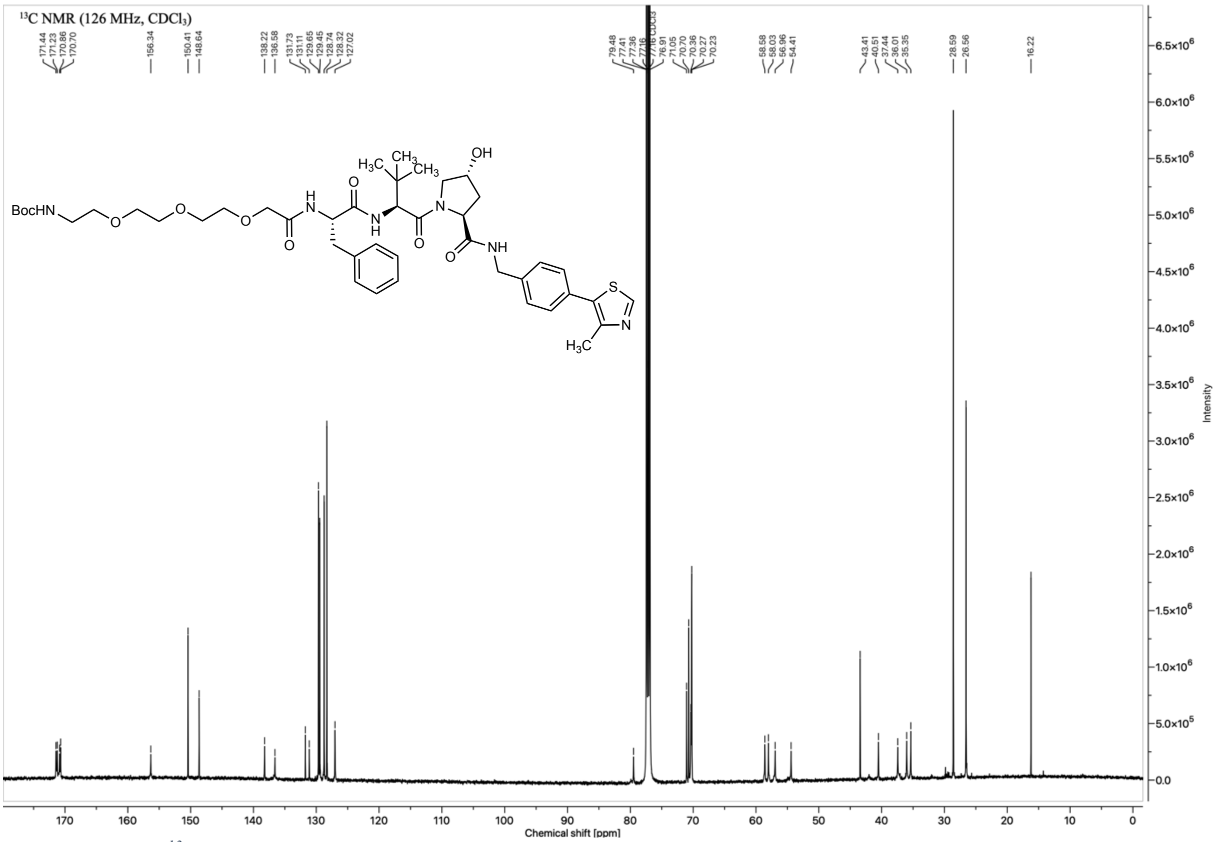

*Figure 54: [13]C NMR spectrum of tert-butyl ((13S,16S)-13-benzyl-16-((2S,4R)-4-hydroxy-2-((4-(4-methylthiazol-5-yl)benzyl)carbamoyl)pyrrolidine-1-carbonyl)-17,17-dimethyl-11,14-dioxo-3,6,9-trioxa-12,15-diazaoctadecyl)carbamate (**19**) in $CDCl_3$*

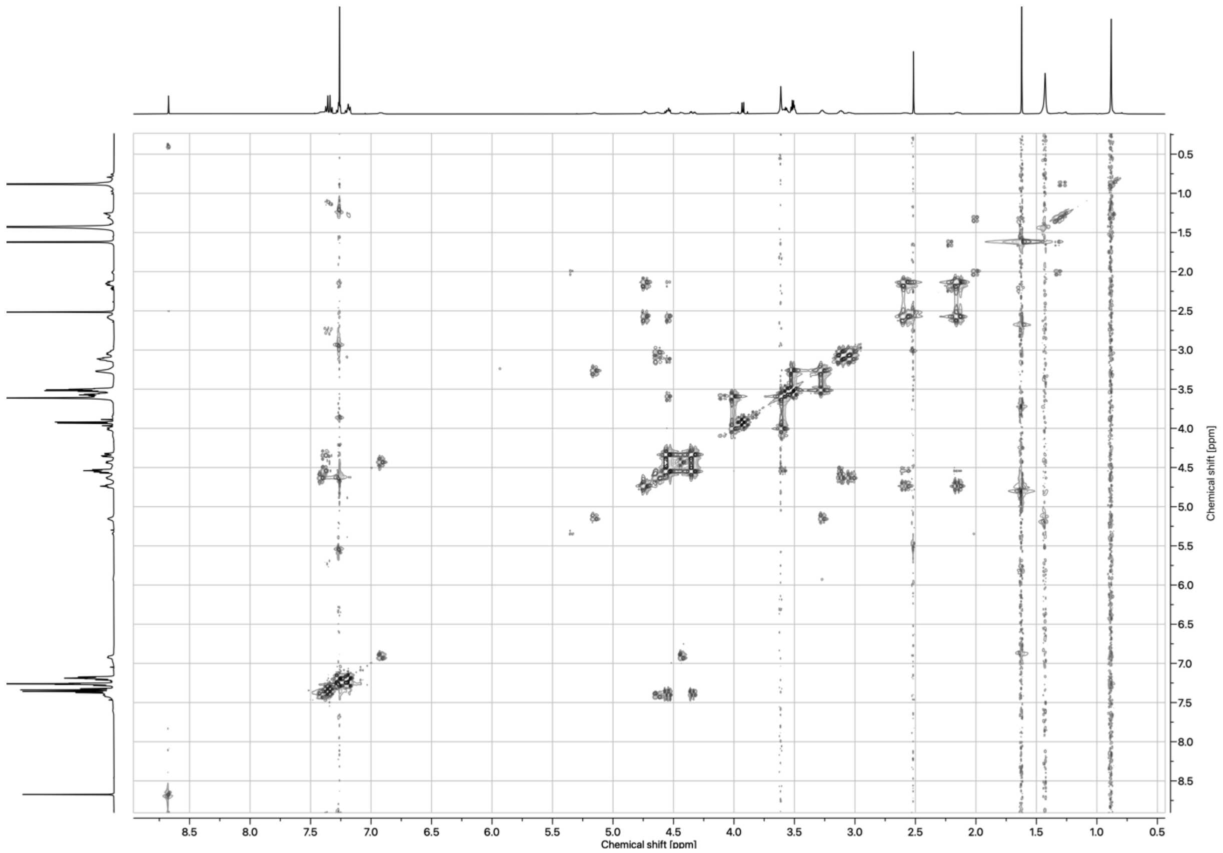

*Figure 55: COSY spectrum of tert-butyl ((13S,16S)-13-benzyl-16-((2S,4R)-4-hydroxy-2-((4-(4-methylthiazol-5-yl)benzyl)carbamoyl)pyrrolidine-1-carbonyl)-17,17-dimethyl-11,14-dioxo-3,6,9-trioxa-12,15-diazaoctadecyl)carbamate (**19**) in $CDCl_3$*

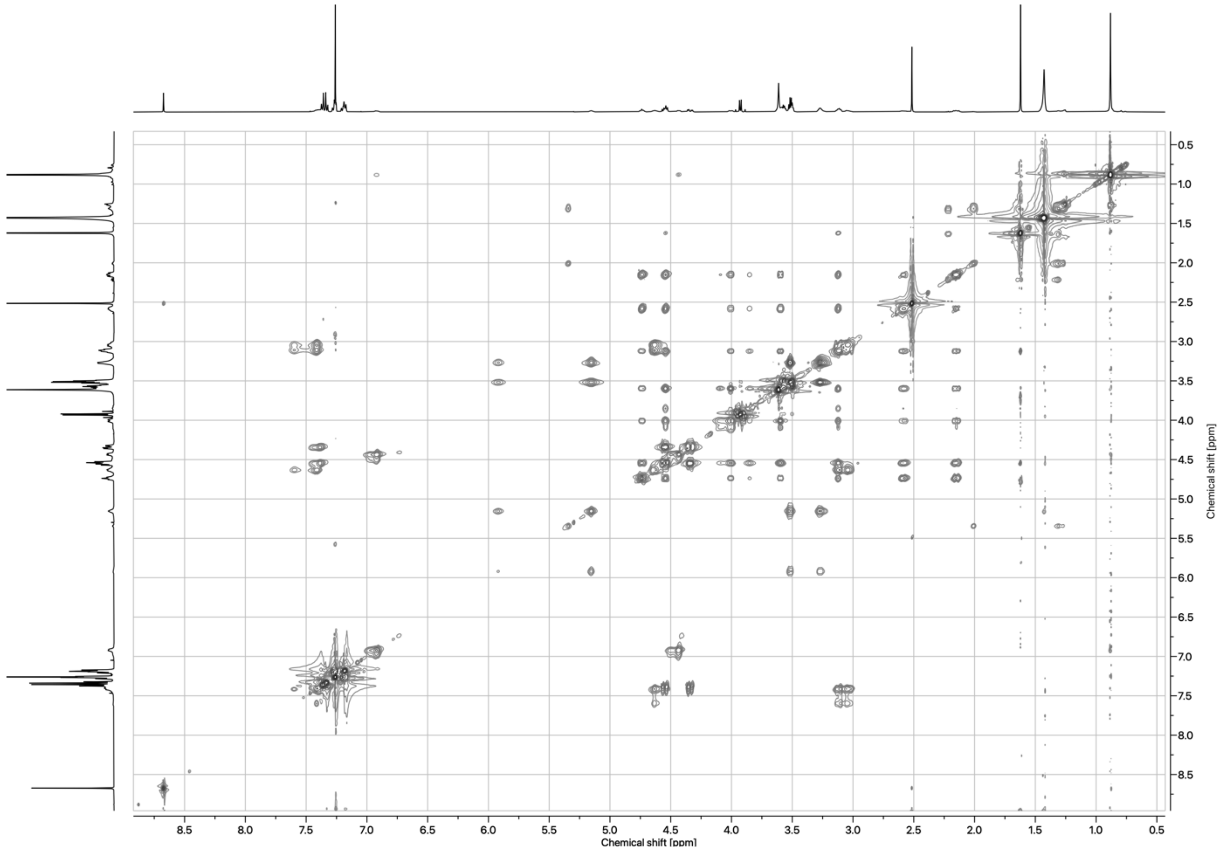

*Figure 56: TOCSY spectrum of tert-butyl ((13S,16S)-13-benzyl-16-((2S,4R)-4-hydroxy-2-((4-(4-methylthiazol-5-yl)benzyl)carbamoyl)pyrrolidine-1-carbonyl)-17,17-dimethyl-11,14-dioxo-3,6,9-trioxa-12,15-diazaoctadecyl)carbamate (**19**) in $CDCl_3$*

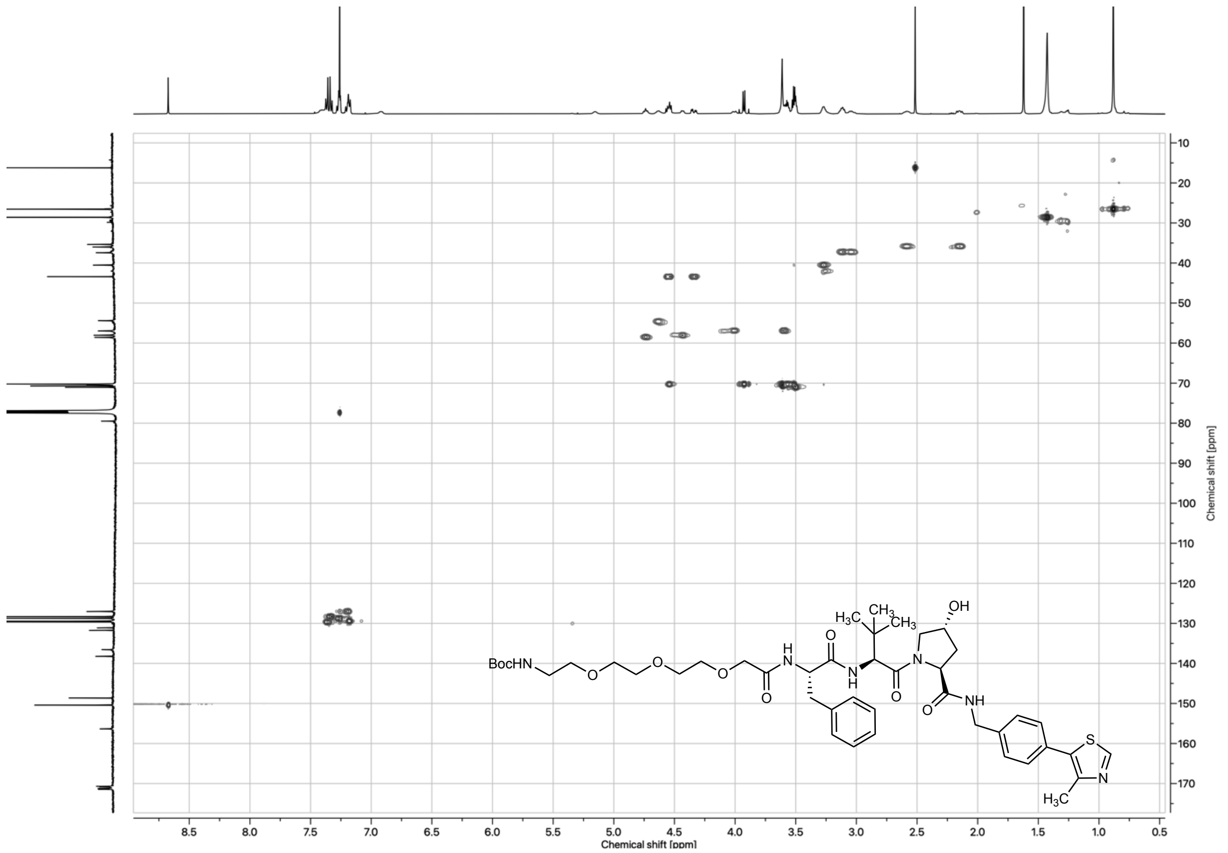

*Figure 57: HSQC spectrum of tert-butyl ((13S,16S)-13-benzyl-16-((2S,4R)-4-hydroxy-2-((4-(4-methylthiazol-5-yl)benzyl)carbamoyl)pyrrolidine-1-carbonyl)-17,17-dimethyl-11,14-dioxo-3,6,9-trioxa-12,15-diazaoctadecyl)carbamate (**19**) in $CDCl_3$*

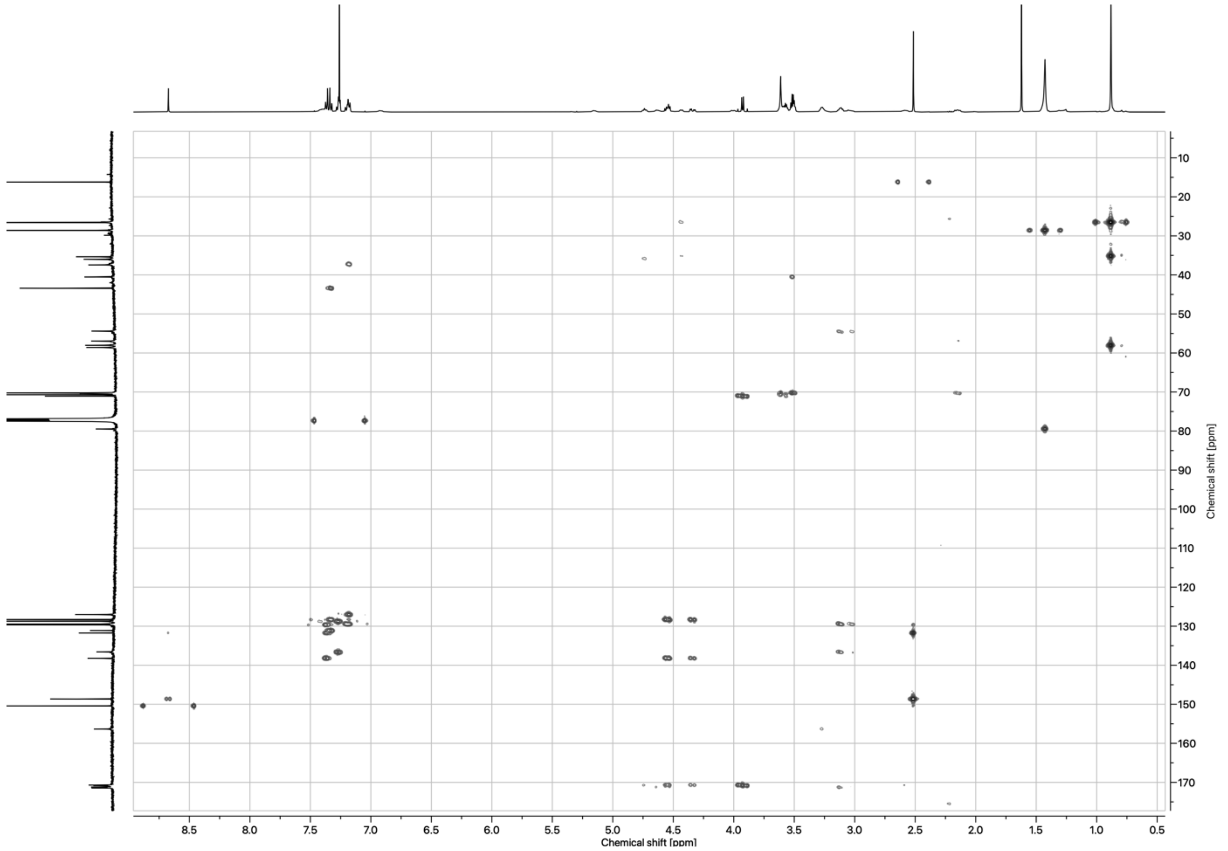

*Figure 58: HMBC spectrum of tert-butyl ((13S,16S)-13-benzyl-16-((2S,4R)-4-hydroxy-2-((4-(4-methylthiazol-5-yl)benzyl)carbamoyl)pyrrolidine-1-carbonyl)-17,17-dimethyl-11,14-dioxo-3,6,9-trioxa-12,15-diazaoctadecyl)carbamate (**19**) in $CDCl_3$*

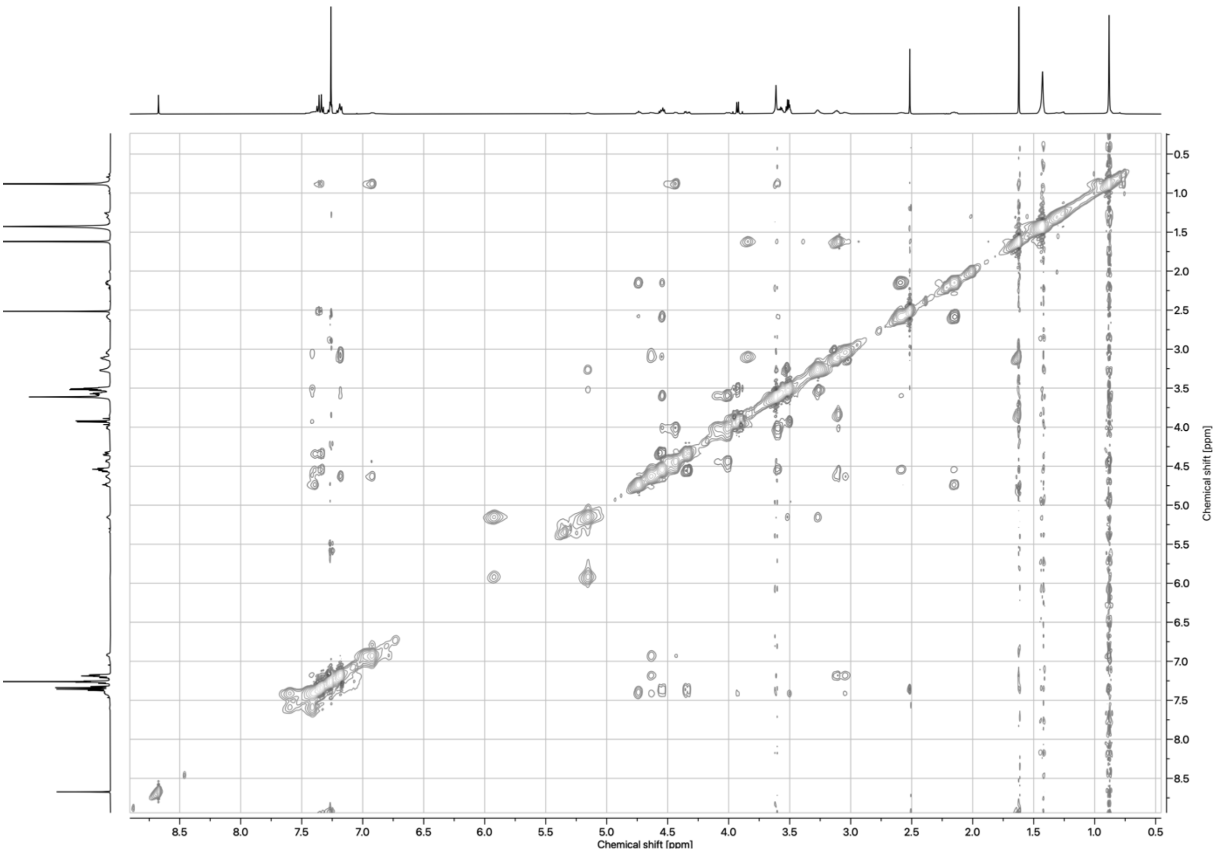

*Figure 59: NOESY spectrum of tert-butyl ((13S,16S)-13-benzyl-16-((2S,4R)-4-hydroxy-2-((4-(4-methylthiazol-5-yl)benzyl)carbamoyl)pyrrolidine-1-carbonyl)-17,17-dimethyl-11,14-dioxo-3,6,9-trioxa-12,15-diazaoctadecyl)carbamate (**19**) in $CDCl_3$*

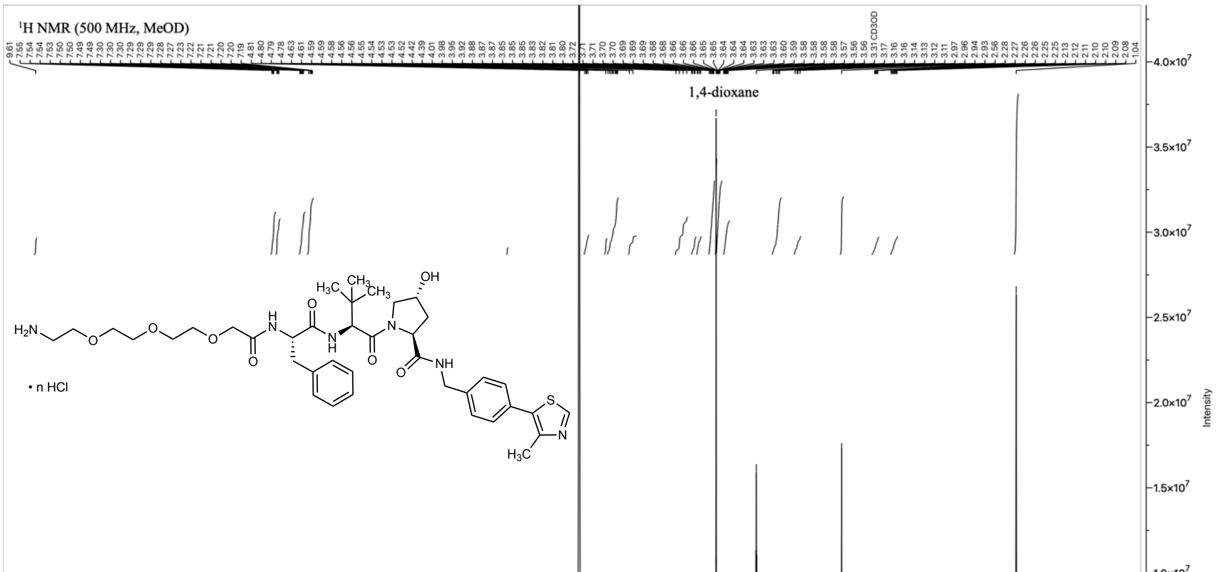

*Figure 60: ¹H NMR spectrum of Boc-deprotected HCl-salt of tert-butyl ((13S,16S)-13-benzyl-16-((2S,4R)-4-hydroxy-2-((4-(4-methylthiazol-5-yl)benzyl)carbamoyl)pyrrolidine-1-carbonyl)-17,17-dimethyl-11,14-dioxo-3,6,9-trioxa-12,15-diazaoctadecyl)carbamate (**20**) in $CD_3OD$*

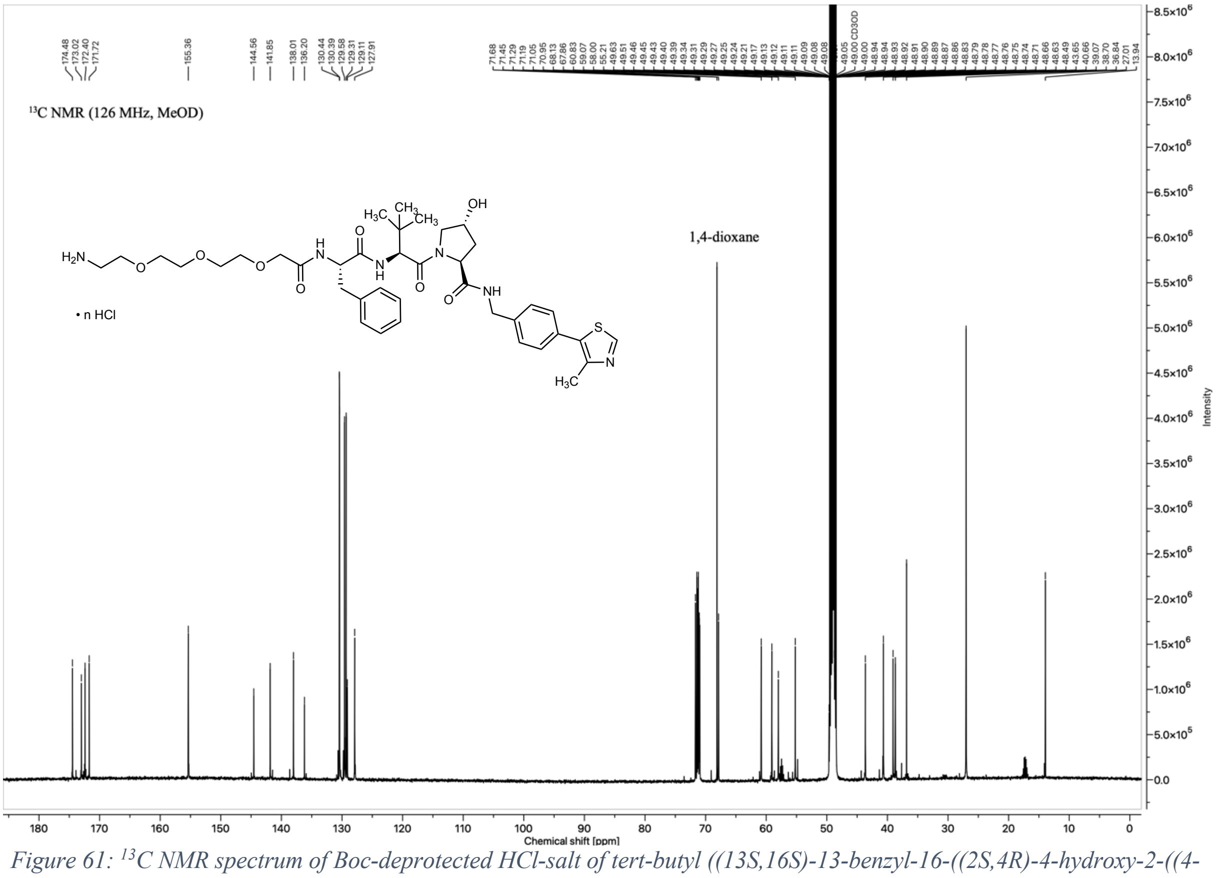

*Figure 61: $^{13}$C NMR spectrum of Boc-deprotected HCl-salt of tert-butyl ((13S,16S)-13-benzyl-16-((2S,4R)-4-hydroxy-2-((4-(4-methylthiazol-5-yl)benzyl)carbamoyl)pyrrolidine-1-carbonyl)-17,17-dimethyl-11,14-dioxo-3,6,9-trioxa-12,15-diazaoctadecyl)carbamate (**20**) in $CD_3OD$*

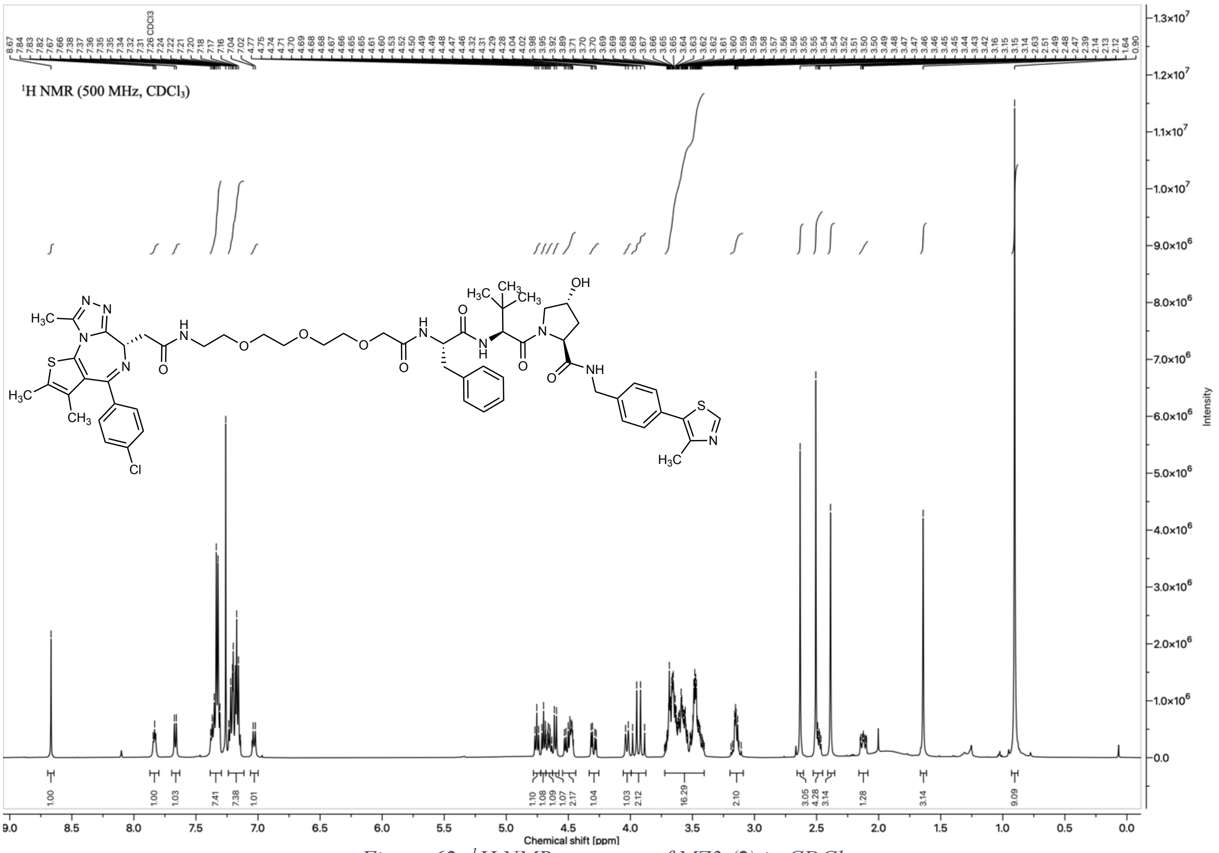

*Figure 62: $^{1}$H NMR spectrum of MZ3 (**2**) in $CDCl_3$*

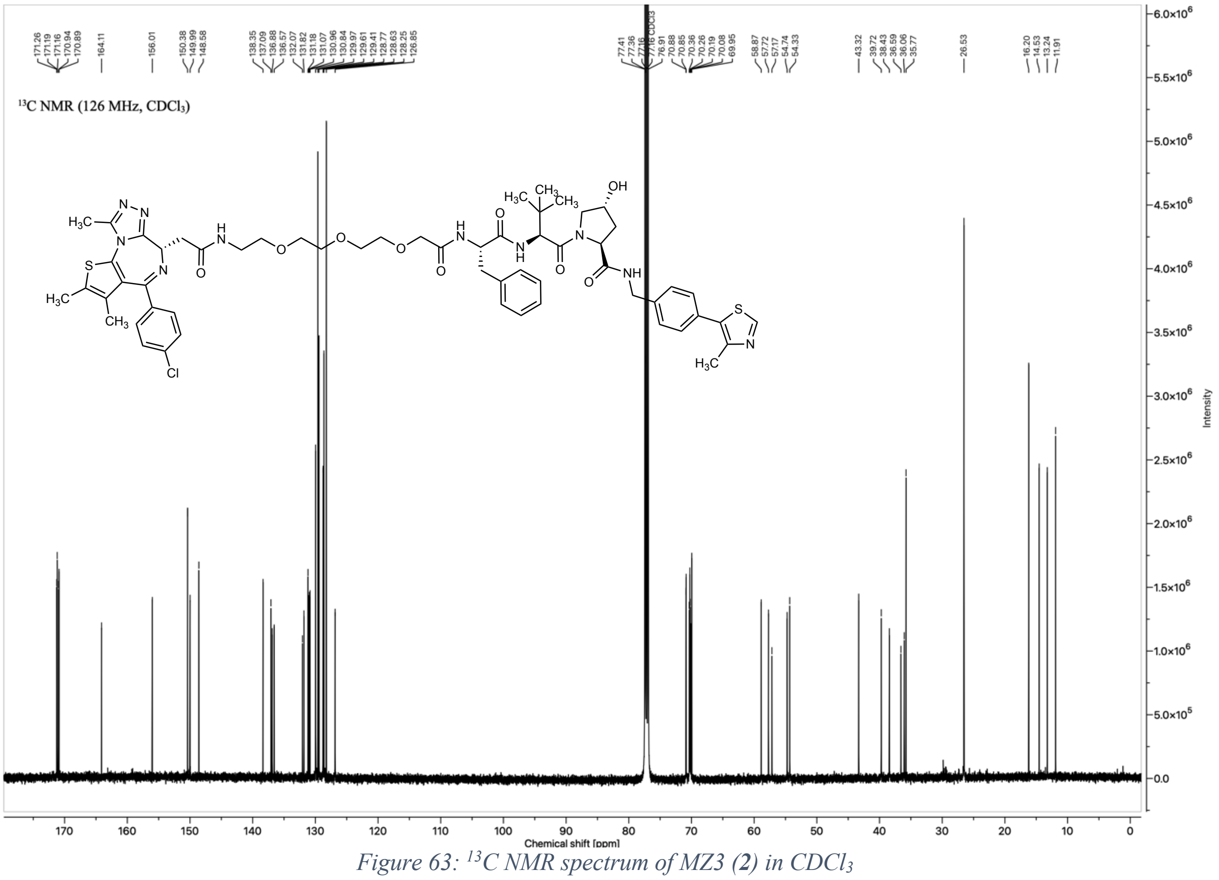

*Figure 63: $^{13}C$ NMR spectrum of MZ3 (**2**) in $CDCl_3$*

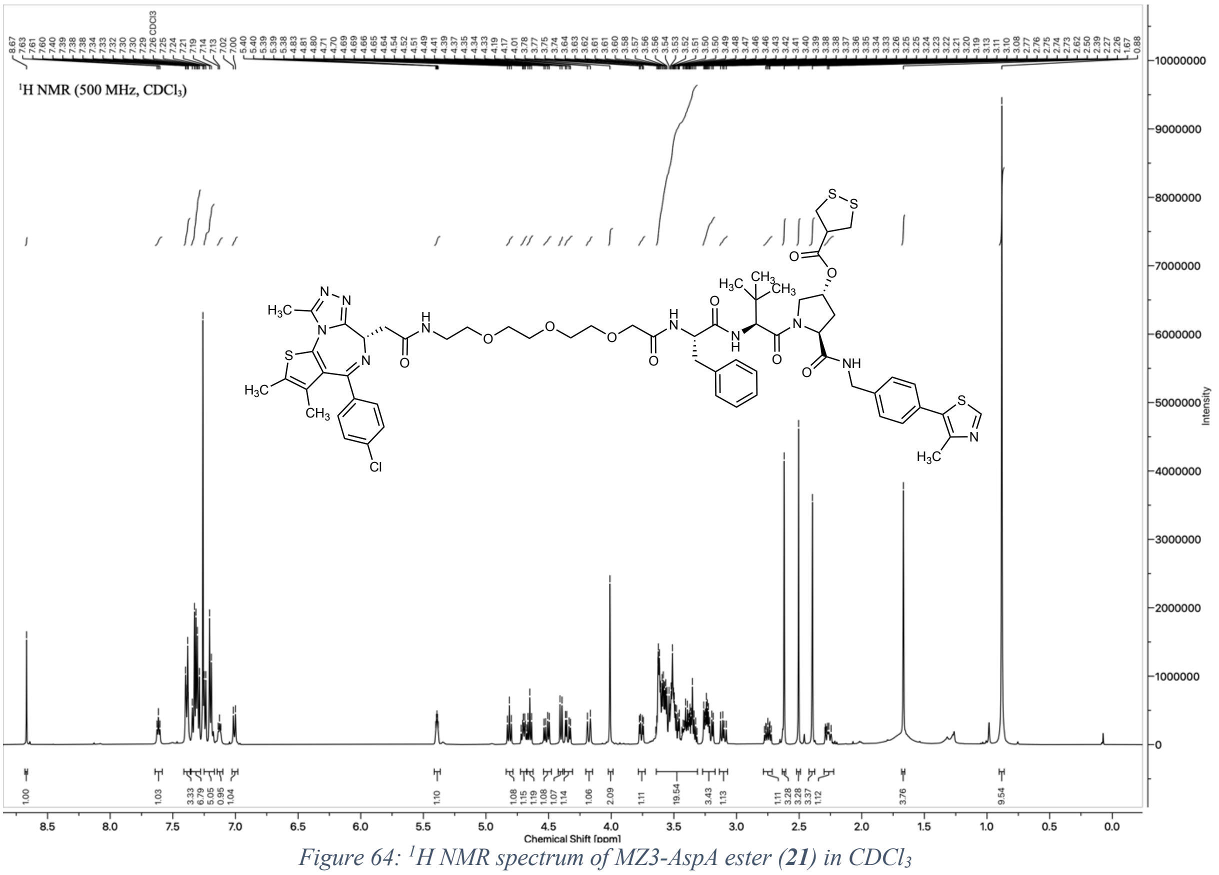

*Figure 64: $^{1}H$ NMR spectrum of MZ3-AspA ester (**21**) in $CDCl_3$*

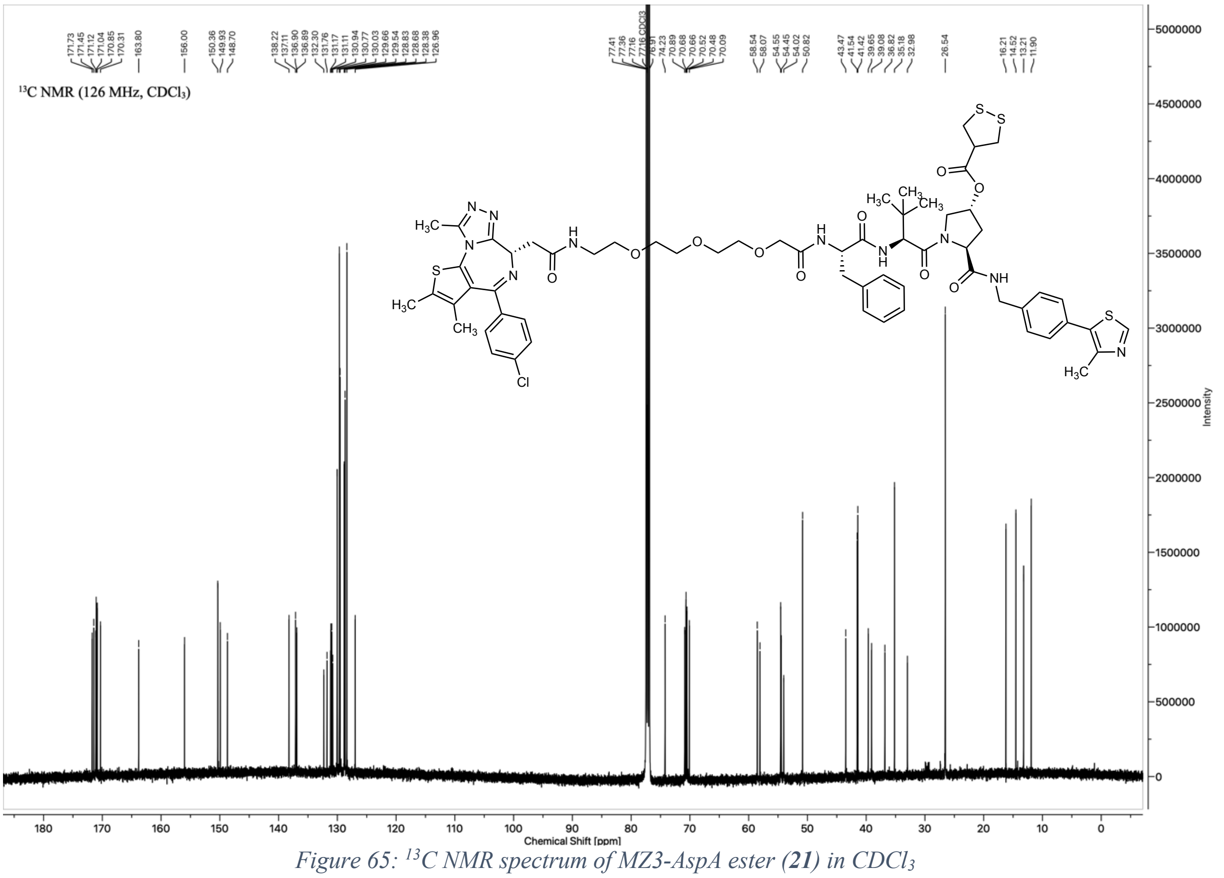

*Figure 65: $^{13}$C NMR spectrum of MZ3-AspA ester (**21**) in $CDCl_3$*

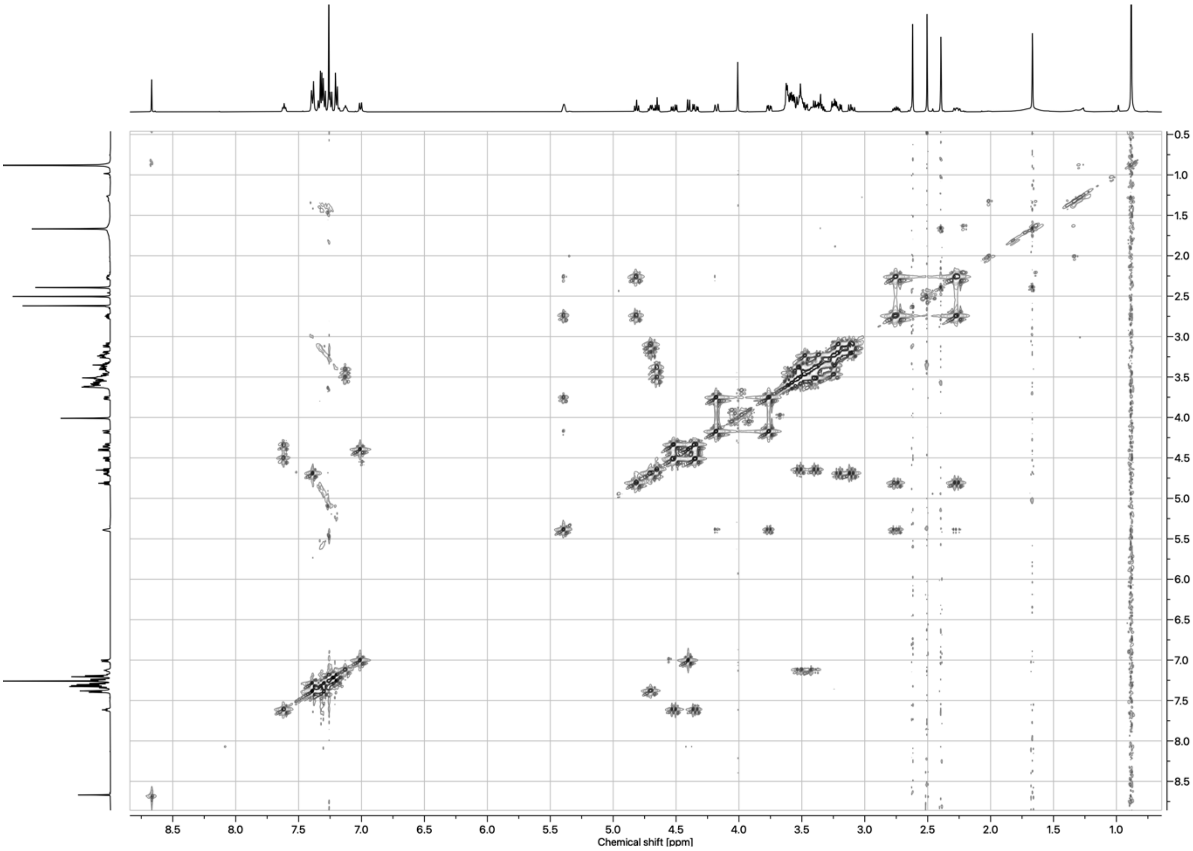

*Figure 66: COSY spectrum of MZ3-AspA ester (**21**) in $CDCl_3$*

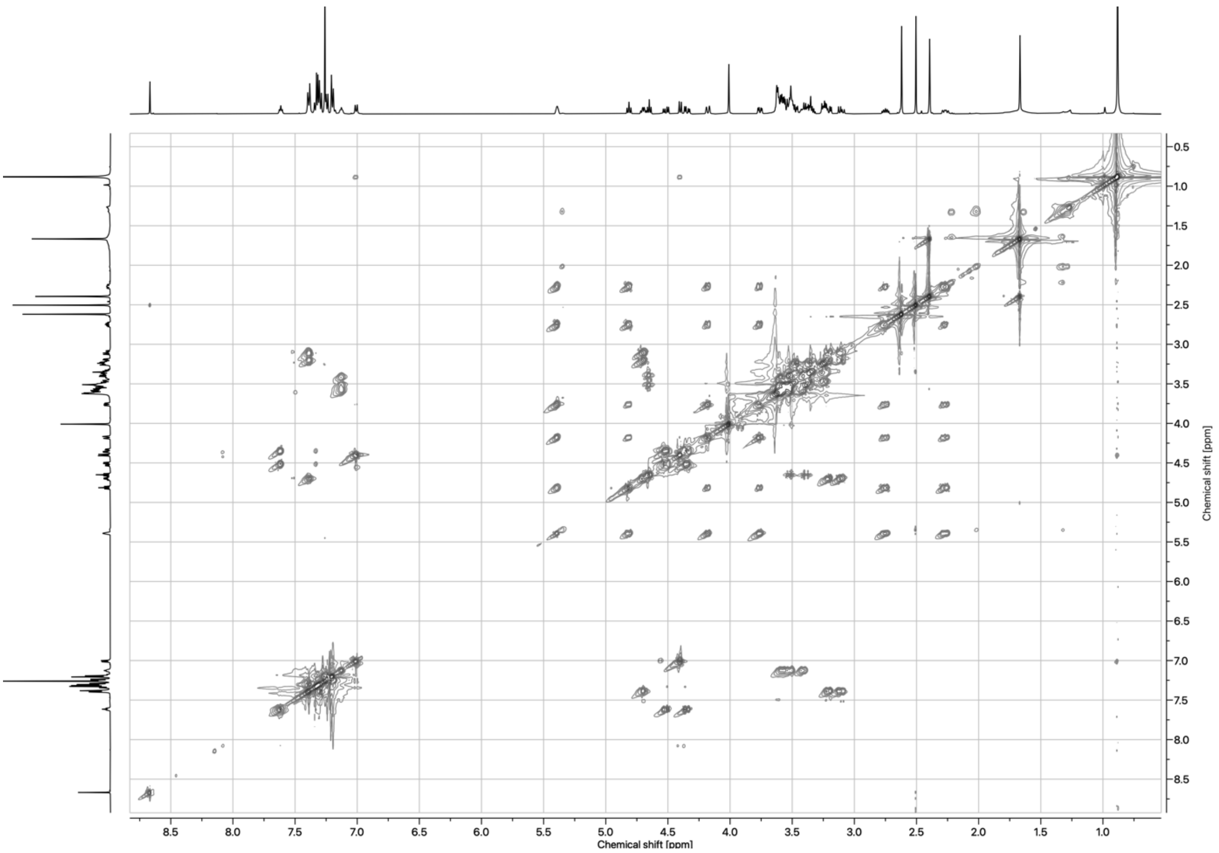

*Figure 67: TOCSY spectrum of MZ3-AspA ester (**21**) in $CDCl_3$*

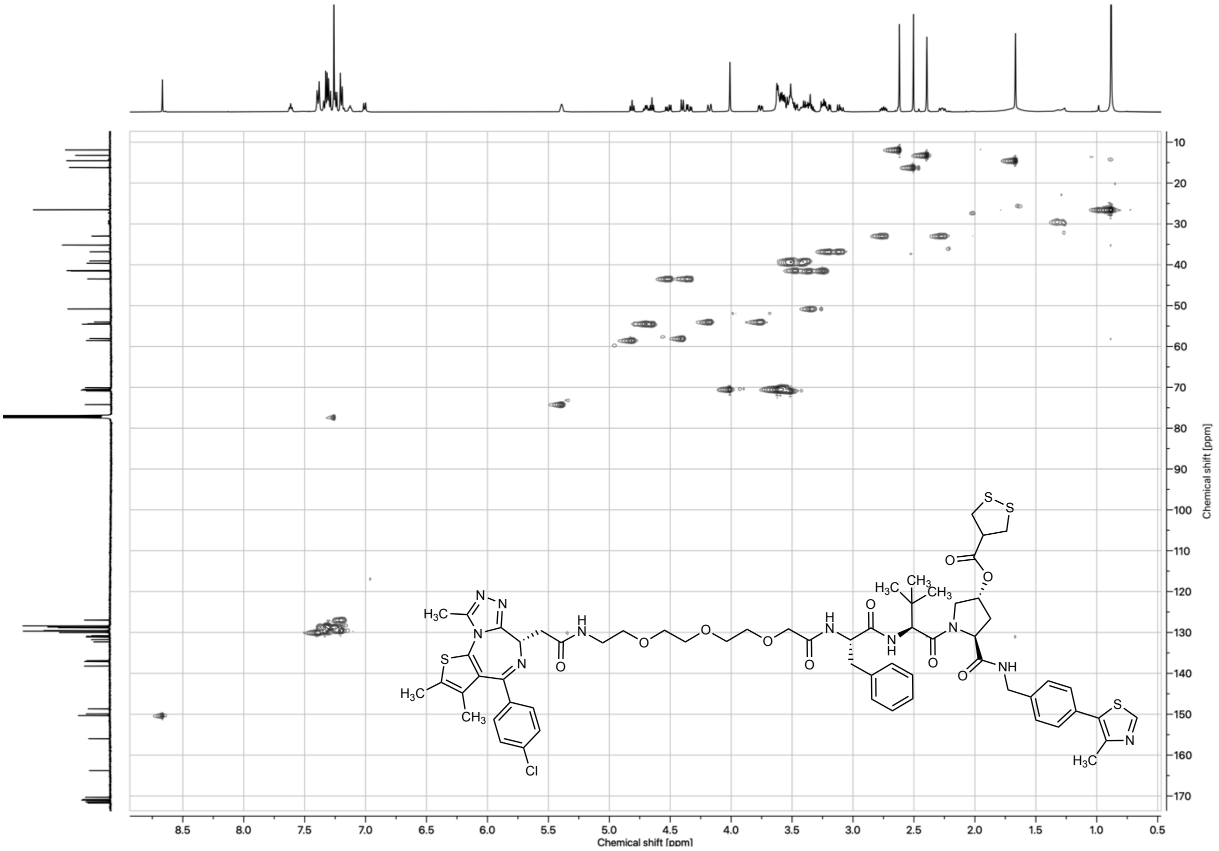

*Figure 68: HSQC spectrum of MZ3-AspA ester (**21**) in $CDCl_3$*

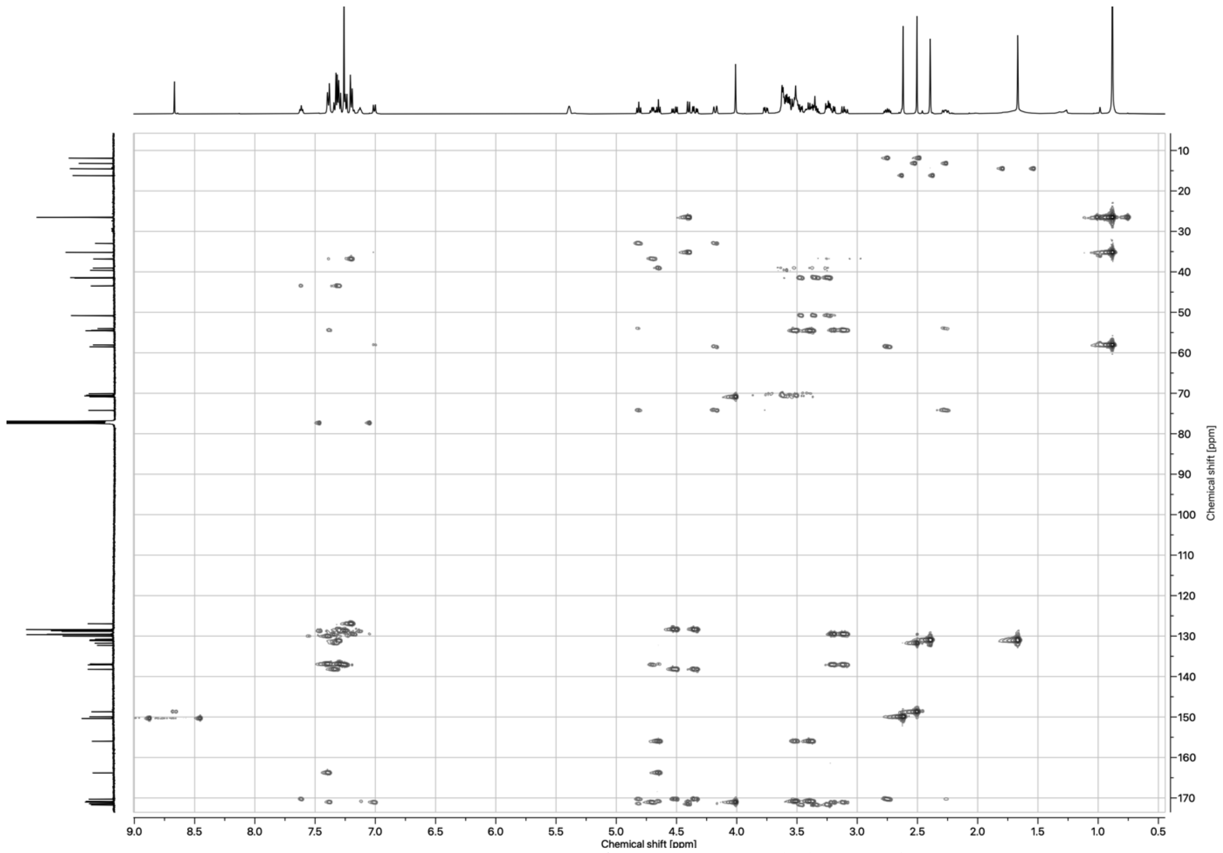

*Figure 69: HMBC spectrum of MZ3-AspA ester (**21**) in $CDCl_3$*

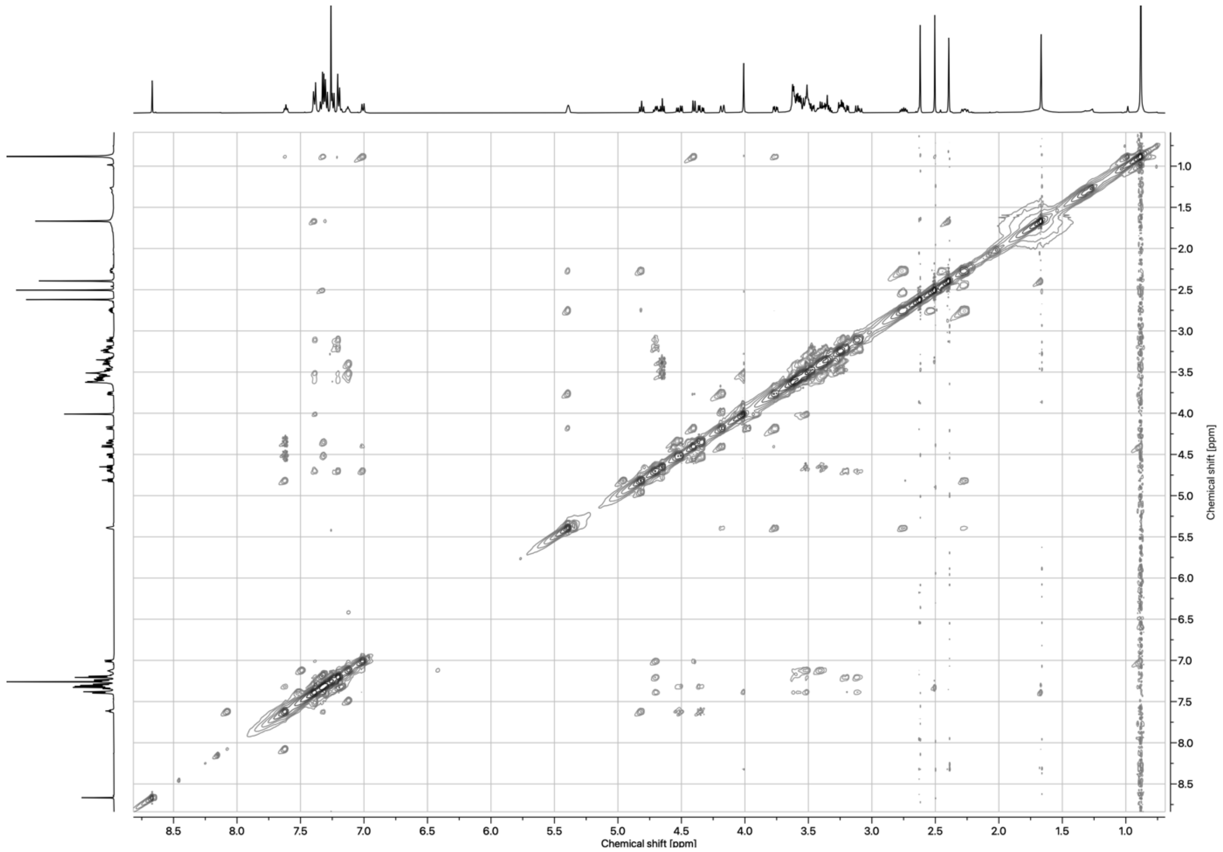

*Figure 70: NOESY spectrum of MZ3-AspA ester (**21**) in $CDCl_3$*

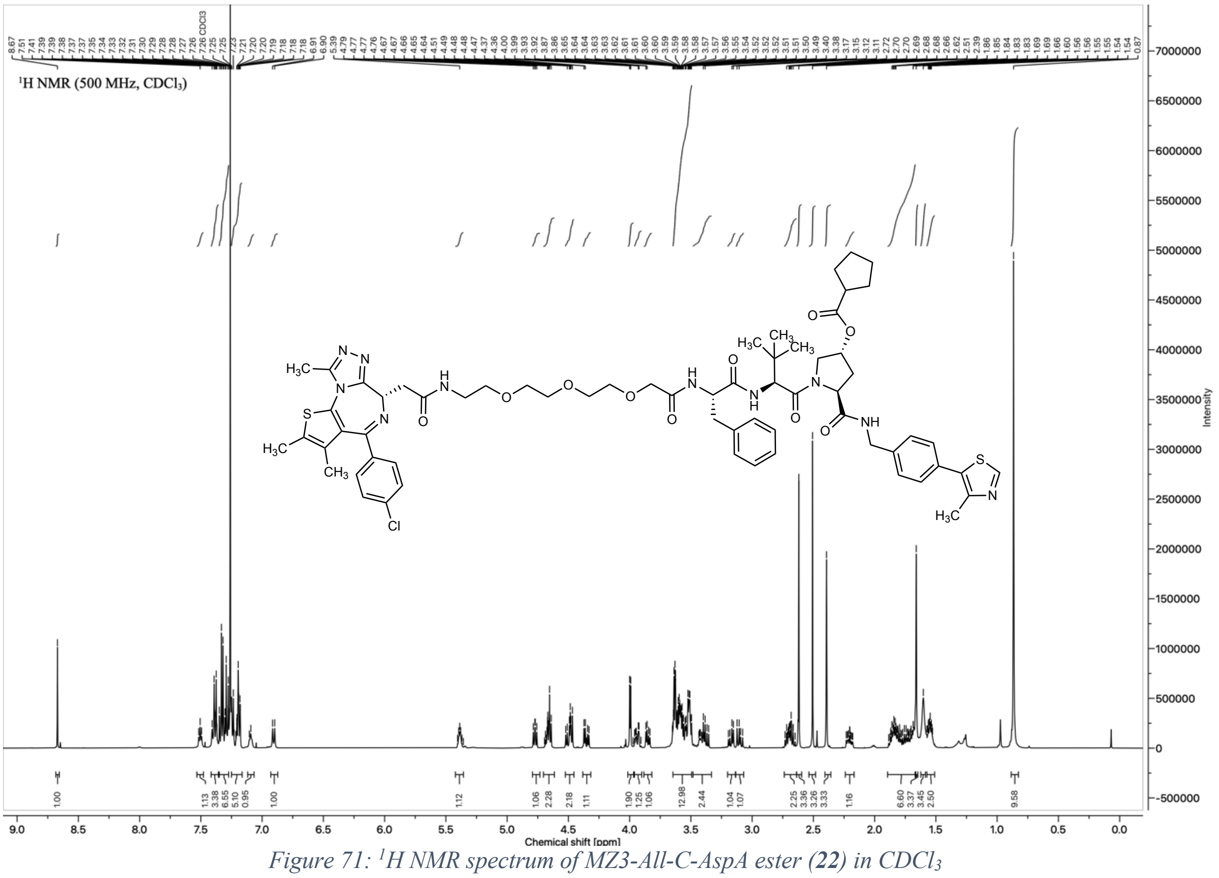

*Figure 71: $^{1}$H NMR spectrum of MZ3-All-C-AspA ester (**22**) in CDCl$_3$*

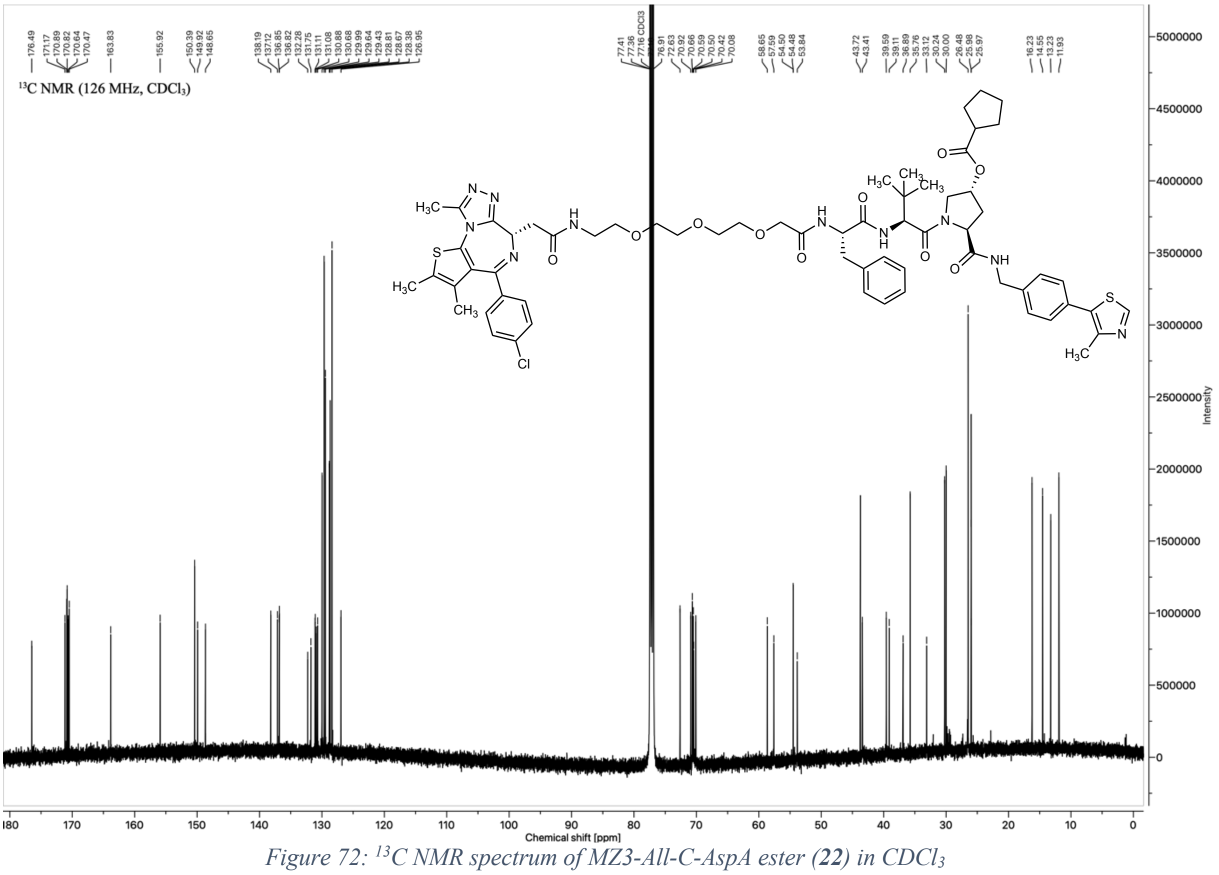

*Figure 72: $^{13}$C NMR spectrum of MZ3-All-C-AspA ester (**22**) in CDCl$_3$*

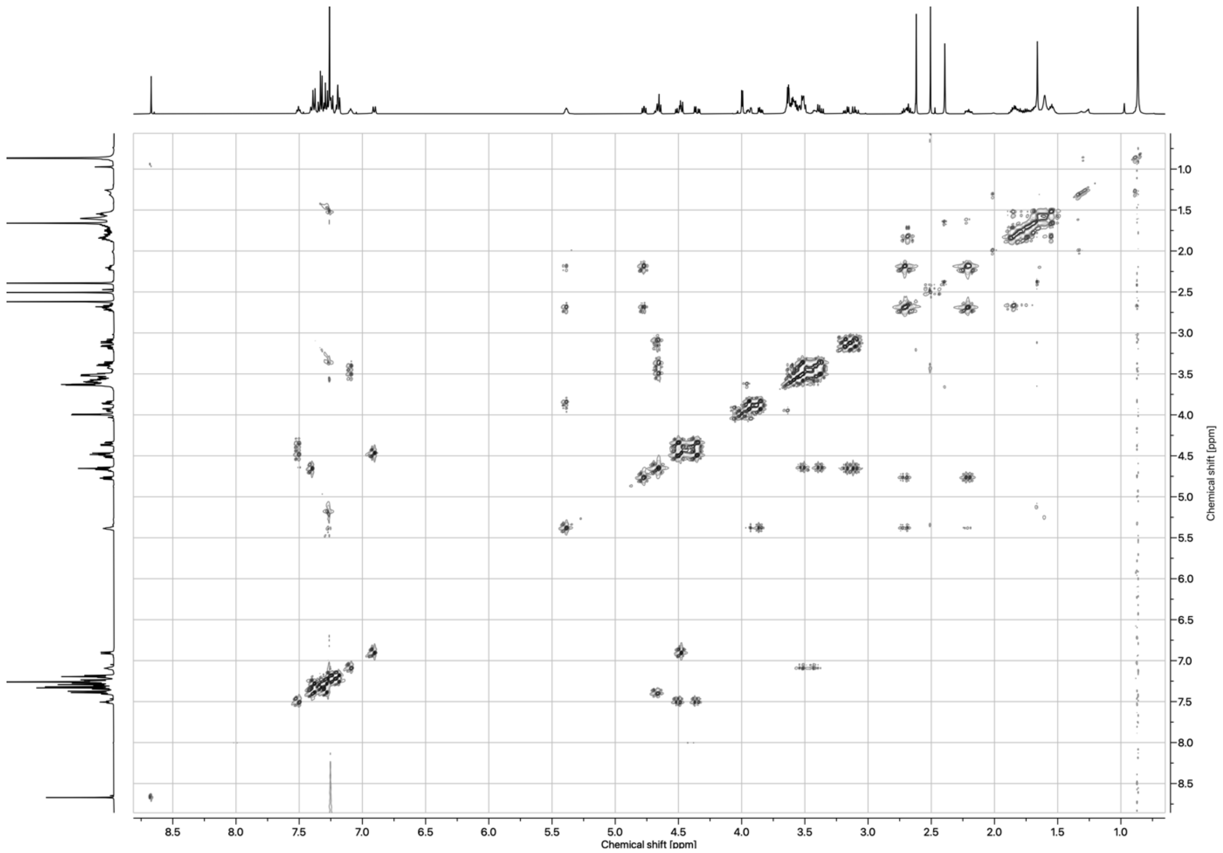

*Figure 73: COSY spectrum of MZ3-All-C-AspA ester (**22**) in $CDCl_3$*

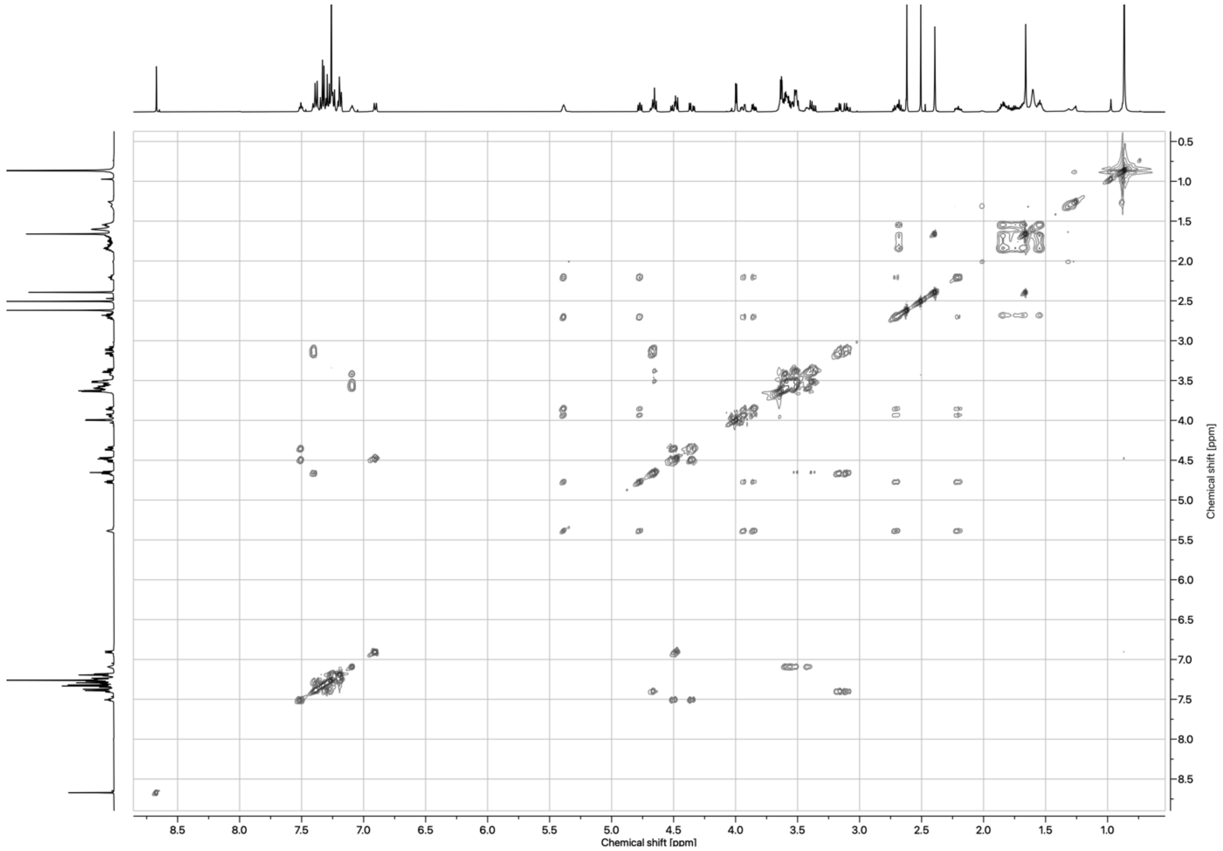

*Figure 74: TOCSY spectrum of MZ3-All-C-AspA ester (**22**) in $CDCl_3$*

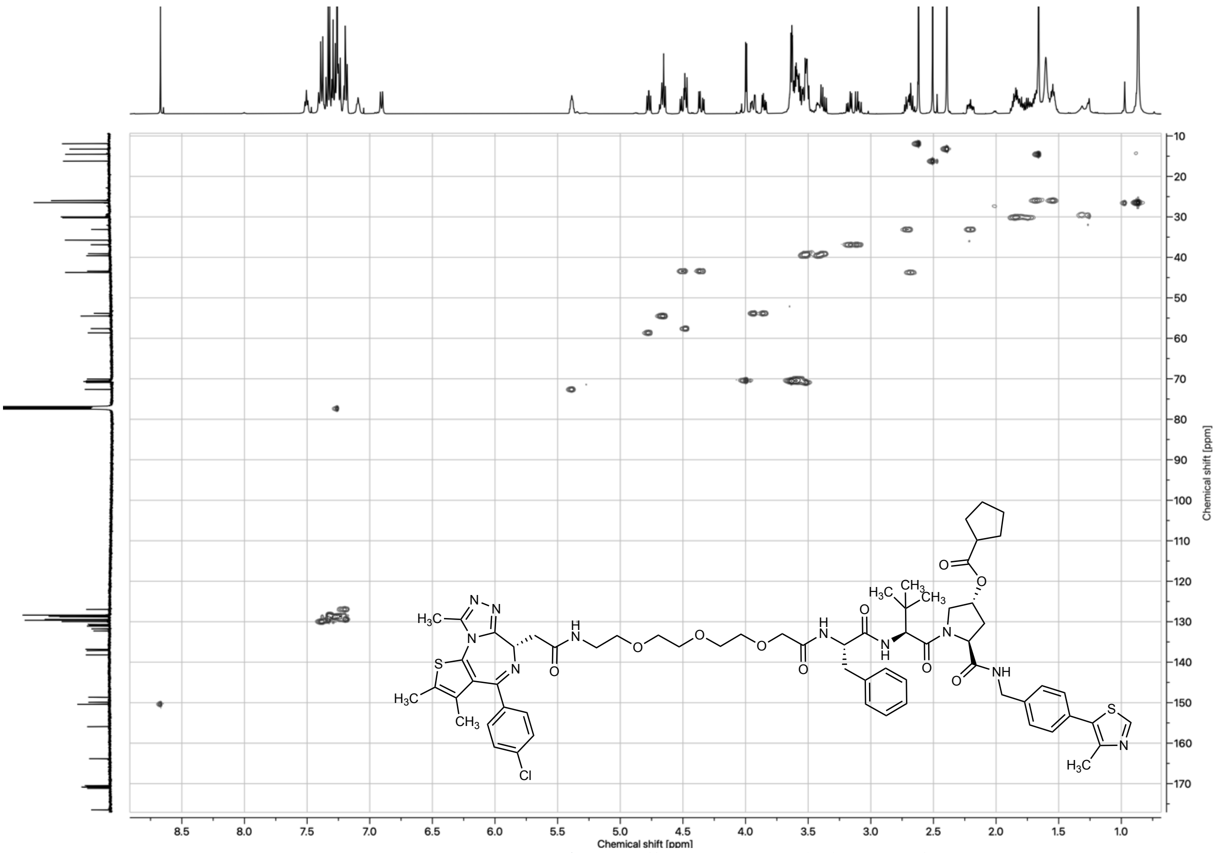

*Figure 75: HSQC spectrum of MZ3-All-C-AspA ester (**22**) in $CDCl_3$*

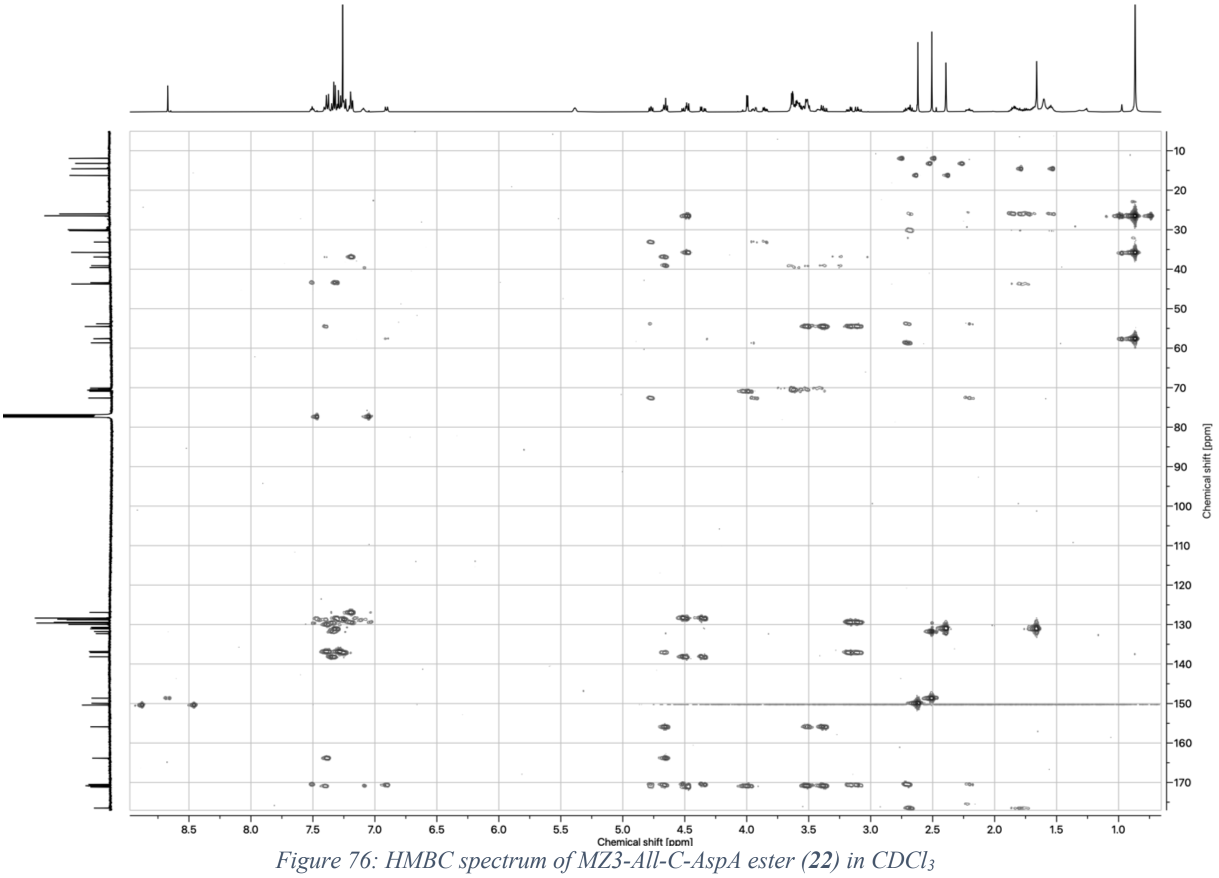

*Figure 76: HMBC spectrum of MZ3-All-C-AspA ester (**22**) in $CDCl_3$*

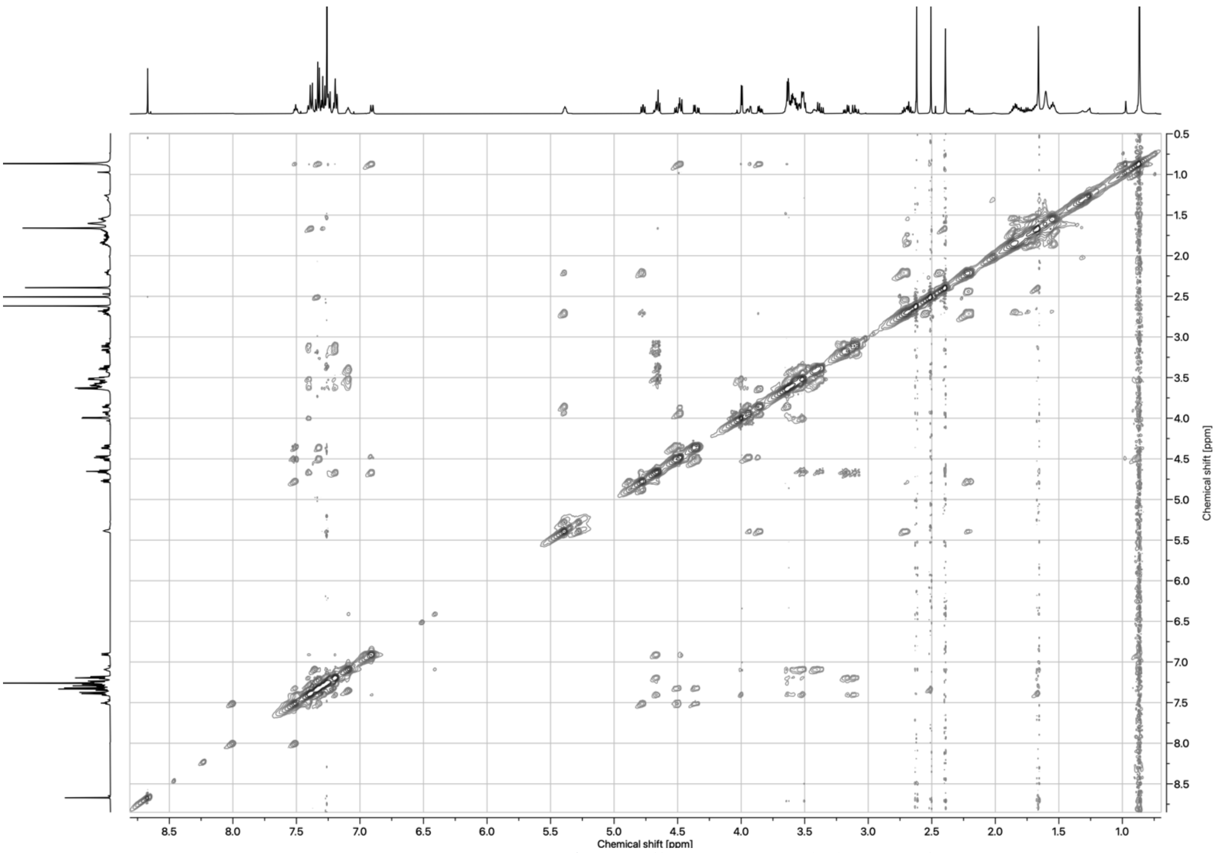

*Figure 77: NOESY spectrum of MZ3-All-C-AspA ester (**22**) in $CDCl_3$*

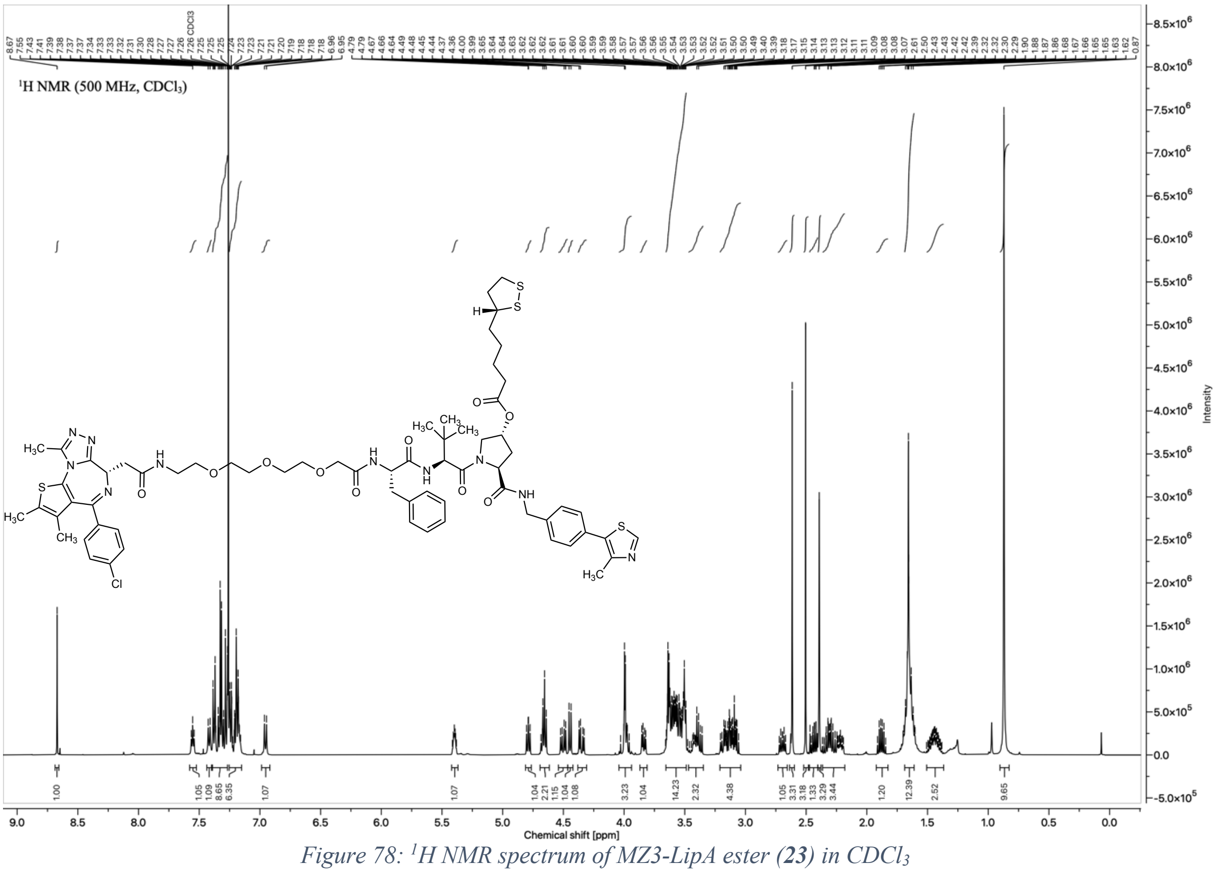

*Figure 78: $^1$H NMR spectrum of MZ3-LipA ester (**23**) in $CDCl_3$*

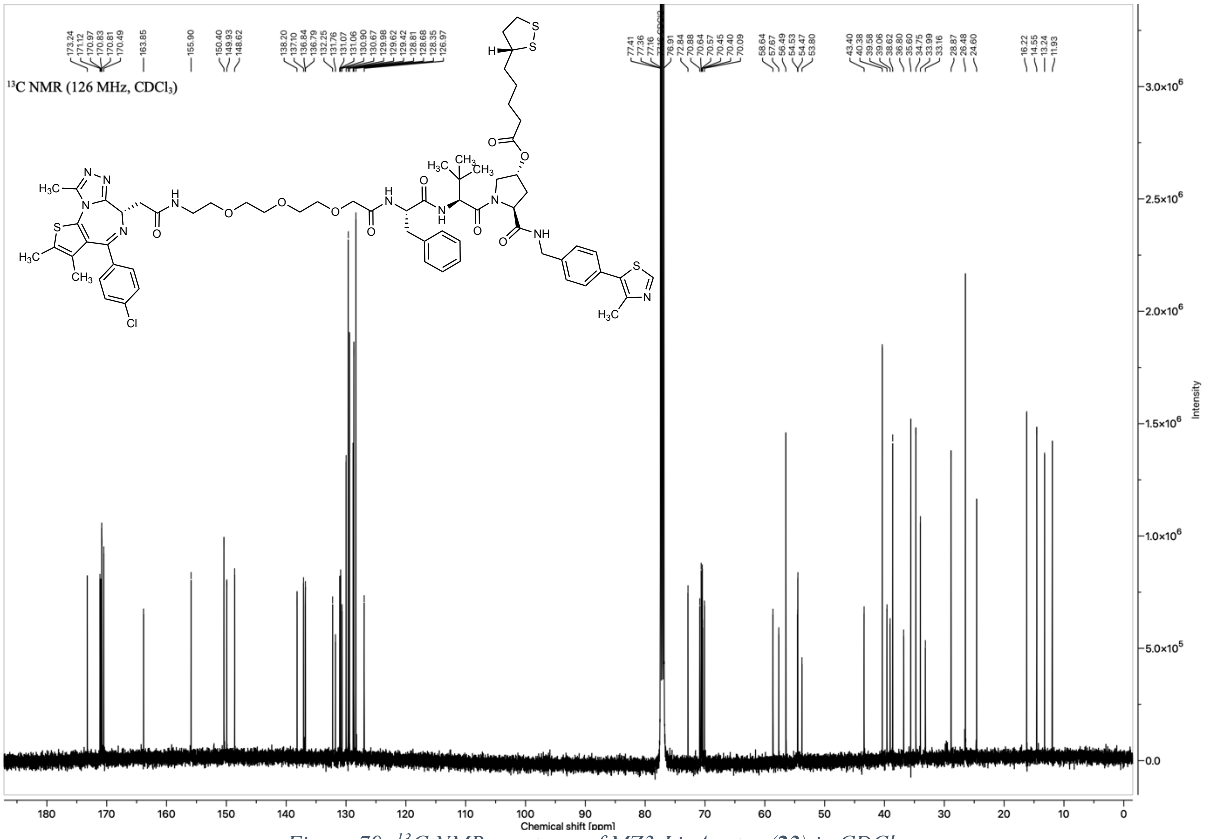

*Figure 79: $^{13}$C NMR spectrum of MZ3-LipA ester (**23**) in $CDCl_3$*

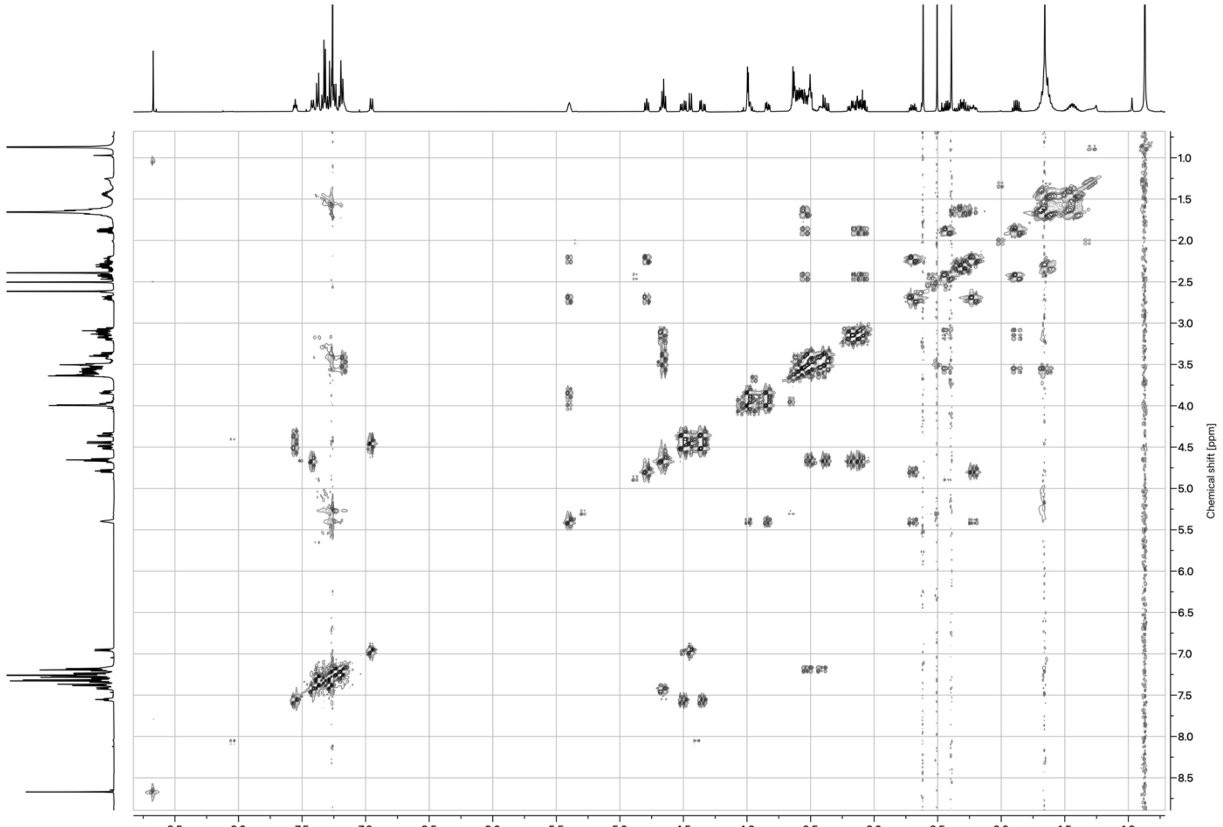

*Figure 80: COSY spectrum of MZ3-LipA ester (**23**) in $CDCl_3$*

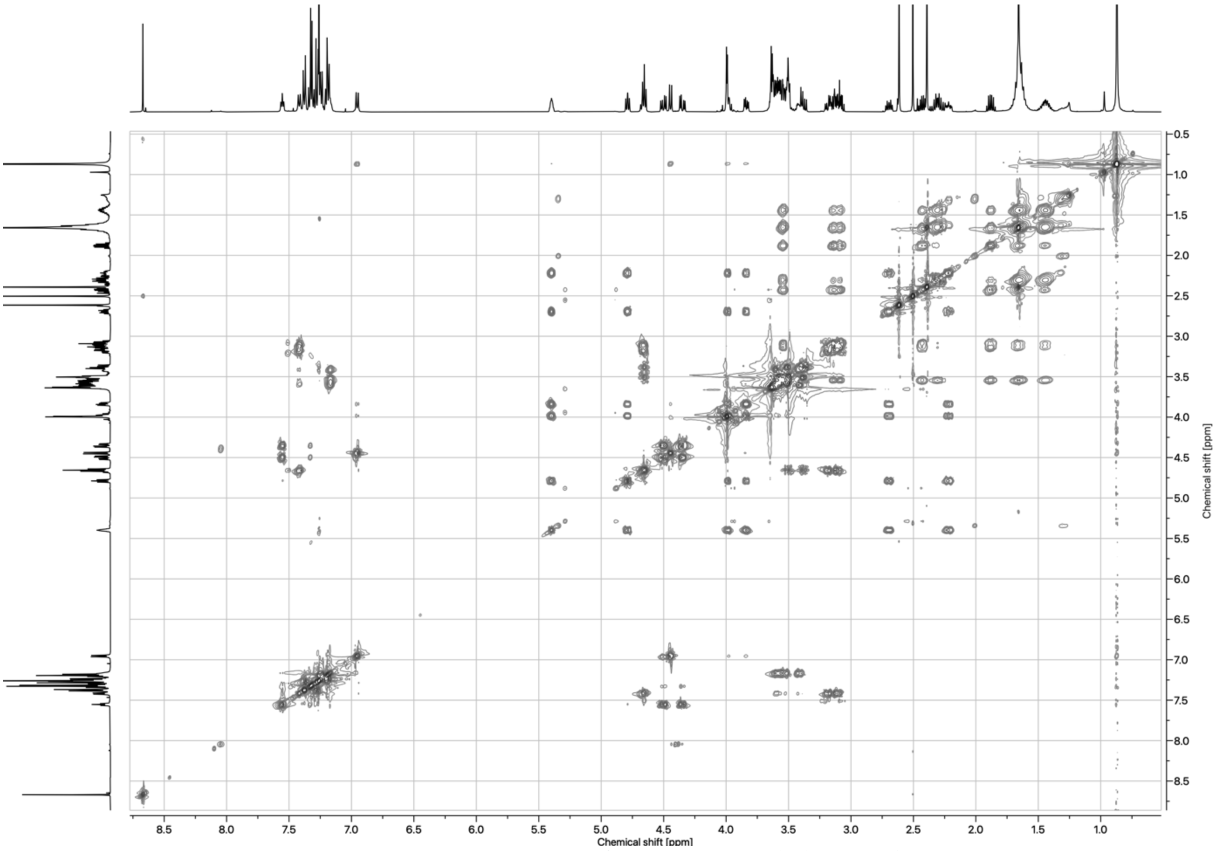

*Figure 81: TOCSY spectrum of MZ3-LipA ester (**23**) in $CDCl_3$*

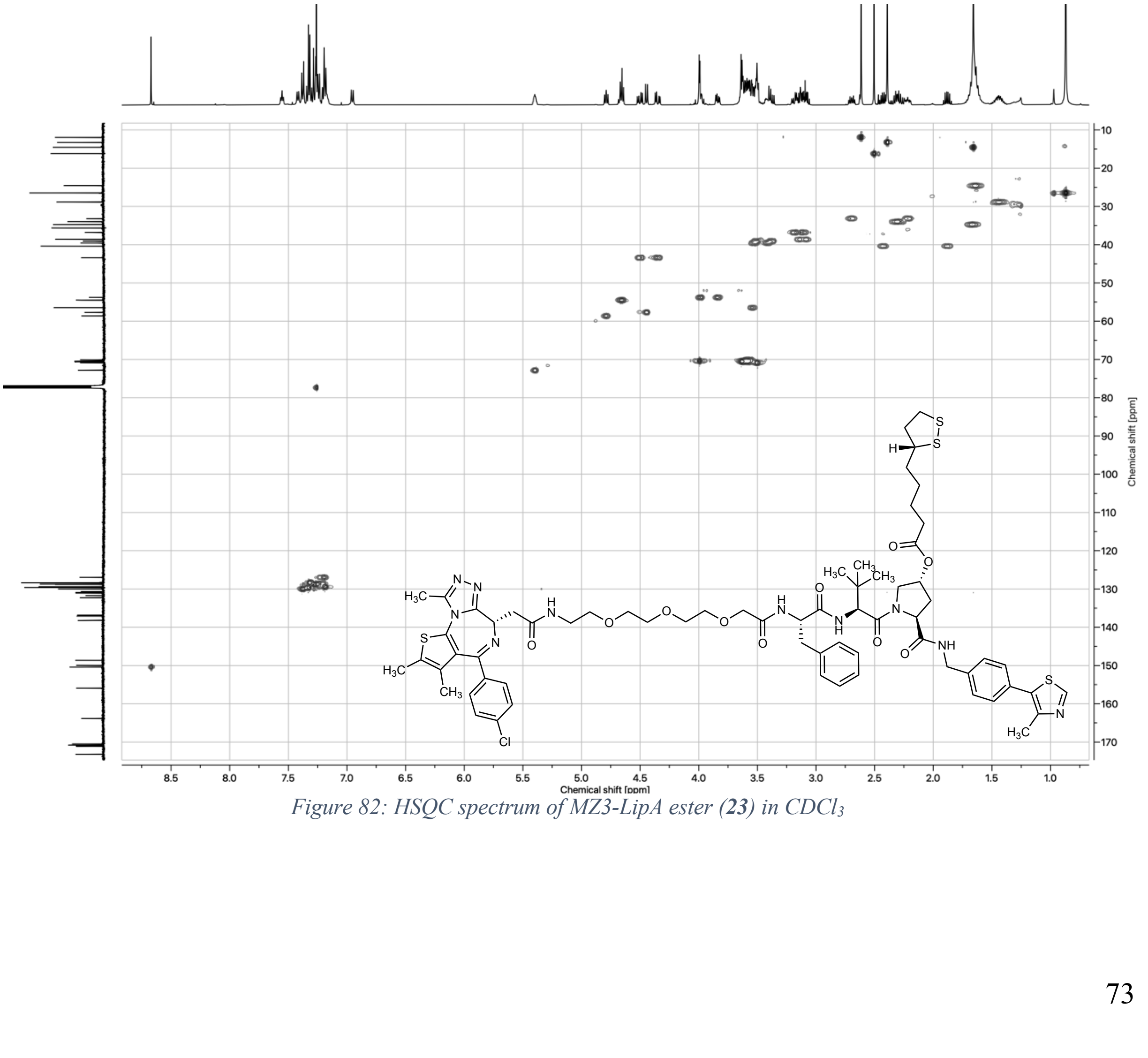

*Figure 82: HSQC spectrum of MZ3-LipA ester (**23**) in $CDCl_3$*

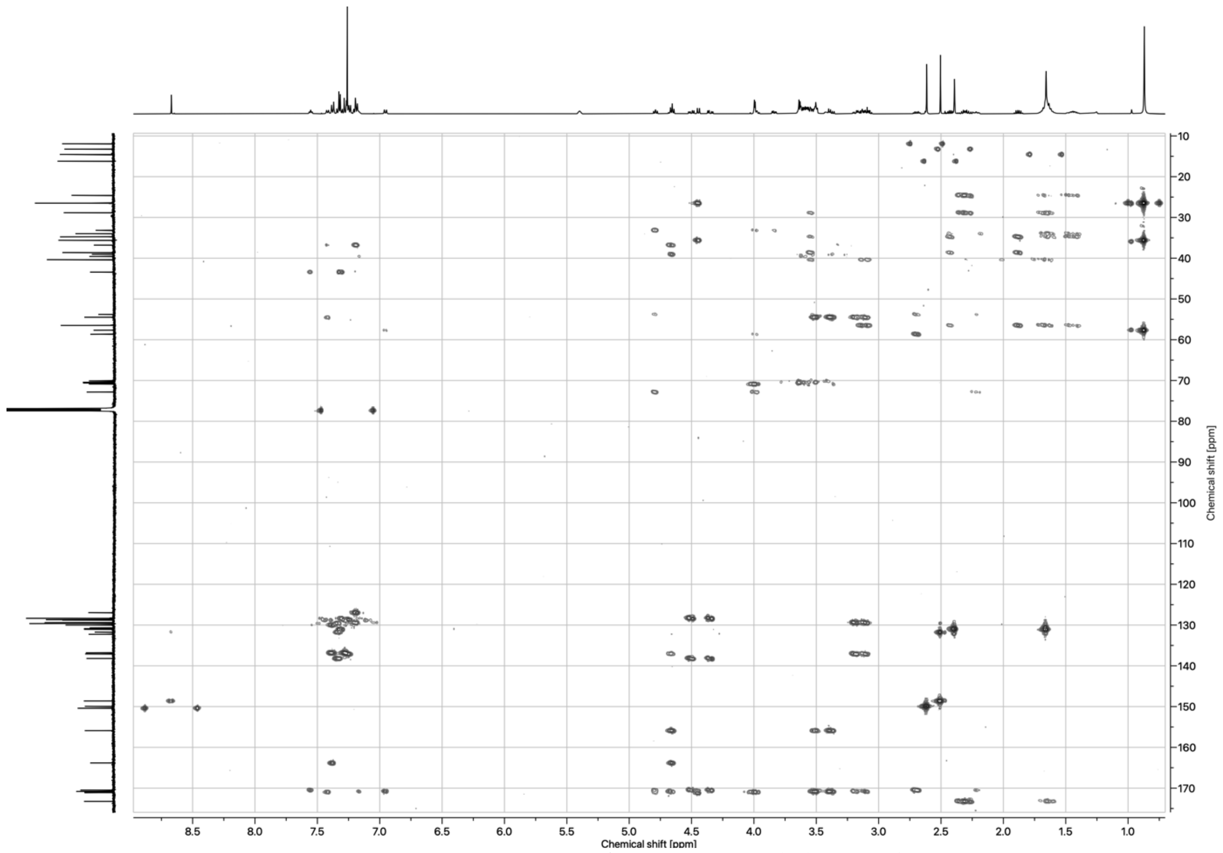

*Figure 83: HMBC spectrum of MZ3-LipA ester (**23**) in $CDCl_3$*

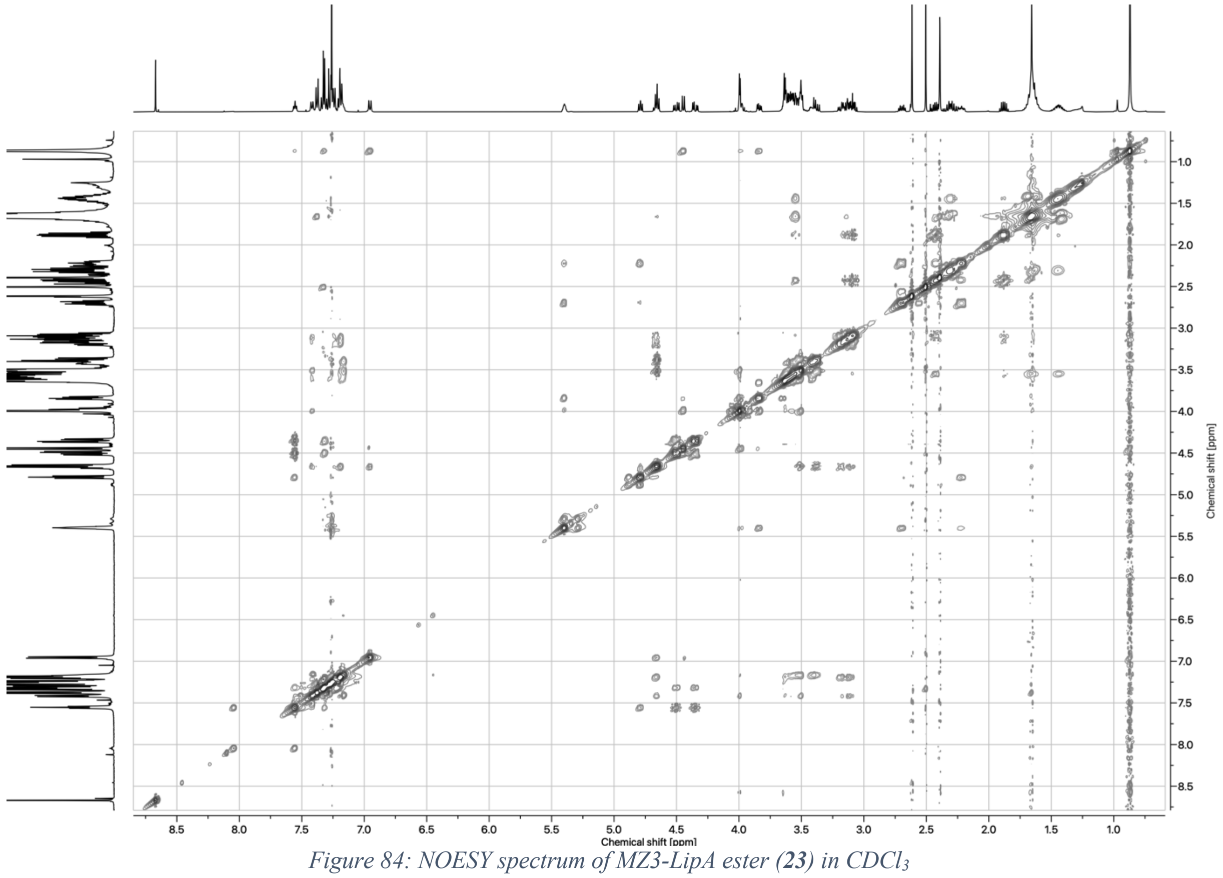

*Figure 84: NOESY spectrum of MZ3-LipA ester (**23**) in $CDCl_3$*

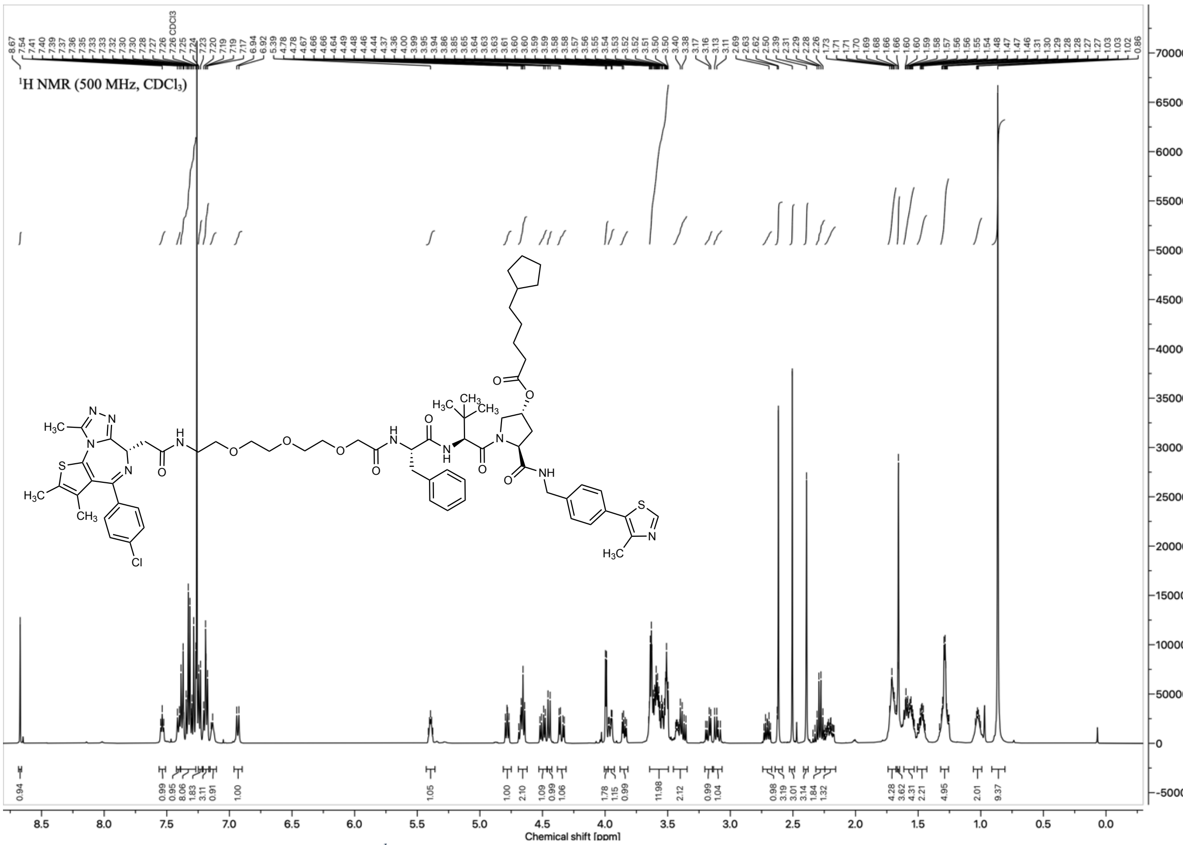

*Figure 85: $^{1}$H NMR spectrum of MZ3-All-C-LipA ester (**24**) in $CDCl_3$*

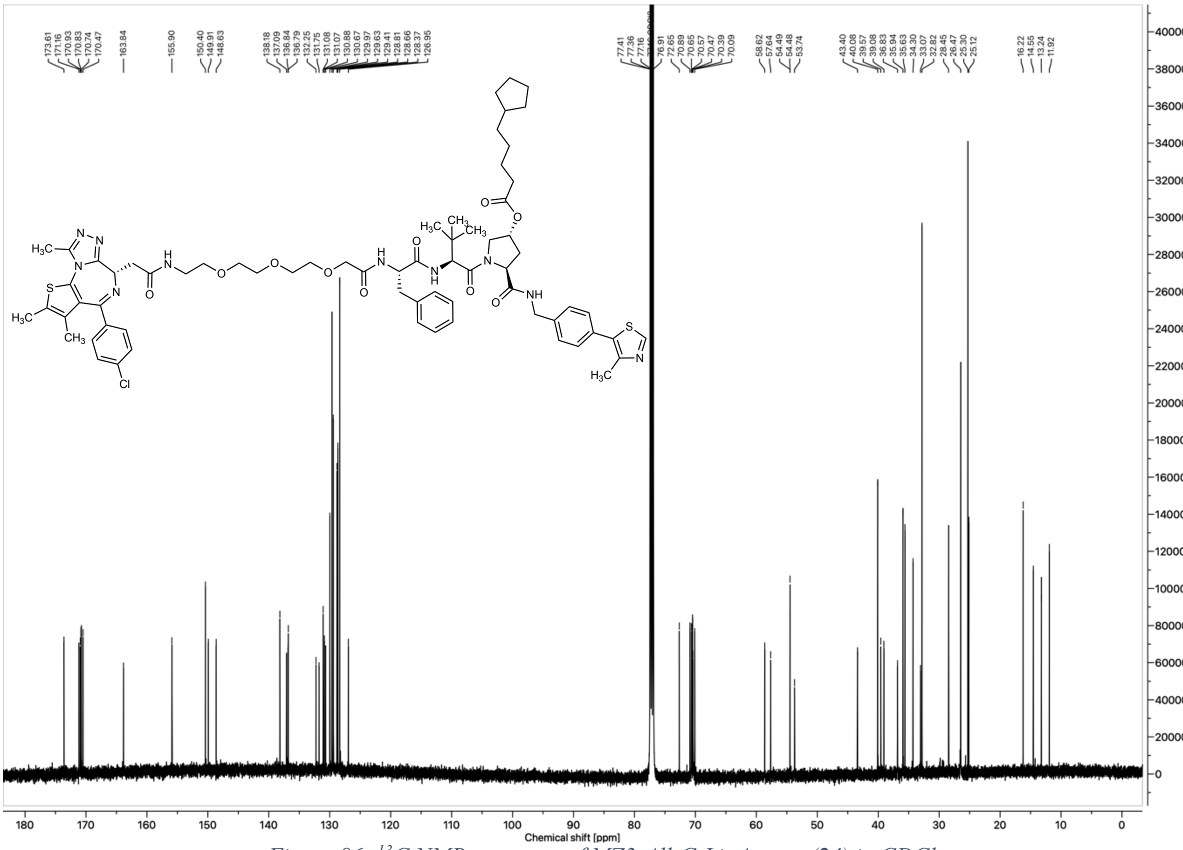

*Figure 86: $^{13}$C NMR spectrum of MZ3-All-C-LipA ester (**24**) in $CDCl_3$*

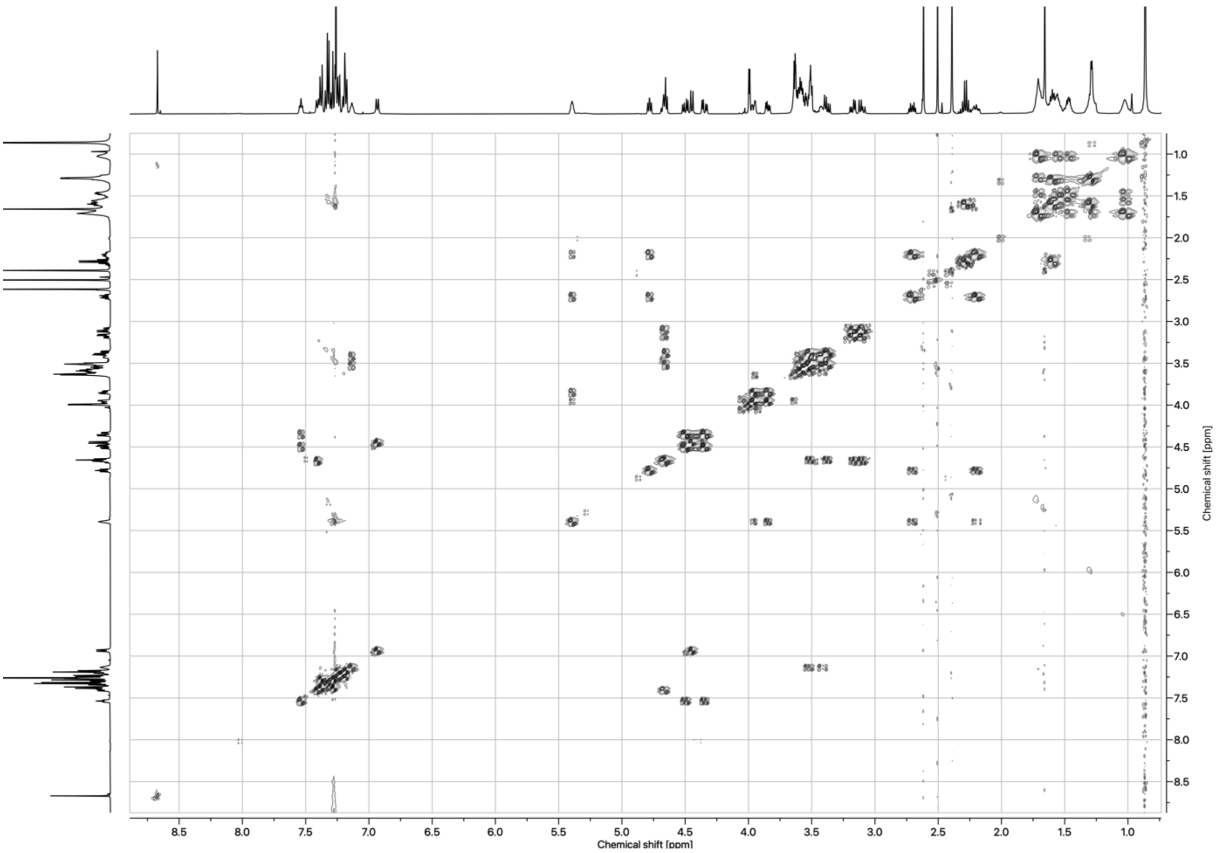

*Figure 87: COSY spectrum of MZ3-All-C-LipA ester (**24**) in $CDCl_3$*

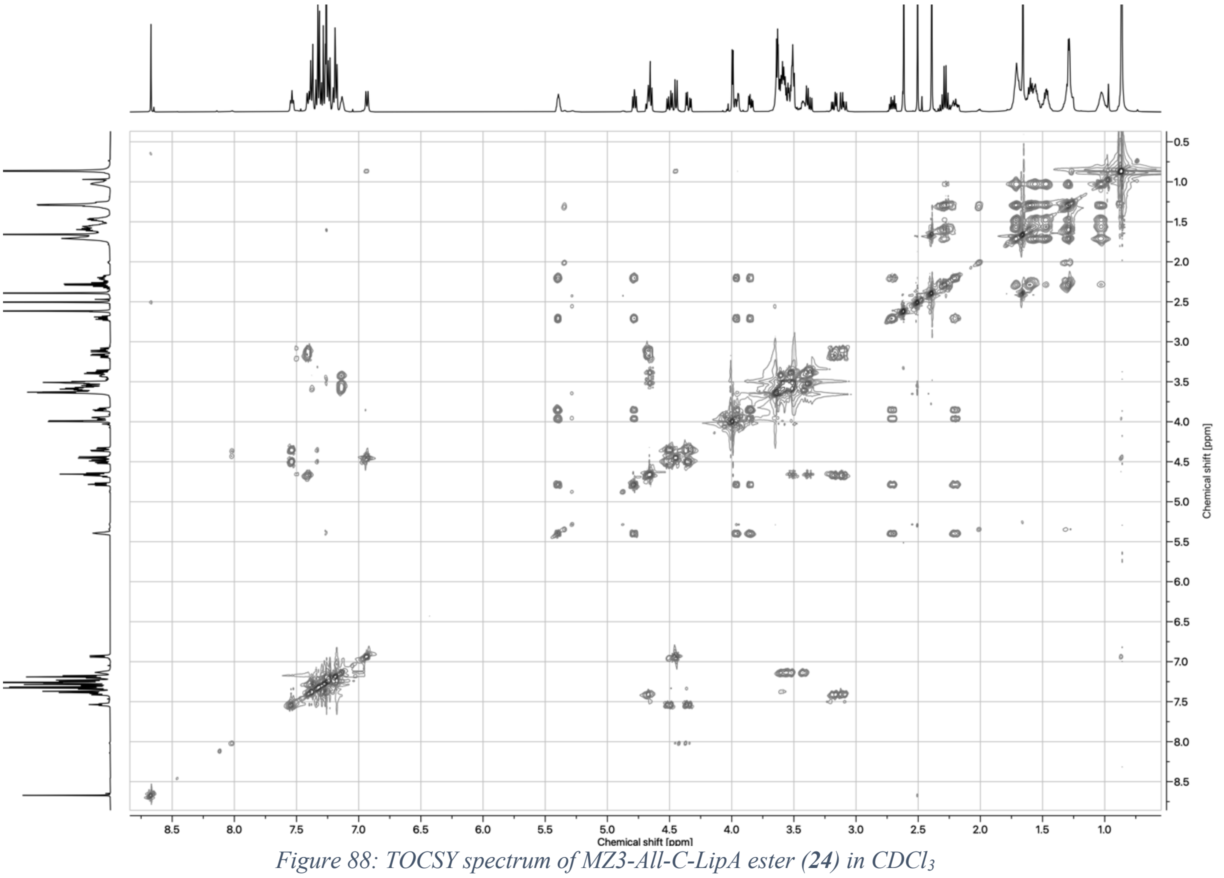

*Figure 88: TOCSY spectrum of MZ3-All-C-LipA ester (**24**) in $CDCl_3$*

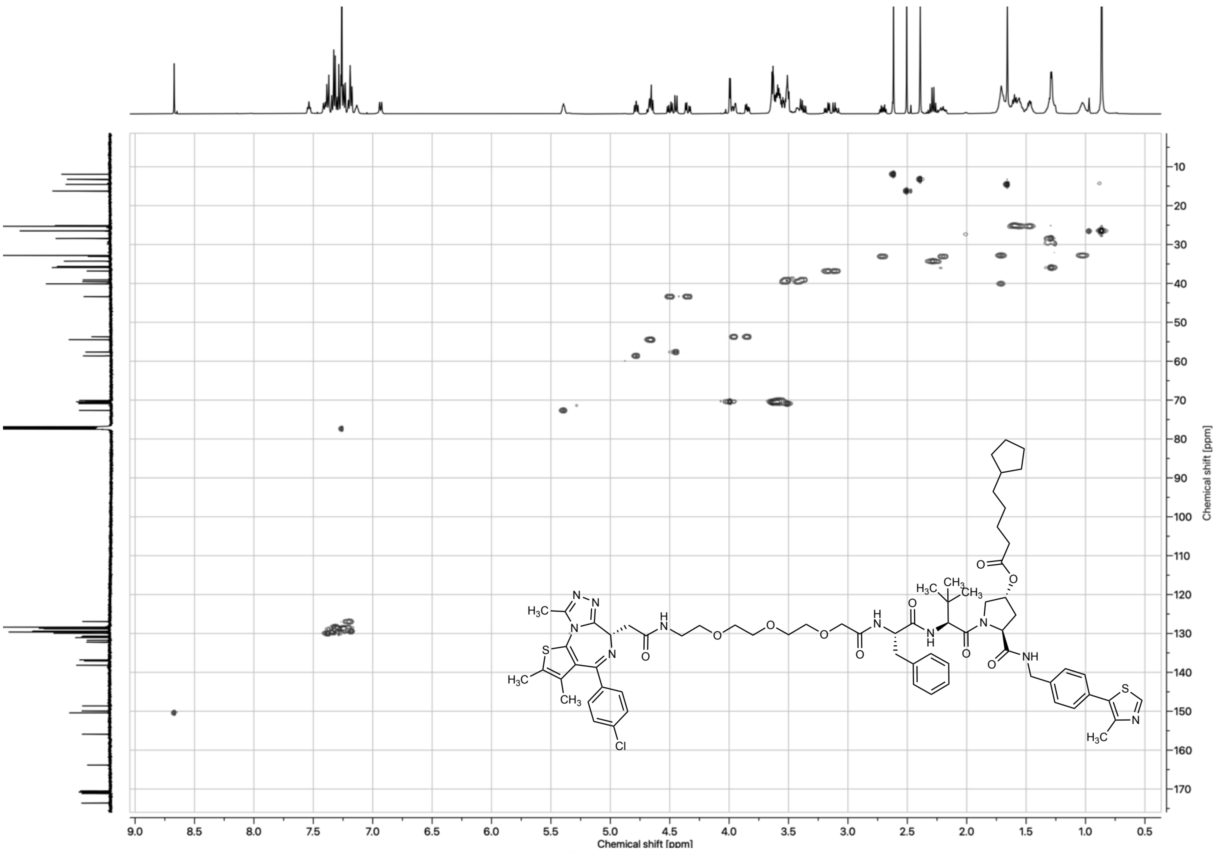

*Figure 89: HSQC spectrum of MZ3-All-C-LipA ester (**24**) in $CDCl_3$*

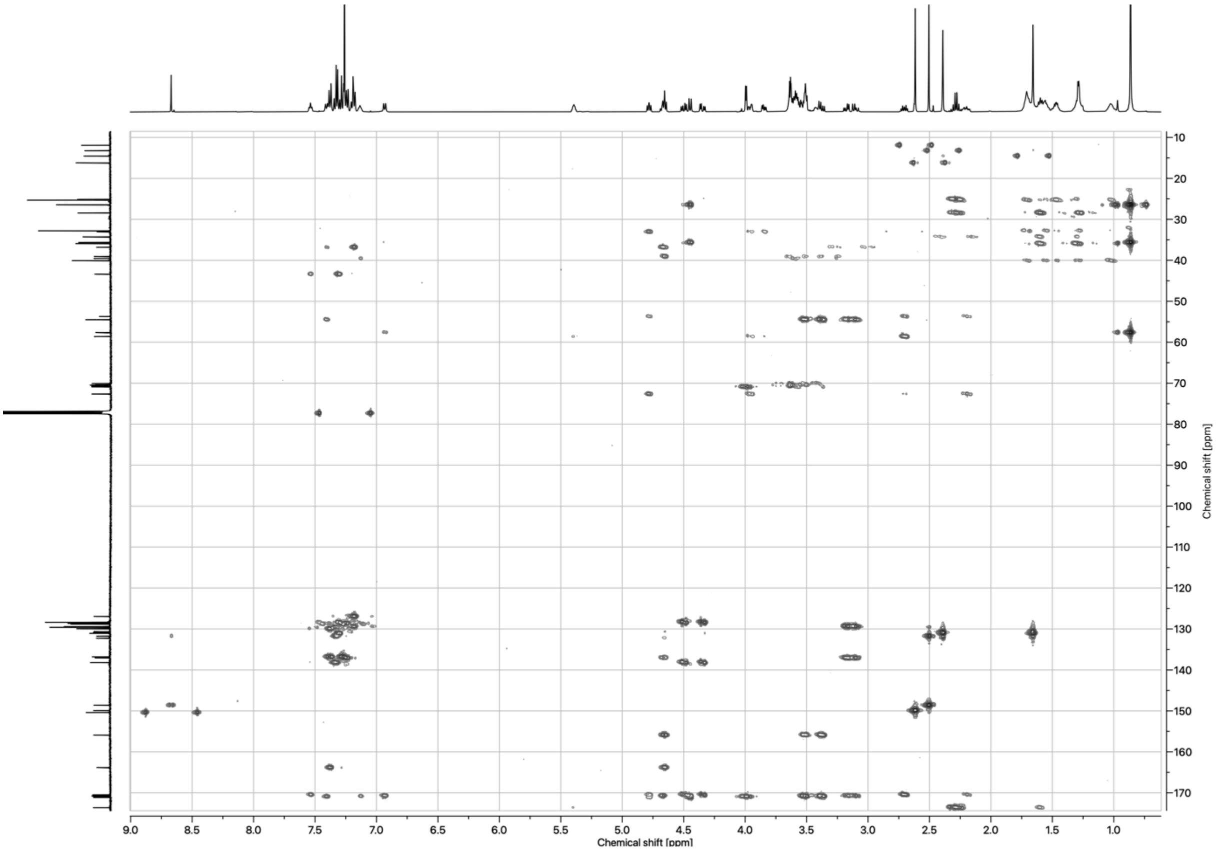

*Figure 90: HMBC spectrum of MZ3-All-C-LipA ester (**24**) in $CDCl_3$*

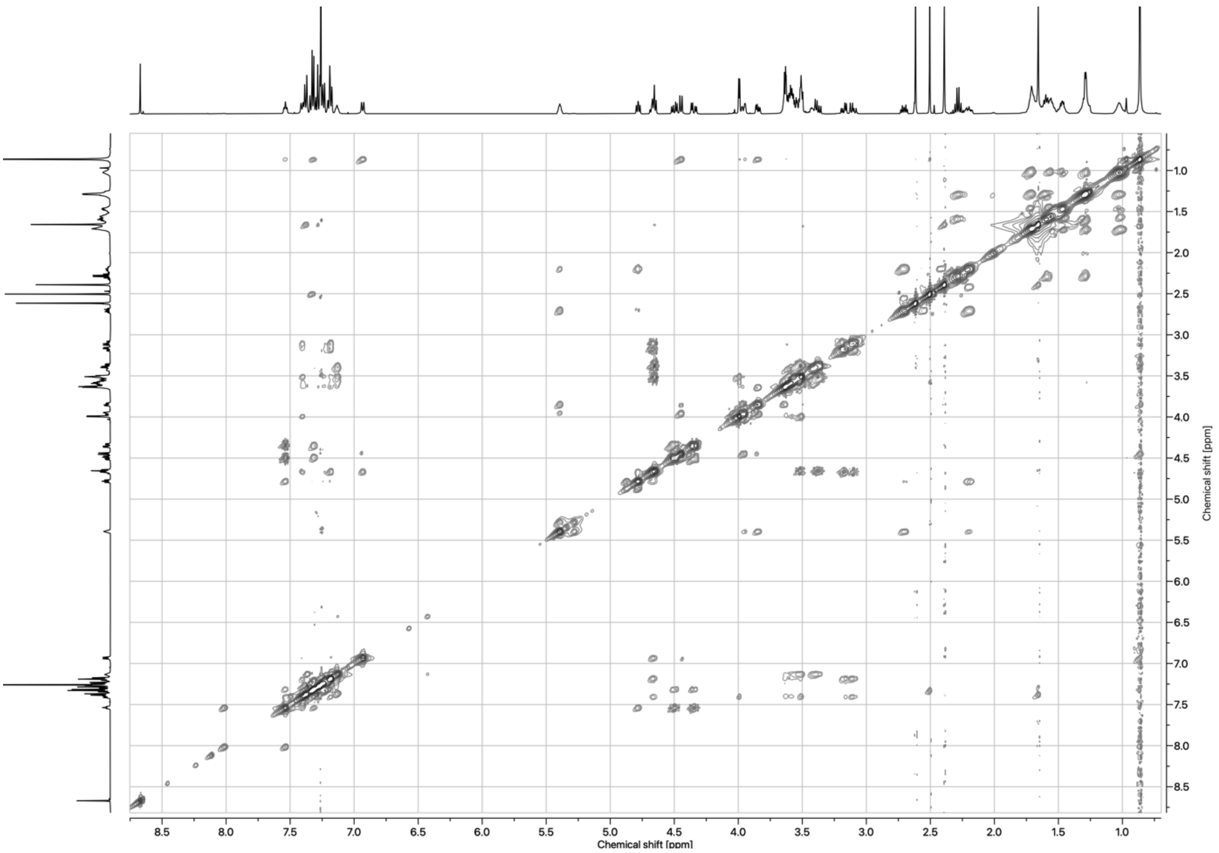

*Figure 91: NOESY spectrum of MZ3-All-C-LipA ester (**24**) in $CDCl_3$*

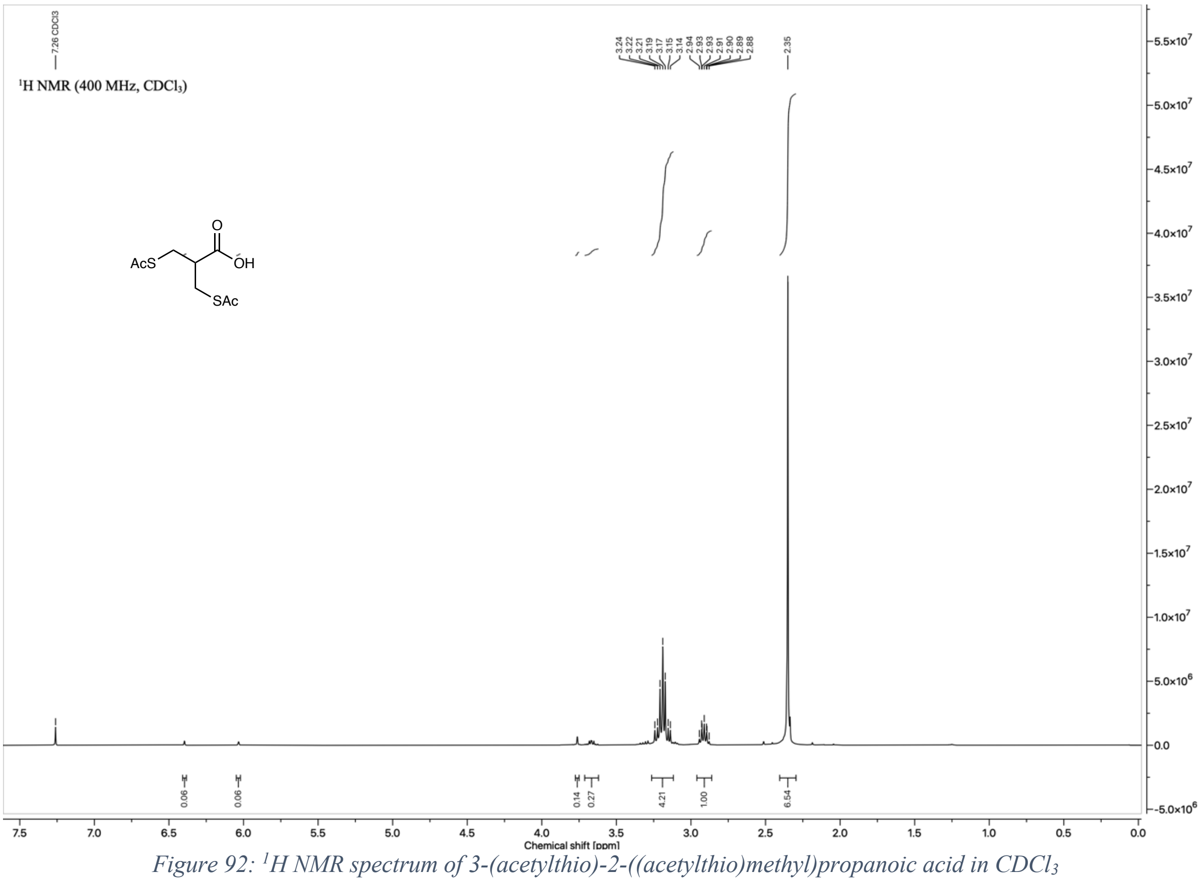

*Figure 92: $^1H$ NMR spectrum of 3-(acetylthio)-2-((acetylthio)methyl)propanoic acid in $CDCl_3$*

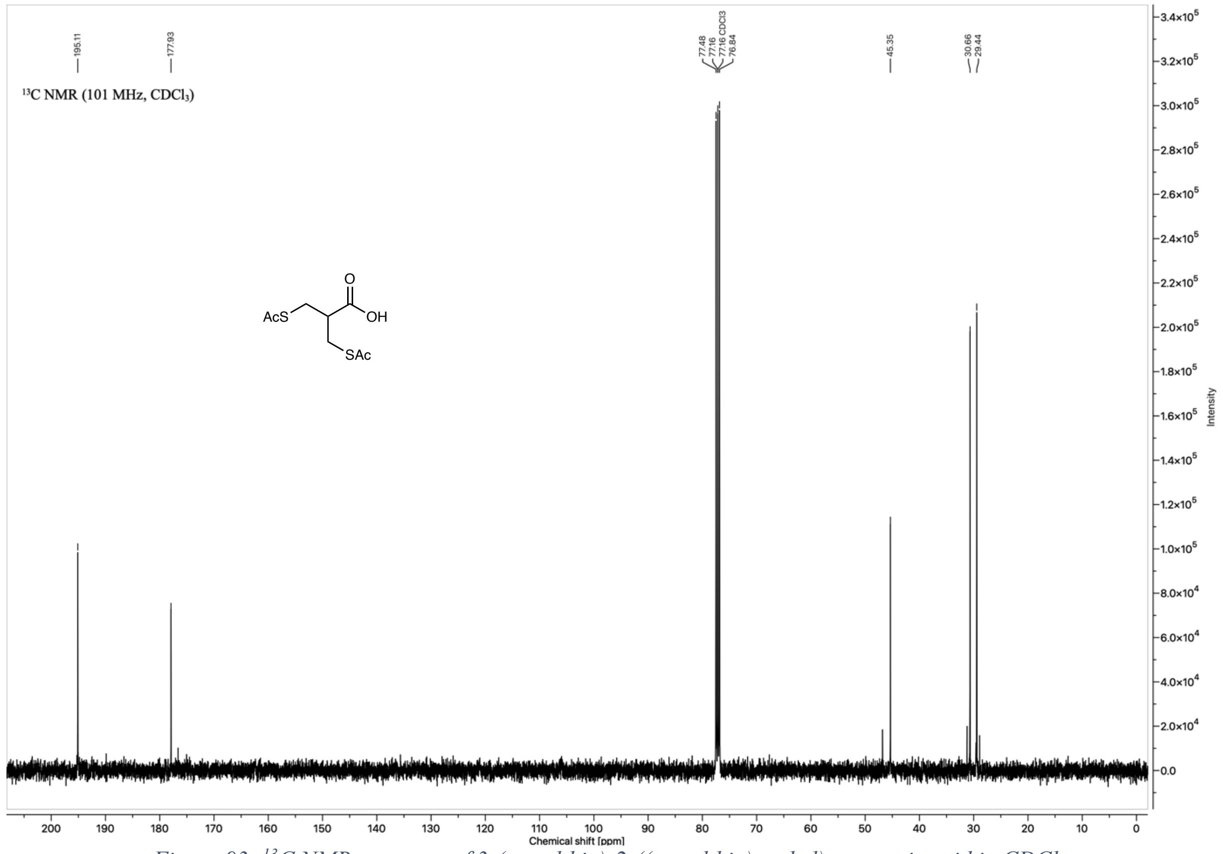

Figure 93: $^{13}C$ NMR spectrum of 3-(acetylthio)-2-((acetylthio)methyl)propanoic acid in $CDCl_3$

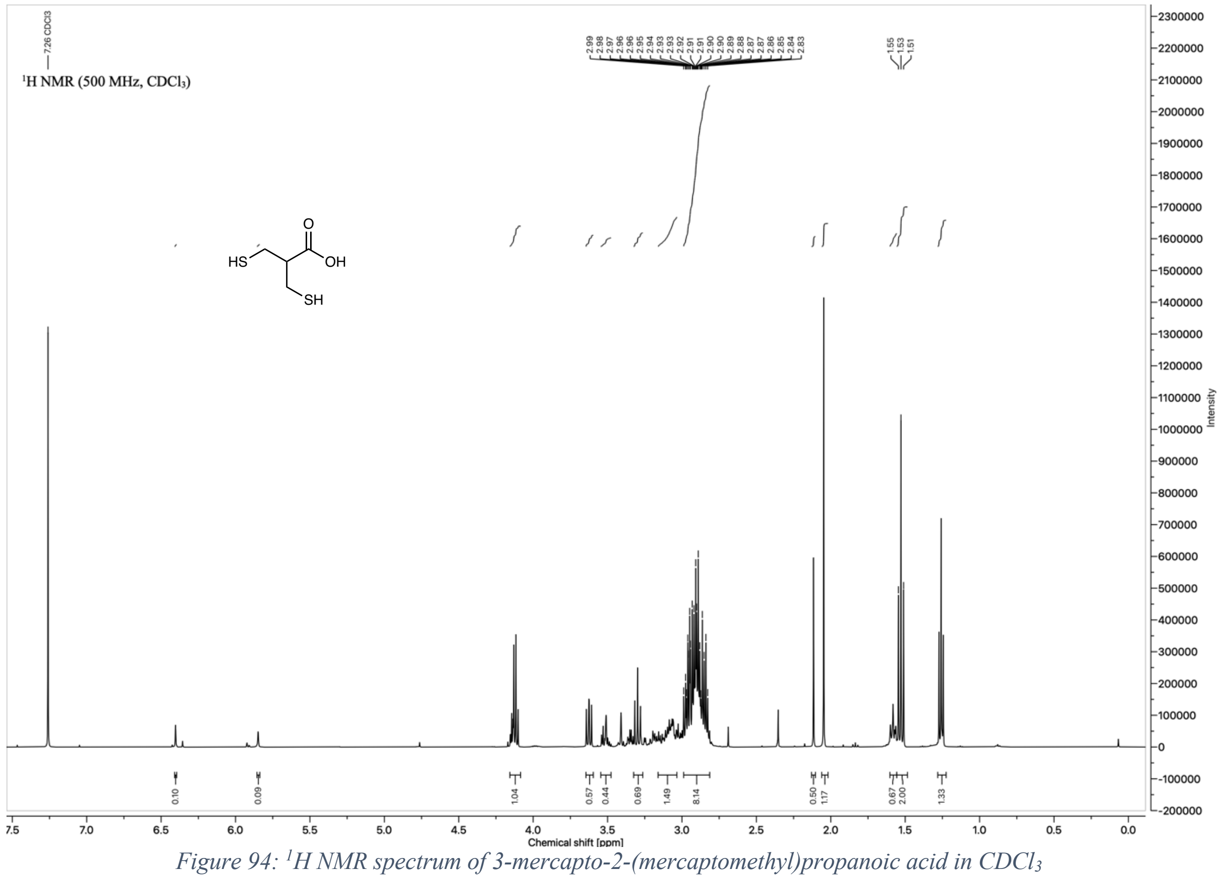

Figure 94: $^{1}H$ NMR spectrum of 3-mercapto-2-(mercaptomethyl)propanoic acid in $CDCl_3$

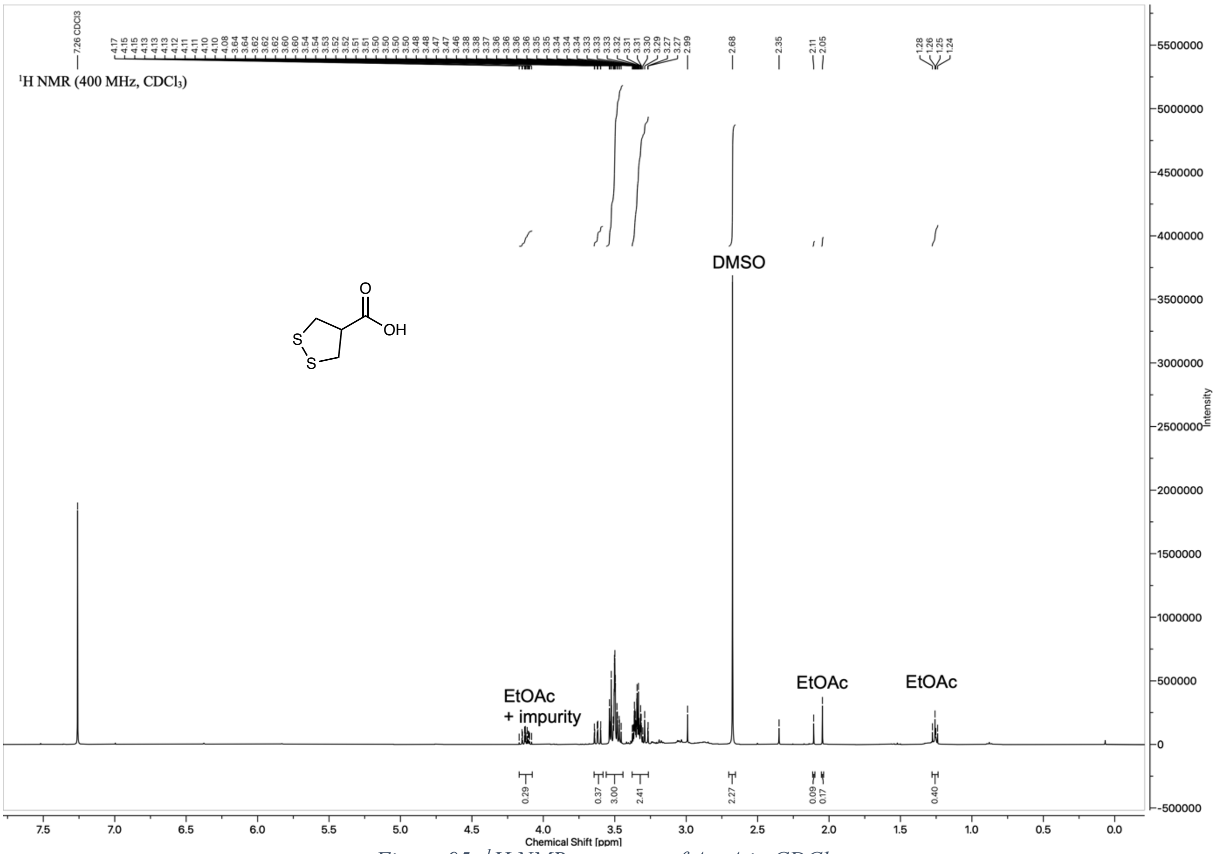

*Figure 95: $^{1}H$ NMR spectrum of AspA in $CDCl_3$*

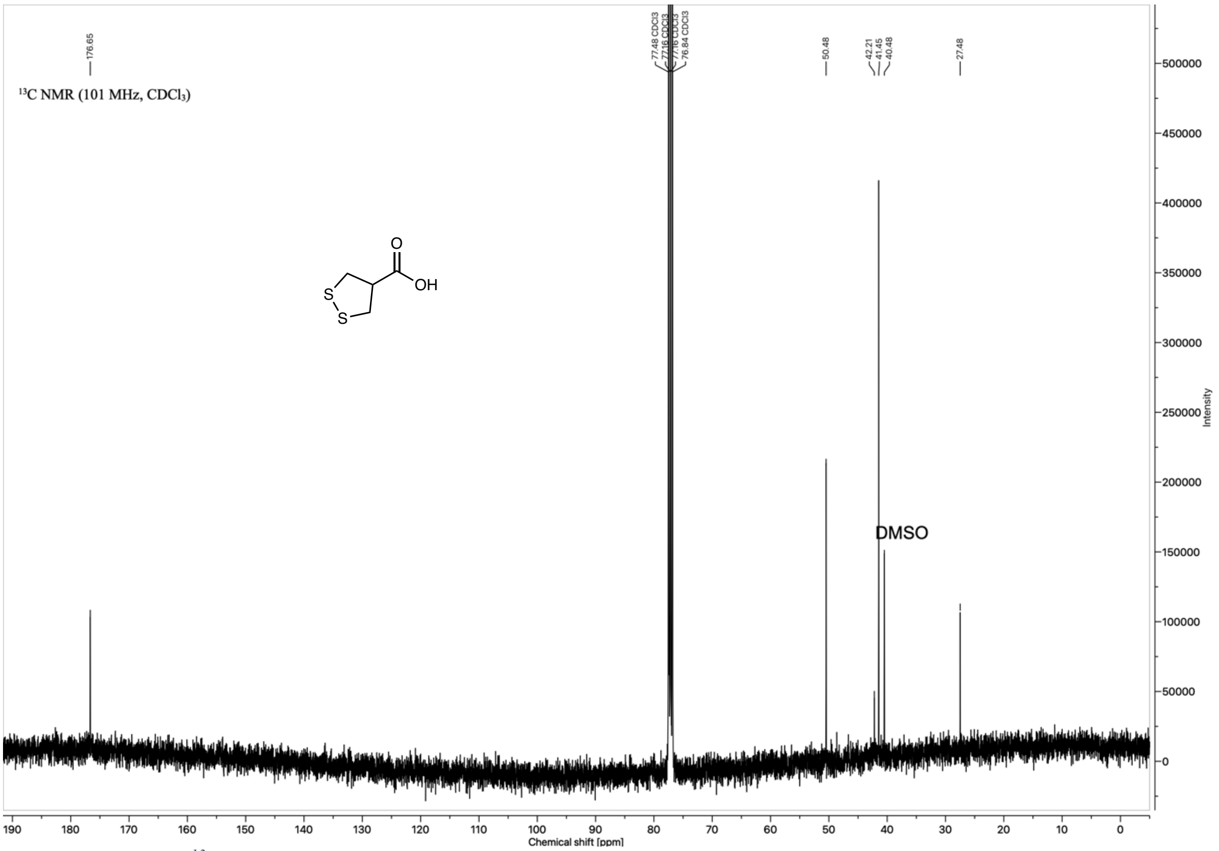

*Figure 96: $^{13}C$ NMR spectrum of AspA in $CDCl_3$. Signals at 27.48 ppm and 42.21 ppm: unidentified impurity.*

#### 3.2. UHPLC-MS/UV spectra

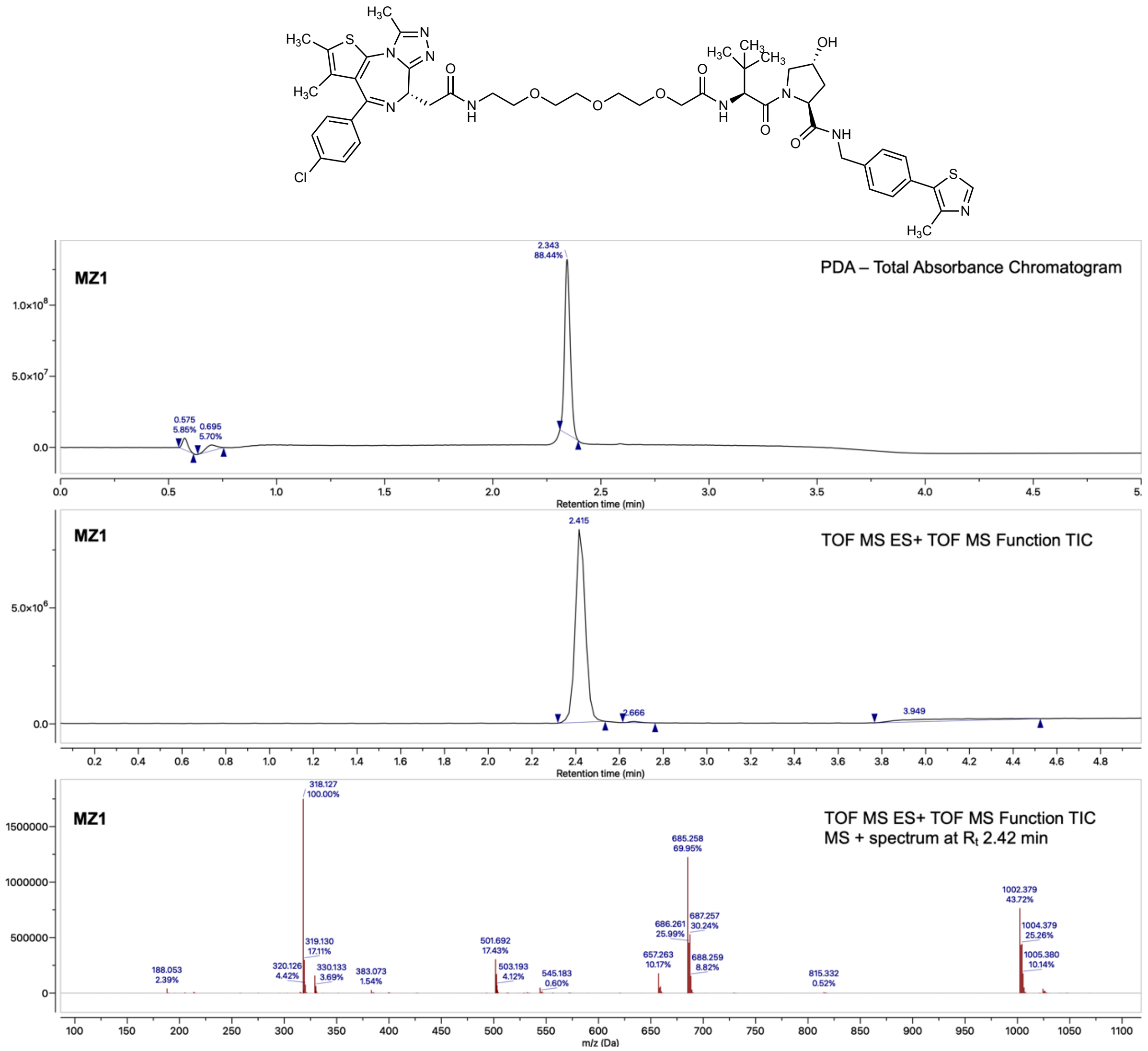

*Figure 97: UHPLC-MS/UV chromatograms of analysis of MZ1 (**1**)*

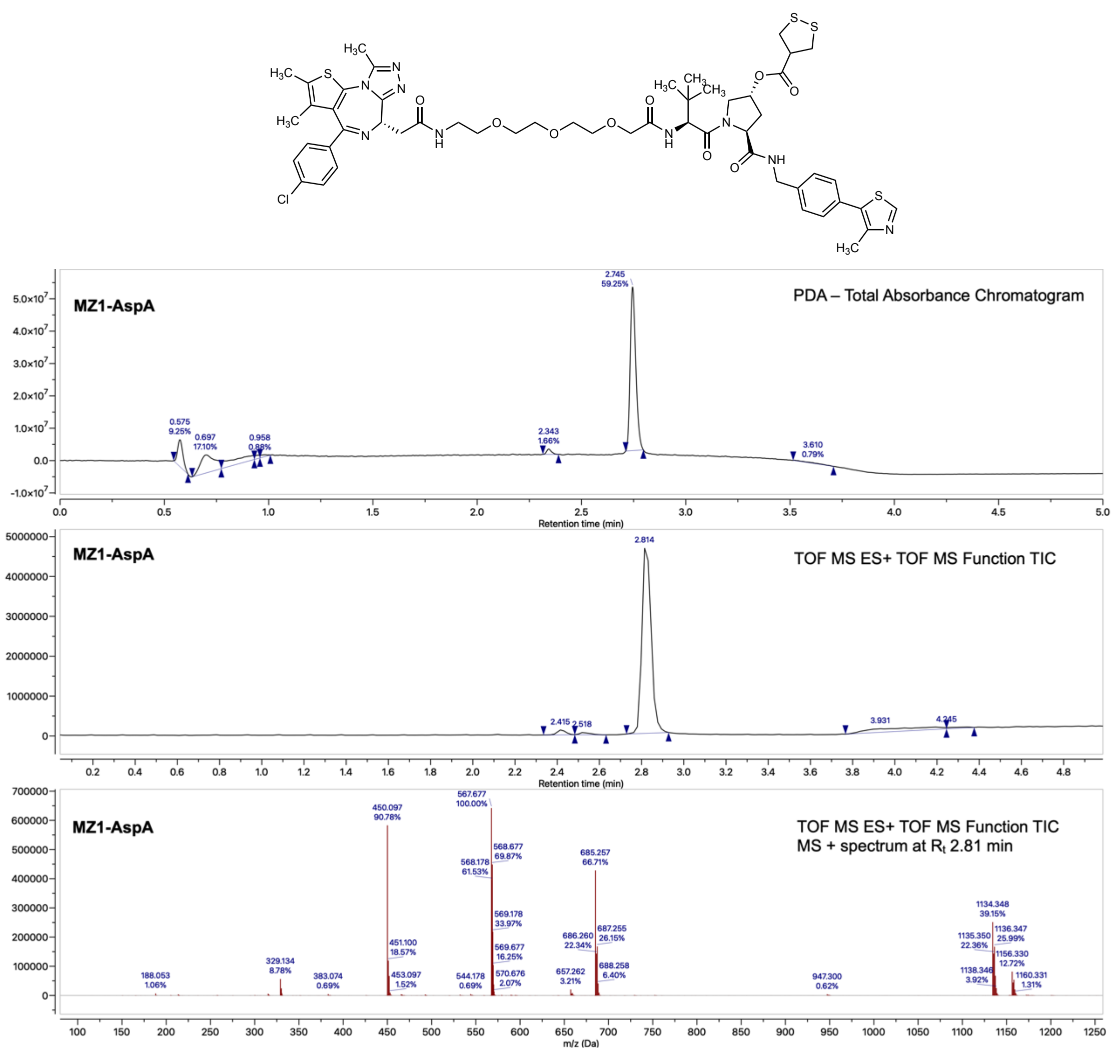

*Figure 98: UHPLC-MS/UV chromatograms of analysis of MZ1-AspA ester (**13**)*

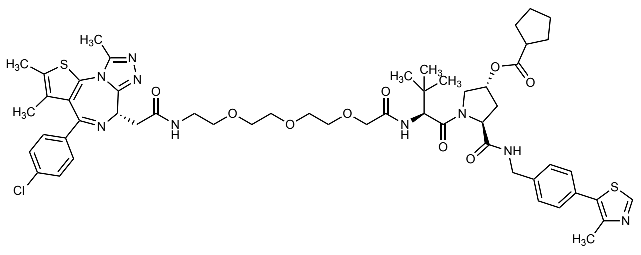

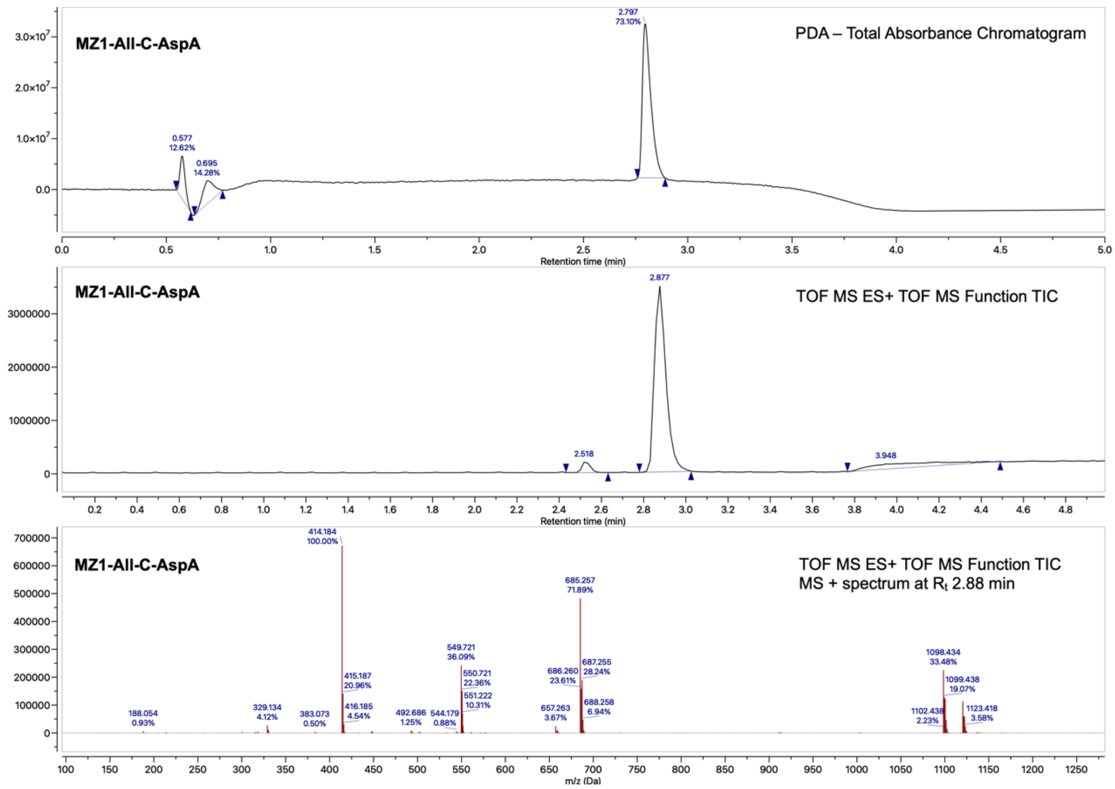

*Figure 99: UHPLC-MS/UV chromatograms of analysis of MZ1-All-C-AspA ester (**14**)*

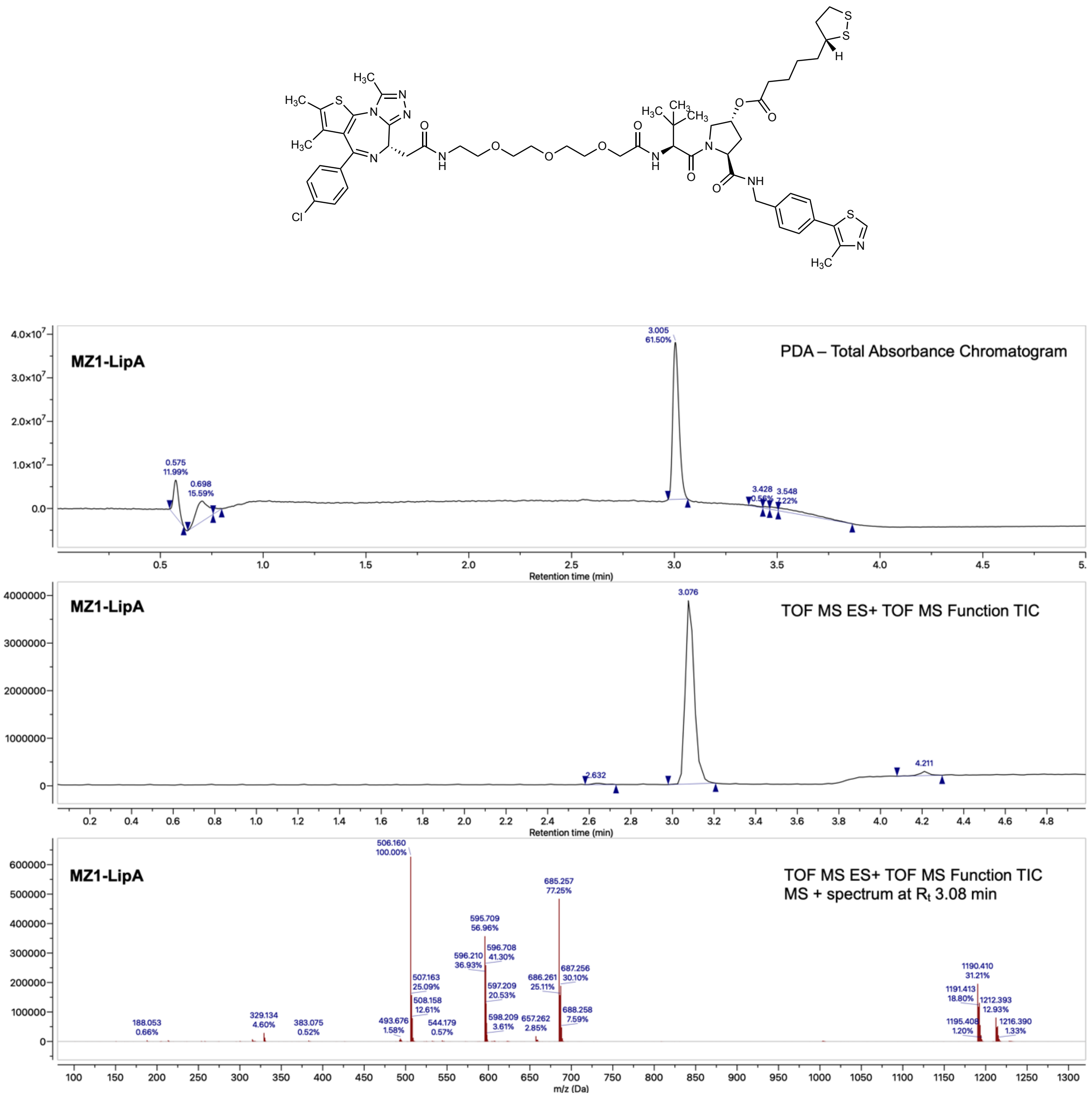

*Figure 100: UHPLC-MS/UV chromatograms of analysis of MZ1-LipA ester (**15**)*

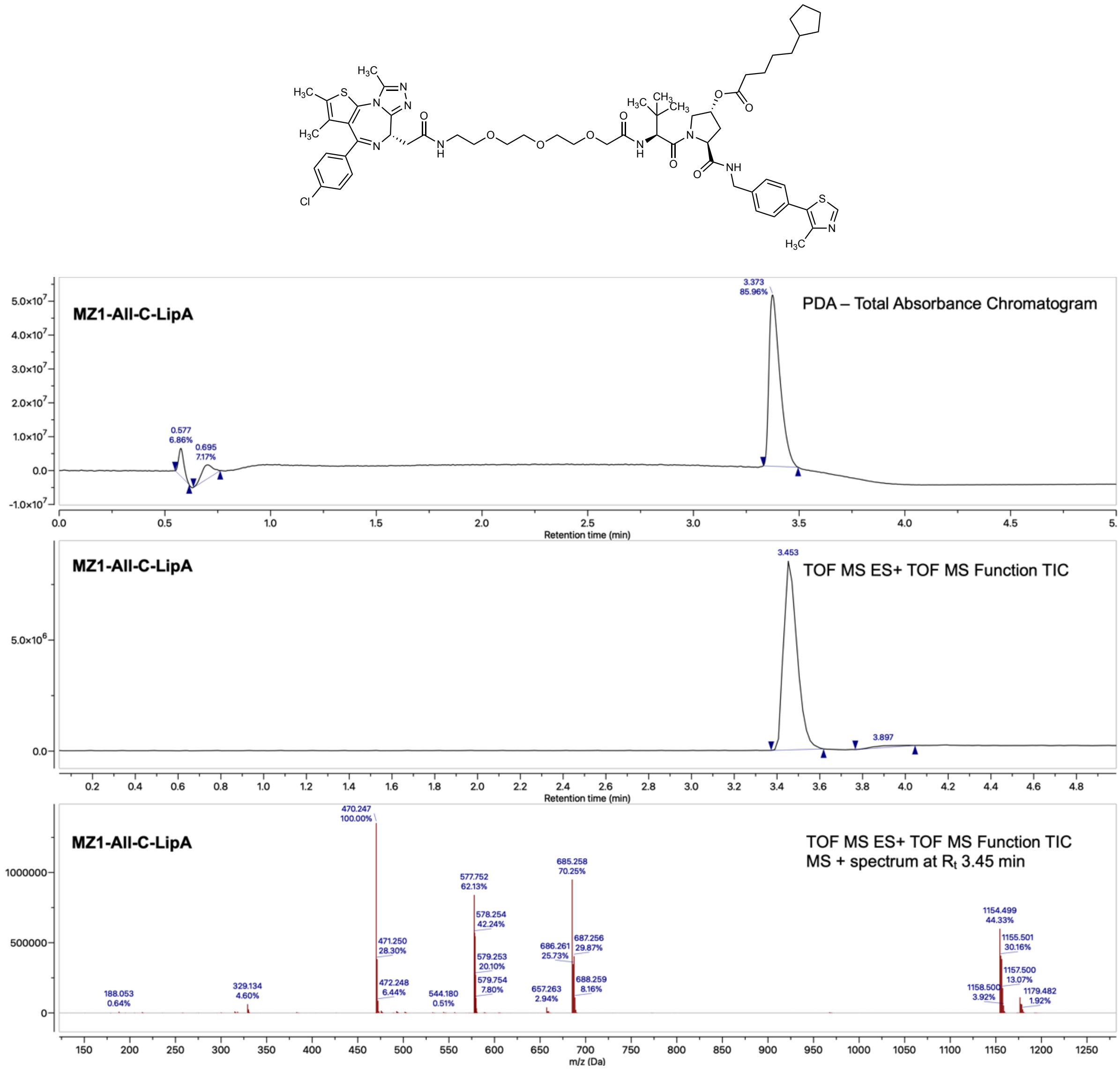

*Figure 101: UHPLC-MS/UV chromatograms of analysis of MZ1-All-C-LipA ester (**16**)*

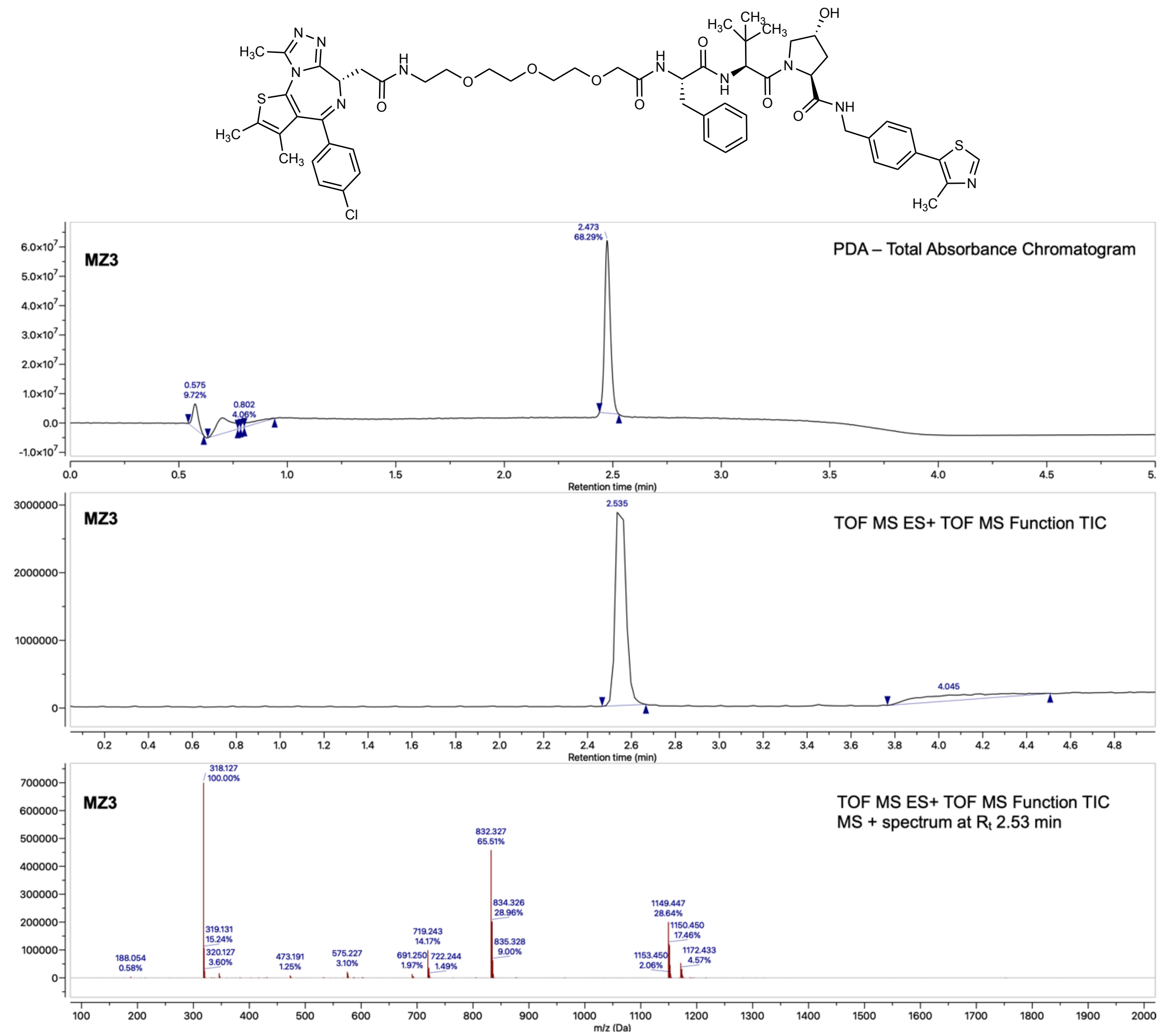

*Figure 102: UHPLC-MS/UV chromatograms of analysis of MZ3 (**2**)*

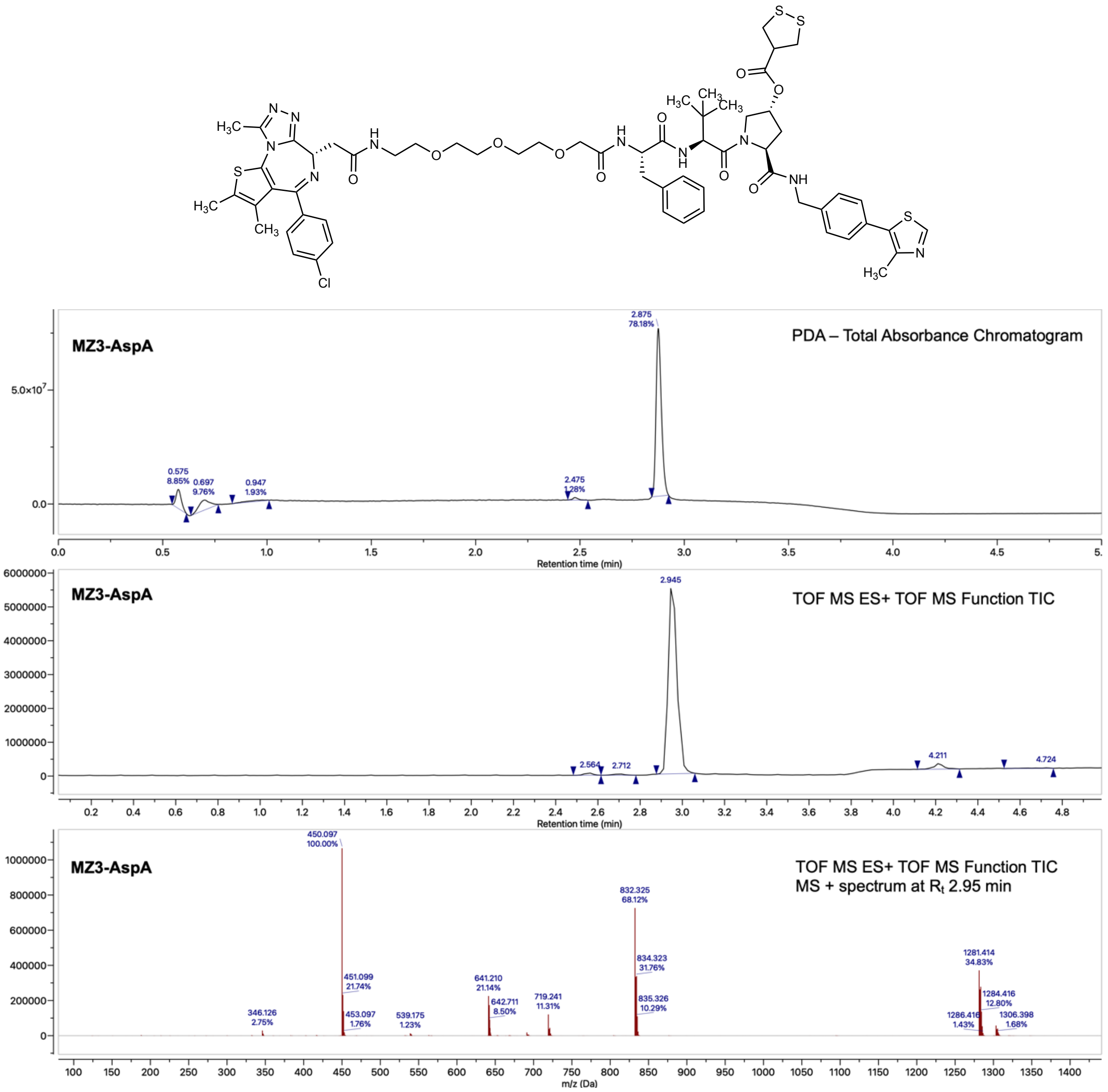

*Figure 103: UHPLC-MS/UV chromatograms of analysis of MZ3-AspA (**21**)*

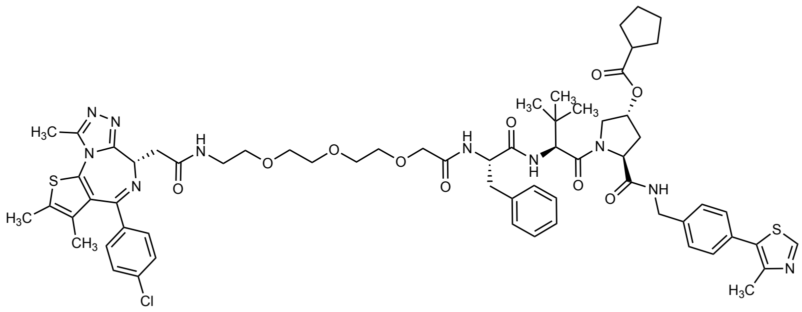

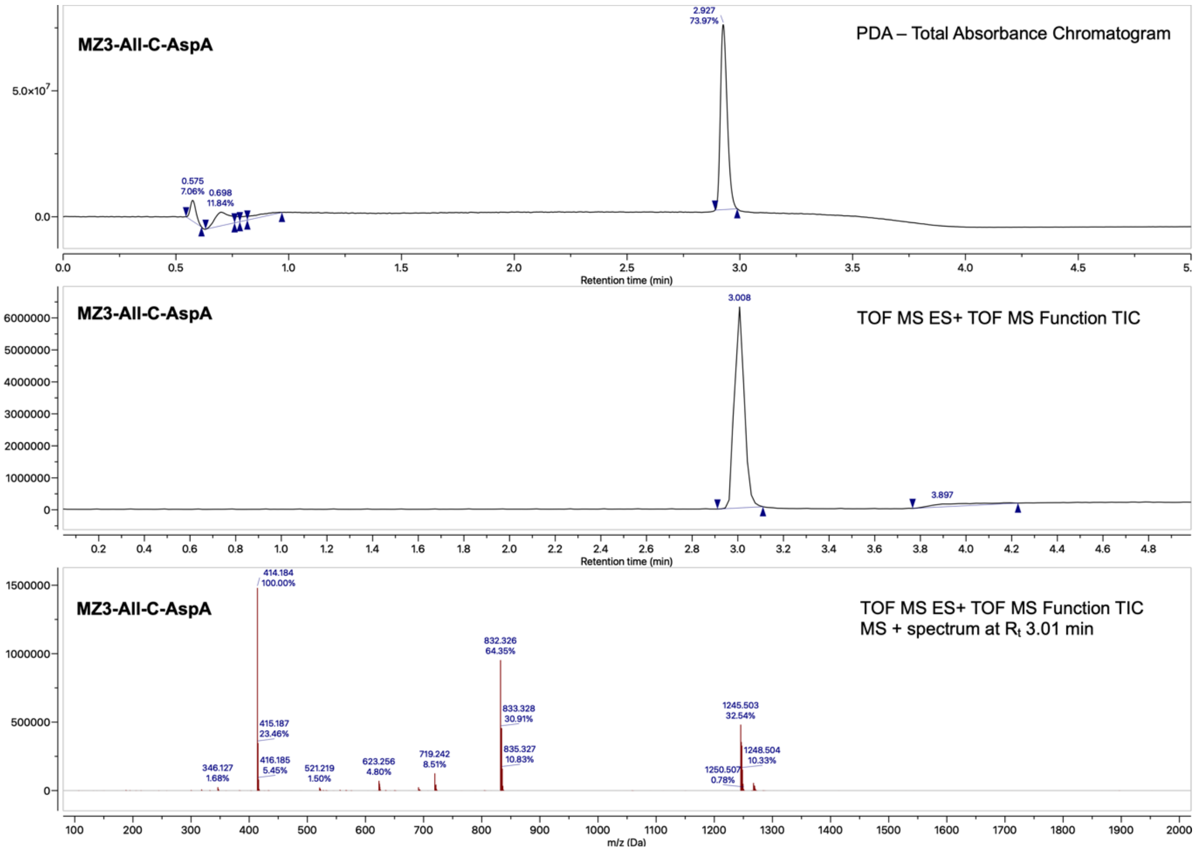

*Figure 104: UHPLC-MS/UV chromatograms of analysis of MZ3-All-C-AspA (**22**)*

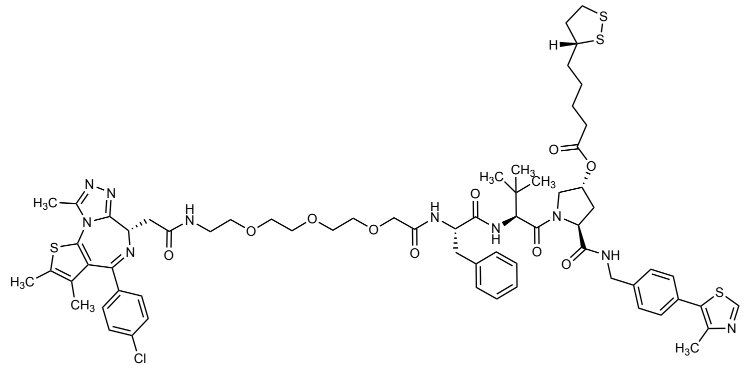

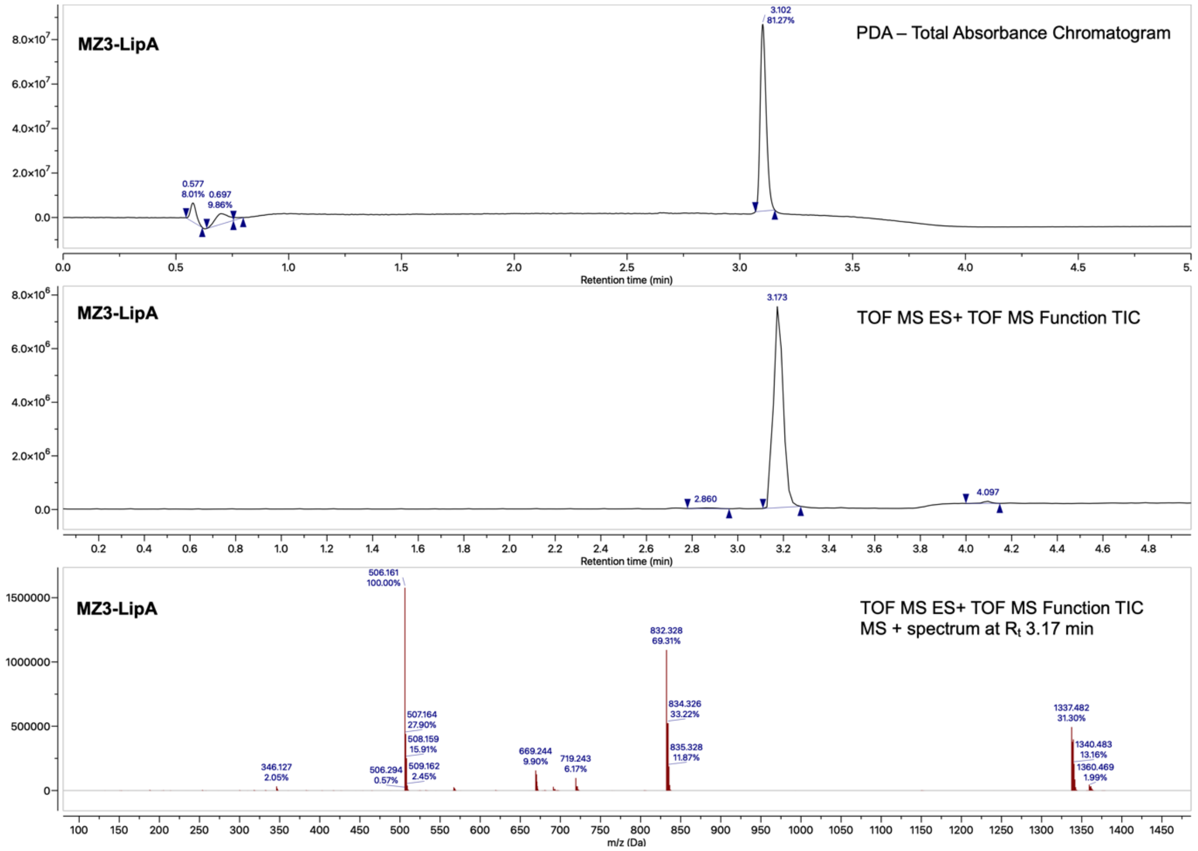

*Figure 105: UHPLC-MS/UV chromatograms of analysis of MZ3-LipA (**23**)*

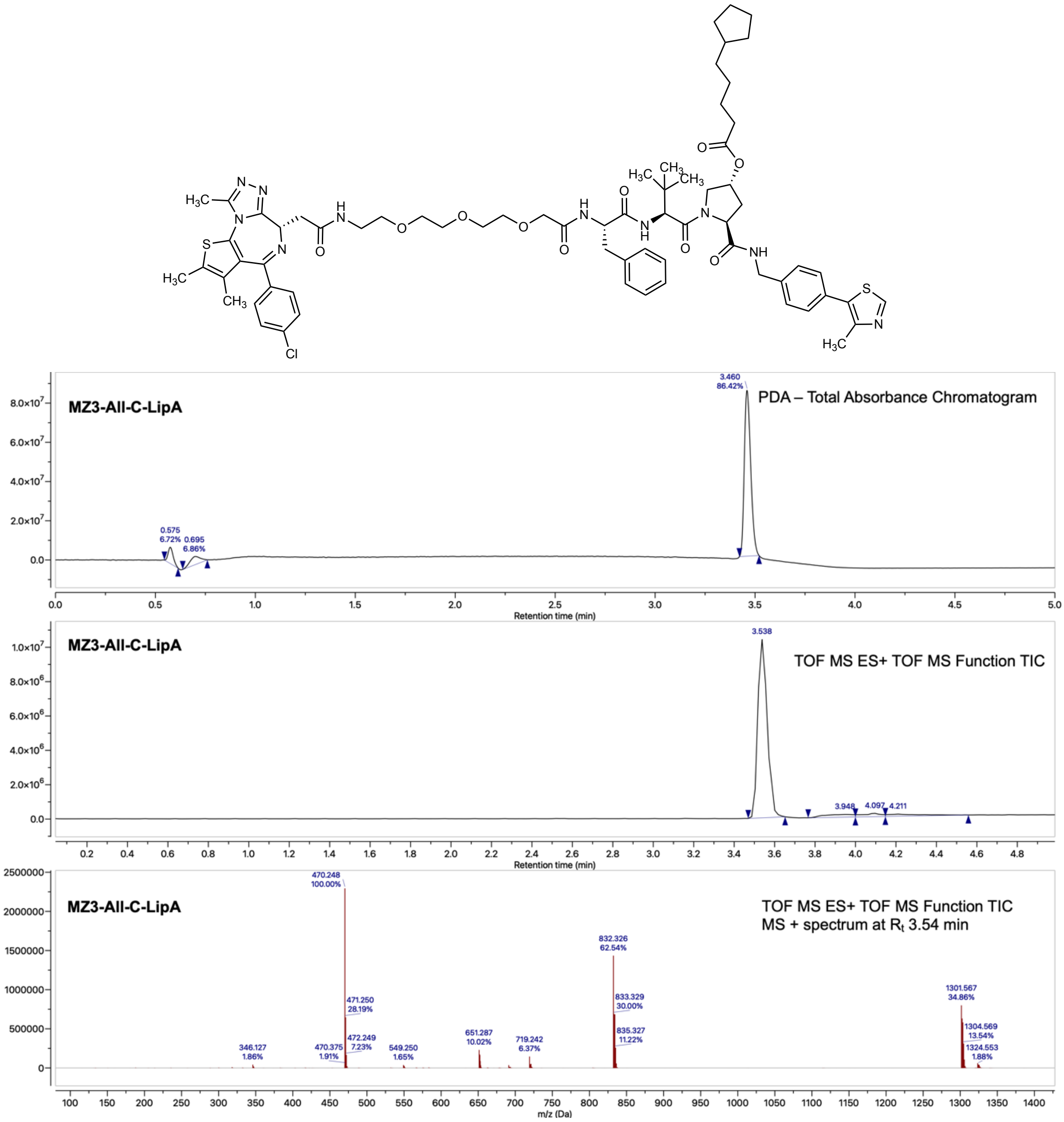

*Figure 106: UHPLC-MS/UV chromatograms of analysis of MZ3-All-C-LipA (**24**)*
